# Idiosyncrasies unveiled: examining the pace, patterns and predictors of biotic diversification in peninsular India

**DOI:** 10.1101/2023.11.15.567174

**Authors:** Pragyadeep Roy, Jahnavi Joshi

**Affiliations:** Academy of Scientific and Innovative Research (AcSIR), Ghaziabad, India; CSIR-Centre for Cellular and Molecular Biology, Uppal Road, Hyderabad, India

**Keywords:** diversification, biogeography, paleoclimate, animals and plants, birth-death models

## Abstract

The Peninsular Indian Plate (PIP), one of the oldest regions of diversification in tropical Asia, harbours highly diverse and endemic biota. However, our understanding of the diversification dynamics of its biota within a quantitative framework remains limited. To address this, we used time-calibrated molecular phylogenies and birth-death models to examine the tempo, mode, and drivers of diversification across 33 well-studied endemic lineages (∼770 species). Among PIP lineages, angiosperms diversified the fastest, invertebrates the slowest and younger lineages of Asian origins diversified more rapidly than the older relictual Gondwanan lineages. Evolutionary relatedness explained the disparities in diversification rates across taxonomic groups and biogeographic origins. A gradual accumulation of diversity was supported in 17 lineages, suggesting that the historical stability of their habitat was an important driver. Miocene intensification of monsoons and aridification and fluctuations in paleotemperature explained diversification patterns in the remaining 16 lineages. Our results highlight the role of regional biogeography, geoclimatic processes, and phylogenetic history in governing diversification dynamics in the tropics.

## Introduction

Diversification, a balance between speciation and extinction, governs biodiversity patterns. The hyperdiverse tropical biome has been central to diversification studies (Cardillo *et al*. 2005; Dagallier *et al*. 2020; Eiserhardt *et al*. 2017). Multiple hypotheses have been invoked to explain the high tropical diversity, including a long time for speciation because of being the oldest biome (Tietje *et al*. 2022), greater climatic stability (Dagallier *et al*. 2020), and habitat and climatic heterogeneity (Willig *et al*. 2003). A recent study from the Neotropics showed that the gradual expansion of diversity was the most common diversification scenario, irrespective of biogeographic subregions, and that diversification rates differed substantially across clades (Meseguer *et al*. 2022). On the other hand, in Australia, aridification and biogeographic barriers played key roles in driving diversification (Pepper & Keogh 2021). In Madagascar, a continental island, wet-forest-adapted species exhibited accelerated diversification (Crottini *et al*. 2012). In addition, tropical islands have highly endemic adaptive radiations primarily driven by niche dynamics and biogeographic processes (Losos & Ricklefs 2009; Wallace 1887). These studies across tropical regions suggest that patterns of diversification depend on regional biogeographic and geoclimatic processes, geographic settings such as being an island or a continent, and vary across the Tree of Life. Therefore, identifying and characterising the processes responsible for differential species diversity among distinct geographical regions and clades is essential for a nuanced understanding of the factors contributing to tropical diversity.

In this regard, the Peninsular India Plate (PIP) remains relatively unexplored despite being one of the oldest regions for diversification in tropical Asia (Karanth 2021; Mani 1974). The PIP was a part of the Gondwanaland supercontinent ∼200 Mya (Ali & Aitchison 2008; Chatterjee *et al*. 2013). Around 120 Mya, it separated from the supercontinent along with Madagascar and began drifting northeastward (Chatterjee *et al*. 2013). On its northward journey, the PIP remained a continental island for ∼50 My until its collision with Asia. It harboured a predominantly Gondwanan biota, as evidenced by fossils and molecular phylogenetic studies of taxa with vicariant origins, many of which (scolopendrid centipedes, *Gerrhopilus* blindsnakes, etc.) continue to persist (Bonaparte 1999; Joshi & Karanth 2011; Prasad *et al*. 2009; Sidharthan & Karanth 2021). Mass extinctions marked this journey due to immense volcanic activity (Fig. 1) at the K-Pg boundary (70–60 Mya), known as the Deccan volcanism (Keller *et al*. 2009; Prasad *et al*. 2009). Subsequently, as the PIP approached the equator, it experienced an expansion of dense perhumid forests, followed by its collision with Asia (Morley 2018) at around ∼35 Mya. Multiple transient land connections with parts of Southeast Asia between 55–35 Mya (Ali & Aitchison 2008) and the prevalence of wet rainforests in the PIP and Southeast Asia facilitated biotic exchanges between these regions, particularly among taxa with similar ecological niches (Grismer *et al*. 2016; Klaus *et al*. 2016; Li *et al*. 2020). Additionally, dispersal events from Africa to the PIP occurred during the Late Maastrichtian (72–66 Mya) and Paleocene (66–56 Mya), involving taxa such as the Dipterocarpaceae plants, *Grypotyphlops*blindsnakes, and natatanuran frogs (Bansal *et al*. 2022; Sidharthan & Karanth 2021; Yuan *et al*. 2019). As a result, the PIP biota has Gondwanan, Asian, and African biogeographic affinities (Karanth 2021; Manivannan *et al*. 2024). The India-Eurasia collision also triggered the Himalayan-Tibetan Orogeny (19–15 Mya) (Chatterjee *et al*. 2013; Ding *et al*. 2017), which induced seasonality in the PIP (Morley 2018) and also created dispersal barriers between the PIP and Southeast Asia (Karanth 2015). The wet evergreen rainforests of PIP further started contracting due to intensified aridification between 11–7 Mya and the expansion of C4 plants across the peninsula (9–2.5 Mya) (Morley 2018; Quade *et al*. 1989). Today, the wet forests are restricted to the western slopes of the Western Ghats, mountaintops in the Eastern Ghats, and fragmented patches within dry forests and savannas of the PIP (Karanth 2003; Mani 1974).

**Fig. 1.**
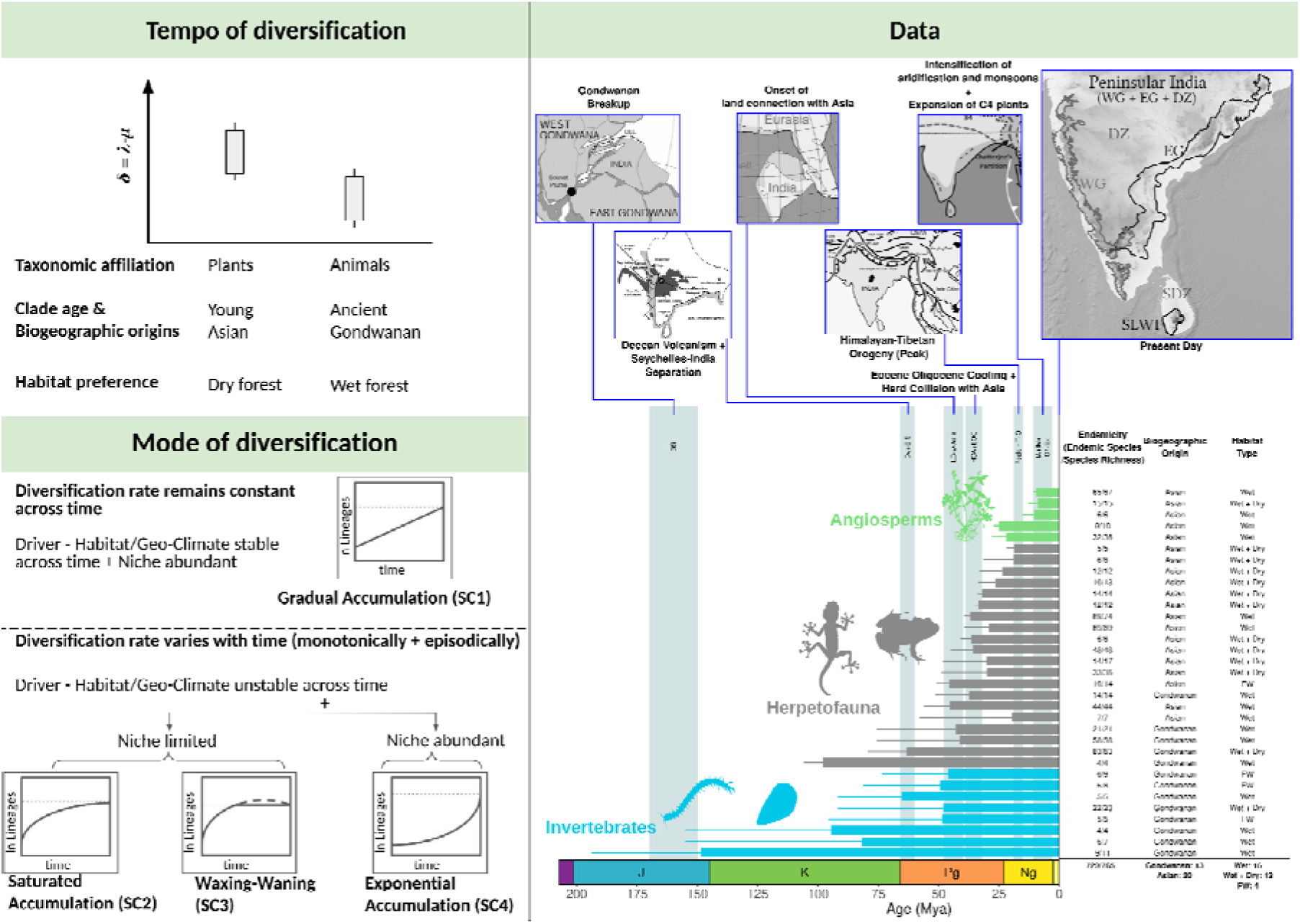
Predictions for the tempo and mode of diversification and the data used in the study. **(Left panel)** The mean diversification rate can be influenced by taxonomic affiliations, clade age, biogeographic origins and habitat preference as illustrated in the box plot (see Box 1 for details). The mode of diversification can follow multiple scenarios due to speciation and extinction rate dynamics as depicted (see Fig S1 for details). **(Right panel)** Our dataset of 33 endemic lineages from the PIP. They vary in clade age, number of species, endemicity, biogeography, and habitat affinities. Key geological events are highlighted by grey vertical bars. Maps used in the figure have been extracted and edited from multiple studies, and the map for the present day has been generated using QGIS(Ali & Aitchison 2008; Chatterjee *et al*. 2013; Morley 2018). WG: Western Ghats; EG: Eastern Ghats; DZ: Dry Zone; SLDZ: Sri Lanka Dry Zone; SLWF: Sri Lanka Wet Forests

Undoubtedly, the complex geo-climatic history of the PIP has played an important role in generating this complex landscape and, in turn, influenced the diversification of flora and fauna both within the PIP and across tropical Asia (Deepak & Karanth 2018; Joshi & Edgecombe 2019; Lajmi & Karanth 2020). Today, the biotic diversity in the PIP is highly endemic with distinct biogeographic origins (Gondwanan, Southeast Asian or Eurasian) (Joshi *et al*. 2020; Karanth 2021; Surveswaran *et al*. 2021). Also, many clades have undergone extensive *in situ* diversification within the PIP during the Cenozoic, establishing it as a distinct biogeographic subregion within tropical Asia (Karanth 2021). However, our understanding of the tempo, mode, and drivers of diversification of the PIP biota remains limited (Agarwal & Ramakrishnan 2017; Cyriac *et al*. 2024; Lajmi & Karanth 2020). Like many tropical areas, the PIP diversity and habitats are threatened by global change. Therefore, understanding diversification dynamics is also crucial for assessing the evolutionary past and predicting the future of the PIP’s highly diverse endemic biota.

Given this, we use a comparative framework to examine the relative roles of geoclimatic processes, biogeographic affinities, and species traits in shaping the extant diversity within the PIP. We use state-of-the-art birth-death models to examine the tempo and temporal mode of *in situ* diversification and their drivers for the endemic biota of PIP (Condamine *et al*. 2013; May *et al*. 2016; Morlon *et al*. 2010). We compiled data on 33 floral and faunal lineages (clades) with available species distribution, molecular phylogenetic relationships, and divergence time estimates (Fig. 1; Table S1). Most of these lineages are endemic to the PIP and have undergone extensive *in situ* diversification, making them ideal for examining diversification patterns and their ecological and evolutionary drivers in a comparative framework. These taxa are distributed across terrestrial (wet or dry forest) and freshwater habitats, with either Asian or Gondwanan biogeographic affinities (Fig. 1; Table S1; Box 1). Building on detailed phylogenetic and biogeographic studies, we infer diversification rates, patterns, and potential drivers of diversification within the PIP. Importantly, we propose diversification hypotheses with testable predictions for the diversification dynamics and their underlying drivers in the PIP’s endemic biota (Fig. 1; Fig. S1).

## Box 1

### “Predictions for the tempo of diversification in the PIP

Based on the role of intrinsic factors (such as pollination strategies and polyploidy), angiosperms tend to have higher diversification rates than animals, as shown using global datasets (Hernández-Hernández et al. 2021; Scholl & Wiens 2016). We tested whether the PIP lineages show a similar pattern. In addition, broader niche width and niche shifts have been suggested to positively correlate with diversification rates (Gómez-Rodríguez *et al*. 2015; Moreira *et al*. 2024). Ecological specialisation and a tendency to retain ancestral niches (i.e., phylogenetic niche conservatism) limit diversification (Cyriac & Kodandaramaiah 2018; Da Silva *et al*. 2020). In the PIP, many clades with Gondwanan affinity are distributed in wet forests, showing niche conservatism, compared to clades with Asian affinity, which tend to occupy wider niches and have distributions in wet and dry habitats. For example, ancient Gondwanan centipedes and caecilians are restricted to wet forests (Gower *et al*. 2016; Joshi & Karanth 2013). Whereas lizards (geckos, agamids, skinks, lacertids) with Asian affinity occupy dry to wet forests, few have been shown to exhibit climatic niche evolution (e.g. *Hemidactylus* geckos) (Agarwal & Ramakrishnan 2017; Lajmi & Karanth 2020). Hence, we hypothesised that Asian clades could have higher diversification rates, owing to their broader climatic niche width than ancient Gondwanan clades, which show niche conservatism (Fig. 1). Additionally, many studies have indicated that diversification rates are negatively associated with the ages of clades. However, the causality of such a pattern has been debated recently. Some studies have attributed this phenomenon to being a statistical artefact (O’Meara & Beaulieu 2024) due to sampling bias (Harmon *et al*. 2021). In contrast, other studies speculate it to be a consequence of a hidden biological phenomenon (Henao Diaz *et al*. 2019). Our predictions for the PIP align with the negative relationship between net-diversification rates and clade age. Therefore, we explicitly test them because of the stark disparity in the age and ecology of clades of varying biogeographic affinity.

### Predictions for mode and drivers of diversification in the PIP

Temporal modes of diversification can be broadly categorised into gradual, saturated, waxing and waning, and exponential accumulation (see Fig 1, Fig S1) (Meseguer *et al*. 2022; Morlon 2014; Morlon *et al*. 2010). These diversity accumulation profiles are combined outcomes of speciation and extinction rate dynamics. Gradual accumulation (Scenario 1 - SC1) is led by constancy in diversification rates over time and is attributed to prolonged environmental stability (Benson *et al*. 2016; Couvreur *et al*. 2011; Derryberry *et al*. 2011). Within the PIP, the southern parts of the Western Ghats and the mountaintops of the Eastern Ghats have remained stable through evolutionary time. They could have served as refugia and, thus, could be the abode of lineages exhibiting gradual accumulation (Das & Chanda 1998; Gower *et al*. 2016; Joshi & Karanth 2013; Prasad *et al*. 2009). Alternatively, lineages can show time-varying changes in diversification rates (Morlon *et al*. 2010). One outcome could be an early increase in diversification followed by saturation (SC2) due to ecological limits imposed by the existing diversity (i.e.,diversity-dependence) or drastic geoclimatic perturbations (Condamine *et al*. 2019a; Moen & Morlon 2014; Rabosky 2009, 2013; Wiens 2011). Also, species diversity can decline following bouts of increase, ultimately leading to a loss in diversity (SC3) (Condamine *et al*. 2019a; Meseguer *et al*. 2022; Moen & Morlon 2014). Such “waxing and waning” diversification patterns are attributed to the inability of lineages to cope with a changing environment and the associated decrease in available niches (Condamine *et al*. 2019a; Moen & Morlon 2014). The PIP has undergone drastic climate changes, e.g., Deccan volcanism and monsoon intensification through the Cenozoic era, so waxing and waning dynamics might have occurred in many lineages (Chatterjee *et al*. 2013; Morley 2018). Lastly, lineages can exhibit an exponential increase of diversity towards the present (SC4) and are primarily attributed to geoclimatic perturbations and suitable environmental conditions promoting adaptive radiations (Meseguer *et al*. 2022; Morlon *et al*. 2010). Some of the critical geoclimatic events, like the mid-Miocene aridification, monsoon intensification and expansion of C4-plants in the PIP, could explain exponential accumulation (Agarwal & Ramakrishnan 2017; Deepak & Karanth 2018). Furthermore, the lineages may also experience episodic shifts in diversification rates, attributed to abrupt geoclimatic events that can cause a rapid increase and/or decrease in diversification rates (Kopperud *et al*. 2023; May *et al*. 2016). Species diversity within a lineage is expected to show fluctuations from general trends, as seen for SC1-4 at time periods relevant to such episodic rate shifts.”

## Materials and Methods

### Data compilation: Endemic lineages from the PIP

We reviewed published molecular phylogenetic studies on the PIP biota using Google Scholar, examining phylogenetic trees published up to December 2022. We identified well-sampled, dated phylogenetic trees for the PIP lineages, with at least 65% of the species being endemic (Table S1). We found 42 time-calibrated phylogenetic trees, of which 34 contained at least four species. These lineages broadly fell into three taxonomic categories: angiosperms, vertebrates, and invertebrates. Since there are a few endemic mammal lineages in the PIP and with no phylogenetic data and only one phylogeny for birds (*Montecincla*; n=4), we focused on time-calibrated phylogenies of well-studied herpetofauna among vertebrates.

They encompass 765 species, of which 95% (729 species) are endemic to the PIP, including 72 species from eight lineages of invertebrates (27 centipede species; 23 scorpion species; 22 mollusc species), 557 species from 20 lineages of herpetofauna (253 frogs and caecilians species; 304 turtles, geckos, agamids, skinks, lacertids, and snakes species), and 136 species from 5 lineages of angiosperms. They are distributed in terrestrial (29 lineages) and freshwater habitats (four lineages). Among terrestrial lineages, 16 are restricted to the wet zones of the Western Ghats, Eastern Ghats, and Sri Lankan Forests, and 13 are distributed across both wet and dry zones in Peninsular India and/or Sri Lanka (which together constitute the PIP). Of the 33 lineages, 20 had Asian, and 13 had Gondwanan biogeographic affinities based on divergence time and biogeographic analyses (Fig. 1; Table S1).

These lineages do not necessarily belong to a single taxonomic rank, as they range from species groups to the family level and represent independent radiations. This approach allowed for the comparison of lineages based on their ages rather than specific taxonomic ranks (such as genus) since time (clade age) is known to be associated with richness patterns (McPeek & Brown 2007). Additionally, taxa of different taxonomic ranks can have similar ages, and taxonomic ranks are arbitrary (Scholl & Wiens 2016).

### Assessing the tempo and mode of species accumulation within each lineage

#### Tempo of diversification

Net diversification rates were calculated to assess the relative pace of diversification for each lineage. For each lineage, the rates were measured with three approaches - 1) by fitting a constant-rate birth-death model from the R package *RPANDA*, 2) by taking the mean speciation and extinction rates from the posterior distribution of a Compound Poisson processes on Mass Extinction Times (CoMET) analysis using the R package *TESS* for each time bin, and 3) by taking the tip-based and mean speciation and extinction rates for all branches and tips constituting distinct clades of interest, estimated using the Cladogenetic Diversification rate Shift (ClaDS) model from the Julia package *PANDA* (Höhna *et al*. 2016; Maliet & Morlon 2022; Morlon *et al*. 2016). The ClaDS model obtained more robust estimates, particularly for a few depauperate clades. This analysis was run on well-sampled larger phylogenies (super trees) to which each clade belonged (Table S2). Then, the branch-specific and tip-rates relevant to our clades of interest were extracted. Details of the parameters and model definitions for each method are provided in the Supporting Information (Section 1.1).

We then compared diversification rates across taxonomic groups, biogeographic origins and habitat types by performing two analyses of variance in R - 1) the non-parametric Kruskal-Wallis test using the *kruskal.test* function, and (2) phylogenetic ANOVA using the *lm.rrpp* function from the *RRPP* package to control for phylogenetic non-independence. Post hoc analyses for respective methods were performed using the *dunn.test* (*dunn.test* package) and *pairwise* (*RRPP* package) (Collyer & Adams 2018; Dinno 2014). We used the tip-rates estimated by the ClaDS model and clade-rates estimated using the three methods to perform the analyses. Phylogenetic ANOVA was performed on the clade-rates and tip-rates using dated phylogenies linking all the 33 lineages and all the species in each lineage, respectively, constructed using information from timetree.org and from phylogenies of all lineages (Fig. S3) (Kumar *et al*. 2017). We also tested if clade age could predict diversification rates by performing a phylogenetic generalised least squares regression (PGLS) using the clade-rates (ClaDS, CoMET, RPANDA) and ages.

#### Mode of diversification

The temporal mode of diversification was evaluated by fitting homogeneous birth-death models (constant-rate and time-varying) to all 33 lineages. Constant-rate models included constant-rate pure birth (CB - only a constant speciation rate, no extinction) and constant-rate birth and death models (CBD - constant and finite speciation and extinction rates). Time-varying models included combinations of linearly (⍰(t) = ⍰_0_ + ⍰·t, ⍰(t) = ⍰_o_ + ⍰·t) or exponentially (⍰ = ⍰_0_ ·*e*^⍰·t^, ⍰ = ⍰_o_ ·*e*^⍰·t^) varying (with time - t) birth (⍰) and death (⍰) rates (⍰_0_ and ⍰_o_ are the expected speciation and extinction rates at t=0, ⍰ and ⍰ are the coefficients of gain or decay in the functions and time, here, was interpreted as time from the present) (Condamine *et al*. 2019a; Morlon *et al*. 2010).

To model the possibility of symmetrical local peaks and dips (sharp increase followed by a decrease in diversification rates or *vice versa*) in the diversification rate in time, we developed a continuous time-varying mathematical function to model symmetrical local peaks in diversification and to perform model fitting in the *RPANDA* framework. This function follows the “Witch of Agnesi” curve, where the diversification rates diverge from a baseline constant rate: 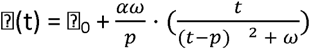, where ⍰_0_ represents a baseline rate of speciation, ⍰ represents the intensity of the peak (or dip), represents the spread of the peak or dip, and *p* represents the position of the peak (or dip) in My (Spencer 1940). Similar rate shifts in extinction were also modelled. This approach also helped qualitatively assess the role of specific geoclimatic events on the diversification of lineages corresponding to the time of occurrence of a peak (or dip). We called this model the “symmetrical episodic shift” (abbreviated as the SES model later and in the supporting information.

Model fitting using constant-rate and time-varying birth-death models was executed in *RPANDA* using the *fit_bd* function (Morlon *et al*. 2016). Best-fitting models were selected based on AICc scores, and ⍰AICc scores were calculated as absolute differences in the AICc scores between each model and the model with the lowest score. The models with ⍰AICc scores less than two were chosen for each lineage (Burnham & Anderson 2004). If multiple models showed ⍰AICc below two i.e., all models were equally likely, then the model with the least parameter complexity was chosen as the best-fitting model. If multiple models of similar AICc and the same complexity existed, the model with the lowest AICc was selected as the best-fitting model. However, if CB or CBD were found along with other models to be equally likely, the corresponding constant-rate model was considered the best-fitting model for a lineage, as the simplest model could not be rejected. We used the parameter values (⍰_0_, ⍰, ⍰_o_ and ⍰) of the selected diversification models to evaluate the scenarios mentioned in Table S2 and Fig. 1, Fig. S1.

Furthermore, we performed the CoMET analysis using the R package *TESS* (Höhna *et al*. 2016; May *et al*. 2016) to detect times of significant rate shifts in a lineage’s history. The CoMET analysis involves binning the history of a clade into 100 equal time bins for speciation and extinction rates to detect rate shifts within each time bin. For each lineage, we extracted the Bayes Factor (BF) supports for rate shifts at these 100 time bins and chose the ones where 2 * ln(BF) scores were substantially higher than constant rates ≥4.6 (Hohna *et al*. 2015; Jeffreys 1998).

### Determining drivers of diversification rates

#### Assessing drivers of diversification patterns

We evaluated the role of important climatic variables on the diversification patterns of biota from the PIP - mainly temperature, changes in seasonality patterns and Miocene aridification. We obtained paleoclimate data for temperature (Zachos *et al*. 2008), reconstructed Himalayan elevations (Ding *et al*. 2017) (as a proxy for the onset of seasonality patterns) and pedogenic carbon (precipitated carbon found in soil) content from the Potwar plateau (Clift & Webb 2019; Quade *et al*. 1989) (as a proxy for aridification and expansion of C4 plants). The paleo-climatic data was smoothened through spline interpolation using the *ss* function from the *npreg* package (Helwig 2020) in R. Subsequently, we used the same combinations for rate-dependencies (birth and death) as on time (except the episodic local shift model), on each of these variables and fitted these models using the *fit_env* function from the *RPANDA* package. Also, we included the functions (⍰(T) = ⍰_0_ ·*e*^-⍰/T^, ⍰(T) = ⍰_o_ ·*e*^-⍰/T^) inspired by the metabolic theory of biodiversity as provided in a recent study (Condamine *et al*. 2019a); (Brown *et al*. 2004). Among biotic drivers of diversification, we checked for diversity-dependence (DD) on speciation and extinction rates, incorporating both linear and exponential dependencies (Condamine *et al*. 2019a; Etienne *et al*. 2023).

A factor analysis of mixed data (FAMD) was performed to assess the correlations among different ecological and evolutionary drivers and to visualise the spread of lineages relative to these axes (Pagès 2004, Table S6). This reduced ecological-evolutionary framework was further used to compare the clusters of diversification scenarios and different categorical predictors while also considering the variations in species richness, diversification rates and clade ages (quantitative variables). We also performed the permutational multivariate analysis of variance (PermANOVA), utilising the dissimilarities (Gower’s distance) between lineages calculated from the matrix used in FAMD (Table S6) to assess the statistical significance of clustering (Anderson 2017). Before performing PermANOVA, we performed a Mantel test (using Pearson’s correlation) using the *mantel* function from the *vegan* package in R to check if the distance matrix was correlated to the covariance matrix generated from the dated phylogeny (Fig. S3.e - upper panel) linking all the 33 lineages. All the analyses were performed using functions from the R packages *FactoMineR* and *factoextra* (Irnawati *et al*. 2021; Lê *et al*. 2008).

### Assessing congruent classes of diversification models

A recent study by Louca and Pennell highlighted the possibility of an infinite class of congruent models which would equally fit a phylogeny, making diversification models unreliable (Louca & Pennell 2020). Hence, in addition to the canonical diversification rates (CDR), we assessed and summarised the trends in pulled diversification rates (PDR) using two approaches.

Firstly, we fitted PDR models present in the *castor* package in R (Louca 2017). Specifically, we used the *fit_hbd_pdr_on_grid* with a NULL value for the age_grid parameter to find the likelihood, AIC and BIC of fitting a model where the birth-death rates have remained “time-independent” (Louca & Pennell 2020). This was followed by assessing the disparity in the pulled diversification rates across taxonomic groups, biogeographic affinities and habitat types. Following this, we used the *fit_hbd_pdr_on_best_grid_size*to assess the best number of time grids (between 1 to 5) that the evolutionary time (clade age) of a lineage can be split into, in each of which rates can vary linearly or exponentially (i.e. rates are piecewise time-dependent) (Louca & Pennell 2020). The AIC and BIC of this time-varying model were then compared with those of the time-independent model to assess the dynamicity in the diversification rates.

Secondly, we performed the Congruent Rate Analyses in Birth–-death Scenarios (CRABS) test to account for the unreliability of the CoMET estimates (Höhna *et al*. 2022). We used the canonical rate through time estimates from the CoMET analysis for each lineage, specified alternate extinction rate functions through time, and computed the corresponding speciation rate functions to construct a set of congruent classes. These extinction rate functions were defined for each time bin, and they could remain constant or vary linearly, exponentially and sigmoidally with time. Also, different values were taken for the parameters (such as slope) that govern the rate of change in those functions to span a wide range of hypotheses. The temporal trends across the congruent classes were then visually assessed. Custom code for this test was written, modifying the relevant code from a recent study (Cyriac *et al*. 2024).

### Assessing reliability of estimates with posterior predictive tests

To assess the reliability of our diversification rate estimates, we performed simulations using the parameters of the best-fit models for each lineage. We calculated errors in model parameters, the timing of significant rate shifts, and in estimating the diversification scenarios. We performed these tests because our dataset was dominated by depauperate clades and small phylogenies, which could affect the robustness of diversification rate estimates. We simulated 100 trees using the best-fit model parameters per lineage using the *tess.sim.age* function from *TESS*. Each simulated tree was constrained to be of the same clade size and age as the empirical tree, while branching patterns were allowed to vary according to the model parameters. We then analysed these simulated trees using the same methods as the empirical phylogenies (RPANDA models and CoMET). We compared the model parameters and diversification dynamics with the empirical trees.

Firstly, the best-fitting birth-death model was assessed in the *RPANDA* framework on each of the 3300 trees (100 simulated trees X 33 sets of model parameters), and the diversification scenarios were assessed. For each set of 100 trees, we calculated the proportion of trees supporting the same diversification scenario as the corresponding empirical diversification scenario. This is equivalent to the proportion of correctness in estimating the diversification scenarios. Secondly, in the CoMET analysis, scenario correctness was determined by comparing the presence or absence of episodic rate shifts in simulated trees relative to empirical trees, as CoMET only provides the timing of such shifts.

The error in model parameter estimation (speciation and extinction rates at present—⍰_0_ and ⍰_0_, and the slopes of speciation and extinction rate functions over time—⍰ and ⍰) was calculated using a method analogous to Cohen’s D (Cohen 2013; Cumming 2013). For each parameter, we subtracted the values of the 100 simulated trees from the empirical tree and normalised each difference by the standard deviation of the parameter across the simulations. Finally, we estimated the timing of significant diversification rate shifts in the simulated trees using the SES model and CoMET analysis and compared them with empirical estimates.

## Results

### Mode and tempo of species accumulation

#### Tempo of diversification

The mean speciation, extinction, and net diversification rates (canonical diversification rates) of 33 lineages were measured using RPANDA models, CoMET analyses and the ClaDS model to assess the tempo of diversification. The trends in net-diversification rates were remarkably consistent across methods (Fig. S3); therefore, only tip-based net-diversification rates estimated using ClaDS have been discussed. We found a significant disparity in the diversification rates among the different taxonomic groups (*p* < 0.05 for pairwise comparisons using the Kruskal-Wallis test and Dunn’s test), where invertebrates showed the lowest diversification rates, and angiosperms had the highest diversification rates (*p* < 0.05, Fig. 2.a; Fig. S3a, b, c; Table S2). The diversification rates were significantly lower among the Gondwanan than in Asian lineages (*p* < 0.05, Fig. 2.b; Fig. S3; Table S2), although diversification rates across habitat types were not significant. However, upon accounting for phylogenetic relatedness (phylogenetic ANOVA), the disparities across taxonomic groups and biogeographic affinities were found to be non-significant (*p* > 0.05; Table S2). Also, net-diversification and speciation rates were significantly associated with clade ages (phylogenetic signal - Pagel’s ⍰ = 0.487, *p* < 0.05; Table S2; Fig. S3) even after controlling for phylogenetic non-independence (*R*^2^ > 0.1, *p* < 0.05, Fig. 2c; Fig. S3; Table S2). Differences in speciation rates mainly contributed to disparities in net-diversification rates because extinction rates across groups were low (Fig. S3). Trends were similar in pulled diversification rates, although the association with clade ages was non-significant (*p* = 0.120, Fig. S3.d).

**Fig. 2.**
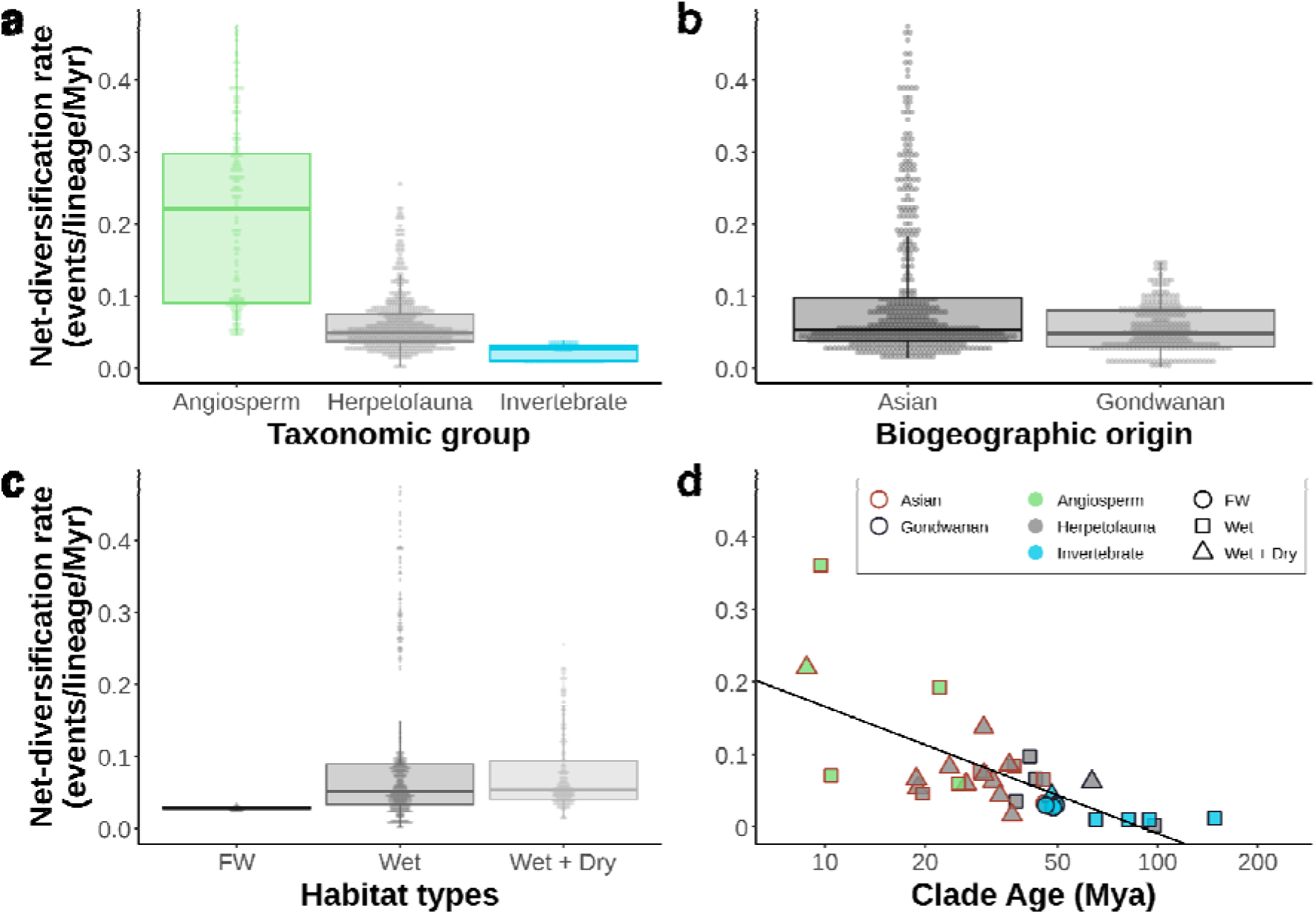
The tempo of diversification. Boxplots represent the net-diversification rates (tip-rates) calculated using the ClaDS model across taxonomic groups **(a)**, biogeographic origins **(b)** and habitat types **(c)**. The scatter and regression plot **(d)** illustrates net-diversification rates (calculated using the CoMET analyses) against the clade age for each lineage. Statistical information relevant to comparisons has been provided in Table S2. Boxplots illustrating the disparities in rates (clade-rates) calculated using all three methods have been provided in Fig. S3.

#### Mode of diversification

Constant and time-varying models of canonical diversification rates (CDR) were fitted to all lineages. We found that 17 lineages supported the constant-rate birth (CB) model, which led to gradual expansion (SC1), and 16 supported time-varying birth-death models that led to diversification slowdowns (SC2) and Waxing and Waning (SC3) (Table 2; Table S1, Fig. 3). These 17 lineages were spread across taxonomic groups, biogeographic origins and habitat types (Fig. S5). Among 16 lineages supporting time-varying birth-death models, 15 also showed episodic shifts in the diversification rates. Only one lineage (*Hemidactylus*) showed saturated accumulation (SC2) without episodic rate shifts. Among the 15 lineages with episodic rate shifts, two showed slowdowns (SC2), five supported waxing and waning (SC3), and eight showed episodic rate shifts. Episodic rate shifts were inferred by CoMET and the symmetrical shift model (SES) in *RPANDA* in 11 and eight lineages, respectively, out of which both methods concordantly inferred four (Fig. 3.b). However, none of the lineages favoured exponential accumulation (SC4). Parameter estimates of birth-death model-fitting (*RPANDA*) and CoMET analysis are provided in Table S3.

**Fig. 3.**
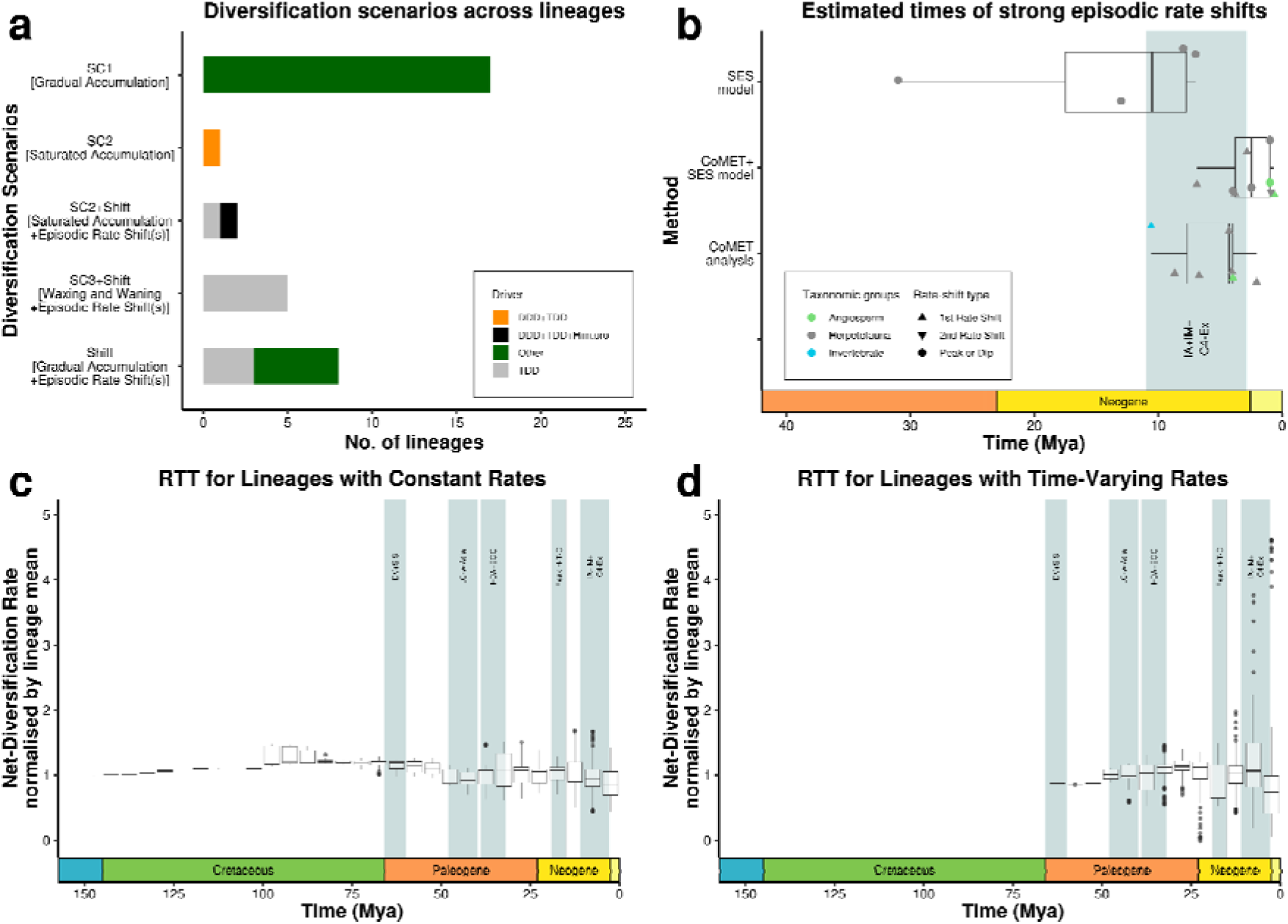
Diversification patterns and drivers. Vertical bar plots **(a)** represent the prevalence of the best-fitted diversification scenarios for the PIP lineages inferred by RPANDA models - SC1 - gradual accumulation, SC2 - slowdowns, SC3 - Waxing and Waning and Episodic Rate Shifts. The drivers of the diversification scenarios (paleo-temperature - TDD, diversity-dependence - DDD and Himalayan orogeny - HDD) have been indicated by different colours. Horizontal box plots **(b)** illustrate the temporal distribution of episodic rate shifts, inferred by the SES model (in RPANDA) and the CoMET analysis, each indicated by the circles and triangles, respectively. Circles (illustrating peaks and dips) and triangles (illustrating one-way rate shifts) represent inferred rate shifts by different methods, and more than one dot can be relevant to one lineage. In case of multiple rate shifts (in the CoMET analysis) in a lineage, an additional inverted triangle has been used. Vertical box plots **(c and d)** represent the distribution of net-diversification rates of lineages for each time bin of 5 My. The rates-through-time (RTT) plot uses the net-diversification rates relative to respective lineage means. The normalised net-diversification rate in 17 lineages favouring constant rate models show less deviation than the 16 lineages favouring time-varying birth-death models.

Pulled Diversification Rates (PDR) suggested temporal non-uniformity (best age grid size >= 2) of diversification rates in 12 lineages and constancy of rates through time in 19 lineages. The remaining two clades were depauperate, and PDR functions could not be successfully fitted (Table 1, S1, S3). We also performed the CRABS test to assess the trends of congruent classes across the evolutionary time of each lineage. We found a high prevalence of constancy in rates through time for 13 lineages, conclusive recent rate shifts (either a declining trend or a peak) in 17 lineages, and a decreasing trend in multiple time bins in one lineage (*Pseudophilautus*) (Fig. S5). The remaining two clades showed ambiguous trends across congruence classes across their evolutionary times (Table 2, Table S1).

**Table 1.**
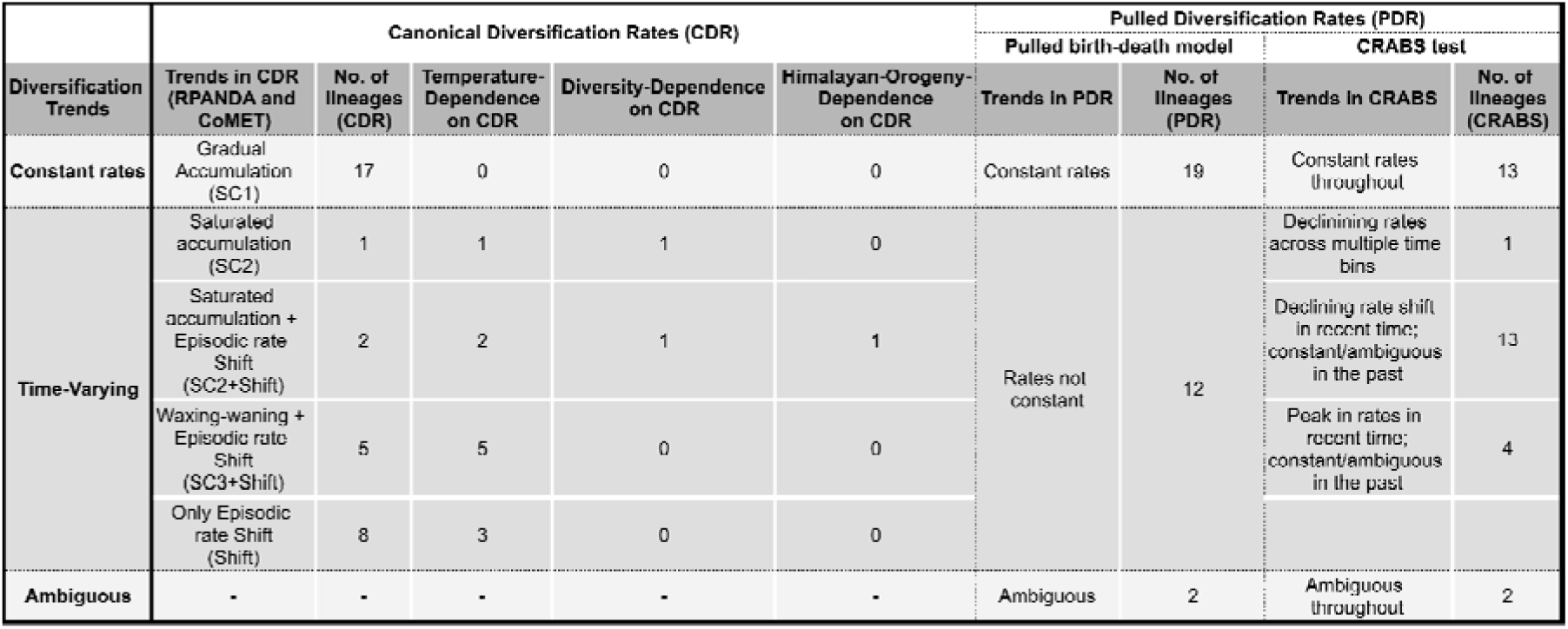
Patterns and drivers of diversification. Numbers indicate the number of lineages favouring each canonical diversification scenario, pulled diversification trends, and diversification dependence on paleotemperature, existing diversity, Himalayan orogeny, and the expansion of C4 Plants.

| Diversification Trends | Canonical Diversification Rates (CDR) |  |  |  |  | Pulled Diversification Rates (PDR) |  |  |  |
| --- | --- | --- | --- | --- | --- | --- | --- | --- | --- |
|  | Trends in CDR (RPANDA and CoMET) | No. of lineages (CDR) | Temperature-Dependence on CDR | Diversity-Dependence on CDR | Himalayan-Orogeny-Dependence on CDR | Pulled birth-death model |  | CRABS test |  |
|  |  |  |  |  |  | Trends in PDR | No. of lineages (PDR) | Trends in CRABS | No. of lineages (CRABS) |
| Constant rates | Gradual Accumulation (SC1) | 17 | 0 | 0 | 0 | Constant rates | 19 | Constant rates throughout | 13 |
| Time-Varying | Saturated accumulation (SC2) | 1 | 1 | 1 | 0 | Rates not constant | 12 | Declining rates across multiple time bins | 1 |
|  | Saturated accumulation + Episodic rate Shift (SC2+Shift) | 2 | 2 | 1 | 1 |  |  | Declining rate shift in recent time; constant/ambiguous in the past | 13 |
|  | Waxing-waning + Episodic rate Shift (SC3+Shift) | 5 | 5 | 0 | 0 |  |  | Peak in rates in recent time; constant/ambiguous in the past | 4 |
|  | Only Episodic rate Shift (Shift) | 8 | 3 | 0 | 0 |  |  |  |  |
| Ambiguous | - | - | - | - | - | Ambiguous | 2 | Ambiguous throughout | 2 |

The errors in the estimates of the model parameters (of the simulated trees) relevant to the birth-death models in RPANDA (speciation and extinction rates at present, respectively - ⍰_0_ and ⍰_0_, and the slopes in speciation and extinction rate functions, respectively, through time - ⍰ and ⍰) existed primarily within +2 and −2 (Fig. S7.c). This indicates that estimated values for the parameters relevant to the simulated trees were within two standard deviations from the empirical values. However, a high error was observed in the simulated trees corresponding to the model parameters of two clades - *Memecylon* (in ⍰_0_) and *Geckoella* (in ⍰). Additionally, the proportion of correctly inferred diversification scenarios varied across lineages (Fig. S7.a, b). Assessments in the RPANDA framework suggested that 13 sets of empirical trees (corresponding to 12 lineages’ best-fit model parameters) had low proportions (< 0.6) of correctly inferred scenarios. The CoMET analyses suggested that eight sets of simulated trees (six same as above and two additional sets) had low proportions (< 0.6) of correctly inferring the episodic rate shifts. Therefore, 15 sets of simulated trees (out of 33) had < 60% recovery of the empirical diversification scenarios. Additionally, in both frameworks, this proportion did not show any general trends with clade size but rather with the type of empirical diversification scenario where many clades with a time-varying diversification scenario had low proportions of correctness. Also, the SES model overestimated episodic rate shifts compared to the CoMET analysis (Fig. S7.b).

### Abiotic and biotic drivers of diversification patterns

There was no strong association between the diversification scenarios and taxonomic groups, habitat types or biogeographic origins (Fig. 3, 5; Fig. S5), except in invertebrates, where seven out of eight lineages supported the gradual accumulation scenario (SC1). The time-varying diversification rates were primarily associated with paleo-temperature and existing diversity through time (i.e. diversity-dependence) (Table 2; Table S1 and S4; Fig. 3, 5). Of 16 lineages that supported time-varying models and/or shifts over constant rates, 11 showed temperature dependence (TD). Among these 11 TD lineages, two lineages equally supported (⍰AICc ≤ 2) diversity-dependence (DD) (Fig. 3; Table 2, S1, S4), one of these lineages supported Himalayan-orogeny-dependence, and none supported C4-plant-expansion-dependence (Fig. 3, Table S4). Speciation rates in all the lineages favouring temperature-dependence were positively associated with paleotemperature (“Temp.alpha” in Table S5). Extinction rates for those lineages were either constant or positively associated (“Temp.beta” in Table S5), except for *Cnemaspis*, which favoured a negative association of extinction rates with paleotemperature.

Episodic diversification rate shifts were inferred for 15 lineages, and 14 occurred in the Neogene (Fig. 3.b). Episodic rate shifts in four lineages were inferred jointly by CoMET analysis and SES model, and rate shifts in four and seven additional lineages were individually inferred by the SES model and CoMET analysis, respectively. Eleven among the 15 lineages above showed diversification rate shifts within the geological period that experienced a mid-Miocene intensification of aridification, monsoons and expansion of C4-plants in the PIP (11 - 3 Mya). Three among these and two of the remaining lineages showed rate shifts younger than 3 Mya in the Neogene and early Quaternary. Sharp declines in global temperatures from around 2 Mya to the present are concurrent with these recent rate shifts in these lineages, all of which are also declines in diversification rates (Fig. 3, 4). Episodic rate shifts in the remaining two lineages - *Gegeneophis* and Uropeltidae - happened around 31 and 13 Mya, respectively.

In the FAMD analysis, the first two dimensions explained ∼43% (Fig. 5 - Axis 1: 24%, Axis 2: 19%) of the total variation (Fig 5). Fig. 4a represents the squared correlation between different quantitative and qualitative (categorical) variables. Clustering patterns of lineages across different categorical predictors have been provided in Fig. 4b. Importantly, taxonomic identity, biogeographic affinity, net-diversification rate, clade age and species richness, contributed to dimension one and diversity-dependence and Himalayan-orogeny-dependence of diversification contributed to dimension two (Fig S6a). Diversification scenarios contributed highly to both ordination dimensions.

**Fig. 4.**
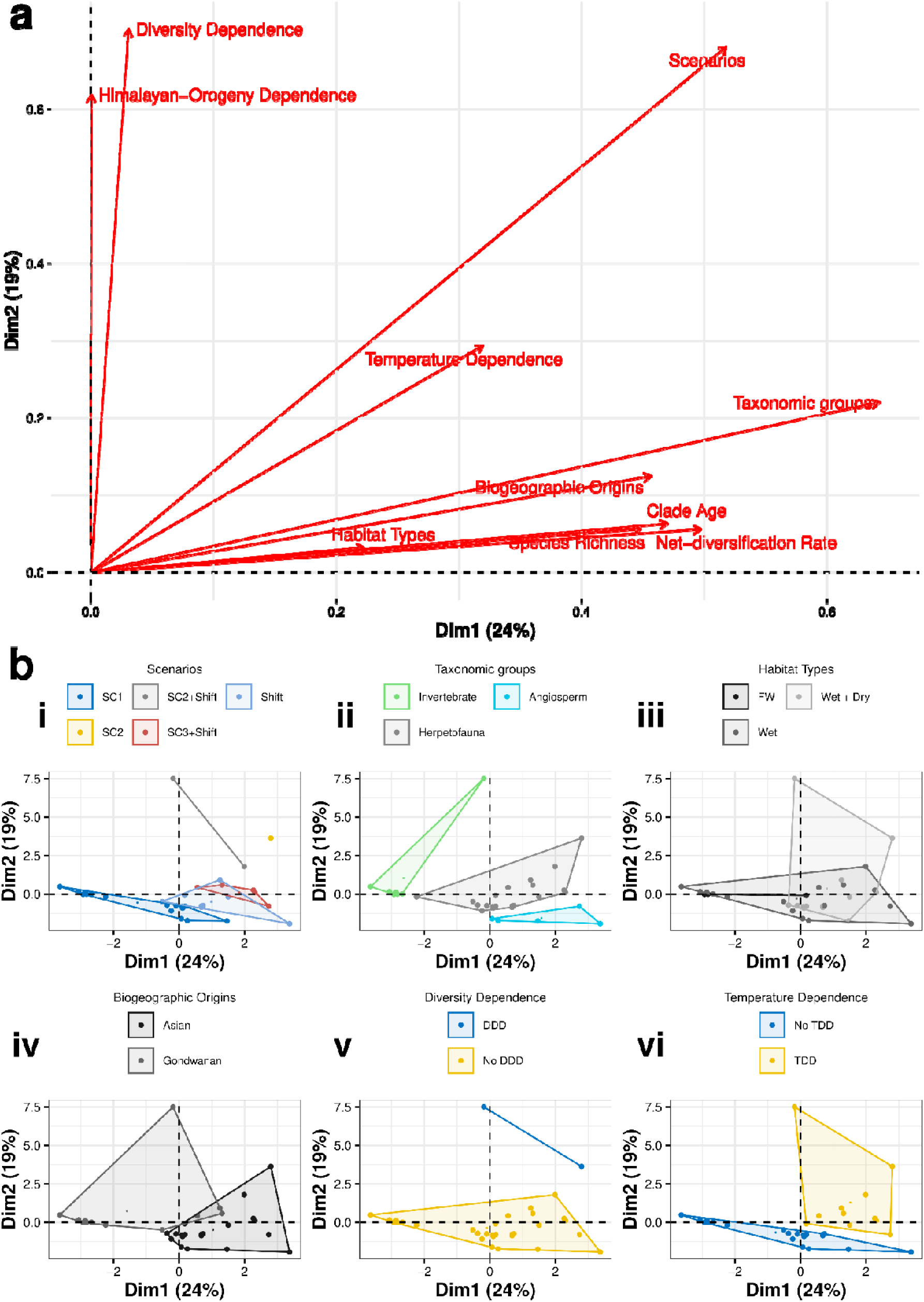
Factor Analysis of Mixed Data ordination plots made using 13 relevant ecological and evolutionary parameters for 33 lineages. **(a)** The first two dimensions of the ordination analysis explain that the variables positioned in the space represent their relative contribution to the relevant axes. The positions of the variables relative to each other represent the strength of their associations; **(b)** each panel represents a categorical variable, and the clusters within them represent the categories. Clusters relevant to diversification scenarios are largely congruent with diversity-dependence and temperature-dependence. Check SI Table 5 for information relevant to PERMANOVA tests for inferring significantly distinct clusters within each categorical variable.

Clusters (convex hulls) for diversification scenarios were found to be the most congruent with that of Temperature-Dependent Diversification (TDD). Lineages showing temperature dependence occupied similar positions in the FAMD-space as the lineages favouring diversification scenarios with time-varying rates (i.e., all scenarios except SC1). Conversely, the lineages that did not follow TDD clustered similarly to those that supported the constant-rate gradual accumulation scenario. Moreover, a few clusters in taxonomic groups, biogeographic origins and habitat types were non-overlapping (Fig. 4b.ii, iii, iv) and significantly distinct (Table S7). Clade age was inversely correlated with net-diversification rate (Fig S6.b - upper panel), as inferred earlier by the PGLS regression and species richness was found to be more closely associated with net-diversification rate than clade age.

## Discussion

Our meta-analyses of 33 lineages demonstrated that species traits, phylogenetic history and global and regional-level geo-climatic history governed the dynamics of diversification of endemic lineages in the PIP. The phylogenetic history and age of lineages were strongly associated with the pace of diversification in the endemic PIP lineages. Gradual accumulation was the most prevalent diversification scenario. Furthermore, global paleotemperature and mid-Miocene aridification influenced the temporal non-uniformity in the rates of many lineages.

### The tempo of diversification in the PIP biota

The PIP harbours biotic assemblages of varying ages, biogeographic affinities and habitat preferences, which vary substantially in their diversification rates. Although trends in diversification rates were observed across taxonomic identities and biogeographic affinities, the phylogeny-informed ANOVA indicated that phylogenetic relatedness primarily drove these disparities, suggesting that unique selection pressures acted on each lineage. While taxonomic identity loosely reflects phylogenetic relatedness, the phylogenetically controlled trend in diversification rates across groups implies that shared ancestry, rather than group-level differences, largely explains similar rates. However, each taxonomic group could have unique species traits shaped by shared ancestry, which can, in turn, influence diversification rates.

Clade-specific traits, such as range size, dispersal abilities, and ecological niches, could influence net diversification rates (Scholl & Wiens 2016). As predicted, angiosperms (flowering plants) had the highest diversification rates, which could be because they typically have larger clade-level range sizes (Hernández-Hernández *et al*. 2021), polyploidy (Heslop-Harrison *et al*. 2023) and higher dispersal ability (Suárez *et al*. 2022). It means that angiosperms can occupy a wide range of ecological niches and are more likely to colonise new areas, promoting their diversification. In contrast, centipedes, scorpions and molluscs had the lowest net diversification rates among invertebrates. Centipedes and scorpions are among the top predators in soil ecosystems and tend to show specialisation for ecological niches, i.e., being both fossorial (burrowing) and predators may limit their opportunities for speciation. They also have relatively low dispersal abilities (Bharti *et al*. 2023; Joshi *et al*. 2020; Loria & Prendini 2020), which could further restrict their ability to colonise new areas and diversify (Wiens 2004). Shieldtails, another group of burrowing predators, exhibited similar patterns (Cyriac & Kodandaramaiah 2018) (Fig. 2, Table S3). For the freshwater molluscs, the diversification rates could be constrained by the limited surface area and niche availability in freshwater ecosystems, relative to their terrestrial counterparts (Román-Palacios *et al*. 2022). Moreover, given that these molluscs inhabit river basins and are not obligate lentic fauna (Liu *et al*. 2022; Tripathy & Mukhopadhayay 2015), they can have high genetic connectivity through rivers, which could limit allopatry and, hence, limit diversification rates.

The PIP biota consists of lineages with either ancient Gondwanan or younger Asian biogeographic affinities, both of which have undergone *in situ* speciation (Karanth 2021). This provided a unique opportunity to examine the influence of biogeographic affinity and lineage age on diversification rates. Interestingly, we observed a negative relationship between net diversification rates and clade ages, with a high phylogenetic signal (i.e closely related lineages have similar rates and ages) across lineages (Fig. 2.d; Fig S3.a, b, c; Fig S6.b - lower panel; Table S2). Consequently, older lineages with Gondwanan affinities tend to have lower diversification rates than younger Asian clades. One possible explanation for this pattern is phylogenetic niche conservatism—the tendency of species to retain ancestral ecological traits over long evolutionary timescales. Most ancient Gondwanan lineages exhibit high ancestral niche retention (Gower *et al*. 2016; Joshi & Karanth 2013). For example, many lineages are restricted to wet forests and are fossorial, such as soil arthropods, blind snakes, and caecilians (Karanth 2021). Also, it has been argued that ecological specialisation may limit diversification rates, resulting in low diversification rates (Cyriac & Kodandaramaiah 2018; Da Silva *et al*. 2020; Wiens 2004). In contrast, many younger lineages occupy diverse ecological and climatic niches, contributing to higher diversification rates. For instance, lizards such as *Hemidactylus*, *Geckoella*, and *Ophisops* are found in wet and dry forests and inhabit various microhabitats, including rocky outcrops, forest floors, leaf litter, and trees (Agarwal & Karanth 2015; Agarwal & Ramakrishnan 2017; Lajmi & Karanth 2020). Thus, lineage age and ecological factors seemed closely linked in the PIP, with ecological constraints potentially shaping diversification rates and contributing to disparities across lineages.

Tropical forests are often described as either "museums" preserving ancient lineages or "cradles" of “new adaptive complexes” (Dagallier et al. 2020; Eiserhardt et al. 2017; Stebbins 1974). A region functions as a museum when old lineages persist while going extinct elsewhere, likely due to environmental stability, large habitat size, and lower extinction rates (Jablonski et al. 2006; Eiserhardt et al. 2017). Conversely, a cradle comprises young species resulting from rapid and dynamic diversification, habitat heterogeneity and climatic instability. Older and younger lineages often align with the museum and cradle models. A recent study on endemic woody plants suggests that the Western Ghats is both a museum and a cradle of diversity (Gopal *et al*. 2023). Given that many endemic lineages (e.g., centipedes, scorpions, geckos) found in the Western Ghats are not restricted to the biodiversity hotspot but have pan-PIP distributions, the entirety of the PIP may harbour museums and cradles. The old lineages studied here follow the museum model, with low diversification rates, longer persistence times and gradual diversification. In contrast, younger lineages show characteristics of the cradle model of more dynamic and higher diversification rates. This could explain the negative relationship between diversification rates and clade ages (Fig. 2d, S3.a, b, c). It would be worth exploring this further to validate this relationship with more lineages with distinct ages, richness patterns and ecologies across tropical regions.

It is important to note that speciation primarily drove the diversification in the PIP lineages, as extinction rates remained low across lineages (Fig. S3b). Most of these lineages have diversified in the Cenozoic after the Deccan volcanism (28 lineages out of 33), which led to massive extinctions (Keller *et al*. 2009; Prasad *et al*. 2009). Hence, the signature of extinctions may not have been captured in the current analyses. Also, low extinctions could be statistical artefacts of the employed methods, owing to the fewer tips in the phylogenies. Old relictual fauna with high diversity, like millipedes, would be appropriate to check signatures of clade-wide mass extinctions. Furthermore, old Gondwanan clades distributed throughout Asia (e.g. Dipterocarpaceae) can be better candidates to capture such signatures of high extinction (Bansal *et al*. 2022).

### Tropical habitat stability and gradual accumulation

More than half of the lineages (∼52%) supported the constant-rate pure-birth model, indicating a gradual accumulation of diversity over time. This gradual accumulation is attributed to climatic stability in regions of diversity expansion (Couvreur *et al*. 2011; Derryberry *et al*. 2011; Meseguer *et al*. 2022), especially in tropical biomes like the tropical forests of the PIP. A recent study on the diversification patterns of endemic frog lineages in the Western Ghats also invoked the role of climatic stability on diversification, as gradual accumulation was found to be the prevalent pattern (Cyriac *et al*. 2024). This is quite intriguing given that the PIP has experienced significant climatic changes throughout the Cenozoic Era, during which predominant diversity has accumulated (Bansal *et al*. 2022; Chatterjee *et al*. 2013; Morley 2018). Particularly, it experienced a climatic shift from perhumid to seasonal wet and also went through intensification in monsoons and aridification in the mid-Miocene. Therefore, we suspect some regions in the PIP may have served as refugia, such as the southern Western Ghats, which acted as refugia during the Deccan volcanism in the Cretaceous and later in the mid-Miocene (Joshi & Karanth 2013; Prasad *et al*. 2009). Future studies should conduct spatially explicit diversification analyses to examine how diversification rates vary across space in the PIP. Like the PIP, 50–67% of endemic plant and tetrapod lineages in the Neotropics supported gradual accumulation (Meseguer *et al*. 2022). This suggests that stable climate refugia supporting gradual diversity accumulation is not unique to the PIP but may have global relevance in tropical regions.

Alternately, the constant rates of diversity accumulation among the PIP lineages also suggest that the habitat/ecosystem could have been stable. However, the angiosperms in the PIP have not been stable until at least the late Paleocene based on fossil pollen data (Bansal *et al*. 2022; Parmar *et al*. 2023; Prasad *et al*. 2009). These extinct floral assemblages of PIP now have extant relatives in Southeast Asia; therefore, the PIP is called an “evolutionary graveyard” (Bacon *et al*. 2022). Both extinct Gondwanan and current Asian floral lineages occupy similar climatic niches, where the current flora in the PIP is shaped by the dispersal events from Southeast Asia rather than the retention of Gondwanan relics (Klaus *et al*. 2016). Therefore, a rapid turnover of the forest ecosystems, where ecologically similar species of Asian origins replaced Gondwanan lineages, could have provided habitat stability, indicating that realised habitats of the lineages have been stable despite the landscape not being stable. A recent global study on angiosperm diversification also inferred the prevalence of rapid turnover at lower taxonomic levels following high extinctions (i.e. similar species took over the niches of extinct species very rapidly) during the Cretaceous-Paleogene (K-Pg) (Thompson & Ramírez-Barahona 2023).

### Paleoclimate, ecological limits and time-varying diversification rates

Lineages with time-varying diversification rates showed a complex interplay between climatic events and fluctuations, episodic rate shifts, and species traits. The Neogene period, especially during the mid-Miocene, witnessed intensified aridification and seasonality in the Indian subcontinent, which is believed to have significantly impacted diversification, particularly among lizards (Agarwal & Ramakrishnan 2017; Deepak & Karanth 2018). While previous studies implicated a connection between climatic events and diversification, a robust statistical framework to test these hypotheses was limited. Of the 15 lineages with episodic rate shifts, 12 showed rate shifts during the Neogene (23–2.6 Mya), and rate shifts younger than 3 Mya occurred in five lineages. These shifts involved an overall decline in diversification rates (SC2+Shift or SC3+Shift) due to ecological limits or symmetrical episodic shifts (peaks or dips in diversification rate from a baseline rate), indicating alternating periods of increased and decreased diversification. Eleven of these lineages showed rate shifts that consistently overlapped with the temporal range of climatic events, suggesting a potential link between climatic changes and diversification in these lineages. We also reported strict declines in diversification rates in five lineages, as mentioned in the results, which appear to be concurrent with drastic declines in global temperatures during the late Neogene and throughout the Quaternary (Fig. 3, 4).

Another important diversification pattern observed among multiple taxa globally and in tropical regions is the slowdown in diversification rates. A global study assessing the drivers of slowdowns inferred the role of diversity-dependence (or ecological limits) and declining global paleotemperature (Condamine *et al*. 2019a; Moen & Morlon 2014). In the PIP, diversification slowdown and diversity-dependence were earlier observed in *Hemidactylus* geckos (Lajmi & Karanth 2020). Apart from this lineage, we found slowdowns in two additional lineages of scorpions (Heterometrinae) and bush frogs (*Pseudophilautus*). These patterns were also linked to diversity-dependence and temperature dependence, suggesting that changes in species diversity and temperature fluctuations play an important role in diversification rates. In five lineages, waxing and waning (diversity declines) were detected, primarily influenced by paleotemperature fluctuations but not ecological limits. Interestingly, speciation rates of most lineages favouring temperature-dependence were found to be positively associated with paleotemperature. Also, extinction rates were either positively associated or were low and constant. This indicates that, like in many taxa globally and in specific regions (like the Neotropics), warmer climates did facilitate higher speciation and extinction events in the PIP (Condamine *et al*. 2019b; Vasconcellos *et al*. 2021). Lastly, variation in speciation rates contributed to diversity declines in these lineages, except in one where the extinction rate was relatively high (*Ophisops*) (Fig. S3b, Table S1).

Mid-Miocene intensification of aridification and monsoons have also been hypothesised to drive exponential species accumulation in the PIP (SC4 - Fig. 1, S1) (Agarwal & Ramakrishnan 2017; Deepak & Karanth 2018). However, none of the lineages in our dataset supported exponential accumulation. Also, the role of Himalayan orogeny and the expansion of C4 plants in explaining diversification among the PIP biota remained limited. Only a few lineages depended on these factors, and their influence may have been episodic and inconsistent throughout lineage history. Hence, derivative curves of these factors, highlighting periods of instability, can be utilised to check the dependence on diversification. Other factors, such as paleo-temperature, have played a more significant role in diversification. Analytical methods to determine the dependence of diversification in specific time intervals (piecewise dependence) on factors like paleo-climate and existing diversity can be useful in these scenarios.

### Diversification in tropical areas: limitations and the way forward

There has been tremendous growth in methods inferring the diversification tempo and mode using time-calibrated phylogenies (Morlon *et al*. 2024). However, one of the limiting factors to applying these across taxa and regions has been the requirement of large datasets, where the number of tips analysed in each lineage should be high. Many clades studied in our study and other areas are inherently species-poor. To overcome this, we used birth-death models implemented in different frameworks, such as homogeneous birth-death models from *RPANDA,* episodically varying models in CoMET, and tip-based rates and mean (tip and branch) rates inferred from the ClaDS model. After performing a meticulous review to verify the sampling of the species in each clade, we assigned a sampling fraction of 1 to all the analyses across all lineages. Even for underrepresented taxa, like invertebrates and freshwater fauna, we mainly focused on the ones with robust phylogenetic species hypotheses with the maximal sampling fractions mentioned in the individual studies. However, with the ever-increasing usage of integrative taxonomy, more species are being described, and species boundaries are being redefined (Joshi & Agarwal 2021). Thus, a sampling fraction of 1 might contribute to an erroneous assessment in diversification models, especially in slowdowns for incompletely sampled taxa. Hence, we could use precise sampling fractions for each lineage in future studies.

Nevertheless, simulations through our parameter estimates largely recovered the branching patterns of the empirical phylogenies (Fig. S7a, b, c, d). The estimated parameters for the simulated trees consistently fell within two standard deviations of those estimated from the empirical trees of the 31 lineages. Since these lineages varied in clade age and size, we can confidently conclude that clade age and size do not significantly affect the accuracy of our estimations. However, the empirical diversification scenarios were recovered with high confidence in only 18 lineages, while 12 had low proportions of correctly inferred scenarios. Further assessments of congruence classes using pulled diversification rates and the CRABS test indicated that many pulled diversification dynamics align with their canonical counterparts.

Our dataset comprising 729 endemic PIP species is strongly biased towards small vertebrates (∼75 %), freshwater biota, and soil arthropods, among the most underrepresented groups in meta-analyses. Notably, birds and mammals are among the most diverse megafauna globally but not in the PIP where endemicity is low (birds ∼5% and mammals ∼15%) as compared to the soil arthropods (∼90%), small vertebrates (∼52% - amphibians and reptiles) and flowering plants (∼26%) (Bharti *et al*. 2021; International Union for Conservation of Nature (IUCN) 2022; Loria & Prendini 2020). This could be because bird and mammal assemblages in the PIP are mainly dispersal-driven and have not undergone *in situ* speciation. Therefore, focusing on herpetofauna, arthropods, and plants will be more insightful in understanding the drivers of diversification processes in landscapes like the PIP.

The current diversity estimates could also be underestimated for soil arthropods, herpetofauna and freshwater biota, as many new species are still being described (Joshi & Agarwal 2021). Also, many studies have explicitly focused on the Western Ghats, the known biodiversity hotspot, while other regions of the PIP, including the savannas, remain unexplored and understudied. Even in global datasets, soil and freshwater habitats are poorly represented, urging us to focus on unexplored taxa and habitats for a holistic understanding of the tempo and mode of diversification across tropical forests. Besides, phylogenetic reconstructions using more sophisticated tree priors like the fossilised birth-death (FBD) may influence the estimates for the diversification dynamics and need to be integrated into future studies.

We show that diversification in the PIP has signatures of regional drivers like intensified mid-Miocene aridification and many known global drivers like tropical ecosystem stability, paleotemperature and existing diversity. The complex tempo and mode of diversification shown in our results suggest that regional biogeographic, geoclimatic and phylogenetic history are critical to understanding the origins of tropical diversity and need to be examined across tropical regions. Exploring spatially structured and trait-based cladogenesis among endemic lineages using heterogeneous birth-death models could shed light on the bioregionalisation within the PIP and other biomes (Helmstetter *et al*. 2023; Martínez-Gómez *et al*. 2024). Studies like this, where multi-taxa comparisons are carried out from different areas, are critical for gaining insights into the evolution of tropical biodiversity and predicting how these groups may respond to changing climatic conditions.

## Acknowledgements

We thank Dr. Anieli Pereira, Dr. Aparna Lajmi, Dr. Ashok Malik, Dr. Chinta Sidharthan, Dr. Ishan Agarwal, Dr. Ivan Bolotov, Dr. Prabha Amarasinghe, Mr. R. Chaitanya, Dr. Sandeep Sen, Mr. Saunak Pal, Dr. Siddharth Surveswaran, Dr. Stephanie Loria, Dr. Varun Torsekar, Dr. Vijay Ramesh, Dr. Vivek Cyriac, Mr. Vladislav Gorin, who shared the maximum clade credibility trees, enabling us to compile the dataset. We also thank Dr. Bharti D. K., Mr. R. Chaitanya, Mr. Abhishek Gopal, Dr. Mihir Kulkarni, Dr. Jun Ying Lim, and Dr. Rohit Naniwadekar for their suggestions on the analyses and comments on the manuscript.

## References

Agarwal, I. & Karanth, K.P. (2015). A phylogeny of the only ground-dwelling radiation of Cyrtodactylus (Squamata, Gekkonidae): diversification of Geckoella across peninsular India and Sri Lanka. Mol. Phylogenet. Evol., 82, 193–199.

Agarwal, I. & Ramakrishnan, U. (2017). A phylogeny of open-habitat lizards (Squamata: Lacertidae: *Ophisops*) supports the antiquity of Indian grassy biomes. J. Biogeogr., 44, 2021–2032.

Ali, J.R. & Aitchison, J.C. (2008). Gondwana to Asia: Plate tectonics, paleogeography and the biological connectivity of the Indian sub-continent from the Middle Jurassic through latest Eocene (166–35 Ma). Earth-Sci. Rev., 88, 145–166.

Anderson, M.J. (2017). Permutational Multivariate Analysis of Variance (PERMANOVA). In: Wiley StatsRef: Statistics Reference Online. John Wiley & Sons, Ltd, pp. 1–15.

Bacon, C.D., Silvestro, D., Hoorn, C., Bogotá-Ángel, G., Antonelli, A. & Chazot, N. (2022). The origin of modern patterns of continental diversity in Mauritiinae palms: the Neotropical museum and the Afrotropical graveyard. Biol. Lett., 18, 20220214.

Bansal, M., Morley, R.J., Nagaraju, S.K., Dutta, S., Mishra, A.K., Selveraj, J., et al. (2022). Southeast Asian Dipterocarp origin and diversification driven by Africa-India floristic interchange. Science, 375, 455–460.

Benson, R.B.J., Butler, R.J., Alroy, J., Mannion, P.D., Carrano, M.T. & Lloyd, G.T. (2016). Near-Stasis in the Long-Term Diversification of Mesozoic Tetrapods. PLOS Biol., 14, e1002359.

Bharti, D.K., Edgecombe, G.D., Karanth, K.P. & Joshi, J. (2021). Spatial patterns of phylogenetic diversity and endemism in the Western Ghats, India: A case study using ancient predatory arthropods. Ecol. Evol., 11, 16499–16513.

Bharti, D.K., Pawar, P.Y., Edgecombe, G.D. & Joshi, J. (2023). Genetic diversity varies with species traits and latitude in predatory soil arthropods (Myriapoda: Chilopoda). Glob. Ecol. Biogeogr., 32, 1508–1521.

Bonaparte, J. (1999). Tetrapod Faunas from South America and India: A Palaeobiogeographic Interpretation. Proc. Indian Natl. Sci. Acad., 65, 427–437.

Brown, J.H., Gillooly, J.F., Allen, A.P., Savage, V.M. & West, G.B. (2004). Toward a metabolic theory of ecology. Ecology, 85, 1771–1789.

Burnham, K.P. & Anderson, D.R. (2004). Multimodel Inference: Understanding AIC and BIC in Model Selection. Sociol. Methods Res., 33, 261–304.

Cardillo, M., Orme, C.D.L. & Owens, I.P.F. (2005). Testing for latitudinal bias in diversification rates: an example using new world birds. Ecology, 86, 2278–2287.

Chatterjee, S., Goswami, A. & Scotese, C.R. (2013). The longest voyage: Tectonic, magmatic, and paleoclimatic evolution of the Indian plate during its northward flight from Gondwana to Asia. Gondwana Res., 23, 238–267.

Clift, P.D. & Webb, A.A.G. (2019). A history of the Asian monsoon and its interactions with solid Earth tectonics in Cenozoic South Asia. Geol. Soc. Lond. Spec. Publ., 483, 631–652.

Cohen, J. (2013). Statistical Power Analysis for the Behavioral Sciences. 2nd edn. Routledge, New York.

Collyer, M.L. & Adams, D.C. (2018). RRPP: An r package for fitting linear models to high-dimensional data using residual randomization. Methods Ecol. Evol., 9, 1772–1779.

Condamine, F.L., Rolland, J. & Morlon, H. (2013). Macroevolutionary perspectives to environmental change. Ecol. Lett., 16, 72–85.

Condamine, F.L., Rolland, J. & Morlon, H. (2019a). Assessing the causes of diversification slowdowns: temperature-dependent and diversity-dependent models receive equivalent support. Ecol. Lett., 22, 1900–1912.

Condamine, F.L., Rolland, J. & Morlon, H. (2019b). Assessing the causes of diversification slowdowns: temperature-dependent and diversity-dependent models receive equivalent support. Ecol. Lett., 22, 1900–1912.

Couvreur, T.L.P., Pirie, M.D., Chatrou, L.W., Saunders, R.M.K., Su, Y.C.F., Richardson, J.E., et al. (2011). Early evolutionary history of the flowering plant family Annonaceae: steady diversification and boreotropical geodispersal. J. Biogeogr., 38, 664–680.

Crottini, A., Madsen, O., Poux, C., Strauß, A., Vieites, D.R. & Vences, M. (2012). Vertebrate time-tree elucidates the biogeographic pattern of a major biotic change around the K–T boundary in Madagascar. Proc. Natl. Acad. Sci., 109, 5358–5363.

Cumming, G. (2013). Understanding The New Statistics: Effect Sizes, Confidence Intervals, and Meta-Analysis. Routledge, New York.

Cyriac, V.P. & Kodandaramaiah, U. (2018). Digging their own macroevolutionary grave: fossoriality as an evolutionary dead end in snakes. J. Evol. Biol., 31, 587–598.

Cyriac, V.P., Mohan, A.V., Dinesh, K.P., Torsekar, V., Jayarajan, A., Swamy, P., et al. (2024). Diversifying in the mountains: spatiotemporal diversification of frogs in the Western Ghats biodiversity hotspot. Evolution, 78, 701–715.

Da Silva, D., Aires, A.E., Zurano, J.P., Olalla-Tárraga, M.A. & Martinez, P.A. (2020). Changing Only Slowly: The Role of Phylogenetic Niche Conservatism in Caviidae (Rodentia) Speciation. J. Mamm. Evol., 27, 713–721.

Dagallier, L.M.J., Janssens, S.B., Dauby, G., Blach-Overgaard, A., Mackinder, B.A., Droissart, V., et al. (2020). Cradles and museums of generic plant diversity across tropical Africa. New Phytol., 225, 2196–2213.

Das, I. & Chanda, S.K. (1998). A new species of Philautus (Anura: Rhacophoridae) from the Eastern Ghats, south-eastern India. J. South Asian Nat. Hist., 3, 103–112.

Deepak, V. & Karanth, P. (2018). Aridification driven diversification of fan-throated lizards from the Indian subcontinent. Mol. Phylogenet. Evol., 120, 53–62.

Derryberry, E.P., Claramunt, S., Derryberry, G., Chesser, R.T., Cracraft, J., Aleixo, A., et al. (2011). Lineage diversification and morphological evolution in a large-scale continental radiation: the neotropical ovenbirds and woodcreepers (Aves: Furnariidae). Evolution, 65, 2973–2986.

Ding, L., Spicer, R.A., Yang, J., Xu, Q., Cai, F., Li, S., et al. (2017). Quantifying the rise of the Himalaya orogen and implications for the South Asian monsoon. Geology, 45, 215–218.

Dinno, A. (2014). dunn.test: Dunn’s Test of Multiple Comparisons Using Rank Sums.

Eiserhardt, W.L., Couvreur, T.L.P. & Baker, W.J. (2017). Plant phylogeny as a window on the evolution of hyperdiversity in the tropical rainforest biome. New Phytol., 214, 1408– 1422.

Etienne, R., Haegeman, B., Hildenbrandt, H. & Laudanno, G. (2023). DDD package for R: Diversity-Dependent Diversification.

Gómez-Rodríguez, C., Baselga, A. & Wiens, J.J. (2015). Is diversification rate related to climatic niche width? Glob. Ecol. Biogeogr., 24, 383–395.

Gopal, A., Bharti, D.K., Page, N., Dexter, K.G., Krishnamani, R., Kumar, A., et al. (2023). Range restricted old and young lineages show the southern Western Ghats to be both a museum and a cradle of diversity for woody plants. Proc. R. Soc. B Biol. Sci., 290, 20222513.

Gower, D.J., Agarwal, I., Karanth, K.P., Datta-Roy, A., Giri, V.B., Wilkinson, M., et al. (2016). The role of wet-zone fragmentation in shaping biodiversity patterns in peninsular India: insights from the caecilian amphibian *Gegeneophis*. J. Biogeogr., 43, 1091–1102.

Grismer, J.L., Schulte, J.A., Alexander, A., Wagner, P., Travers, S.L., Buehler, M.D., et al. (2016). The Eurasian invasion: phylogenomic data reveal multiple Southeast Asian origins for Indian Dragon Lizards. BMC Evol. Biol., 16, 43.

Harmon, L.J., Pennell, M.W., Henao-Diaz, L.F., Rolland, J., Sipley, B.N. & Uyeda, J.C. (2021). Causes and Consequences of Apparent Timescaling Across All Estimated Evolutionary Rates. Annu. Rev. Ecol. Evol. Syst., 52, 587–609.

Helmstetter, A.J., Zenil-Ferguson, R., Sauquet, H., Otto, S.P., Méndez, M., Vallejo-Marin, M., et al. (2023). Trait-dependent diversification in angiosperms: Patterns, models and data. Ecol. Lett., 26, 640–657.

Helwig, N.E. (2020). Multiple and Generalized Nonparametric Regression. SAGE Publications Ltd.

Henao Diaz, L.F., Harmon, L.J., Sugawara, M.T.C., Miller, E.T. Pennell, M.W. (2019). Macroevolutionary diversification rates show time dependency. Proc. Natl. Acad. Sci., 116, 7403–7408.

Hernández-Hernández, T., Miller, E.C., Román-Palacios, C. & Wiens, J.J. (2021). Speciation across the Tree of Life. Biol. Rev., 96, 1205–1242.

Heslop-Harrison, J.S. (Pat), Schwarzacher, T. & Liu, Q. (2023). Polyploidy: its consequences and enabling role in plant diversification and evolution. Ann. Bot., 131, 1–10.

Höhna, S., Kopperud, B.T. & Magee, A.F. (2022). CRABS: Congruent rate analyses in birth–death scenarios. Methods Ecol. Evol., 13, 2709–2718.

Hohna, S., May, M.R. & Moore, B.R. (2015). Phylogeny Simulation and Diversification Rate Analysis with TESS.

Höhna, S., May, M.R. & Moore, B.R. (2016). TESS: an R package for efficiently simulating phylogenetic trees and performing Bayesian inference of lineage diversification rates. Bioinformatics, 32, 789–791.

International Union for Conservation of Nature (IUCN). (2022). The IUCN Red List of Threatened Species, Version 2022-2.

Irnawati, I., Riswanto, F.D.O., Riyanto, S., Martono, S. & Rohman, A. (2021). The use of software packages of R factoextra and FactoMineR and their application in principal component analysis for authentication of oils. Indones. J. Chemom. Pharm. Anal., 1–10.

Jeffreys, H. (1998). The Theory of Probability. OUP Oxford.

Joshi, J. & Agarwal, I. (2021). Integrative Taxonomy in the Indian Subcontinent: Current Progress and Prospects. J. Indian Inst. Sci., 101, 125–149.

Joshi, J. & Edgecombe, G.D. (2019). Evolutionary biogeography of the centipede genus Ethmostigmus from Peninsular India: testing an ancient vicariance hypothesis for Old World tropical diversity. BMC Evol. Biol., 19, 1–1.

Joshi, J. & Karanth, K.P. (2011). Cretaceous–Tertiary diversification among select Scolopendrid centipedes of South India. Mol. Phylogenet. Evol., 60, 287–294.

Joshi, J. & Karanth, P. (2013). Did southern Western Ghats of peninsular India serve as refugia for its endemic biota during the Cretaceous volcanism? Ecol. Evol., 3275–3282.

Joshi, J., Karanth, P.K. & Edgecombe, G.D. (2020). The out-of-India hypothesis: evidence from an ancient centipede genus, Rhysida (Chilopoda: Scolopendromorpha) from the Oriental Region, and systematics of Indian species. Zool. J. Linn. Soc., 189, 828–861.

Karanth, K.P. (2003). Evolution of disjunct distributions among wet-zone species of the Indian subcontinent: Testing various hypotheses using a phylogenetic approach. Curr. Sci., 85.

Karanth, K.P. (2015). An island called India: phylogenetic patterns across multiple taxonomic groups reveal endemic radiations. Curr. Sci., 108, 1847–1851.

Karanth, K.P. (2021). Dispersal vs. vicariance: the origin of India’s extant tetrapod fauna. Front. Biogeogr., 13.

Keller, G., Sahni, A. & Bajpai, S. (2009). Deccan volcanism, the KT mass extinction and dinosaurs. J. Biosci., 34, 709–728.

Klaus, S., Morley, R.J., Plath, M., Zhang, Y.-P. & Li, J.-T. (2016). Biotic interchange between the Indian subcontinent and mainland Asia through time. Nat. Commun., 7, 12132.

Kopperud, B.T., Magee, A.F. & Höhna, S. (2023). Rapidly changing speciation and extinction rates can be inferred in spite of nonidentifiability. Proc. Natl. Acad. Sci., 120, e2208851120.

Kumar, S., Stecher, G., Suleski, M. & Hedges, S.B. (2017). TimeTree: A Resource for Timelines, Timetrees, and Divergence Times. Mol. Biol. Evol., 34, 1812–1819.

Lajmi, A. & Karanth, P.K. (2020). Eocene–Oligocene cooling and the diversification of Hemidactylus geckos in Peninsular India. Mol. Phylogenet. Evol., 142, 106637.

Lê, S., Josse, J. & Husson, F. (2008). FactoMineR: An R Package for Multivariate Analysis. J. Stat. Softw., 25, 1–18.

Li, F., Shao, L. & Li, S. (2020). Tropical Niche Conservatism Explains the Eocene Migration from India to Southeast Asia in Ochyroceratid Spiders. Syst. Biol., 69, 987–998.

Liu, X., Liu, Y., Wu, R., Zanatta, D.T., Lopes-Lima, M., Gonçalves, D.V., et al. (2022). Systematics, distribution, biology, and conservation of freshwater mussels (Bivalvia: Unionida) in China. Aquat. Conserv. Mar. Freshw. Ecosyst., 32, 859–895.

Loria, S.F. & Prendini, L. (2020). OPEN Out of India, thrice: diversification of Asian forest scorpions reveals three colonizations of Southeast Asia. Sci. Rep., 10, 22301.

Losos, J.B. & Ricklefs, R.E. (2009). Adaptation and diversification on islands. Nature, 457, 830– 836.

Louca, S. (2017). castor: Efficient Phylogenetics on Large Trees.

Louca, S. & Pennell, M.W. (2020). Extant timetrees are consistent with a myriad of diversification histories. Nature, 580, 502–505.

Maliet, O. & Morlon, H. (2022). Fast and Accurate Estimation of Species-Specific Diversification Rates Using Data Augmentation. Syst. Biol., 71, 353–366.

Mani, M.S. (1974). Biogeographical Evolution in India. In: Ecology and Biogeography in India, Monographiae Biologicae (ed. Mani, M.S.). Springer Netherlands, Dordrecht, pp. 698–724.

Manivannan, M., Gurung, N., Edgecombe, G.D. & Joshi, J. (2024). A passage through India: The biotic ferry model supports the build-up of Indo-Australian biodiversity of an ancient soil arthropod clade. J. Biogeogr., 51, 2395–2411.

Martínez-Gómez, J., Song, M.J., Tribble, C.M., Kopperud, B.T., Freyman, W.A., Höhna, S., et al. (2024). Commonly used Bayesian diversification methods lead to biologically meaningful differences in branch-specific rates on empirical phylogenies. Evol. Lett., 8, 189–199.

May, M.R., Höhna, S. & Moore, B.R. (2016). A Bayesian approach for detecting the impact of mass-extinction events on molecular phylogenies when rates of lineage diversification may vary. Methods Ecol. Evol., 7, 947–959.

McPeek, M.A. & Brown, J.M. (2007). Clade Age and Not Diversification Rate Explains Species Richness among Animal Taxa. Am. Nat., 169, E97–E106.

Meseguer, A.S., Michel, A., Fabre, P.-H., Pérez Escobar, O.A., Chomicki, G., Riina, R., et al. (2022). Diversification dynamics in the Neotropics through time, clades, and biogeographic regions. eLife, 11, e74503.

Moen, D. & Morlon, H. (2014). Why does diversification slow down? Trends Ecol. Evol., 29, 190– 197.

Moreira, M.O., Wiens, J.J., Fonseca, C. & Rojas, D. (2024). Climatic-niche breadth, niche position, and speciation in lizards and snakes. J. Biogeogr., 51, 969–981.

Morley, R.J. (2018). Assembly and division of the South and South-East Asian flora in relation to tectonics and climate change. J. Trop. Ecol., 34, 209–234.

Morlon, H. (2014). Phylogenetic approaches for studying diversification. Ecol. Lett., 17, 508–525.

Morlon, H., Andréoletti, J., Barido-Sottani, J., Lambert, S., Perez-Lamarque, B., Quintero, I., et al. (2024). Phylogenetic Insights into Diversification. Annu. Rev. Ecol. Evol. Syst., 55, 1–21.

Morlon, H., Lewitus, E., Condamine, F.L., Manceau, M., Clavel, J. & Drury, J. (2016). RPANDA1: an R package for macroevolutionary analyses on phylogenetic trees. Methods Ecol. Evol., 7, 589–597.

Morlon, H., Potts, M.D. & Plotkin, J.B. (2010). Inferring the Dynamics of Diversification: A Coalescent Approach. PLoS Biol., 8, e1000493.

O’Meara, B.C. & Beaulieu, J.M. (2024). Noise leads to the perceived increase in evolutionary rates over short time scales. PLOS Comput. Biol., 20, e1012458.

Pagès, J. (2004). Analyse factorielle de donnees mixtes: principe et exemple d’application. Rev. Stat. Appliquée, 52, 93–111.

Parmar, S., Morley, R.J., Bansal, M., Singh, B.P., Morley, H. & Prasad, V. (2023). Evolution of family Arecaceae on the Indian Plate modulated by the Early Palaeogene climate and tectonics. Rev. Palaeobot. Palynol., 313, 104890.

Pepper, M. & Keogh, J.S. (2021). Life in the “dead heart” of Australia: The geohistory of the Australian deserts and its impact on genetic diversity of arid zone lizards. J. Biogeogr., 48, 716–746.

Prasad, V., Farooqui, A., Tripathi, S.K.M., Garg, R. & Thakur, B. (2009). Evidence of Late Palaeocene-Early Eocene equatorial rain forest refugia in southern Western Ghats, India. J. Biosci., 34, 777–797.

Quade, J., Cerling, T.E. & Bowman, J.R. (1989). Development of Asian monsoon revealed by marked ecological shift during the latest Miocene in northern Pakistan. Nature, 342, 163–166.

Rabosky, D.L. (2009). Ecological limits and diversification rate: alternative paradigms to explain the variation in species richness among clades and regions. Ecol. Lett., 12, 735–743.

Rabosky, D.L. (2013). Diversity-Dependence, Ecological Speciation, and the Role of Competition in Macroevolution. Annu. Rev. Ecol. Evol. Syst., 44, 481–502.

Román-Palacios, C., Moraga-López, D. & Wiens, J.J. (2022). The origins of global biodiversity on land, sea and freshwater. Ecol. Lett., 25, 1376–1386.

Scholl, J.P. & Wiens, J.J. (2016). Diversification rates and species richness across the Tree of Life. Proc. R. Soc. B Biol. Sci., 283, 20161334.

Sidharthan, C. & Karanth, K.P. (2021). India’s biogeographic history through the eyes of blindsnakes-filling the gaps in the global typhlopoid phylogeny. Mol. Phylogenet. Evol., 157, 107064.

Spencer, R.C. (1940). Properties of the Witch of Agnesi—Application to Fitting the Shapes of Spectral Lines. J. Opt. Soc. Am., 30, 415–419.

Suárez, D., Arribas, P., Jiménez-García, E. & Emerson, B.C. (2022). Dispersal ability and its consequences for population genetic differentiation and diversification. Proc. R. Soc. B Biol. Sci., 289, 20220489.

Surveswaran, S., Kambale, S.S., Srivastav, M., Punekar, S.A., Yadav, S.R. & Karanth, K.P. (2021). Origin and diversification of Indian Ceropegieae (Apocynaceae) and its possible relation to the Indian monsoon. J. Syst. Evol., 59, 93–112.

Thompson, J.B. & Ramírez-Barahona, S. (2023). No phylogenetic evidence for angiosperm mass extinction at the Cretaceous–Palaeogene (K-Pg) boundary. Biol. Lett., 19, 20230314.

Tietje, M., Antonelli, A., Baker, W.J., Govaerts, R., Smith, S.A. & Eiserhardt, W.L. (2022). Global variation in diversification rate and species richness are unlinked in plants. Proc. Natl. Acad. Sci., 119, e2120662119.

Tripathy, B. & Mukhopadhayay, A. (2015). Freshwater Molluscs of India: An Insight of into Their Diversity, Distribution and Conservation. In: Aquatic Ecosystem: Biodiversity, Ecology and Conservation (eds. Rawat, M., Dookia, S. & Sivaperuman, C.). Springer India, New Delhi, pp. 163–195.

Vasconcellos, M.M., Colli, G.R. & Cannatella, D.C. (2021). Paleotemperatures and recurrent habitat shifts drive diversification of treefrogs across distinct biodiversity hotspots in sub-Amazonian South America. J. Biogeogr., 48, 305–320.

Wallace, A.R. (1887). Oceanic Islands: Their Physical and Biological Relations. J. Am. Geogr. Soc. N. Y., 19, 1.

Wiens, J.J. (2004). Speciation and ecology revisited: Phylogenetic niche conservatism and the origin of species. Evolution, 58, 193–197.

Wiens, J.J. (2011). The Causes Of Species Richness Patterns Across Space, Time, And Clades And The Role Of “Ecological Limits.” Q. Rev. Biol., 86, 75–96.

Willig, M.R., Kaufman, D.M. & Stevens, R.D. (2003). Latitudinal Gradients of Biodiversity: Pattern, Process, Scale, and Synthesis. Annu. Rev. Ecol. Evol. Syst., 34, 273–309.

Yuan, Z.-Y., Zhang, B.-L., Raxworthy, C.J., Weisrock, D.W., Hime, P.M., Jin, J.-Q., et al. (2019). Natatanuran frogs used the Indian Plate to step-stone disperse and radiate across the Indian Ocean. Natl. Sci. Rev., 6, 10–14.

Zachos, J.C., Dickens, G.R. & Zeebe, R.E. (2008). An early Cenozoic perspective on greenhouse warming and carbon-cycle dynamics. Nature, 451, 279–283.

