## Supplementary figures and tables for "Idiosyncrasies unveiled: examining the pace, patterns and predictors of biotic diversification in peninsular India"

**This PDF includes:**

Detailed Methodology

SI Tables 1 to 7

SI Figures 1 to 7

### 1 Detailed Methodology

#### 1.1 Assessing diversification rates

Speciation, extinction and net-diversification rates were calculated to assess the pace in diversification of lineages. Clade-specific and species-specific rates were calculated and compared using multiple analyses of variance across taxonomic groups, biogeographic origins and habitat types. Three distinct approaches were employed in calculating these rates.

Firstly, we defined a constant-rate birth-death model and the parameters were optimsed in a maximum-likelihood framework of the R package *RPANDA* [1]. Specifically, the birth (i.e. speciation) model needed two parameters - t (indicating time) and $\lambda_0$(speciation rate at the present as per the CBD model), however, time was excluded in the mathematical construct of the function and only the optimsed value of $\lambda_0$ would be returned. This single optimised value which is constant through time, can essentially be equivalent to the mean speciation rate across time and species for a lineage. Similary, an optimsed value for the mean extinction rate (which is also equal to the extinction rate at present - $\mu_0$) was calculated. The difference between the mean speciation and extinction rates was then calculated to infer the mean net-diversification rate for each lineage. Distinct model-fitting-optimisation exercises were performed for each of the 33 phylogenies (i.e. lineages) to obtain the relevant diversification rates.

Secondly, we performed the CoMET (Compound poisson process on Mass ExTinction) analysis on each of the 33 lineages using the R package *TESS* [2]. The CoMET analysis involves binning the history of a clade into 100 time points at equal intervals. We explicitly estimated empirical hyperpriors and estimation of mass extinctions was disabled. Initially, The analysis was ran by setting 1000 MCMC iterations. Following this, we checked effective sample size (ESS) for speciation and extinction rate in each time bin. If the ESS had not reached 200 for at least one time bin, then the analysis was re-run with five times the previous number of generations. It was looped until all the time bins showed a minimum ESS of 200 for speciation and extinction rates. Next, from the posterior distribution, speciation and extinction rates pertaining to each time bin were extracted. Furthermore, the mean speciation and extinction rate of a lineage was calculated from the respective distributions. The difference between these mean rates further provided the mean net-diversification rates for all lineages.

Thirdly, we used the ClaDS (Cladogenetic Diversification rate Shift) model to estimate branch-specific and tip-specific (i.e. species-specific) diversification rates (speciation and extinction rates) for each lineage. This analysis was run on relevant well-sampled larger phylogenies (super trees) to which each lineage belonged (details of the super trees are provided in Table S2 and can be found in the deposited data in the Zenodo repository) and was performed using the *infer_ClaDS* function from the Julia package *PANDA* [3]. A default setting of 3 MCMC chains was used and the analysis continued for as many iterations as it would take to converge. Convergence was assessed using the gelman statistic and the run would stop when the value reached a value below 1.05. Further, the output for each lineage was saved in ".Rdata" format and the relevant output variables were loaded in R for further analyses. Branch and tip rates for relevant sub-clades (pertaining to each of the 33 lineages) were extracted from the loaded "CladsOutput" object for the super trees and mean speciation, extinction and net-diversification rates were calculated from these relevant rates. This method was employed to obtain more robust estimates, particularly for a few depauperate clades.

For comparing diversification rates across taxonomic groups, biogeographic origins and habitat types, we performed two analyses of variance - 1) the non-parametric Kruskal-Wallis test and 2) phylogenetic ANOVA, to additionally control for phylogenetic non-independence.

The Kruskal-Wallis test was chosen over standard ANOVA because the distribution of diversification rates (estimated using different methods) failed the test for normality - the Shapiro-Wilk test ($p < 0.05$ - indicating it is significantly different from a normal distribution), which was performed using the *shapiro.test* function in R [4]. The *kruskal.test* function was used to perform the Kruskal-Wallis test and the relevant post hoc analysis - Dunn's test, applying Bonferroni corrections, was also performed using the *dunn.test* function from *dunn.test* package in R [4, 5]. Separate analyses were performed on diversification rates estimated through different methods. Besides clade-rates (i.e. mean rates of speciation, extinction and net-diversification for each clade/lineage), tip-rates estimated using the ClaDS model were compared as well.

Phylogenetic ANOVA was performed using the *lm.rrpp* function from the R package - *RRPP* [6]. This analysis required a phylogenetic tree linking all the 33 lineages. Hence, we constructed a dated phylogeny using information from timetree.org and from phylogenies of the relevant lineages (Fig. S3.e - upper panel) [7]. This tree was further converted to a variance-covariance matrix using the *vcv.phylo* function from the *ape* package in R, which was used in the regression to control for phylogenetic relatedness [8]. Furthermore, post hoc comparisons were performed using the *pairwise* function from the *RRPP* package in R to obtain corrected $p$-values for comparing rates across categories within each predictor (taxonomic group, biogeographic origin, habitat type). Both clade-rates and tip-rates (from ClaDS results) were compared using this method. A larger dated phylogeny was constructed by imputing the individual phylogeny of each lineage to the tree generated earlier (that linked the lineages) by manually editing its newick file. This tree was used to particularly perform phylogenetic ANOVA on the tip-rates (Fig. S3.e - lower panel). Phylogenetic signal in each association was assessed by calcuting the Pagel's $\lambda$ [9] for the residuals in each dataset using the *phylosig* function from the *phytools* package in R [10]. from Fig. S3.e .

Lastly, we tested if clade ages could predict diversification rates by performing a phylogenetic generalised least squares regression (PGLS) using the clade-rates (ClaDS, CoMET, *RPANDA*) and the respective clade-ages. PGLS was performed using the *pgls* function from the *caper* package in R [11]. The earlier mentioned dated phylogeny linking all the lineages (Fig. S3.e - upper panel) was used in this analysis and the estimation for Pagel's $\lambda$ was enabled.

Boxplots for the comparisons were made using the *ggbetweenstats* function from the *ggstatsplot* package in R [12] and the scatter plots representating the results of PGLS were made using the functions from the *ggplot2* package [13].

#### 1.2 Assessing diversification patterns

The temporal mode of diversification was evaluated by first fitting homogeneous birth-death models (constant-rate and time-varying) to all 33 lineages. For this class of birth-death models, speciation (birth) and extinction (death) rates remain homogeneous across the branches of a phylogeny at one time point but can vary across time. Constant-rate models included constant-rate pure birth (CB - only a constant speciation rate across time, no extinction) and constant-rate birth and death models (CBD - constant and finite speciation and extinction rates across time) [14, 15]. Time-varying models included combinations of linearly ($\lambda$(t)=$\lambda_0$+$\alpha$t) or exponentially ($\lambda$(t)=$\lambda_0\cdot e^{\alpha t}$) varying (with time - t) birth ($\lambda$) and death ($\mu$) rates ($\lambda_0$ and $\mu_0$ are the expected speciation and extinction rates at t=0, $\alpha$ and $\beta$ are the coefficients of gain or decay in the functions and time, here, is interpreted as time from the present) [14, 15]. These combinations also included constant-rate models for scenarios where one rate (birth or death) varies with time, and the other remains constant. In the models above, the absolute values of speciation and extinction rates are considered even when the models infer negative values. The negative values for speciation and extinction rates are not biologically informative but are feasible mathematically. Therefore, we followed the approach described by [16] to include a set of models with the same mathematical formulations where we take a value of 0 in cases where rates become negative and refer to them as "non-negative models".

Additionally, to model the possibility of symmetrical local peaks and dips (sharp increase followed by a decrease in diversification rates or vice versa) in diversification rates across time, we developed a continuous time-varying mathematical function to model symmetrical local peaks in diversification and to perform model fitting in the same framework. This function follows the "Witch of Agnesi" curve, where the diversification rates diverge from a baseline constant rate: $\lambda\left(t\right) = \lambda_0 + \frac{\alpha\cdot\omega}{\mathrm{p}}\cdot\left(\frac{\mathrm{t}}{(\mathrm{t}\text{-}\mathrm{p})^2+\omega}\right)$, where $\lambda_0$ represents a baseline rate of speciation (which is also the rate at present), $\alpha$ represents the intensity of the peak (or dip), represents the spread of the peak or dip, and p represents the position of the peak (or dip) in My [17]. Similar rate shifts in extinction were also modelled. This helped examine the prevalence of symmetrical local peaks (or dips) in diversification. Additionally, using this function, we qualitatively assessed the role of specific geoclimatic events (highlighted in Fig. 1) on the diversification of lineages corresponding to the time of occurrence of a peak (or dip). We called this model the "symmetrical local shift"/"symmetrical episodic shift" (abbreviated as SES) model and implemented it in the *RPANDA* model-fitting framework. Model fitting using constant-rate and time-varying birth-death models was executed in *RPANDA* using the *fit_bd* function to determine the diversification pattern of each lineage .

Best-fitting models were selected based on AICc scores to account for over-fitting. ΔAICc scores were calculated as absolute differences in the AICc scores between each model and the model with the lowest AICc score. The models with ΔAICc scores lesser than two were chosen for each lineage following [18]. If multiple models showed ΔAICc below two, i.e., all models were equally likely, then the model with the least parameter complexity was chosen as the best-fitting model. If multiple models of similar AICc and the same complexity existed, the model with the least AICc was selected as the best-fitting model. However, if CB or CBD were found along with other models to be equally likely, the corresponding constant-rate model was considered the best-fitting model for a lineage, as the simplest model could not be rejected. We used the parameter values ($\lambda_0$, $\alpha$, $\mu_0$ and $\beta$) of the selected diversification models to evaluate the scenarios mentioned in Table S2 and Fig. 1, Fig. S1.

In addition, we used the results from the CoMET (CPP on Mass-Extinction Times) analysis [19, 20] mentioned earlier to detect times of significant rate shifts in a lineage's history and generate rates-through-time (RTT) plots. In the CoMET analysis, the lineage history is split into 100 time bins and using a Bayesian framework the speciation and extinction rates for each time bin are calculated. Bayes Factor supports providing the statistical strength in assessing if episodic rate shift happened in a time bin relative to its previous time bin are also calculated for each time bin. For each lineage, we extracted the Bayes Factor (BF) supports for rate shifts for all the 100 time points and chose the ones where 2 * ln(BF) scores were substantially higher than that of constant rates ($\geq$4.6) [19, 21]. The corresponding time bins were then considered as the times of episodic rate shifts in speciation and extinction rates.

#### 1.3 Credibility of estimates

To assess the reliability of our diversification rate estimates, we performed simulations using the best-fit parameters from constant and time-varying models for each lineage. We calculated errors in model parameters, the timing of significant rate shifts, and diversification scenarios. We performed these tests because our dataset was dominated by depauperate clades and small phylogenies can affect the robustness of diversification rate estimates. We simulated 100 trees per lineage using the best model parameter estimates conditioned on crown ages. We conducted the simulations with the *tess.sim.age* function in the R package *TESS*, ensuring that each tree retained the same clade size and age while allowing branching patterns to vary according to the model parameters. We then analysed these simulated trees using the same methods as the empirical phylogenies (*RPANDA* models and CoMET).

We first assessed the diversification scenarios of the simulated phylogenies and compared them with empirical phylogenies to determine the proportion of correctly estimated scenarios. This assessment was conducted using two methods: RPANDA and CoMET. The RPANDA framework, along with the SES model (developed by us), enabled comparison across all scenarios while also detecting episodic rate shifts. In this approach, for a set of 100 simulated trees generated using the best-fit model parameters of an empirical tree, if 80 trees matched the empirical scenario, the proportion of correctness would be 0.8. For the CoMET analysis, scenario correctness was determined by comparing the presence or absence of episodic rate shifts in simulated trees relative to empirical trees, as CoMET only provides the timing of such shifts. If an empirical tree lacked an episodic rate shift, and in a corresponding set of 100 simulated trees, 20 exhibited at least one episodic rate shift, the proportion of correctness would also be 0.8.

The error in model parameter estimation (speciation and extinction rates at present— $\lambda_0$ and $\mu_0$, and the slopes of speciation and extinction rate functions over time— $\alpha$ and $\beta$) was calculated using a method analogous to Cohen's D [22, 23]. For each parameter, we subtracted the values of the 100 simulated trees from the empirical tree and normalised each difference by the standard deviation of the parameter across the simulations. Finally, we estimated the timing of significant diversification rate shifts in the simulated trees using the SES model and CoMET analysis and compared them with empirical estimates.

### 2 Tables

| | | |
|---|---|---|
| **Supplementary Tables** | | |
| **MS Title:** | Idiosyncrasies unveiled: examining the pace, patterns and predictors of biotic diversification in peninsular India | |
| **Authors:** | Pragyadeep Roy* | Jahnavi Joshi* |
| **Affiliations** | Academy of Scientific and Innovative Research (AcSIR), Ghaziabad, India | Academy of Scientific and Innovative Research (AcSIR), Ghaziabad, India |
| | CSIR-Centre for Cellular and Molecular Biology, Uppal Road, Hyderabad, India | CSIR-Centre for Cellular and Molecular Biology, Uppal Road, Hyderabad, India |
| **Email IDs for correspondence*:** | | |
| **Table legends** | | |
| **Table S1** | **page 8** | **The dataset used.** The 33 lineages and their taxonomic identities, constitution, ages, species richness, biogeographic affinities and habitat preferences and information about the phylogenetic trees used, estimates for their diversification rates, the diversification scenarios they follow and the relevant drivers of diversification. Trees used for the ClaDS analyses were either the same trees used for RPANDA models and CoMET (as only focal clades were thoroughly sampled) or were well-sampled larger clades that relevant clades of interest were part of. |
| **Table S2** | **page 9** | **Analysis of variance and regression.** Kruskal-Wallis and Dunn test (post hoc test) were performed to check if there is significant structure in the diversification rates across taxonomic groups, biogeographic origins and habitat types. Additionally, it order to account for phylogenetic non-independence (as indicated by the Pagel's λ values), we performed phylogenetic ANOVA. Diversification rates estimated using three methods (RPANDA, CoMET and ClaDS) were used in both the analyses of variance. To check the effect of clade ages on the diversification rates, we performed linear and phylogenetically generalised least squares regression. Relevants statistical information for the analyses of variance and regression have been provided. Significant *p*-values have been highlighted in bold |
| **Table S3** | **page 10** | **Criteria for parameters of time-varying birth-death models corresponding to each diversification scenario assessed in the study.** $\alpha$ and $\beta$ indicates the coefficient of gain or loss with time used in the birth and death models respectively. $\lambda 0$ and $\mu 0$ are the speciation (birth) and extinction (death) rates at present time (t=0). Additional remarks define unique intricacies within the corresponding sub-scenarios; see the SI figure 1 for detailed scenarios and predictions. |
| **Table S4** | **page 11** | **Parameter estimates for constant and time-varying models (CDR and PDR).** Estimates for the times of significant episodic rates shift inferred by CoMET and the parameters for constant-rate and time-varying (only the best-fitting time-varying model) birth-death models (RPANDA), their absolute and relative fits (AICc and delta-AICC) and the best-fitting model among all, ultimately to infer the diversification scenarios that the 34 lineages followed. CB.aicc, CBD.aicc and Time.aicc refer to the AICc scores of the constant-rate birth, constant-rate birth-death and the best-fit time-varying models, respectively. Similar prefixes have been used before parameter names in the columns to denote the relevance of the parameter to the respective models. Examples of names for time-varying models as "b.lin.max_d.exp.abs" translates to a combination of linear birth model that takes the "non-negative" formulation (refer to main text methods - section 2.2) and a exponential death model that takes absolute values. Examples of names for time-varying models incorporating symmetrical rate shifts as "b.epi.var.8_d.epi.fix.15" translates to a combination of birth model with an episodic rate shift at 8 Mya with varying width of peak or dip and a death model with an episodic rate shift at 15 Mya with fixed (default width parameter = 1) width of peak or dip. |
| **Table S5** | **page 12** | **Drivers of diversification.** AICc scores, parameter estimates of the constant-rate and different factor-dependent models used for inferring the best driver(s) explaining the diversification regimes of the lineages. Factors used in the study include paleo-environmental variables (Temp - Global paleotemperature, Him.oro - Reconstructed Elevations of the Himalayan orogen and C4-Exp - pedogenic Carbon-content indicating expansion of C4 plants with intensified aridification) and existing species diversity through time (DDD - Diversity Dependent Diversification). Lowest AICc scores among all and the relative fits (delta.AICc - ΔAICc) of the aforementioned birth-death models were calculated to infer the best driver(s) of diversification for each of the 34 lineages in the dataset. |
| **Table S6** | **page 13** | **Input matrix for FAMD.** The matrix used for the FAMD analysis included information about the identity of the lineages, the taxonomic group each belonged to, their clade ages, species richness, net-diversification rates (ClaDS), biogeographic origins, habitat types, diversification scenarios they followed, and whether they showed strong support for diversity-dependent, temperature-dependent and Himalayan-orogeny dependent diversification |
| **Table S7** | **page 14** | **PermANOVA results.** The fit and the significance of differences between clusters corresponding to sub-categories within each qualitative variable (diversification scenarios, taxonomic groups, habitat types, etc.) used in the FAMD analysis. Adjusted *p*-values indicate the significance in distinctness of two clusters. Standard PermANOVA was performed on a distance (Gower's distance) matrix utilising the FAMD coordinates. Additionally we weighted this distance matrix by the phylogenetic distance matrix of the lineages generated from the dated phylogenetic tree linking all the lineages, and re-performed PermANOVA on the weighted matrix. |
| **N.B. The contents might not be visible at the first look, but please zoom them and they would be legible.** | | |

| Lineage | Reference | Taxonomic group | Taxonomic hierarchy | Details on the taxonomic scale of a clade | Details of super-tree used for ClaDS analysis from original study | Crown age (Mya) | No. of species (Clade size of lineage of interest) | No. of endemic species | Biogeographic origin | Habitat Type | Tree file description |
|---|---|---|---|---|---|---|---|---|---|---|---|
| Acanthaceae | **Surveswaran, S.**, Tiwari, N., Karanth, P. K., Deshmukh, P. V. & Lekhak, M. M. Molecular phylogenetics and character evolution in Haplanthodes (Acanthaceae), an endemic genus from peninsular India. Nord. J. Bot. 2022, njb.03238 (2022). | Angiosperm | Plantae -> Angiospermae -> Lamiales -> Acanthaceae | Clade comprising multiple species of 3 genera: *Andrographis, Haplanthodes, Haplanthus* | Size: 15 (Fig. 4 of original study) | 8.81 | 15 | 15 | Asian | Wet + Dry | recieved from author |
| Ahaetullinae | **Mallik, A. K.** et al. Discovery of a deeply divergent new lineage of vine snake (Colubridae: Ahaetuliinae: Proahaetulla gen. nov.) from the southern Western Ghats of Peninsular India with a revised key for Ahaetuliinae. PLOS ONE 14, e0218851 (2019). | Herpetofauna | Animalia -> Chordata -> Reptilia- > Squamata -> Colubridae | Clade comprising 2 entire genera: *Ahaetulla, Proahaetulla* | Size: 13 (Fig. 1 of original study) | 26.57 | 13 | 10 | Asian | Wet + Dry | recieved from author |
| Ceropegiae | **Surveswaran, S.** et al. Origin and diversification of Indian Ceropegieae (Apocynaceae) and its possible relation to the Indian monsoon. J. Syst. Evol. 59, 93–112 (2021). | Angiosperm | Plantae -> Angiospermae -> Gentianales -> Apocynaceae | Clade comprising multiple species from 2 genera: *Ceropegia, Brachystelma* | Size: 67 (Fig. 5 of original study) | 9.69 | 67 | 65 | Asian | Wet | recieved from author |
| *Cnemaspis* | **Pal, S.** et al. Diversifying on the Ark: multiple new endemic lineages of dwarf geckos from the Western Ghats provide insights into the systematics and biogeography of South Asian Cnemaspis (Reptilia: Squamata). Zool. Res. 42, 675–691 (2021). | Herpetofauna | Animalia -> Chordata -> Reptilia- > Squamata -> Gekkonidae | Clade comprising 1 entire genus: *Cnemaspis* | Size: 83 (Fig. S2 of original study) | 63.43 | 83 | 83 | Gondwanan | Wet + Dry | recieved from author |
| *Digitipes* | **Joshi, J.**, Karanth, P. K. & Edgecombe, G. D. The out-of-India hypothesis: evidence from an ancient centipede genus, Rhysida (Chilopoda: Scolopendromorpha) from the Oriental Region, and systematics of Indian species. Zool. J. Linn. Soc. 189, 828–861 (2020). | Invertebrate | Animalia -> Arthropoda -> Myriapoda -> Chilopoda -> Scolopendridae | Clade comprising 1 entire genus: *Digitipes* | Size: 66 (Fig. 4 of original study) | 148.66 | 11 | 9 | Gondwanan | Wet | recieved from author |
| Dravidogecko | **Chaitanya, R.** et al. Diversification in the mountains: a generic reappraisal of the Western Ghats endemic gecko genus Dravidogecko Smith, 1933 (Squamata: Gekkonidae) with descriptions of six new species. Zootaxa 4688, (2019). | Herpetofauna | Animalia -> Chordata -> Reptilia- > Squamata -> Gekkonidae | Clade comprising 1 entire genus: *Dravidogecko* | Size: 7 (Fig. 3 of original study) | 19.68 | 7 | 7 | Asian | Wet | recieved from author |
| Ethmostigmus | **Joshi, J.** & Edgecombe, G. D. Evolutionary biogeography of the centipede genus Ethmostigmus from Peninsular India: testing an ancient vicariance hypothesis for Old World tropical diversity. BMC Evol. Biol. 19, 41 (2019). | Invertebrate | Animalia -> Arthropoda -> Myriapoda -> Chilopoda -> Scolopendridae | Clade comprising multiple species of 1 genus: *Ethmostigmus* | Size: 66 (Fig. 4 of original study) | 65.21 | 5 | 5 | Gondwanan | Wet | recieved from author |
| *Eutropis* | **Barley, A. J.**, Datta-Roy, A., Karanth, K. P. & Brown, R. M. Sun skink diversification across the Indian-Southeast Asian biogeographical interface. J. Biogeogr. 42, 292–304 (2015). | Herpetofauna | Animalia -> Chordata -> Reptilia- > Squamata -> Scincidae | Clade comprising multiple species of 1 genus: *Eutropis* | Size: 12 (Fig. 3 of original study) | 33.57 | 12 | 12 | Asian | Wet + Dry | recieved from author |
| Fan-Throated | **Deepak, V.** & Karanth, P. Aridification driven diversification of fan-throated lizards from the Indian subcontinent. Mol. Phylogenet. Evol. 120, 53–62 (2018). | Herpetofauna | Animalia -> Chordata -> Reptilia- > Squamata -> Agamidae | Clade comprising 3 entire genera: *Sitana, Sarada, Otocryptis* | Size: 17 (Fig. 4 of original study) | 30.05 | 17 | 14 | Asian | Wet + Dry | wrote newick using date estimates |
| *Geckoella* | **Agarwal, I.** & Karanth, K. P. A phylogeny of the only ground-dwelling radiation of Cyrtodactylus (Squamata, Gekkonidae): diversification of Geckoella across peninsular India and Sri Lanka. Mol. Phylogenet. Evol. 82, 193–199 (2015). | Herpetofauna | Animalia -> Chordata -> Reptilia- > Squamata -> Gekkonidae | Clade comprising 1 entire genus: *Geckoella* | Size: 14 (Fig. 2 of original study) | 32 | 14 | 14 | Asian | Wet + Dry | wrote newick using date estimates |
| *Gegeneophis* | Gower, D. J., **Agarwal I.**, et al. The role of wet-zone fragmentation in shaping biodiversity patterns in peninsular India: insights from the caecilian amphibian Gegeneophis. J. Biogeogr. 43, 1091–1102 (2016). | Herpetofauna | Animalia -> Chordata -> Amphibia - > Gymphiona -> Grandisoniidae | Clade comprising 1 entire genus: *Gegeneophis* | Size: 14 (Fig. 3 of original study) | 37.5 | 14 | 14 | Gondwanan | Wet | wrote newick using date estimates |
| *Gerrhopilus* | **Sidharthan, C.** & Karanth, K. P. India's biogeographic history through the eyes of blindsnakes- filling the gaps in the global typhlopoid phylogeny. Mol. Phylogenet. Evol. 157, 107064 (2021). | Herpetofauna | Animalia -> Chordata -> Reptilia- > Squamata -> Gerrhopilidae | Clade comprising 1 entire genus: *Gegeneophis* | Size: 4 (Fig. 2 of original study) | 97.93 | 4 | 4 | Gondwanan | Wet | recieved from author |
| *Hemidactylus* | **Lajmi, A.** & Karanth, P. K. Eocene–Oligocene cooling and the diversification of Hemidactylus geckos in Peninsular India. Mol. Phylogenet. Evol. 142, 106637 (2020). | Herpetofauna | Animalia -> Chordata -> Reptilia- > Squamata -> Gekkonidae | Clade comprising multiple species of 1 genus: *Hemidactylus* | Size: 48 (Fig. 4 of original study) | 35.79 | 48 | 48 | Asian | Wet + Dry | recieved from author |
| *Hemiphyllodactylus* | **Agarwal, I.**, Khandekar, A., Giri, V. B., Ramakrishnan, U. & Karanth, K. P. The hills are alive with geckos! A radiation of a dozen species on sky islands across peninsular India (Squamata: Gekkonidae, Hemiphyllodactylus) with the description of three new species. Org. Divers. Evol. 19, 341–361 (2019). | Herpetofauna | Animalia -> Chordata -> Reptilia- > Squamata -> Gekkonidae | Clade comprising multiple species of 1 genus: *Hemiphyllodactylus* | Size: 12 (Fig. 1 of original study) | 23.66 | 12 | 12 | Asian | Wet + Dry | recieved from author |
| Heterometrinae | **Loria, S. F.** & Prendini, L. Out of India, thrice: diversification of Asian forest scorpions reveals three colonizations of Southeast Asia. Sci. Rep. (2020). | Invertebrate | Animalia -> Arthropoda -> Arachnida -> Scorpiones -> Scorpionidae | Clade comprising multiple species of 6 genera: *Sahyadimetrus, Srilankametrus, Javanimetrus, Deccanometrus, Gigantometrus, Chersonesometrus* | Size: 23 (Fig. 4 of original study) | 48.06 | 23 | 22 | Gondwanan | Wet + Dry | recieved from author |
| *Indonaia* | **Bolotov, I. N.** et al. Oriental freshwater mussels arose in East Gondwana and arrived to Asia on the Indian Plate and Burma Terrane. Sci. Rep. 12, 1–26 (2022). | Invertebrate | Animalia -> Mollusca -> Bivalvia -> Unionida -> Unionidae | Clade comprising 1 entire genus: *Indonaia* | Size: 57 (Fig. 2 of original study) | 49.44 | 8 | 6 | Gondwanan | FW | recieved from author |
| *Lamellidens* | **Bolotov, I. N.** et al. Oriental freshwater mussels arose in East Gondwana and arrived to Asia on the Indian Plate and Burma Terrane. Sci. Rep. 12, 1–26 (2022). | Invertebrate | Animalia -> Mollusca -> Bivalvia -> Unionida -> Unionidae | Clade comprising 1 entire genus: *Lamellidens* | Size: 57 (Fig. 2 of original study) | 46.08 | 9 | 6 | Gondwanan | FW | recieved from author |
| Memecylon | **Amarasinghe, P.** et al. Evolution and biogeography of Memecylon. Am. J. Bot. 108, 628–646 (2021). | Angiosperm | Plantae -> Angiospermae -> Myrtales -> Melastomataceae | Clade comprising multiple species of 1 genus: *Memecylon* | Size: 38 (Fig. 2 of original study) | 22.13 | 38 | 32 | Asian | Wet | recieved from author |
| *Microhyla1* | **Gorin, V. A.** et al. A little frog leaps a long way: compounded colonizations of the Indian Subcontinent discovered in the tiny Oriental frog genus Microhyla (Amphibia: Microhylidae). PeerJ 8, e9411 (2020). | Herpetofauna | Animalia -> Chordata -> Amphibia - > Anura -> Microhylidae | Clade comprising multiple species of 1 genus: *Microhyla* | Size: 51 (Fig. 2 of original study) | 18.82 | 5 | 5 | Asian | Wet + Dry | recieved from author |
| *Microhyla2* | **Gorin, V. A.** et al. A little frog leaps a long way: compounded colonizations of the Indian Subcontinent discovered in the tiny Oriental frog genus Microhyla (Amphibia: Microhylidae). PeerJ 8, e9411 (2020). | Herpetofauna | Animalia -> Chordata -> Amphibia - > Anura -> Microhylidae | Clade comprising multiple species of 1 genus: *Microhyla* | Size: 51 (Fig. 2 of original study) | 19.02 | 6 | 6 | Asian | Wet + Dry | recieved from author |
| Nyctibatrachidae | **Torsekar, V. R.** Lineage delimitation and diversification in Nyctibatrachus, an endemic frog genus from the Western Ghats. (2020). | Herpetofauna | Animalia -> Chordata -> Amphibia - > Anura -> Nyctibatrachidae | Clade comprising 2 entire genera: *Nyctibtrachus, Lankanectes* | Size: 58 (Fig. 2 of original study) | 41.26 | 58 | 58 | Gondwanan | Wet | recieved from author |
| *Ophisops* | **Agarwal, I.** & Ramakrishnan, U. A phylogeny of open-habitat lizards (Squamata: Lacertidae: Ophisops ) supports the antiquity of Indian grassy biomes. J. Biogeogr. 44, 2021–2032 (2017). | Herpetofauna | Animalia -> Chordata -> Reptilia- > Squamata -> Lacertidae | Clade comprising 1 entire genus: *Ophisops* | Size: 36 (Appendix S2.5 of original study) | 30 | 36 | 33 | Asian | Wet + Dry | wrote newick using date estimates |
| *Parreysia* | **Bolotov, I. N.** et al. Oriental freshwater mussels arose in East Gondwana and arrived to Asia on the Indian Plate and Burma Terrane. Sci. Rep. 12, 1–26 (2022). | Invertebrate | Animalia -> Mollusca -> Bivalvia -> Unionida -> Unionidae | Clade comprising 1 entire genus: *Parreysia* | Size: 57 (Fig. 2 of original study) | 48.55 | 5 | 5 | Gondwanan | FW | recieved from author |
| *Piper*_group_1 | **Sen, S.** et al. Origin and evolution of the genus Piper in Peninsular India. Mol. Phylogenet. Evol. 138, 102–113 (2019). | Angiosperm | Plantae -> Angiospermae -> Piperales -> Piperaceae | Clade comprising multiple species of 1 genus: *Piper* | Size: 155 (Fig. 3 of original study) | 10.43 | 6 | 6 | Asian | Wet | recieved from author |
| *Piper*_group_2 | **Sen, S.** et al. Origin and evolution of the genus Piper in Peninsular India. Mol. Phylogenet. Evol. 138, 102–113 (2019). | Angiosperm | Plantae -> Angiospermae -> Piperales -> Piperaceae | Clade comprising multiple species of 1 genus: *Piper* | Size: 155 (Fig. 3 of original study) | 25.18 | 10 | 9 | Asian | Wet | recieved from author |
| *Pseudophilautus* | **Meegaskumbura, M.** et al. Diversification of shrub frogs (Rhacophoridae, Pseudophilautus) in Sri Lanka – Timing and geographic context. Mol. Phylogenet. Evol. 132, 14–24 (2019). + Ellepola, G., and Meegaskumbura, M. "Diversification and biogeography of Rhacophoridae–a model testing approach." Frontiers in Ecology and Evolution 11 (2023): 1195689. | Herpetofauna | Animalia -> Chordata -> Amphibia - > Anura -> Rhacophoridae | Clade comprising 1 entire genus: *Pseudophilautus* | Size: 69 (Fig. 2 of original study) | 29.21 | 69 | 69 | Asian | Wet | intially wrote the newick tree using date estimates from Meegaskumbura et al, 2019; but later found it on an online repository of the data from Ellepola and Meegaskumbura, 2023 |
| Ranixalidae | **Ramesh, V.**, Vijayakumar, S. P., Gopalakrishna, T., Jayarajan, A. & Shanker, K. Determining levels of cryptic diversity within the endemic frog genera, Indirana and Walkerana, of the Western Ghats, India. PLOS ONE 15, e0237431 (2020). | Herpetofauna | Animalia -> Chordata -> Amphibia - > Anura -> Ranixalidae | Clade comprising entire family of 2 genera: *Indirana, Walkarana* | Size: 21 (Fig. 3 of original study) | 43.01 | 21 | 21 | Gondwanan | Wet | recieved from author |
| *Raorchestes* | **Vijayakumar, S. P.**, Menezes, R. C., Jayarajan, A. & Shanker, K. Glaciations, gradients, and geography: multiple drivers of diversification of bush frogs in the Western Ghats Escarpment. Proc. R. Soc. B Biol. Sci. 283, 20161011 (2016). + Cyriac, V.P., Mohan, A.V., Dinesh, K.P., Torsekar, V., Jayarajan, A., Swamy, P., et al. (2024). Diversifying in the mountains: spatiotemporal diversification of frogs in the Western Ghats biodiversity hotspot. Evolution, 78, 701–715. | Herpetofauna | Animalia -> Chordata -> Amphibia - > Anura -> Rhacophoridae | Clade comprising 1 entire genus: *Raorchestes* | Size: 74 (Fig. 1 of original study) | 36.98 | 74 | 69 | Asian | Wet | intially wrote the newick tree using date estimates from Vijayakumar et al, 2016; but later found it on an online repository of the data from Cyriac et al, 2024 |
| Rhysida_immarginata_complex | **Joshi, J.**, Karanth, P. K. & Edgecombe, G. D. The out-of-India hypothesis: evidence from an ancient centipede genus, Rhysida (Chilopoda: Scolopendromorpha) from the Oriental Region, and systematics of Indian species. Zool. J. Linn. Soc. 189, 828–861 (2020). | Invertebrate | Animalia -> Arthropoda -> Myriapoda -> Chilopoda -> Scolopendridae | Clade comprising multiple species of 1 genus: *Rhysida* | Size: 66 (Fig. 4 of original study) | 81.83 | 7 | 6 | Gondwanan | Wet | recieved from author |
| Rhysida_longipes_complex | **Joshi, J.**, Karanth, P. K. & Edgecombe, G. D. The out-of-India hypothesis: evidence from an ancient centipede genus, Rhysida (Chilopoda: Scolopendromorpha) from the Oriental Region, and systematics of Indian species. Zool. J. Linn. Soc. 189, 828–861 (2020). | Invertebrate | Animalia -> Arthropoda -> Myriapoda -> Chilopoda -> Scolopendridae | Clade comprising multiple species of 1 genus: *Rhysida* | Size: 66 (Fig. 4 of original study) | 94.5 | 4 | 4 | Gondwanan | Wet | recieved from author |
| Testudines | **Pereira, A. G.**, Sterli, J., Moreira, F. R. R. & Schrago, C. G. Multilocus phylogeny and statistical biogeography clarify the evolutionary history of major lineages of turtles. Mol. Phylogenet. Evol. 113, 59–66 (2017). | Herpetofauna | Animalia -> Chordata -> Reptilia- > Testudines -> Geoemydidae | Clade comprising multiple species of 5 genera: *Batagur, Geoclemys, Pangshura, Morenia, Hardella* | Size: 14 (Fig. 1 of original study) | 45.65 | 14 | 10 | Asian | FW | recieved from author |
| Uperodon | **Garg, S.** & Biju, S. D. New microhylid frog genus from Peninsular India with Southeast Asian affinity suggests multiple Cenozoic biotic exchanges between India and Eurasia. Sci. Rep. 9, 1906 (2019). | Herpetofauna | Animalia -> Chordata -> Amphibia - > Anura -> Microhylidae | Clade comprising 1 entire genus: *Uperodon* | Size: 6 (Fig. 4 of original study) | 36.5 | 6 | 6 | Asian | Wet + Dry | wrote newick using date estimates |
| Uropeltids | **Cyriac, V. P.** & Kodandaramaiah, U. Paleoclimate determines diversification patterns in the fossorial snake family Uropeltidae Cuvier, 1829. Mol. Phylogenet. Evol. 116, 97–107 (2017). | Herpetofauna | Animalia -> Chordata -> Reptilia- > Squamata -> Uropeltidae | Clade comprising 7 entire genera: *Melanophidium, Platyplecturus, Teretrurus, Pseudoplectrurus, Plectrurus, Rhinophis, Uropeltis* | Size: 44 (Fig. 1 of original study) | 45.29 | 44 | 44 | Asian | Wet | recieved from author |

| Lineage | Speciation rate (RPANDA) | Extinction rate (RPANDA) | Net diversification rate (RPANDA) | Speciation rate (CoMET) | Extinction rate (CoMET) | Net diversification rate (CoMET) | Speciation rate (ClaDS) | Extinction rate (ClaDS) | Net diversification rate (ClaDS) | Pulled Diversification Rate | Results (CDR) | Driver | Results (PDR) | Result (CRABS) |
|---|---|---|---|---|---|---|---|---|---|---|---|---|---|---|
| Acanthaceae | 0.234 | 0 | 0.234 | 0.278 | 0.032 | 0.245 | 0.249 | 0.029 | 0.22 | 0.25 | SC1 | Other | SC1 | declining rate shift in recent time |
| Ahaetullinae | 0.064 | 0 | 0.064 | 0.081 | 0.006 | 0.075 | 0.066 | 0.007 | 0.059 | 0.165 | SC1 | Other | Not SC1 | declining rate shift in recent time |
| Ceropegiae | 0.346 | 0 | 0.347 | 0.387 | 0.024 | 0.363 | 0.405 | 0.045 | 0.361 | 0.388 | Shift | Other | SC1 | declining rate shift in recent time |
| *Cnemaspis* | 0.064 | 0 | 0.064 | 0.073 | 0.006 | 0.067 | 0.065 | 0.003 | 0.063 | 0.081 | Shift | TDD | Not SC1 | declining rate shift in recent time |
| *Digitipes* | 0.01 | 0 | 0.01 | 0.013 | 0.001 | 0.012 | 0.013 | 0.001 | 0.012 | 0.024 | SC1 | Other | SC1 | constant throughout |
| Dravidogecko | 0.061 | 0 | 0.061 | 0.084 | 0.007 | 0.077 | 0.058 | 0.012 | 0.046 | 0.155 | SC1 | Other | Not SC1 | declining rate shift in recent time |
| Ethmostigmus | 0.012 | 0 | 0.012 | 0.018 | 0.001 | 0.017 | 0.011 | 0.001 | 0.011 | 0.052 | SC1 | Other | SC1 | constant throughout |
| *Eutropis* | 0.045 | 0 | 0.045 | 0.056 | 0.005 | 0.051 | 0.049 | 0.005 | 0.044 | 0.08 | SC1 | Other | SC1 | majorly constant throughout |
| Fan-Throated | 0.106 | 0.045 | 0.061 | 0.104 | 0.015 | 0.09 | 0.08 | 0.008 | 0.072 | 0.061 | SC1 | Other | SC1 | majorly constant throughout |
| *Geckoella* | 0.069 | 0.008 | 0.061 | 0.079 | 0.01 | 0.07 | 0.071 | 0.008 | 0.063 | 0.061 | Shift | Other | Not SC1 | declining rate shift in recent time |
| *Gegeneophis* | 0.04 | 0 | 0.04 | 0.049 | 0.003 | 0.046 | 0.04 | 0.005 | 0.036 | 0.108 | Shift | Other | SC1 | majorly constant throughout |
| *Gerrhopilus* | 0.006 | 0 | 0.006 | 0.009 | 0.001 | 0.009 | 0.003 | 0.001 | 0.002 | 0.042 | SC1 | Other | Not SC1 | constant throughout |
| *Hemidactylus* | 0.064 | 0 | 0.064 | 0.08 | 0.003 | 0.077 | 0.094 | 0.009 | 0.086 | 0.146 | SC2 | DD+TD | SC1 | declining rate shift in recent time |
| *Hemiphyllodactylus* | 0.092 | 0 | 0.092 | 0.114 | 0.014 | 0.1 | 0.095 | 0.012 | 0.083 | 0.095 | Shift | TDD | Not SC1 | rate shifts (peak) in recent time |
| Heterometrinae | 0.041 | 0 | 0.041 | 0.054 | 0.003 | 0.051 | 0.048 | 0.005 | 0.043 | 0.102 | SC2+Shift | TDD+H | SC1 | declining rate shift in recent time |
| *Indonaia* | 0.028 | 0 | 0.028 | 0.037 | 0.004 | 0.034 | 0.032 | 0.002 | 0.03 | 0.05 | SC1 | Other | SC1 | majorly constant throughout |
| *Lamellidens* | 0.033 | 0 | 0.033 | 0.041 | 0.004 | 0.038 | 0.032 | 0.002 | 0.03 | 0.053 | SC1 | Other | | majorly constant throughout |
| Memecylon | 0.124 | 0 | 0.124 | 0.13 | 0.006 | 0.124 | 0.206 | 0.014 | 0.193 | 0.529 | SC3+Shift | TDD | Not SC1 | rate shifts (peak) in recent time |
| *Microhyla1* | 0.045 | 0 | 0.045 | 0.063 | 0.005 | 0.057 | 0.07 | 0.004 | 0.066 | 0.16 | SC1 | Other | SC1 | ambiguous throughout |
| *Microhyla2* | 0.061 | 0 | 0.061 | 0.083 | 0.009 | 0.074 | 0.057 | 0.003 | 0.054 | 0.096 | Shift | Other | Not SC1 | rate shifts (peak) in recent time |
| Nyctibatrachidae | 0.091 | 0 | 0.091 | 0.103 | 0.006 | 0.097 | 0.103 | 0.006 | 0.097 | 0.134 | SC3+Shift | TDD | Not SC1 | declining rate shift in recent time |
| *Ophisops* | 0.249 | 0.178 | 0.071 | 0.233 | 0.183 | 0.05 | 0.151 | 0.014 | 0.137 | 0.071 | SC3+Shift | TDD | Not SC1 | rate shifts (peak) in recent time |
| *Parreysia* | 0.049 | 0.074 | -0.025 | 0.032 | 0.004 | 0.028 | 0.029 | 0.002 | 0.027 | -0.025 | SC1 | Other | Not SC1 | majorly constant throughout |
| *Piper*_group_1 | 0.148 | 0.092 | 0.057 | 0.158 | 0.018 | 0.141 | 0.074 | 0.003 | 0.071 | 0.057 | SC1 | Other | SC1 | ambiguous throughout |
| *Piper*_group_2 | 0.06 | 0 | 0.06 | 0.075 | 0.007 | 0.068 | 0.063 | 0.003 | 0.06 | 0.105 | SC1 | Other | SC1 | majorly constant throughout |
| *Pseudophilautus* | 0.077 | 0 | 0.077 | 0.093 | 0.003 | 0.09 | 0.184 | 0.108 | 0.076 | 0.154 | SC2 | TDD | SC1 | decreasing trend in multiple time bins |
| Ranixalidae | 0.065 | 0 | 0.065 | 0.075 | 0.008 | 0.067 | 0.073 | 0.006 | 0.067 | 0.096 | SC3+Shift | TDD | Not SC1 | declining rate shift in recent time |
| *Raorchestes* | 0.069 | 0 | 0.069 | 0.082 | 0.003 | 0.079 | 0.109 | 0.025 | 0.084 | 0.13 | SC1 | TDD | SC1 | declining rate shift in recent time |
| Rhysida_immarginata_complex | 0.014 | 0 | 0.014 | 0.019 | 0.002 | 0.017 | 0.011 | 0.001 | 0.01 | 0.041 | SC1 | Other | SC1 | constant throughout |
| Rhysida_longipes_complex | 0.007 | 0 | 0.007 | 0.01 | 0.001 | 0.01 | 0.011 | 0.001 | 0.011 | NA | SC1 | Other | | constant throughout |
| Testudines | 0.036 | 0 | 0.036 | 0.046 | 0.003 | 0.043 | 0.036 | 0.004 | 0.033 | 0.09 | Shift | TDD | SC1 | declining rate shift in recent time |
| Uperodon | 0.024 | 0 | 0.024 | 0.033 | 0.003 | 0.03 | 0.02 | 0.004 | 0.017 | 0.145 | SC1 | Other | SC1 | declining rate shift in intermediate time |
| Uropeltids | 0.065 | 0 | 0.065 | 0.077 | 0.004 | 0.073 | 0.071 | 0.006 | 0.065 | 0.099 | Shift | Other | SC1 | majorly constant throughout |

| | Diversification Method | Statistical test/method | *p*-values | | | Phylogenetic signal | |
|---|---|---|---|---|---|---|---|
| | | | Angiosperms - Herpetofauna | Angiosperms - Invertebrates | Herpetofauna - Invertebrates | Pagel's λ of residuals (inference) | *p*-value for λ (inference) |
| Diversification rate vs. Taxonomic groups | RPANDA | **Kruskal-Wallis + Dunn Test** | 0.621 | **0.001** | **0.003** | - | - |
| | | **PhyloANOVA + Post Hoc** | 0.39 | 0.332 | 0.681 | < 0.001 (low phylogenetic signal) | 1 (i.e. not significantly deviating from λ=0) |
| | CoMET | **Kruskal-Wallis + Dunn Test** | 0.09 | **< 0.001** | **0.009** | - | - |
| | | **PhyloANOVA + Post Hoc** | 0.386 | 0.336 | 0.74 | < 0.001 (low phylogenetic signal) | 1 (i.e. not significantly deviating from λ=0) |
| | ClaDS (clade rates) | **Kruskal-Wallis + Dunn Test** | 0.242 | **< 0.001** | **0.006** | - | - |
| | | **PhyloANOVA + Post Hoc** | 0.386 | 0.336 | 0.74 | < 0.001 (low phylogenetic signal) | 1 (i.e. not significantly deviating from λ=0) |
| | ClaDS (tip rates) | **Kruskal-Wallis + Dunn Test** | **< 0.001** | **< 0.001** | **< 0.001** | - | - |
| | | **PhyloANOVA + Post Hoc** | 0.973293 | 0.95764 | 0.99457 | 0.994766 | **< 0.001 (i.e. significantly deviating from λ=0)** |
| | | | | **Asian - Gondwanan** | | **Pagel's λ of residuals (inference)** | ***p*-value for λ (inference)** |
| Diversification rate vs. Biogeographic origins | RPANDA | **Kruskal-Wallis + Dunn Test** | | **0.002** | | - | - |
| | | **PhyloANOVA + Post Hoc** | | 0.454 | | 1.018 | **< 0.001** |
| | CoMET | **Kruskal-Wallis + Dunn Test** | | **< 0.001** | | - | - |
| | | **PhyloANOVA + Post Hoc** | | 0.454 | | 0.654 | **< 0.001** |
| | ClaDS (clade rates) | **Kruskal-Wallis + Dunn Test** | | **< 0.001** | | - | - |
| | | **PhyloANOVA + Post Hoc** | | 0.454 | | 1.018 | **< 0.001** |
| | ClaDS (tip rates) | **Kruskal-Wallis + Dunn Test** | | **< 0.001** | | - | - |
| | | **PhyloANOVA + Post Hoc** | | 0.854 | | 0.994911 | **< 0.001** |
| | | | **FW - Wet** | **FW - Wet + Dry** | **Wet - Wet + Dry** | **Pagel's λ of residuals (inference)** | ***p*-value for λ (inference)** |
| Diversification rate vs. Habitat types | RPANDA | **Kruskal-Wallis + Dunn Test** | 0.169 | 0.065 | 1 | - | - |
| | | **PhyloANOVA + Post Hoc** | 0.682 | 0.634 | 0.877 | 1.018 | **< 0.001** |
| | CoMET | **Kruskal-Wallis + Dunn Test** | 0.288 | 0.132 | 1 | - | - |
| | | **PhyloANOVA + Post Hoc** | 0.682 | 0.634 | 0.877 | 0.69 | **< 0.001** |
| | ClaDS (clade rates) | **Kruskal-Wallis + Dunn Test** | 0.39 | 0.112 | 1 | - | - |
| | | **PhyloANOVA + Post Hoc** | 0.682 | 0.634 | 0.877 | 1.018 | **< 0.001** |
| | ClaDS (tip rates) | **Kruskal-Wallis + Dunn Test** | **< 0.001** | **< 0.001** | 0.19 | - | - |
| | | **PhyloANOVA + Post Hoc** | 0.996128 | 0.977268 | 0.908642 | 0.994819 | **< 0.001** |
| | | | ***p*-value** | ***R^2 (Mean R squared)*** | | **Pagel's λ of residuals (inference)** | ***p*-value for λ (inference)** |
| Diversification rate vs. Clade age | RPANDA | **LM** | **0.002** | 0.262 | | 1.02 | **0.006** |
| | | **PGLS** | 0.053 | 0.116 | | 0.453 | **0.042** |
| | CoMET | **LM** | **< 0.001** | 0.324 | | 0.41 | **0.017** |
| | | **PGLS** | **0.019** | 0.165 | | 0.501 | **0.008** |
| | ClaDS (clade rates) | **LM** | **0.002** | 0.274 | | 1.015 | **0.026** |
| | | **PGLS** | 0.051 | 0.1176 | | 0.487 | **0.023** |

| **Scenarios** | **$\alpha$** | **$\beta$** | **$\lambda 0$ - $\mu o$** | **Additional remarks** |
|---|---|---|---|---|
| SC 1a | 0 | 0 | >0 | $\mu o=0$ |
| SC 1b | 0 | 0 | >=0 | |
| SC 1c | <0 | <0 | >=0 | $\alpha=\beta$ |
| SC 1d | >0 | >0 | >=0 | $\alpha=\beta$ |
| SC 2a | >0 | 0 | >0 | $\mu o=0$ |
| SC 2b | >0 | 0 | >=0 | |
| SC 2c | 0 | <0 | >=0 | |
| SC 2d | >0 | <0 | >=0 | |
| SC 3a | >0 | 0 | <0 | $\lambda(t) - \mu(t) < 0$ for any t |
| SC 3b | 0 | <0 | <0 | $\lambda(t) - \mu(t) < 0$ for any t |
| SC 3c | >0 | >0 | <0 | $\alpha>\beta$; $\lambda(t) - \mu(t) < 0$ for any t |
| SC 3d | >0 | <0 | <0 | $\lambda(t) - \mu(t) < 0$ for any t |
| SC 4a | <0 | 0 | >0 | $\mu o=0$ |
| SC 4b | <0 | 0 | >=0 | |
| SC 4c | 0 | >0 | >=0 | |
| SC 4d | <0 | >0 | >=0 | |

| Lineage names | No. of tips | Clade age (Mya) | Speciation rate shift times (CoMET) (Mya) | CB.AICc | CB.lamb_par | CBD.AICc | CBD.lamb_par | CBD.mu_par | Best.Time.Model | Time.AICc | Time.lambda | Time.alpha | Time.omega | Time.mu | Time.beta | Time.pi | lowest.AICc | delta.AICc.CB | delta.AICc.CBD | delta.AICc.Time |
|---|---|---|---|---|---|---|---|---|---|---|---|---|---|---|---|---|---|---|---|---|
| Acanthaceae | 15 | 8.81 | | 66.081 | 0.234 | 68.773 | 0.234 | -8.36E-07 | b.exp.abs_d.0 | 68.772 | 0.237 | -0.004 | NA | NA | NA | NA | 66.081 | 0 | 2.692 | 2.691 |
| Ahaetullinae | 13 | 26.57 | | 84.795 | 0.064 | 87.632 | 0.064 | 2.17E-08 | b.epi.fix.1_d.0 | 83.238 | 0.093 | -0.098 | NA | NA | NA | NA | 83.238 | 1.557 | 4.394 | 0 |
| Ceropegiae | 67 | 9.69 | 0.679 | 269.833 | 0.347 | 271.959 | 0.346 | -1.14E-06 | b.epi.fix.1_d.0 | 267.796 | 0.205 | 0.241 | NA | NA | NA | NA | 267.796 | 2.037 | 4.163 | 0 |
| *Cnemaspis* | 83 | 63.43 | 3.806, 3.172 | 608.684 | 0.064 | 610.784 | 0.064 | -1.20E-07 | b.epi.fix.1_d.0 | 601.949 | 0.085 | -0.08 | NA | NA | NA | NA | 601.949 | 6.735 | 8.835 | 0 |
| *Digitipes* | 11 | 148.66 | | 102.54 | 0.01 | 105.596 | 0.01 | 1.32E-07 | b.epi.fix.113.5_d.0 | 102.223 | 0.009 | 0.128 | NA | NA | NA | NA | 102.223 | 0.317 | 3.372 | 0 |
| Dravidogecko | 7 | 19.68 | | 40.804 | 0.061 | 45.004 | 0.061 | 1.25E-08 | b.epi.fix.7_d.0 | 41.255 | 0.009 | 0.297 | NA | NA | NA | NA | 40.804 | 0 | 4.2 | 0.451 |
| Ethmostigmus | 5 | 65.21 | | 35.692 | 0.012 | 42.359 | 0.012 | 2.00E-08 | b.epi.fix.53_d.0 | 39.027 | 0.008 | 0.131 | NA | NA | NA | NA | 35.692 | 0 | 6.667 | 3.335 |
| *Eutropis* | 12 | 33.57 | | 84.23 | 0.045 | 87.163 | 0.045 | -4.30E-08 | b.epi.fix.31_d.0 | 84.498 | 0.038 | 0.205 | NA | NA | NA | NA | 84.23 | 0 | 2.933 | 0.268 |
| Fan-Throated | 17 | 30.05 | | 106.039 | 0.086 | 108.345 | 0.106 | 0.045 | b.epi.fix.10_d.0 | 106.244 | 0.069 | 0.166 | NA | NA | NA | NA | 106.039 | 0 | 2.306 | 0.205 |
| *Geckoella* | 14 | 32 | | 91.677 | 0.066 | 94.423 | 0.069 | 0.008 | b.epi.var.8_d.cst | 85.262 | 0.072 | 625.38 | 0.00E+00 | 0.024 | NA | NA | 85.262 | 6.415 | 9.16 | 0 |
| *Gegeneophis* | 14 | 37.5 | | 103.825 | 0.04 | 106.583 | 0.04 | 3.34E-08 | b.epi.fix.31_d.0 | 99.451 | 0.028 | 0.335 | NA | NA | NA | NA | 99.451 | 4.375 | 7.132 | 0 |
| *Gerrhopilus* | 4 | 97.93 | | 28.443 | 0.006 | 40.443 | 0.006 | -2.74E-09 | NA | NA | NA | NA | NA | NA | NA | NA | 28.443 | 0 | 12 | NA |
| *Hemidactylus* | 48 | 35.79 | | 346.292 | 0.065 | 348.472 | 0.064 | 3.00E-08 | b.lin.abs_d.0 | 336.273 | 0.016 | 0.005 | NA | NA | NA | NA | 336.273 | 10.019 | 12.199 | 0 |
| *Hemiphyllodactylus* | 12 | 23.66 | 3.076, 2.839 | 70.175 | 0.092 | 73.108 | 0.092 | 7.25E-07 | b.epi.fix.4_d.0 | 66.026 | 0.018 | 0.336 | NA | NA | NA | NA | 66.026 | 4.148 | 7.082 | 0 |
| Heterometrinae | 23 | 48.06 | 10.573, 9.612, 9.132, 8.651 | 178.809 | 0.041 | 181.218 | 0.041 | -1.15E-08 | b.lin.abs_d.0 | 174.434 | -0.013 | 0.003 | NA | NA | NA | NA | 174.434 | 4.375 | 6.785 | 0 |
| *Indonaia* | 8 | 49.44 | | 57.468 | 0.028 | 61.201 | 0.028 | 9.07E-09 | b.lin.abs_d.0 | 61.012 | 0.017 | 0.001 | NA | NA | NA | NA | 57.468 | 0 | 3.733 | 3.544 |
| *Lamellidens* | 9 | 46.08 | | 64.285 | 0.033 | 67.714 | 0.033 | 3.31E-07 | b.lin.abs_d.0 | 67.616 | 0.026 | 0 | NA | NA | NA | NA | 64.285 | 0 | 3.429 | 3.331 |
| Memecylon | 38 | 22.13 | 3.983, 3.762 | 224.579 | 0.124 | 226.811 | 0.124 | 3.86E-08 | b.lin.max_d.lin.max | 173.289 | -0.184 | 0.074 | NA | -4.212 | 0.506 | NA | 173.289 | 51.291 | 53.522 | 0 |
| *Microhyla1* | 5 | 18.82 | | 27.927 | 0.045 | 34.593 | 0.045 | -3.61E-08 | b.epi.fix.10_d.0 | 31.319 | 0.00E+00 | 0.314 | NA | NA | NA | NA | 27.927 | 0 | 6.667 | 3.392 |
| *Microhyla2* | 6 | 19.02 | | 33.408 | 0.061 | 38.408 | 0.061 | 7.34E-09 | b.epi.fix.7_d.0 | 28.685 | -0.01 | 0.419 | NA | NA | NA | NA | 28.685 | 4.723 | 9.723 | 0 |
| Nyctibatrachidae | 58 | 41.26 | 2.063, 1.650 | 382.202 | 0.091 | 384.349 | 0.091 | -7.18E-08 | b.lin.max_d.exp.abs | 373.77 | -0.025 | 0.056 | NA | 0.224 | 0.075 | NA | 373.77 | 8.432 | 10.578 | 0 |
| *Ophisops* | 36 | 30 | 6.9, 6.6, 6, 5.7, 1.2, 0.9 | 194.383 | 0.161 | 192.958 | 0.249 | 0.178 | b.epi.fix.2.5_d.0 | 175.146 | 0.032 | 0.513 | NA | NA | NA | NA | 175.146 | 19.238 | 17.812 | 0 |
| *Parreysia* | 5 | 48.55 | | 31.906 | 0.023 | 37.647 | 0.049 | 0.074 | b.epi.fix.10_d.0 | 31.896 | -0.001 | 0.302 | NA | NA | NA | NA | 31.896 | 0.01 | 5.751 | 0 |
| *Piper*_group_1 | 6 | 10.43 | | 28.121 | 0.118 | 33.03 | 0.148 | 0.092 | b.lin.abs_d.0 | 32.49 | 0.195 | -0.019 | NA | NA | NA | NA | 28.121 | 0 | 4.909 | 4.369 |
| *Piper*_group_2 | 10 | 25.18 | | 63.611 | 0.06 | 66.825 | 0.06 | 9.69E-08 | b.epi.fix.16_d.0 | 63.984 | 0.043 | 0.195 | NA | NA | NA | NA | 63.611 | 0 | 3.214 | 0.373 |
| *Pseudophilautus* | 69 | 29.21 | 6.719 | 479.135 | 0.077 | 481.257 | 0.077 | 1.78E-08 | b.exp.max_d.0 | 467.371 | 0.041 | 0.062 | NA | NA | NA | NA | 467.371 | 11.763 | 13.885 | 0 |
| Ranixalidae | 21 | 43.01 | 4.301, 3.871, 3.441 | 143.974 | 0.065 | 146.43 | 0.065 | -9.66E-08 | b.lin.max_d.exp.abs | 136.73 | -0.085 | 0.038 | NA | 0.095 | 0.098 | NA | 136.73 | 7.244 | 9.7 | 0 |
| *Raorchestes* | 74 | 36.98 | 4.068, 3.699, 3.328, 2.959 | 530.054 | 0.07 | 532.167 | 0.069 | -2.62E-08 | b.lin.max_d.lin.abs | 509.136 | -0.115 | 0.067 | NA | 0.09 | 0.053 | NA | 509.136 | 20.918 | 23.031 | 0 |
| Rhysida_immarginata_complex | 7 | 81.83 | | 55.773 | 0.014 | 59.973 | 0.014 | 9.15E-09 | b.epi.fix.34.5_d.0 | 56.323 | 0.009 | 0.123 | NA | NA | NA | NA | 55.773 | 0 | 4.2 | 0.55 |
| Rhysida_longipes_complex | 4 | 94.5 | | 28.044 | 0.007 | 40.044 | 0.007 | 2.82E-09 | NA | NA | NA | NA | NA | NA | NA | NA | 28.044 | 0 | 12 | NA |
| Testudines | 14 | 45.65 | 8.674, 8.218 | 106.423 | 0.036 | 109.18 | 0.036 | -1.92E-08 | b.lin.max_d.0 | 104.962 | -0.003 | 0.002 | NA | NA | NA | NA | 104.962 | 1.46 | 4.218 | 0 |
| Uperodon | 6 | 36.5 | | 40.949 | 0.024 | 45.949 | 0.024 | -1.40E-08 | b.lin.max_d.0 | 41.047 | -0.039 | 0.004 | NA | NA | NA | NA | 40.949 | 0 | 5 | 0.098 |
| Uropeltids | 44 | 45.29 | | 315.566 | 0.065 | 317.763 | 0.065 | -7.12E-09 | b.epi.fix.13_d.0 | 313.463 | 0.054 | 0.123 | NA | NA | NA | NA | 313.463 | 2.103 | 4.3 | 0 |

| | CONTINUED . . . | | | | | | | | | | | | | | | |
|---|---|---|---|---|---|---|---|---|---|---|---|---|---|---|---|---|
| Lineage names | best.model | lambda0 | alpha | omega | mu0 | beta | pi | additional remark | Pulled diversification rate estimated by fitting time-independent model | AIC of time-independent PDR model | BIC of time-independent PDR model | Best_grid_size for time-dependent PDR model | AIC of time-dependent PDR model | BIC of time-dependent PDR model | delta.AIC (time-dependent - time-independent) | delta.BIC (time-dependent - time-independent) |
| Acanthaceae | CB | 0.234 | 0 | 0 | 0 | 0 | 0 | | 0.25 | 67.76 | 69.038 | 1 | 67.76 | 69.038 | 0 | 0 |
| Ahaetullinae | CB | 0.064 | 0 | 0 | 0 | 0 | 0 | | 0.165 | 83.648 | 84.618 | 2 | 80.46 | 81.914 | -3.188 | -2.704 |
| Ceropegiae | b.epi.fix.1_d.0 | 0.205 | 0.241 | 0 | 0 | 0 | 0 | | 0.388 | 271.503 | 275.882 | 1 | 271.503 | 275.882 | 0 | 0 |
| *Cnemaspis* | b.epi.fix.1_d.0 | 0.085 | -0.08 | 0 | 0 | 0 | 0 | | 0.081 | 609.289 | 614.102 | 2 | 606.324 | 613.544 | -2.965 | -0.558 |
| *Digitipes* | CB | 0.01 | 0 | 0 | 0 | 0 | 0 | | 0.024 | 102.494 | 103.099 | 1 | 102.494 | 103.099 | 0 | 0 |
| Dravidogecko | CB | 0.061 | 0 | 0 | 0 | 0 | 0 | | 0.155 | 41.141 | 40.725 | 2 | 38.011 | 37.386 | -3.13 | -3.339 |
| Ethmostigmus | CB | 0.012 | 0 | 0 | 0 | 0 | 0 | | 0.052 | 35.107 | 33.88 | 1 | 35.107 | 33.88 | 0 | 0 |
| *Eutropis* | CB | 0.045 | 0 | 0 | 0 | 0 | 0 | | 0.08 | 85.025 | 85.82 | 1 | 85.025 | 85.82 | 0 | 0 |
| Fan-Throated | CB | 0.086 | 0 | 0 | 0 | 0 | 0 | | 0.061 | 107.488 | 109.033 | 1 | 107.488 | 109.033 | 0 | 0 |
| *Geckoella* | b.epi.var.8_d.cst | 0.072 | 625.38 | 0 | 0.024 | 0 | 0 | | 0.061 | 93.332 | 94.462 | 5 | 90.249 | 93.638 | -3.083 | -0.824 |
| *Gegeneophis* | b.epi.fix.31_d.0 | 0.028 | 0.335 | 0 | 0 | 0 | 0 | | 0.108 | 101.409 | 102.539 | 1 | 101.409 | 102.539 | 0 | 0 |
| *Gerrhopilus* | CB | 0.006 | 0 | 0 | 0 | 0 | 0 | results couldn't reach convergence for time-varying and DDD models | 0.042 | 26.888 | 25.086 | 5 | 14.339 | 8.931 | -12.549 | -16.155 |
| *Hemidactylus* | b.lin.abs_d.0 | 0.016 | 0.005 | 0 | 0 | 0 | 0 | | 0.146 | 336.155 | 339.855 | 1 | 336.155 | 339.855 | 0 | 0 |
| *Hemiphyllodactylus* | b.epi.fix.4_d.0 | 0.018 | 0.336 | 0 | 0 | 0 | 0 | | 0.095 | 71.773 | 72.569 | 2 | 64.905 | 66.099 | -6.868 | -6.47 |
| Heterometrinae | b.lin.abs_d.0 | -0.013 | 0.003 | 0 | 0 | 0 | 0 | | 0.102 | 174.322 | 176.504 | 1 | 174.322 | 176.504 | 0 | 0 |
| *Indonaia* | CB | 0.028 | 0 | 0 | 0 | 0 | 0 | | 0.05 | 58.465 | 58.356 | 2 | 56.924 | 56.761 | -1.541 | -1.595 |
| *Lamellidens* | CB | 0.033 | 0 | 0 | 0 | 0 | 0 | | 0.053 | 65.45 | 65.609 | NA | NA | NA | NA | NA |
| Memecylon | b.lin.max_d.lin.max | -0.184 | 0.074 | 0 | -4.212 | 0.506 | 0 | | 0.529 | 197.647 | 200.869 | 4 | 170.718 | 178.773 | -26.929 | -22.096 |
| *Microhyla1* | CB | 0.045 | 0 | 0 | 0 | 0 | 0 | | 0.16 | 27.767 | 26.54 | 2 | 25.87 | 24.029 | -1.897 | -2.511 |
| *Microhyla2* | b.epi.fix.7_d.0 | -0.01 | 0.419 | 0 | 0 | 0 | 0 | | 0.096 | 34.314 | 33.533 | 2 | 25.67 | 24.498 | -8.644 | -9.035 |
| Nyctibatrachidae | b.lin.max_d.exp.abs | -0.025 | 0.056 | 0 | 0.224 | 0.075 | 0 | | 0.134 | 381.478 | 385.564 | 4 | 370.648 | 380.864 | -10.83 | -4.7 |
| *Ophisops* | b.epi.fix.2.5_d.0 | 0.032 | 0.513 | 0 | 0 | 0 | 0 | | 0.071 | 192.594 | 195.705 | 4 | 179.295 | 187.071 | -13.299 | -8.634 |
| *Parreysia* | CB | 0.023 | 0 | 0 | 0 | 0 | 0 | | -0.025 | 31.647 | 30.42 | 2 | 26.527 | 24.686 | -5.12 | -5.734 |
| *Piper*_group_1 | CB | 0.118 | 0 | 0 | 0 | 0 | 0 | | 0.057 | 29.03 | 28.249 | 2 | 28.823 | 27.652 | -0.207 | -0.597 |
| *Piper*_group_2 | CB | 0.06 | 0 | 0 | 0 | 0 | 0 | | 0.105 | 64.575 | 64.97 | 1 | 64.575 | 64.97 | 0 | 0 |
| *Pseudophilautus* | b.exp.max_d.0 | 0.041 | 0.062 | 0 | 0 | 0 | 0 | | 0.154 | 468.551 | 472.99 | 1 | 468.551 | 472.99 | 0 | 0 |
| Ranixalidae | b.lin.max_d.exp.abs | -0.085 | 0.038 | 0 | 0.095 | 0.098 | 0 | | 0.096 | 145.108 | 147.099 | 2 | 137.499 | 140.486 | -7.609 | -6.613 |
| *Raorchestes* | b.lin.max_d.lin.abs | -0.115 | 0.067 | 0 | 0.09 | 0.053 | 0 | | 0.13 | 520.764 | 525.345 | 1 | 520.764 | 525.345 | 0 | 0 |
| Rhysida_immarginata_complex | CB | 0.014 | 0 | 0 | 0 | 0 | 0 | | 0.041 | 55.6 | 55.184 | 2 | 55.733 | 55.109 | 0.133 | -0.075 |
| Rhysida_longipes_complex | CB | 0.007 | 0 | 0 | 0 | 0 | 0 | results couldn't reach convergence for time-varying and DDD models | NA | NA | NA | NA | NA | NA | NA | NA |
| Testudines | CB | 0.036 | 0 | 0 | 0 | 0 | 0 | | 0.09 | 104.815 | 105.945 | 1 | 104.815 | 105.945 | 0 | 0 |
| Uperodon | CB | 0.024 | 0 | 0 | 0 | 0 | 0 | | 0.145 | 38.039 | 37.257 | 1 | 38.039 | 37.257 | 0 | 0 |
| Uropeltids | b.epi.fix.13_d.0 | 0.054 | 0.123 | 0 | 0 | 0 | 0 | | 0.099 | 315.194 | 318.716 | 1 | 315.194 | 318.716 | 0 | 0 |

| Lineage names | N.tips | Clade.age | CB.AICc | CB.lamb_par | CBD.AICc | CBD.lamb_par | CBD.mu_par | Best.DDD.model | DDD.aicc | DDD.lambda | DDD.mu | DDD.K | Best.Temp.Model | Temp.aicc | Temp.lambda | Temp.alpha | Temp.mu | Temp.beta | Best.Him.oro.Model | Him.oro.aicc | Him.oro.lambda | Him.oro.alpha |
|---|---|---|---|---|---|---|---|---|---|---|---|---|---|---|---|---|---|---|---|---|---|---|
| Acanthaceae | 15 | 8.81 | 66.081 | 0.234 | 68.773 | 0.234 | 0 | DDL + E | 70.465 | 1.411 | 0.325 | 14.138 | b.mtb.max_d.0 | 68.603 | 0.289 | 0.54 | NA | NA | b.lin.abs_d.0 | 68.773 | 0.13 | 0 |
| Ahaetullinae | 13 | 26.57 | 84.795 | 0.064 | 87.632 | 0.064 | 0 | DDL + E | 85.722 | 0.213 | 0 | 13.185 | b.mtb.abs_d.0 | 83.03 | 0.306 | 8.238 | NA | NA | b.lin.max_d.lin.abs | 95.141 | 0.08 | 0 |
| Ceropegiae | 67 | 9.69 | 269.833 | 0.347 | 271.959 | 0.347 | 0 | DDL + E | 273.495 | 0.414 | 0 | 221.285 | b.mtb.max_d.0 | 271.573 | 0.405 | 0.355 | NA | NA | b.cst_d.exp.max | 274.659 | 0.381 | NA |
| *Cnemaspis* | 83 | 63.43 | 608.684 | 0.064 | 610.791 | 0.065 | 0 | DDL + E | 610.734 | 0.085 | 0 | 186.891 | b.mtb.max_d.lin.abs | 601.548 | 0.217 | 4.834 | 0.032 | -0.013 | NA | NA | NA | NA |
| *Digitipes* | 11 | 148.66 | 102.54 | 0.01 | 105.596 | 0.01 | 0 | DDL + E | 106.24 | 0.033 | 0.00E+00 | 11.27 | NA | NA | NA | NA | NA | NA | NA | NA | NA | NA |
| Dravidogecko | 7 | 19.68 | 40.804 | 0.061 | 45.004 | 0.061 | 0 | DDL + E | 52.272 | 0.237 | 0.00E+00 | 7 | NA | NA | NA | NA | NA | NA | NA | NA | NA | NA |
| Ethmostigmus | 5 | 65.21 | 35.692 | 0.012 | 42.359 | 0.012 | 0 | DDL + E | 34.251 | 0.067 | 0.00E+00 | 5 | NA | NA | NA | NA | NA | NA | NA | NA | NA | NA |
| *Eutropis* | 12 | 33.57 | 84.23 | 0.045 | 87.163 | 0.045 | 0 | DDL + E | 86.514 | 0.956 | 0.054 | 11.317 | b.exp.max_d.0 | 85.867 | 0.024 | 0.071 | NA | NA | b.cst_d.exp.abs | 90.83 | 0.045 | NA |
| Fan-Throated | 17 | 30.05 | 106.039 | 0.086 | 108.345 | 0.106 | 0.045 | DDL + E | 110.388 | 0.436 | 0.144 | 15.722 | b.exp.max_d.0 | 108.134 | 0.114 | -0.044 | NA | NA | NA | NA | NA | NA |
| *Geckoella* | 14 | 32 | 91.677 | 0.066 | 94.423 | 0.069 | 0.008 | DDL + E | 95.675 | 0.5 | 0.1 | 13.013 | b.mtb.max_d.0 | 92.831 | 0.121 | 2.954 | NA | NA | NA | NA | NA | NA |
| *Gegeneophis* | 14 | 37.5 | 103.825 | 0.04 | 106.583 | 0.04 | 0 | DDL + E | 103.141 | 0.138 | 0.00E+00 | 14.09 | b.mtb.max_d.0 | 102.686 | 0.189 | 9.733 | NA | NA | NA | NA | NA | NA |
| *Gerrhopilus* | 4 | 97.93 | 28.443 | 0.006 | 40.443 | 0.006 | 0 | NA | NA | NA | NA | NA | NA | NA | NA | NA | NA | NA | NA | NA | NA | NA |
| *Hemidactylus* | 48 | 35.79 | 346.292 | 0.064 | 348.472 | 0.064 | 0 | DDL + E | 336.613 | 0.154 | 0 | 54.55 | b.mtb.max_d.0 | 334.732 | 0.213 | 6.475 | NA | NA | b.exp.abs_d.0 | 338.438 | 0.349 | 0 |
| *Hemiphyllodactylus* | 12 | 23.66 | 70.175 | 0.092 | 73.108 | 0.092 | 0 | DDL + E | 76.122 | 0.221 | 0.055 | 12.981 | b.lin.max_d.exp.abs | 67.384 | -1.153 | 0.335 | 0.037 | 0.527 | b.lin.max_d.lin.abs | 81.956 | 0.137 | 0 |
| Heterometrinae | 23 | 48.06 | 178.809 | 0.041 | 181.218 | 0.041 | 0 | DDL + E | 174.836 | 0.116 | 0 | 24.447 | b.lin.abs_d.0 | 175.757 | -0.006 | 0.007 | NA | NA | b.lin.max_d.0 | 174.222 | 0.098 | 0 |
| *Indonaia* | 8 | 49.44 | 57.468 | 0.028 | 61.201 | 0.028 | 0 | DDL + E | 66.619 | 0.088 | 0.00E+00 | 8.103 | NA | NA | NA | NA | NA | NA | NA | NA | NA | NA |
| *Lamellidens* | 9 | 46.08 | 64.285 | 0.033 | 67.714 | 0.033 | 0 | DDL + E | 72.027 | 0.084 | 0.00E+00 | 9.844 | NA | NA | NA | NA | NA | NA | NA | NA | NA | NA |
| Memecylon | 38 | 22.13 | 224.579 | 0.124 | 226.811 | 0.124 | 0 | DDL + E | 194.118 | 0.533 | 0 | 38.066 | b.lin.max_d.lin.abs | 178.157 | -5.444 | 2.517 | -8.932 | 3.074 | b.cst_d.exp.abs | 229.513 | 0.136 | NA |
| *Microhyla1* | 5 | 18.82 | 27.927 | 0.045 | 34.593 | 0.045 | 0 | DDL + E | 26.906 | 0.229 | 0.00E+00 | 5 | NA | NA | NA | NA | NA | NA | NA | NA | NA | NA |
| *Microhyla2* | 6 | 19.02 | 33.408 | 0.061 | 38.408 | 0.061 | 0 | DDL + E | 57.874 | 0.226 | 0.00E+00 | 6 | NA | NA | NA | NA | NA | NA | NA | NA | NA | NA |
| Nyctibatrachidae | 58 | 41.26 | 382.202 | 0.091 | 384.349 | 0.091 | 0 | DDL + E | 382.617 | 0.143 | 0 | 93.867 | b.mtb.max_d.0 | 376.383 | 0.171 | 2.773 | NA | NA | NA | NA | NA | NA |
| *Ophisops* | 36 | 30 | 194.383 | 0.161 | 192.958 | 0.249 | 0.179 | DDL + E | 194.403 | 0.516 | 0.292 | 31.479 | b.lin.max_d.lin.max | 171.55 | -1.251 | 0.825 | -0.781 | 0.767 | NA | NA | NA | NA |
| *Parreysia* | 5 | 48.55 | 31.906 | 0.023 | 37.647 | 0.049 | 0.074 | NA | NA | NA | NA | NA | NA | NA | NA | NA | NA | NA | NA | NA | NA | NA |
| *Piper*_group_1 | 6 | 10.43 | 28.121 | 0.118 | 33.03 | 0.148 | 0.092 | DDL + E | 51.807 | 2560879389 | 0.337 | 5 | NA | NA | NA | NA | NA | NA | NA | NA | NA | NA |
| *Piper*_group_2 | 10 | 25.18 | 63.611 | 0.06 | 66.825 | 0.06 | 0 | DDL + E | 69.86 | 0.152 | 0 | 10.947 | b.mtb.abs_d.0 | 65.923 | -0.105 | 2.701 | NA | NA | NA | NA | NA | NA |
| *Pseudophilautus* | 69 | 29.21 | 479.135 | 0.077 | 481.257 | 0.077 | 0 | DDL + E | 469.049 | 0.161 | 0.00E+00 | 84.216 | b.exp.abs_d.0 | 465.392 | 0.036 | 0.098 | NA | NA | b.lin.max_d.lin.max | 473.258 | 0.216 | 0.00E+00 |
| Ranixalidae | 21 | 43.01 | 143.974 | 0.065 | 146.43 | 0.065 | 0 | DDL + E | 147.123 | 0.117 | 0 | 28.858 | b.lin.max_d.lin.max | 139.168 | -0.984 | 0.231 | -0.765 | 0.21 | NA | NA | NA | NA |
| *Raorchestes* | 74 | 36.98 | 530.054 | 0.069 | 532.167 | 0.069 | 0 | DDL + E | 521.07 | 0.138 | 0.00E+00 | 93.383 | b.mtb.abs_d.0 | 510.51 | 0.221 | 6.151 | NA | NA | b.lin.max_d.0 | 523.875 | 0.153 | 0.00E+00 |
| Rhysida_immarginata_complex | 7 | 81.83 | 55.773 | 0.014 | 59.973 | 0.014 | 0 | DDL + E | 66.712 | 0.056 | 0.00E+00 | 7 | NA | NA | NA | NA | NA | NA | NA | NA | NA | NA |
| Rhysida_longipes_complex | 4 | 94.5 | 28.044 | 0.007 | 40.044 | 0.007 | 0 | NA | NA | NA | NA | NA | NA | NA | NA | NA | NA | NA | NA | NA | NA | NA |
| Testudines | 14 | 45.65 | 106.423 | 0.036 | 109.18 | 0.036 | 0 | DDL + E | 106.749 | 0.114 | 0 | 14.319 | b.lin.max_d.0 | 102.539 | -0.058 | 0.014 | NA | NA | b.lin.abs_d.0 | 105.321 | 0.093 | 0 |
| Uperodon | 6 | 36.5 | 40.949 | 0.024 | 45.949 | 0.024 | 0 | DDL + E | 60.692 | 0.159 | 0.00E+00 | 6 | NA | NA | NA | NA | NA | NA | NA | NA | NA | NA |
| Uropeltids | 44 | 45.29 | 315.566 | 0.065 | 317.763 | 0.065 | 0 | DDL + E | 316.711 | 0.105 | 0 | 68.16 | b.lin.max_d.0 | 314.82 | 0.026 | 0.007 | NA | NA | b.cst_d.exp.abs | 320.071 | 0.065 | NA |

**CONTINUED . . .**

| Lineage names | Him.oro.mu | Him.oro.beta | Best.C4.Exp.Model | C4.Exp.aicc | C4.Exp.lambda | C4.Exp.alpha | C4.Exp.mu | C4.Exp.beta | lowest.aicc | delta.aicc.CB | delta.aicc.CBD | delta.aicc.DDD | delta.aicc.Temp | delta.aicc.Him.oro | delta.aicc.C4.Exp | best.driver | additional remark |
|---|---|---|---|---|---|---|---|---|---|---|---|---|---|---|---|---|---|
| Acanthaceae | NA | NA | b.lin.abs_d.0 | 67.362 | 0.179 | 0.054 | NA | NA | 66.081 | 0 | 2.692 | 4.384 | 2.523 | 2.693 | 1.282 | Other | |
| Ahaetullinae | 0.012 | 0 | NA | NA | NA | NA | NA | NA | 83.03 | 1.765 | 4.601 | 2.691 | 0 | 12.11 | NA | Other | |
| Ceropegiae | 0 | 0.002 | b.exp.abs_d.0 | 271.893 | 0.349 | -0.017 | NA | NA | 269.833 | 0 | 2.126 | 3.662 | 1.739 | 4.826 | 2.06 | Other | |
| *Cnemaspis* | NA | NA | NA | NA | NA | NA | NA | NA | 601.548 | 7.136 | 9.244 | 9.186 | 0 | NA | NA | Temp | |
| *Digitipes* | NA | NA | NA | NA | NA | NA | NA | NA | 102.54 | 0 | 3.056 | 3.7 | NA | NA | NA | Other | |
| Dravidogecko | NA | NA | NA | NA | NA | NA | NA | NA | 40.804 | 0 | 4.2 | 11.468 | NA | NA | NA | Other | |
| Ethmostigmus | NA | NA | NA | NA | NA | NA | NA | NA | 34.251 | 1.441 | 8.108 | 0 | NA | NA | NA | Other | |
| *Eutropis* | 0 | 0 | NA | NA | NA | NA | NA | NA | 84.23 | 0 | 2.933 | 2.285 | 1.638 | 6.6 | NA | Other | |
| Fan-Throated | NA | NA | NA | NA | NA | NA | NA | NA | 106.039 | 0 | 2.306 | 4.349 | 2.095 | NA | NA | Other | |
| *Geckoella* | NA | NA | NA | NA | NA | NA | NA | NA | 91.677 | 0 | 2.745 | 3.998 | 1.154 | NA | NA | Other | |
| *Gegeneophis* | NA | NA | NA | NA | NA | NA | NA | NA | 102.686 | 1.139 | 3.896 | 0.454 | 0 | NA | NA | Other | |
| *Gerrhopilus* | NA | NA | NA | NA | NA | NA | NA | NA | NA | NA | NA | NA | NA | NA | NA | Other | could not reach convergence for DDD, Temp, Him Oro and C4 Exp |
| *Hemidactylus* | NA | NA | NA | NA | NA | NA | NA | NA | 334.732 | 11.56 | 13.74 | 1.88 | 0 | 3.706 | NA | DDD+Temp | |
| *Hemiphyllodactylus* | 0.017 | 0 | NA | NA | NA | NA | NA | NA | 67.384 | 2.791 | 5.724 | 8.738 | 0 | 14.572 | NA | Temp | |
| Heterometrinae | NA | NA | NA | NA | NA | NA | NA | NA | 174.222 | 4.586 | 6.996 | 0.614 | 1.534 | 0 | NA | DDD+Temp+Him.oro | |
| *Indonaia* | NA | NA | NA | NA | NA | NA | NA | NA | 57.468 | 0 | 3.733 | 9.151 | NA | NA | NA | Other | |
| *Lamellidens* | NA | NA | NA | NA | NA | NA | NA | NA | 64.285 | 0 | 3.429 | 7.742 | NA | NA | NA | Other | |
| Memecylon | 0 | 0.002 | NA | NA | NA | NA | NA | NA | 178.157 | 46.422 | 48.654 | 15.961 | 0 | 51.356 | NA | Temp | |
| *Microhyla1* | NA | NA | NA | NA | NA | NA | NA | NA | 26.906 | 1.021 | 7.687 | 0 | NA | NA | NA | Other | |
| *Microhyla2* | NA | NA | NA | NA | NA | NA | NA | NA | 33.408 | 0 | 5 | 24.466 | NA | NA | NA | Other | |
| Nyctibatrachidae | NA | NA | NA | NA | NA | NA | NA | NA | 376.383 | 5.82 | 7.966 | 6.234 | 0 | NA | NA | Temp | |
| *Ophisops* | NA | NA | NA | NA | NA | NA | NA | NA | 171.55 | 22.834 | 21.408 | 22.853 | 0 | NA | NA | Temp | |
| *Parreysia* | NA | NA | NA | NA | NA | NA | NA | NA | NA | NA | NA | NA | NA | NA | NA | Other | could not reach convergence for DDD, Temp, Him Oro and C4 Exp |
| *Piper*_group_1 | NA | NA | NA | NA | NA | NA | NA | NA | 28.121 | 0 | 4.909 | 23.686 | NA | NA | NA | Other | |
| *Piper*_group_2 | NA | NA | NA | NA | NA | NA | NA | NA | 63.611 | 0 | 3.214 | 6.249 | 2.313 | NA | NA | Other | |
| *Pseudophilautus* | -9.80E-02 | 0.00E+00 | NA | NA | NA | NA | NA | NA | 465.392 | 13.743 | 15.865 | 3.657 | 0 | 7.867 | NA | Temp | |
| Ranixalidae | NA | NA | NA | NA | NA | NA | NA | NA | 139.168 | 4.807 | 7.263 | 7.956 | 0 | NA | NA | Temp | |
| *Raorchestes* | NA | NA | NA | NA | NA | NA | NA | NA | 510.51 | 19.544 | 21.657 | 10.56 | 0 | 13.365 | NA | Temp | |
| Rhysida_immarginata_complex | NA | NA | NA | NA | NA | NA | NA | NA | 55.773 | 0 | 4.2 | 10.939 | NA | NA | NA | Other | |
| Rhysida_longipes_complex | NA | NA | NA | NA | NA | NA | NA | NA | NA | NA | NA | NA | NA | NA | NA | Other | could not reach convergence for DDD, Temp, Him Oro and C4 Exp |
| Testudines | NA | NA | NA | NA | NA | NA | NA | NA | 102.539 | 3.884 | 6.642 | 4.211 | 0 | 2.782 | NA | Temp | |
| Uperodon | NA | NA | NA | NA | NA | NA | NA | NA | 40.949 | 0 | 5 | 19.743 | NA | NA | NA | Other | |
| Uropeltids | 0 | 0.001 | NA | NA | NA | NA | NA | NA | 314.82 | 0.746 | 2.943 | 1.891 | 0 | 5.251 | NA | Other | |

| Lineage | Taxonomic groups | Clade Age | Species Richness | Net-diversification Rate | Biogeographic Origins | Habitat Types | Scenarios | Diversity Dependence | Temperature Dependence | Himalayan-Orogeny Dependence |
|---|---|---|---|---|---|---|---|---|---|---|
| Acanthaceae | Angiosperm | 8.80558937 | 15 | 0.2198099826 | Asian | Wet + Dry | SC1 | No DDD | No TDD | No Him.oro |
| Ahaetullinae | Herpetofauna | 26.56505916 | 13 | 0.0594197912 | Asian | Wet + Dry | SC1 | No DDD | No TDD | No Him.oro |
| Ceropegiae | Angiosperm | 9.694371791 | 67 | 0.3606064998 | Asian | Wet | Shift | No DDD | No TDD | No Him.oro |
| *Cnemaspis* | Herpetofauna | 63.434892 | 83 | 0.06259263632 | Gondwanan | Wet + Dry | Shift | No DDD | TDD | No Him.oro |
| *Digitipes* | Invertebrate | 148.6598285 | 11 | 0.01181438806 | Gondwanan | Wet | SC1 | No DDD | No TDD | No Him.oro |
| Dravidogecko | Herpetofauna | 19.6755314 | 7 | 0.0460255148 | Asian | Wet | SC1 | No DDD | No TDD | No Him.oro |
| Ethmostigmus | Invertebrate | 65.20786887 | 5 | 0.01066123135 | Gondwanan | Wet | SC1 | No DDD | No TDD | No Him.oro |
| *Eutropis* | Herpetofauna | 33.56926215 | 12 | 0.0436472644 | Asian | Wet + Dry | SC1 | No DDD | No TDD | No Him.oro |
| Fan-Throated | Herpetofauna | 30.05 | 17 | 0.07212630156 | Asian | Wet + Dry | SC1 | No DDD | No TDD | No Him.oro |
| *Geckoella* | Herpetofauna | 32 | 14 | 0.06308338833 | Asian | Wet + Dry | Shift | No DDD | No TDD | No Him.oro |
| *Gegeneophis* | Herpetofauna | 37.5 | 14 | 0.03551053658 | Gondwanan | Wet | Shift | No DDD | No TDD | No Him.oro |
| *Gerrhopilus* | Herpetofauna | 97.930502 | 4 | 0.002280595666 | Gondwanan | Wet | SC1 | No DDD | No TDD | No Him.oro |
| *Hemidactylus* | Herpetofauna | 35.786529 | 48 | 0.08551833033 | Asian | Wet + Dry | SC2 | DDD | TDD | No Him.oro |
| *Hemiphyllodactylus* | Herpetofauna | 23.661538 | 12 | 0.0826757853 | Asian | Wet + Dry | Shift | No DDD | TDD | No Him.oro |
| Heterometrinae | Invertebrate | 48.0613186 | 23 | 0.04327173881 | Gondwanan | Wet + Dry | SC2+Shift | DDD | TDD | Him.oro |
| *Indonaia* | Invertebrate | 49.44461517 | 8 | 0.02999435841 | Gondwanan | FW | SC1 | No DDD | No TDD | No Him.oro |
| *Lamellidens* | Invertebrate | 46.08268875 | 9 | 0.02974241178 | Gondwanan | FW | SC1 | No DDD | No TDD | No Him.oro |
| Memecylon | Angiosperm | 22.127469 | 38 | 0.1925180794 | Asian | Wet | SC3+Shift | No DDD | TDD | No Him.oro |
| *Microhyla1* | Herpetofauna | 18.818149 | 5 | 0.0662073636 | Asian | Wet + Dry | SC1 | No DDD | No TDD | No Him.oro |
| *Microhyla2* | Herpetofauna | 19.016731 | 6 | 0.05401188905 | Asian | Wet + Dry | Shift | No DDD | No TDD | No Him.oro |
| Nyctibatrachidae | Herpetofauna | 41.256085 | 58 | 0.09746028458 | Gondwanan | Wet | SC3+Shift | No DDD | TDD | No Him.oro |
| *Ophisops* | Herpetofauna | 30 | 36 | 0.1368874588 | Asian | Wet + Dry | SC3+Shift | No DDD | TDD | No Him.oro |
| *Parreysia* | Invertebrate | 48.54884912 | 5 | 0.02694135914 | Gondwanan | FW | SC1 | No DDD | No TDD | No Him.oro |
| *Piper*_group_1 | Angiosperm | 10.4289128 | 6 | 0.07070009421 | Asian | Wet | SC1 | No DDD | No TDD | No Him.oro |
| *Piper*_group_2 | Angiosperm | 25.18193901 | 10 | 0.05993468044 | Asian | Wet | SC1 | No DDD | No TDD | No Him.oro |
| *Pseudophilautus* | Herpetofauna | 29.2112471 | 69 | 0.07634348001 | Asian | Wet | SC2+Shift | No DDD | TDD | No Him.oro |
| Ranixalidae | Herpetofauna | 43.009832 | 21 | 0.06684204214 | Gondwanan | Wet | SC3+Shift | No DDD | TDD | No Him.oro |
| *Raorchestes* | Herpetofauna | 36.9816 | 74 | 0.08375941625 | Asian | Wet | SC3+Shift | No DDD | TDD | No Him.oro |
| Rhysida_immarginata_complex | Invertebrate | 81.82658306 | 7 | 0.01025730002 | Gondwanan | Wet | SC1 | No DDD | No TDD | No Him.oro |
| Rhysida_longipes_complex | Invertebrate | 94.50163191 | 4 | 0.0108218796 | Gondwanan | Wet | SC1 | No DDD | No TDD | No Him.oro |
| Testudines | Herpetofauna | 45.652964 | 14 | 0.03272018266 | Asian | FW | Shift | No DDD | TDD | No Him.oro |
| Uperodon | Herpetofauna | 36.5 | 6 | 0.01668524552 | Asian | Wet + Dry | SC1 | No DDD | No TDD | No Him.oro |
| Uropeltids | Herpetofauna | 45.29276 | 44 | 0.0649799335 | Asian | Wet | Shift | No DDD | No TDD | No Him.oro |

| | pairs | SumOfSqs | F.Model | *R^2* | *p*-value | Adjusted *p*-values | *p-value* of Mantel test comparing FAMD distance matrix and phylogenetic distance matrix |
|---|---|---|---|---|---|---|---|
| **Scenarios** | SC1 vs Shift | 0.171 | 6.769 | 0.227 | 0.003 | 0.03 | 0.754 |
| | SC1 vs SC2 | 0.071 | 2.542 | 0.137 | 0.058 | 0.58 | |
| | SC1 vs SC2+Shift | 0.119 | 4.161 | 0.197 | 0.01 | 0.1 | |
| | SC1 vs SC3+Shift | 0.242 | 9.86 | 0.33 | 0.001 | 0.01 | |
| | Shift vs SC2 | 0.031 | 1.638 | 0.19 | 0.203 | 1 | |
| | Shift vs SC2+Shift | 0.052 | 2.507 | 0.239 | 0.091 | 0.91 | |
| | Shift vs SC3+Shift | 0.062 | 3.911 | 0.262 | 0.013 | 0.13 | |
| | SC2 vs SC2+Shift | 0.01 | 0.274 | 0.215 | 1 | 1 | |
| | SC2 vs SC3+Shift | 0.02 | 1.926 | 0.325 | 0.167 | 1 | |
| | SC2+Shift vs SC3+Shift | 0.023 | 1.47 | 0.227 | 0.276 | 1 | |
| **Taxonomic groups** | Angiosperm vs Herpetofauna | 0.075 | 2.953 | 0.114 | 0.033 | 0.099 | 0.763 |
| | Angiosperm vs Invertebrate | 0.15 | 9.646 | 0.467 | 0.002 | 0.006 | |
| | Herpetofauna vs Invertebrate | 0.376 | 16.218 | 0.384 | 0.001 | 0.003 | |
| **Biogeographic origins** | Asian vs Gondwanan | 0.33 | 13.066 | 0.297 | 0.001 | 0.001 | 0.759 |
| **Habitat types** | Wet + Dry vs Wet | 0.183 | 6.262 | 0.188 | 0.001 | 0.003 | 0.758 |
| | Wet + Dry vs FW | 0.143 | 5.337 | 0.262 | 0.003 | 0.009 | |
| | Wet vs FW | 0.048 | 1.693 | 0.086 | 0.185 | 0.555 | |
| **Temperature dependence** | No TDD vs TDD | 0.332 | 13.215 | 0.299 | 0.001 | 0.001 | 0.74 |
| **Diversity-dependence** | No DDD vs DDD | 0.092 | 2.782 | 0.082 | 0.009 | 0.009 | 0.74 |

### List of Figures

S1 **(Upper panel) The dynamics of diversification and diversity accumulation.** The broad diversification scenarios or the general trends in diversity accumulation – gradual accumulation, saturated accumulation, waxing-waning and exponential accumulation. **(Lower panel)** Alternative speciation and extinction dynamics that could generate the illustrated diversity accumulation profiles. Linear functions for both speciation and extinction (time-varying) have been illustrated here for simplicity, however, exponential formulations for the time-varying diversification rates were also used in the study. This figure has been made by modifying conceptual figures from two earlier studies ([24, 14]). . . . . . . . . . . . . . . . . . . . . . . . . . . . . . . . . . . . . . . . 19

S2 **Density plots illustrating the frequency distributions of endemicity (left), species richness (middle) and clade age (right) of PIP biota in our dataset.** These plots were generated using the *geom_density* function from the *ggplot2* package in R [13]. Different colours correspond to different taxonomic groups. Values on the Y-axis indicate the frequency (i.e. probability) of any value on the X-axis and sum of the values on the Y-axis for all the points on the curve would give a total of 1. . . . . . . . . . . . . . . . . . . . . . . 20

S3 **Comparisons of (a) speciation (page 21), (b) extinction rates (page 22), (c) net-diversification rates (page 23) and (d)pulled diversification rates (page 24)** across taxonomic groups (angiosperms, herpetofauna and invertebrates), habitat types (FW - Freshwater habitat, Wet - Wet forests, Wet+Dry - Wet and dry habitats) and biogeographic origins (Asian and Gondwanan) in 33 lineages estimated using three methods - *RPANDA* models, CoMET analysis and the ClaDS model [3, 20, 1]. Bonferroni-corrected *p*-values (inferred by Dunn's Test using Kruskal Wallis test parameters) on top of the boxplots indicate statistical significance in comparisons and only significant comparisons have been highlighted. Numbers in the x-axis labels indicate sample size within each category and the dotted line indicates the overall mean rate of speciation, extinction or net-diversification. In addition to the Kruskal-Wallis test, phylogenetic ANOVA was performed (results in page - 9), first, using the clade-rates and then, using the tip rates of species in each lineage, to correct for phylogenetic relatedness while comparing rates. Two phylogenetic trees **(e)** (page 25) – one, for comparing clade-rates **(upper panel)** and another, for comparing tip-rates **(lower panel)** were utilised for performing phylogenetic ANOVA. Coloured bands beside the tips in the larger tree (lower panel) indicate the extent of each lineage and alternate colours have been used todistinguish between two adjacent clades. . . . . . . . . . . . . . . . . . . . . . . . . 26

S4 **Prevalence of different diversification scenarios across PIP lineages.** The abundance of different diversification scenarios across all lineages, lineages with a minimum of 10 species, 20 species and 30 species, respectively with varying taxonomic and biogeographic affinities, and habitat preferences. Gradual accumulation is the most prevalent scenario when clades with less than 20 species are considered. More speciose clades favour time-varying diversification scenarios, many of which also show a strong episodic rate shift in the Neogene. 27

S5 **The priors (page 28) and results of the CRABS analysis (pages 29 - 124)** [25]. The different prior functions and values were used for extinction rates to obtain the corresponding pulled speciation rates and vice-versa that constitute the congruence class and this further helped generate the pulled diversification rates. Functions include constant rates and time-varying rates (linearly, exponentially and sigmoidally). Within pages 29 - 124, each consecutive triplet of pages correspond to one lineage, the name of which is written on the top right. Illustrations of the dynamics in the different congruent speciation ($\lambda$), extinction ($\mu$) and diversification ($\delta$) models have been provided. The upper panel plots the change in the slopes of each model and the lower panel summarises the trends in the change of slope (i.e. flat or constant, increasing and decreasing) for each time bin. . . . . . . . . . . . . . . . . . . . . . . . . . 125

S6 **Results from the Factor Analysis of Mixed Data** [26, 27] **(a) (page 126) (upper panel)** Scree plot indicates the percentages of total variance in the dataset that each ordination dimension explains, **(middle and bottom panel)** variables used in the analysis and their individual contributions to dimension-1 and dimension-2 respectively. Red dotted line indicates the average contribution by each variable to the corresponding ordination dimension. **(b) (page 127)** Quantitative and Qualitative variables on the reduced two-dimensional ordination space. (upper panel) Three quantitative variables – species richness, net-diversification rates and clade ages represented by arrows. Colours indicate the contribution (percentage) of each variable to the total variance of the dataset (legend provided in figure). Species richness seems to be more strongly correlated to net-diversification rate than to clade age and as has been shown in earlier figures, clade age and net-diversification rates seem to be inversely correlated. (lower panel) Positions of different categorical(qualitative) variables in the ordination space indicating their correlations in with each other. **(c) (page 128)** The FAMD analysis indicates high congruence in the clusters for temperature-dependence in diversification and for different diversification scenarios. And, temperature-dependence and scenarios have high contribution to Dimension 1 of the ordination which also is substantially contributed by net-diversification rates and species richness. Hence, we expected to observe some structure in species richness and diversification rates across different scenarios and groups of lineages showing varying support for temperature-dependence. **(upper panel)** Boxplots comparing species richness and net-diversifcation rates (Mann-Whitney U test) of lineages across lineages that show temperature-dependent diversification and the ones that do not. Lineages showing TDD seem to show significantly higher species richness and net-diversification rates than lineages that do not (p-values have been provided at the top the respective plots). **(bottom panel)** Boxplots comparing species richness and net-diversification rates across different diversification scenarios. Lineages following gradual accumulation (SC1) seem to show lower species richness and net-diversification rates than the ones following other diversification scenarios (tested using the Dunn's test following Kruskal-Wallis test). Significant comparisons have been highlighted and their corresponding corrected p-values have been provided. . . . . . . . . . . . . . . . . . . . . . . . . 129

S7 **Credibility in estimates through simulations**. Credibility of the best-fitting models was inferred through simulating 100 trees with the corresponding parameter estimates, conditioning on the crown ages of the lineages. Proportion of correctness in scenario estimations were assessed by analysing the diversification dynamics of the simulated trees using **(a) (page 130)** the birth-death models in RPANDA and by performing the **(b) (page 131)** CoMET analysis. Stacked bar plots in **(a)** and **(b)** indicate the proportions of correctly (black) and incorrectly (white) inferred diversification scenarios. Scatter plots indicate no apparent trend in the proportion of correctly inferred scenarios with the number of tips. **(c) (page 132)** Boxplots indicates the distributions of errors in estimating the empirical parameters - $\lambda_0$, $\alpha$, $\mu_0$ and $\beta$ through simulations. **(d) (page 133)** Dots indicate estimated times of significant diversification rate shifts in the simulated trees estimated using the SES model and CoMET analysis and the triangles and circles indicate estimated times of episodic rate shifts in empirical trees estimated using CoMET and the SES models, respectively. . 134

### 3 Figures

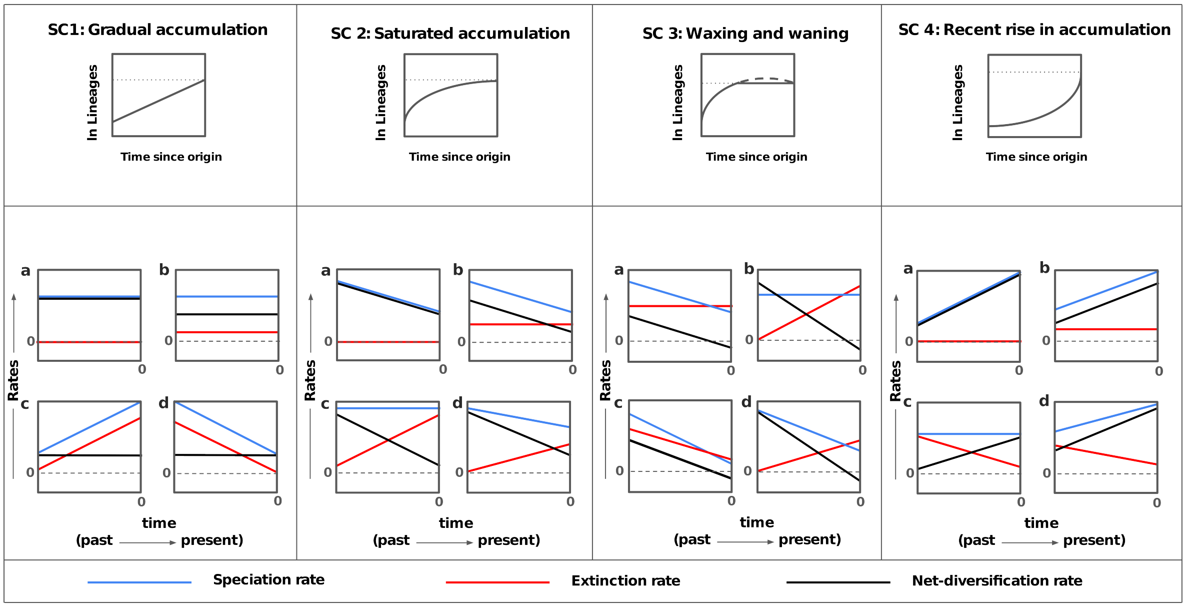

**Figure S1: (Upper panel) The dynamics of diversification and diversity accumulation.** The broad diversification scenarios or the general trends in diversity accumulation – gradual accumulation, saturated accumulation, waxing-waning and exponential accumulation. **(Lower panel)** Alternative speciation and extinction dynamics that could generate the illustrated diversity accumulation profiles. Linear functions for both speciation and extinction (time-varying) have been illustrated here for simplicity, however, exponential formulations for the time-varying diversification rates were also used in the study. This figure has been made by modifying conceptual figures from two earlier studies ([24, 14]).

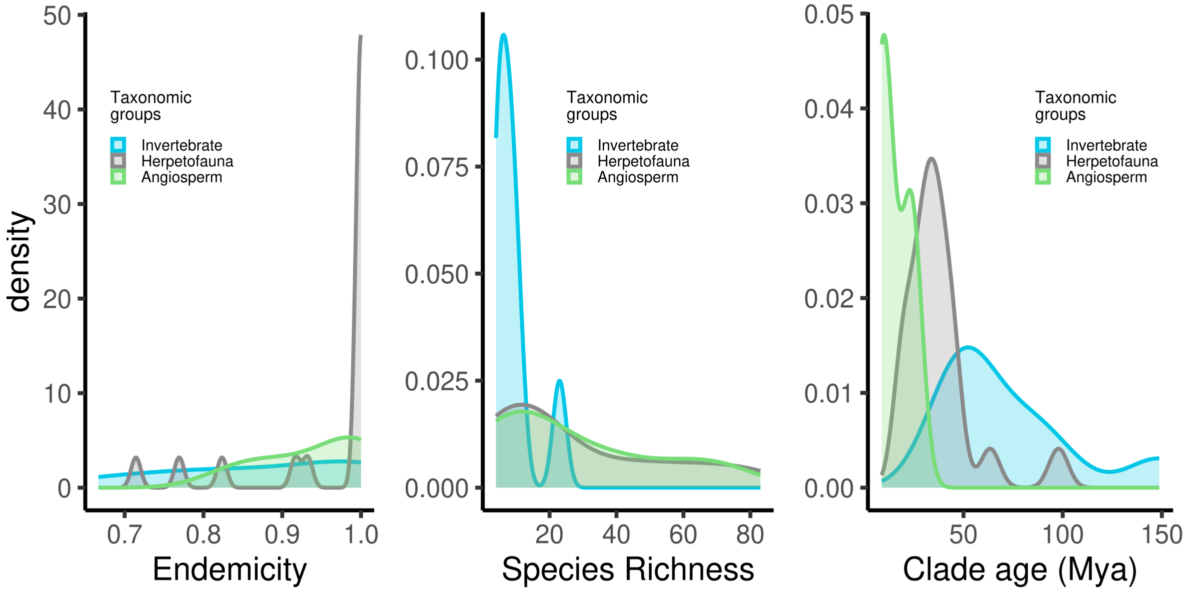

**Figure S2: Density plots illustrating the frequency distributions of endemicity (left), species richness (middle) and clade age (right) of PIP biota in our dataset.** These plots were generated using the *geom_density* function from the *ggplot2* package in R [13]. Different colours correspond to different taxonomic groups. Values on the Y-axis indicate the frequency (i.e. probability) of any value on the X-axis and sum of the values on the Y-axis for all the points on the curve would give a total of 1.

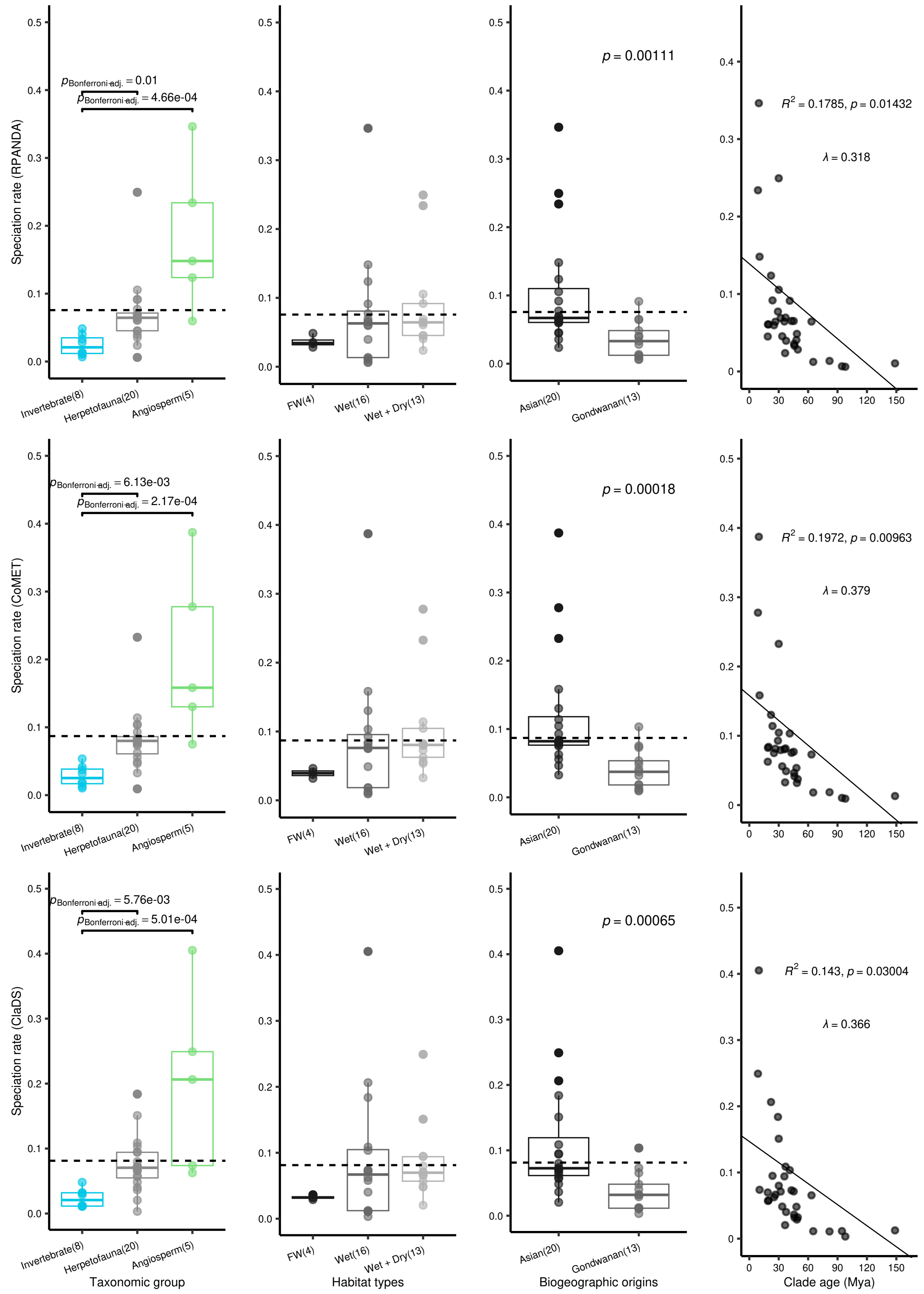

**(a)** Speciation rates

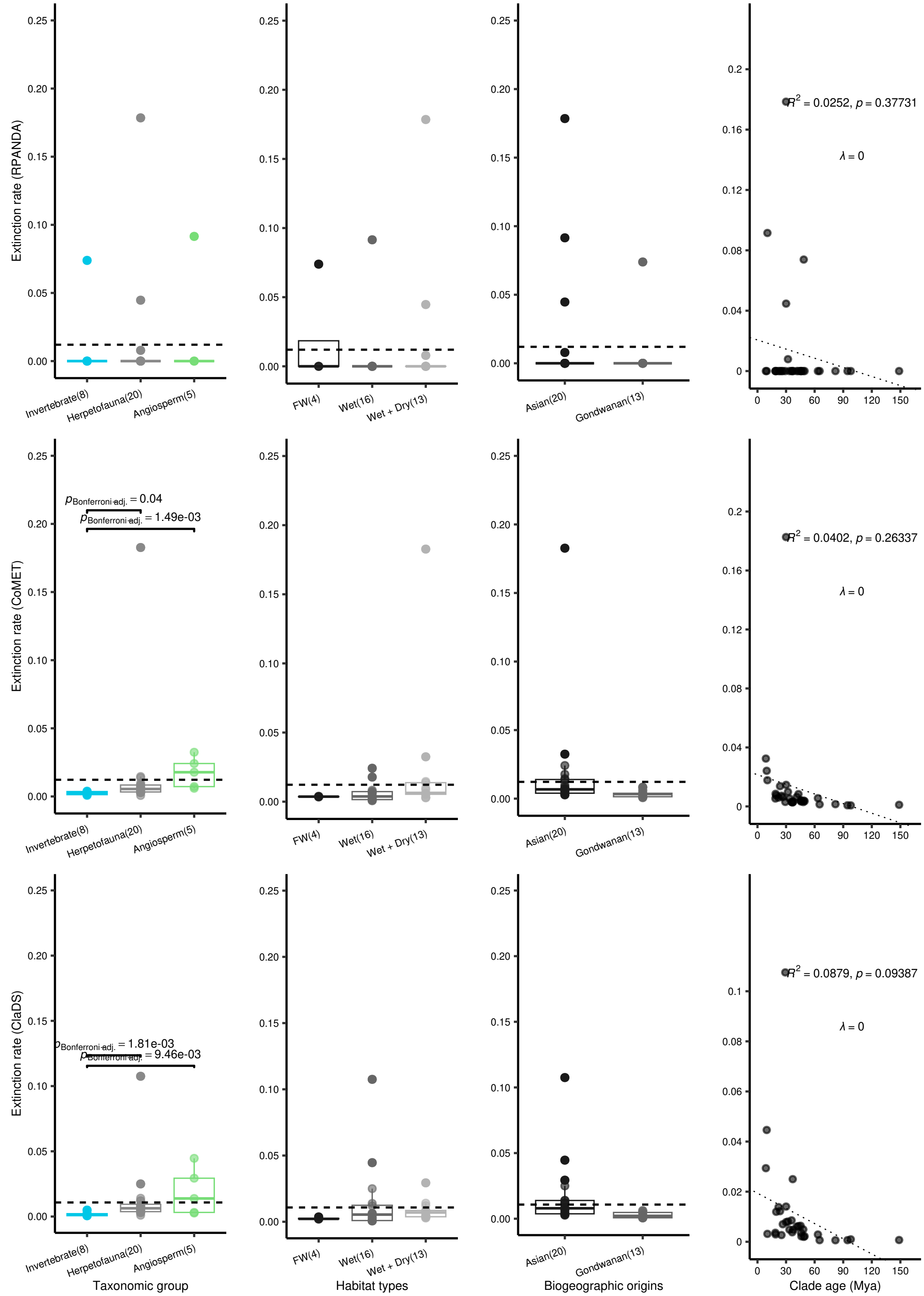

**(b)** Extinction rates

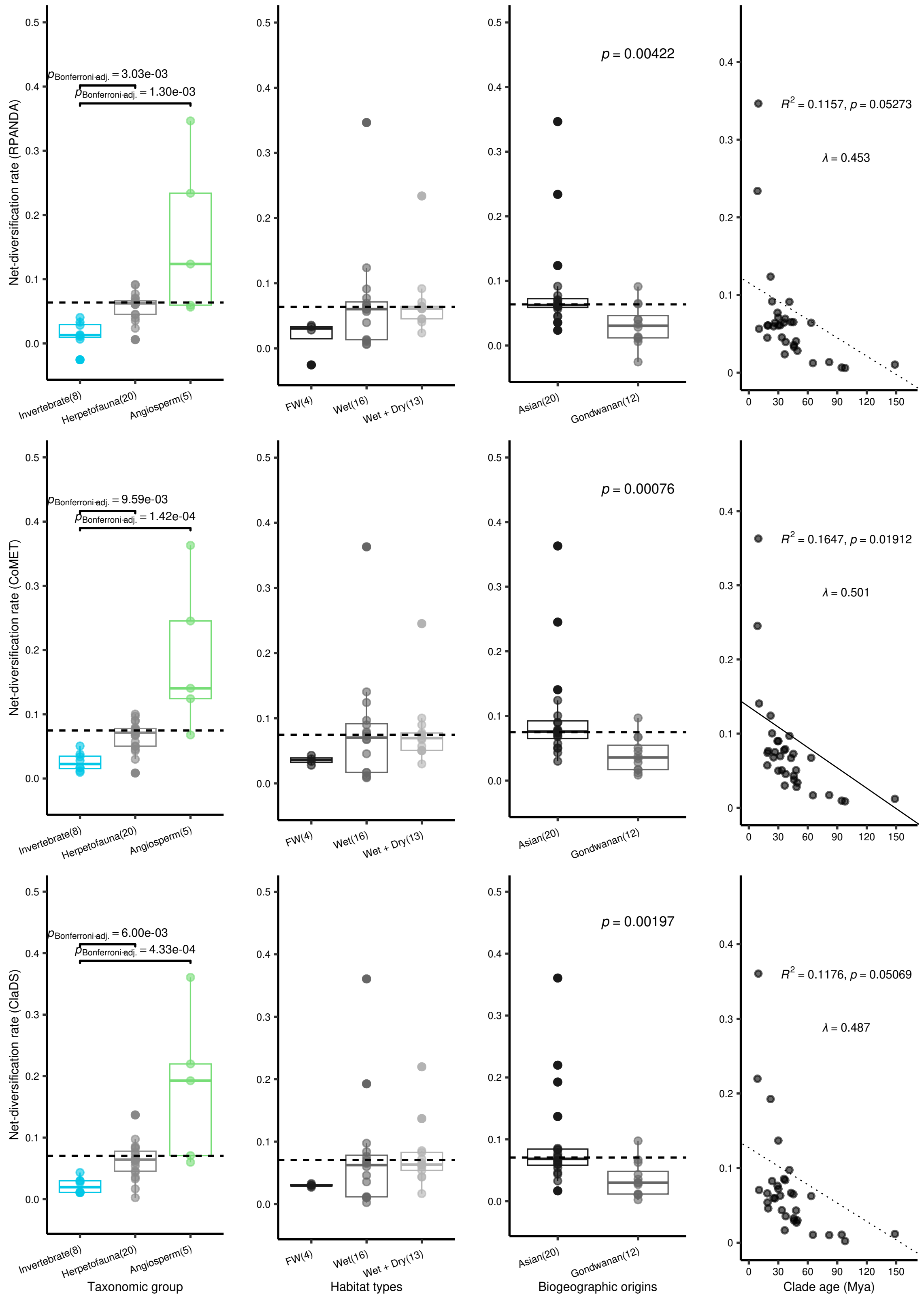

**(c)** Net-diversification rates

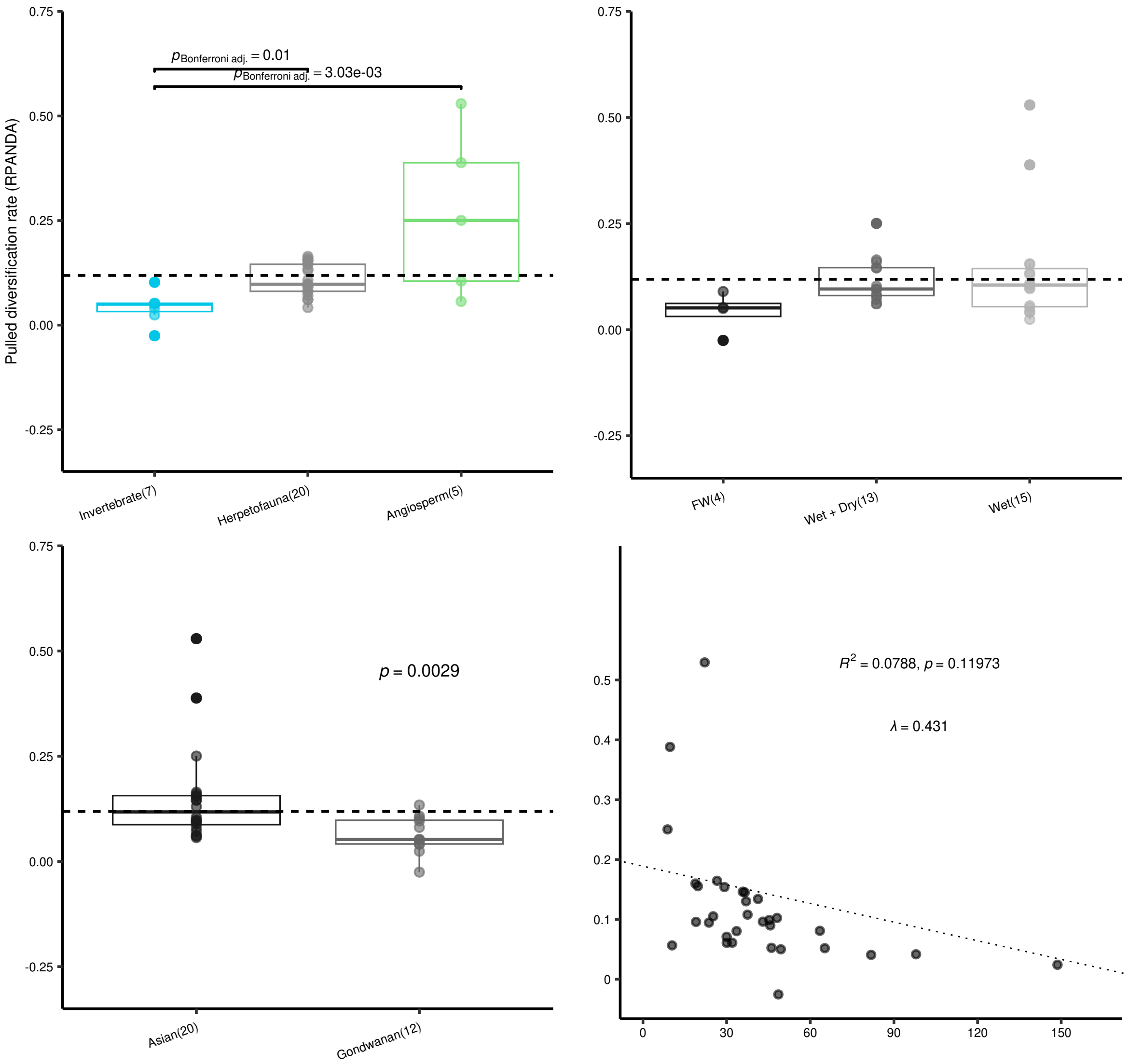

323 **(d)** Pulled diversification rates

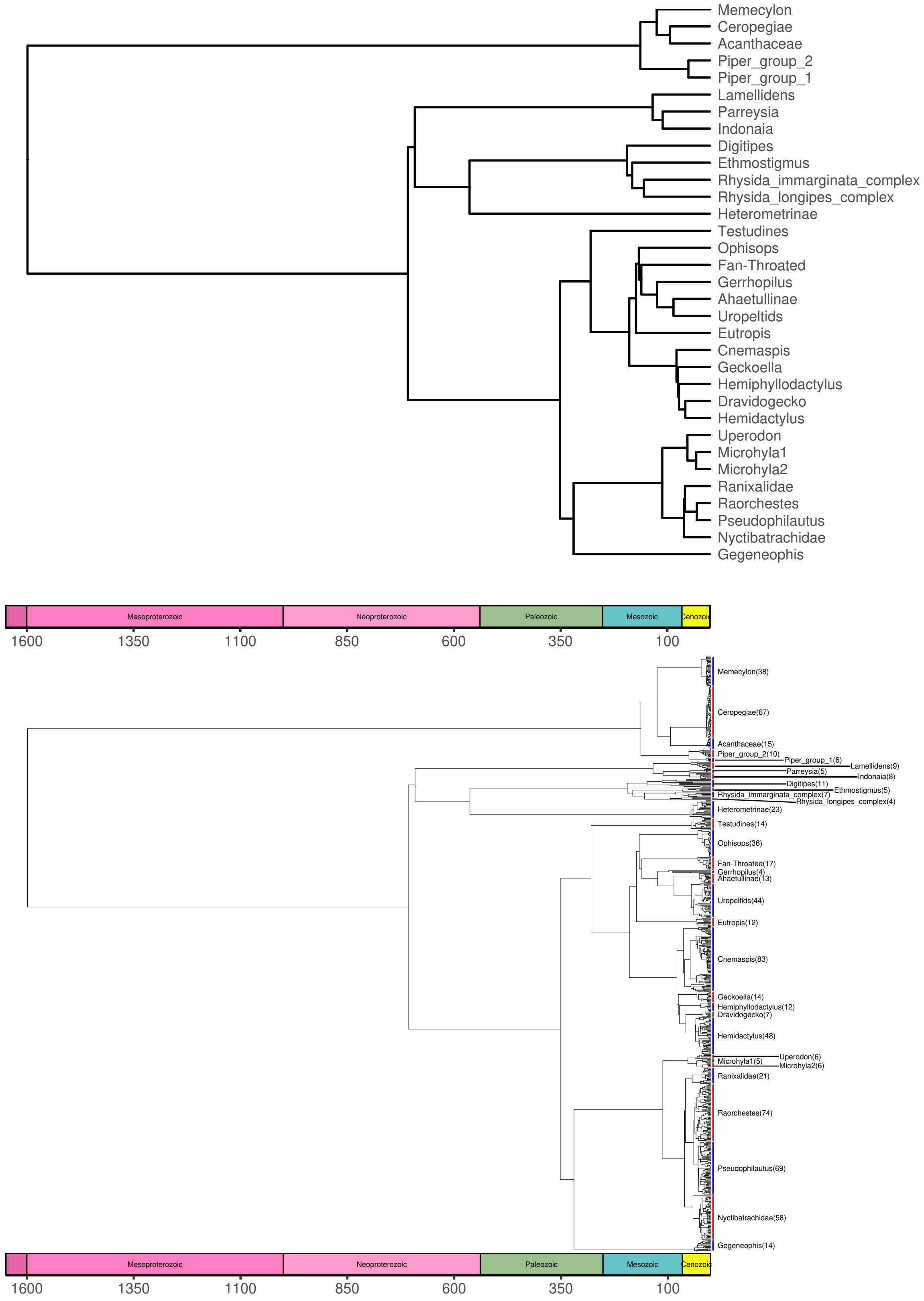

**(e)** Trees used in phylogenetic regressions

**Figure S3: Comparisons of (a) speciation (page 21), (b) extinction rates (page 22), (c) net-diversification rates (page 23) and (d)pulled diversification rates (page 24)** across taxonomic groups (angiosperms, herpetofauna and invertebrates), habitat types (FW - Freshwater habitat, Wet - Wet forests, Wet+Dry - Wet and dry habitats) and biogeographic origins (Asian and Gondwanan) in 33 lineages estimated using three methods - *RPANDA* models, CoMET analysis and the ClaDS model [3, 20, 1]. Bonferroni-corrected $p$-values (inferred by Dunn's Test using Kruskal Wallis test parameters) on top of the boxplots indicate statistical significance in comparisons and only significant comparisons have been highlighted. Numbers in the x-axis labels indicate sample size within each category and the dotted line indicates the overall mean rate of speciation, extinction or net-diversification. In addition to the Kruskal-Wallis test, phylogenetic ANOVA was performed (results in page - 9), first, using the clade-rates and then, using the tip rates of species in each lineage, to correct for phylogenetic relatedness while comparing rates. Two phylogenetic trees **(e)** (page 25) – one, for comparing clade-rates **(upper panel)** and another, for comparing tip-rates **(lower panel)** were utilised for performing phylogenetic ANOVA. Coloured bands beside the tips in the larger tree (lower panel) indicate the extent of each lineage and alternate colours have been used todistinguish between two adjacent clades.

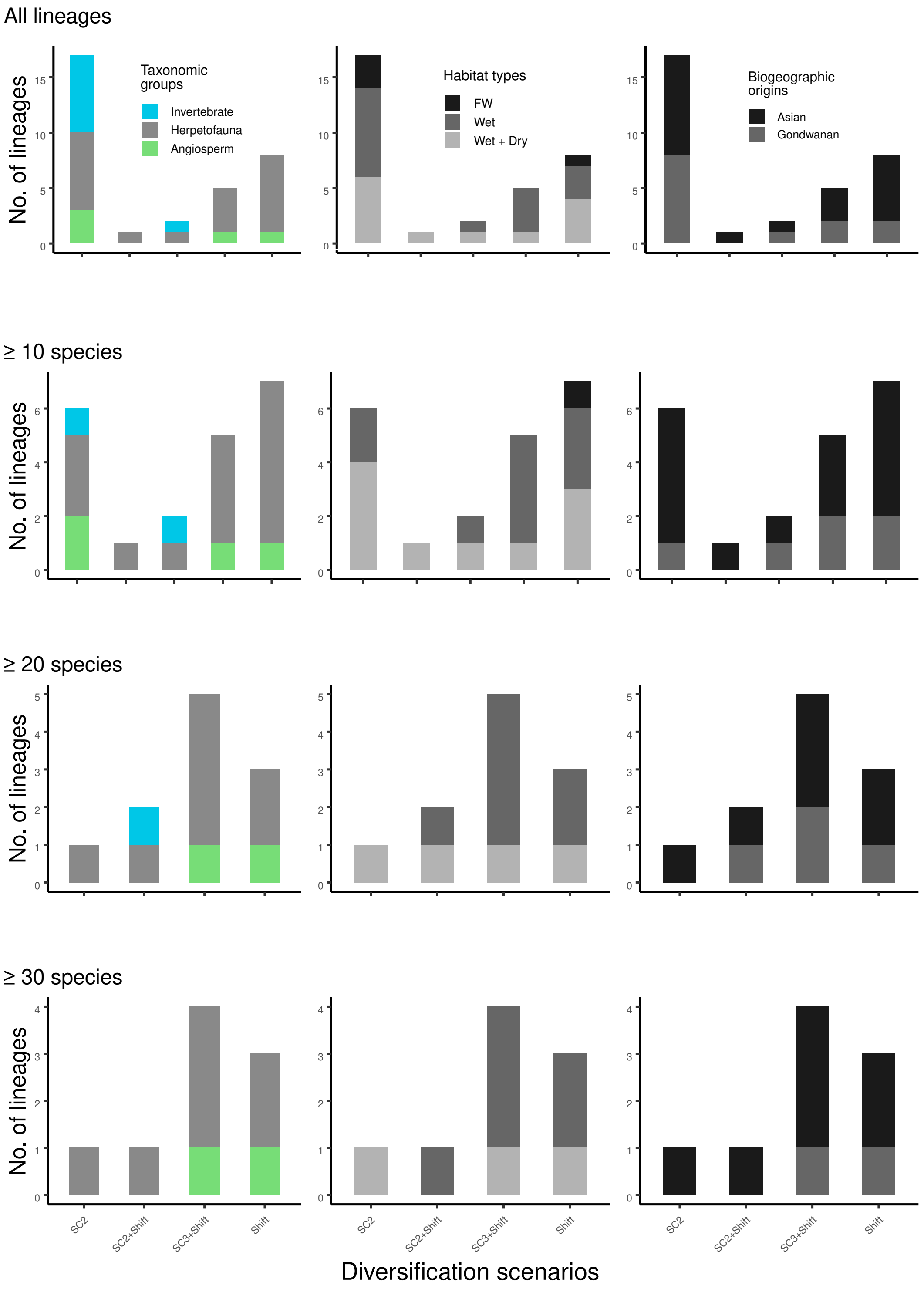

**Figure S4: Prevalence of different diversification scenarios across PIP lineages.** The abundance of different diversification scenarios across all lineages, lineages with a minimum of 10 species, 20 species and 30 species, respectively with varying taxonomic and biogeographic affinities, and habitat preferences. Gradual accumulation is the most prevalent scenario when clades with less than 20 species are considered. More speciose clades favour time-varying diversification scenarios, many of which also show a strong episodic rate shift in the Neogene.

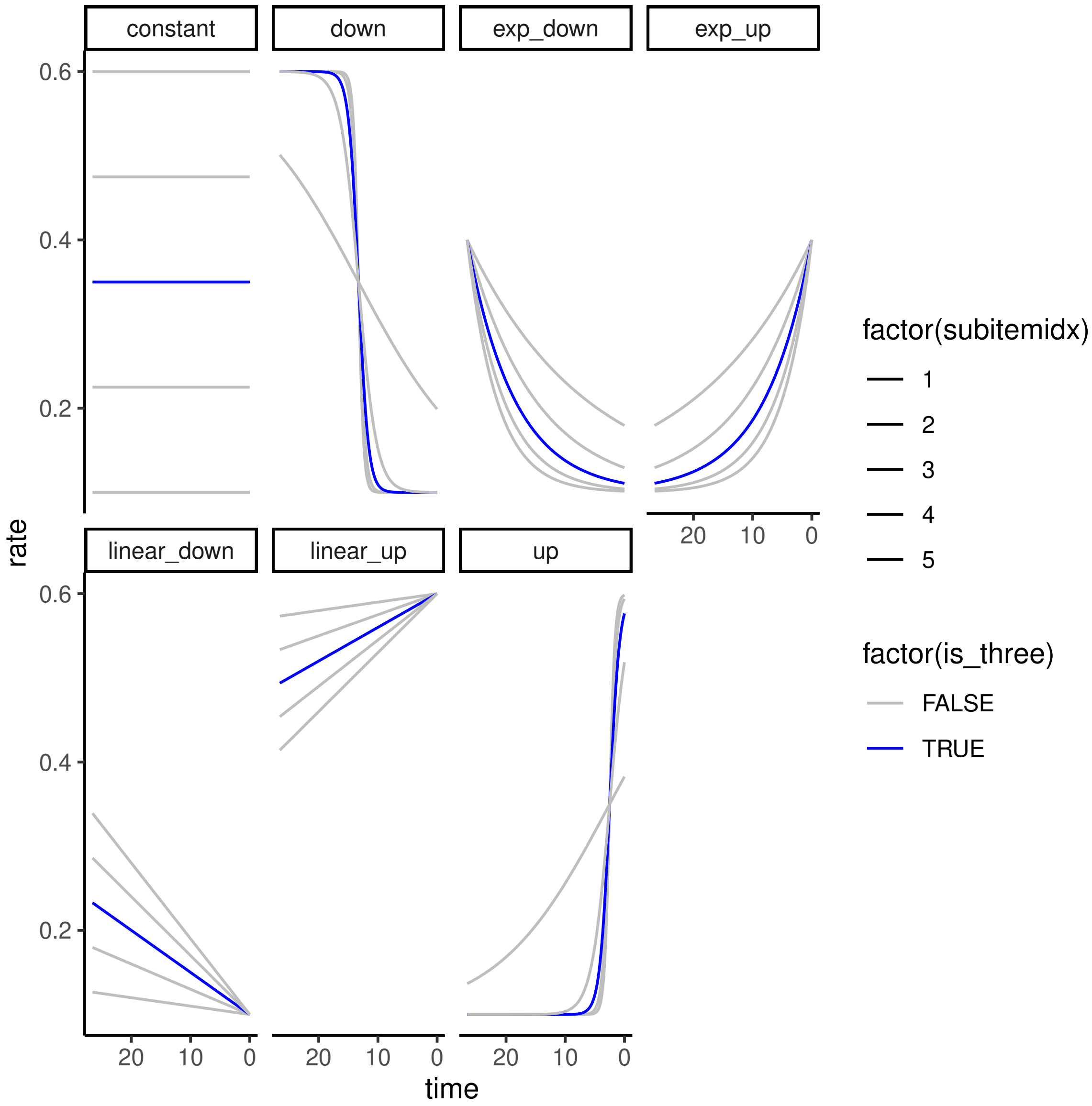

**(a)** Prior functions with varying parameters used in the CRABS analyses

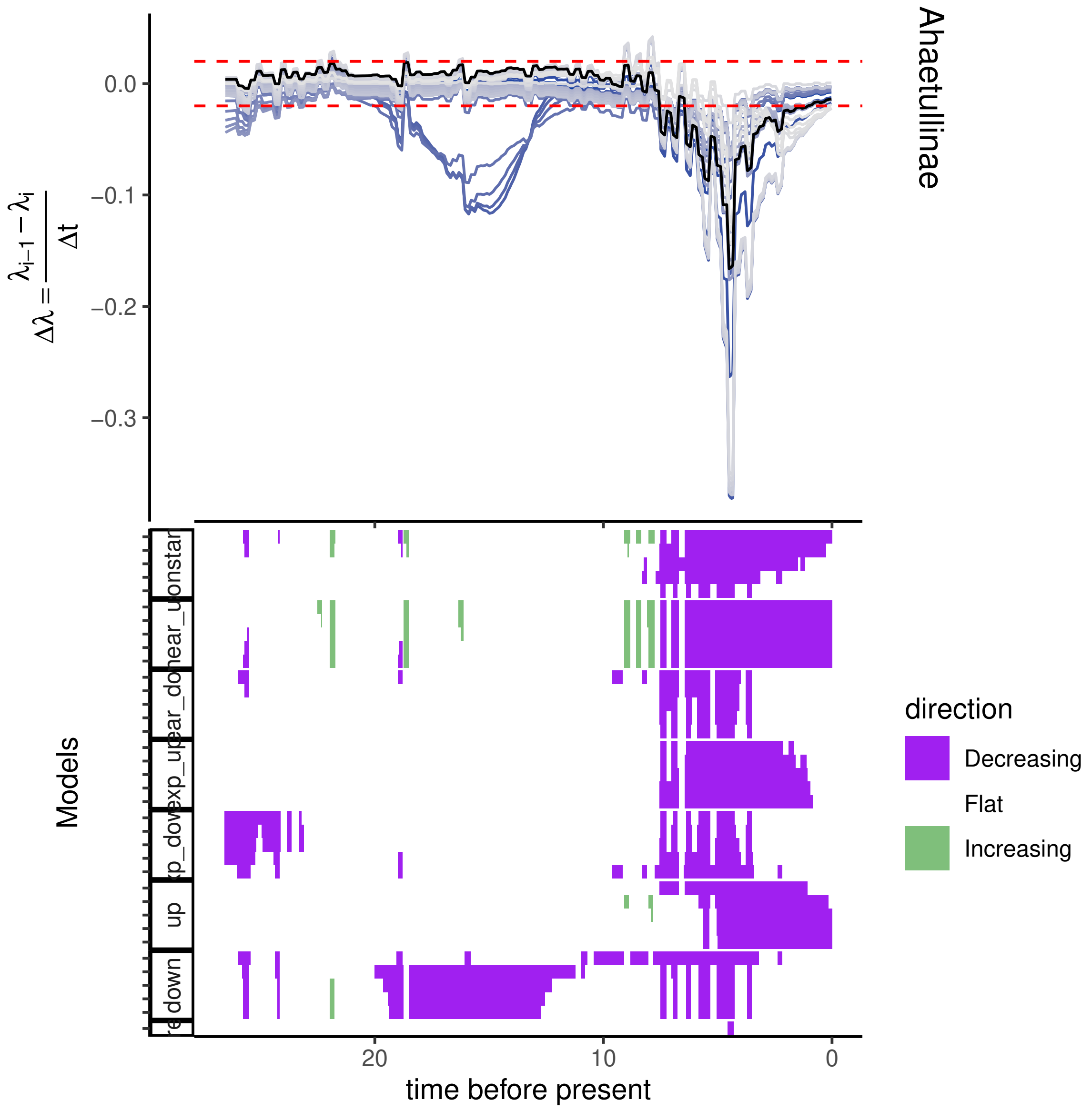
Ahaetullinae
$\Delta\lambda = \frac{\lambda_{i-1} - \lambda_i}{\Delta t}$
0.0
−0.1
−0.2
−0.3
Models
constant
linear_up
linear_down
exp_up
exp_down
up
down
20
10
0
time before present
direction
Decreasing
Flat
Increasing

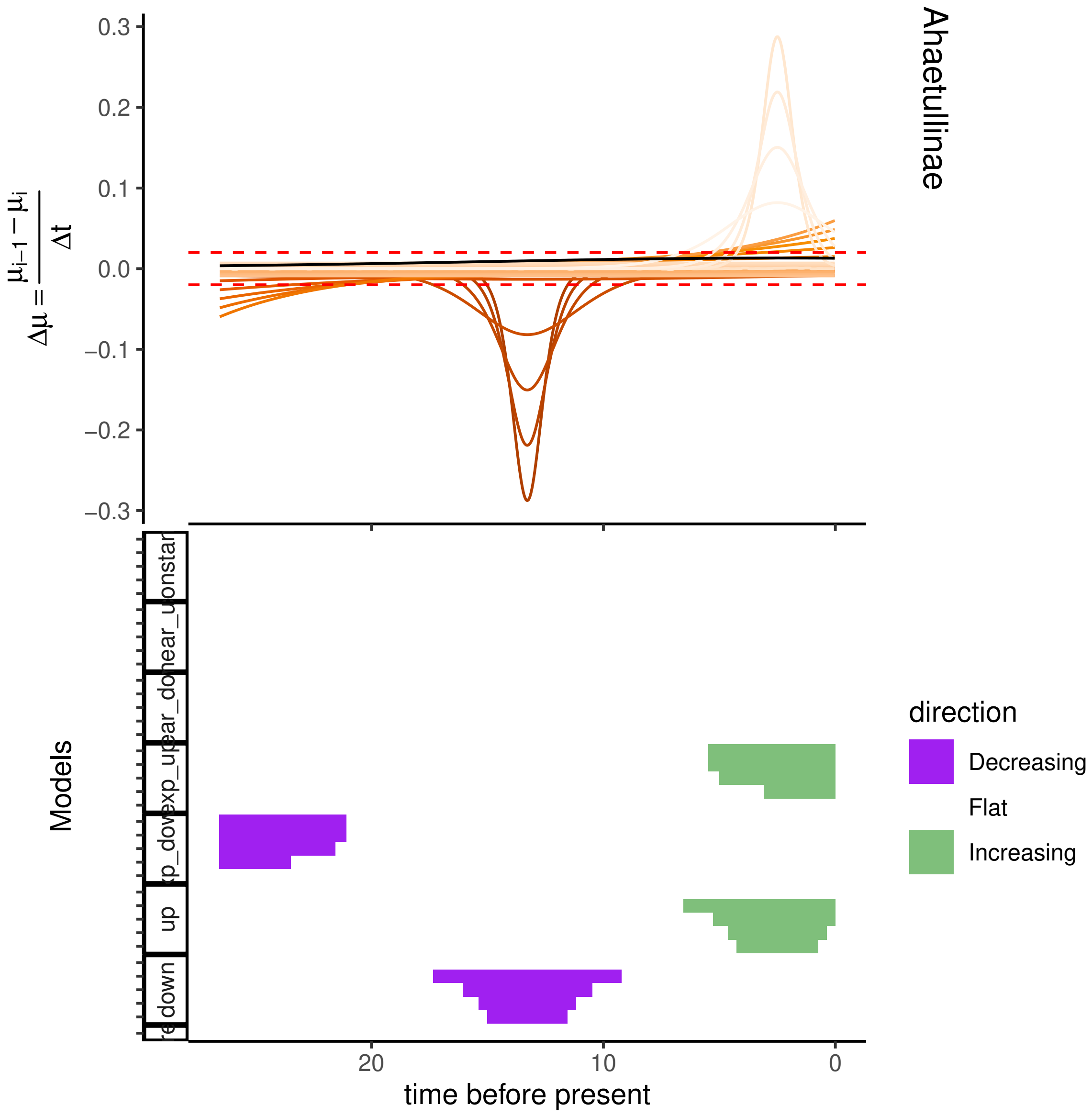
Ahaetullinae
0.3
0.2
0.1
0.0
−0.1
−0.2
−0.3
$\Delta\mu = \frac{\mu_{i-1} - \mu_i}{\Delta t}$
Models
direction
Decreasing
Flat
Increasing
20
10
0
time before present
up
down

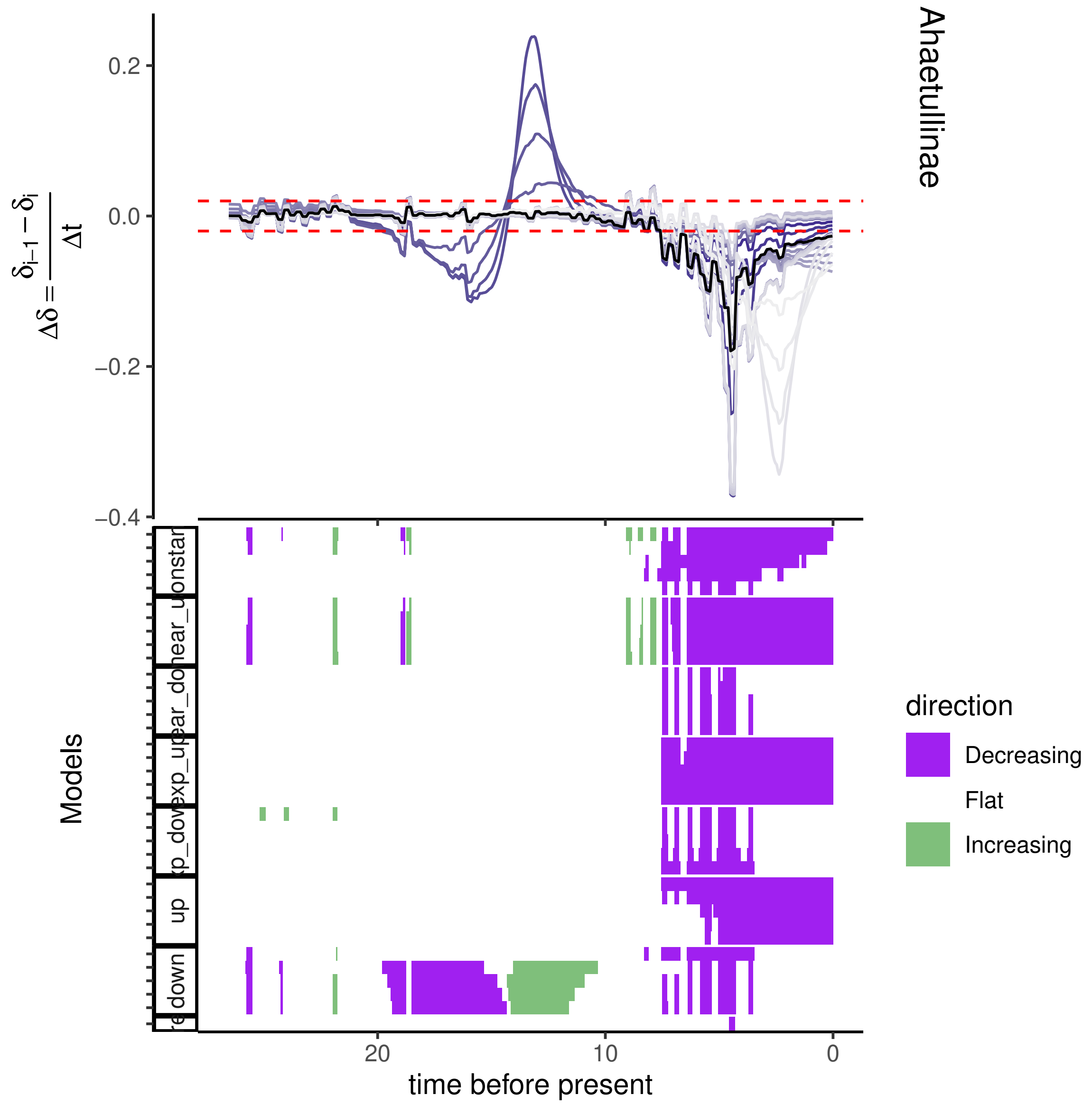

Ahaetullinae
$\Delta\delta = \frac{\delta_{i-1} - \delta_i}{\Delta t}$
0.2
0.0
−0.2
−0.4
Models
down
up
time before present
20
10
0
direction
Decreasing
Flat
Increasing

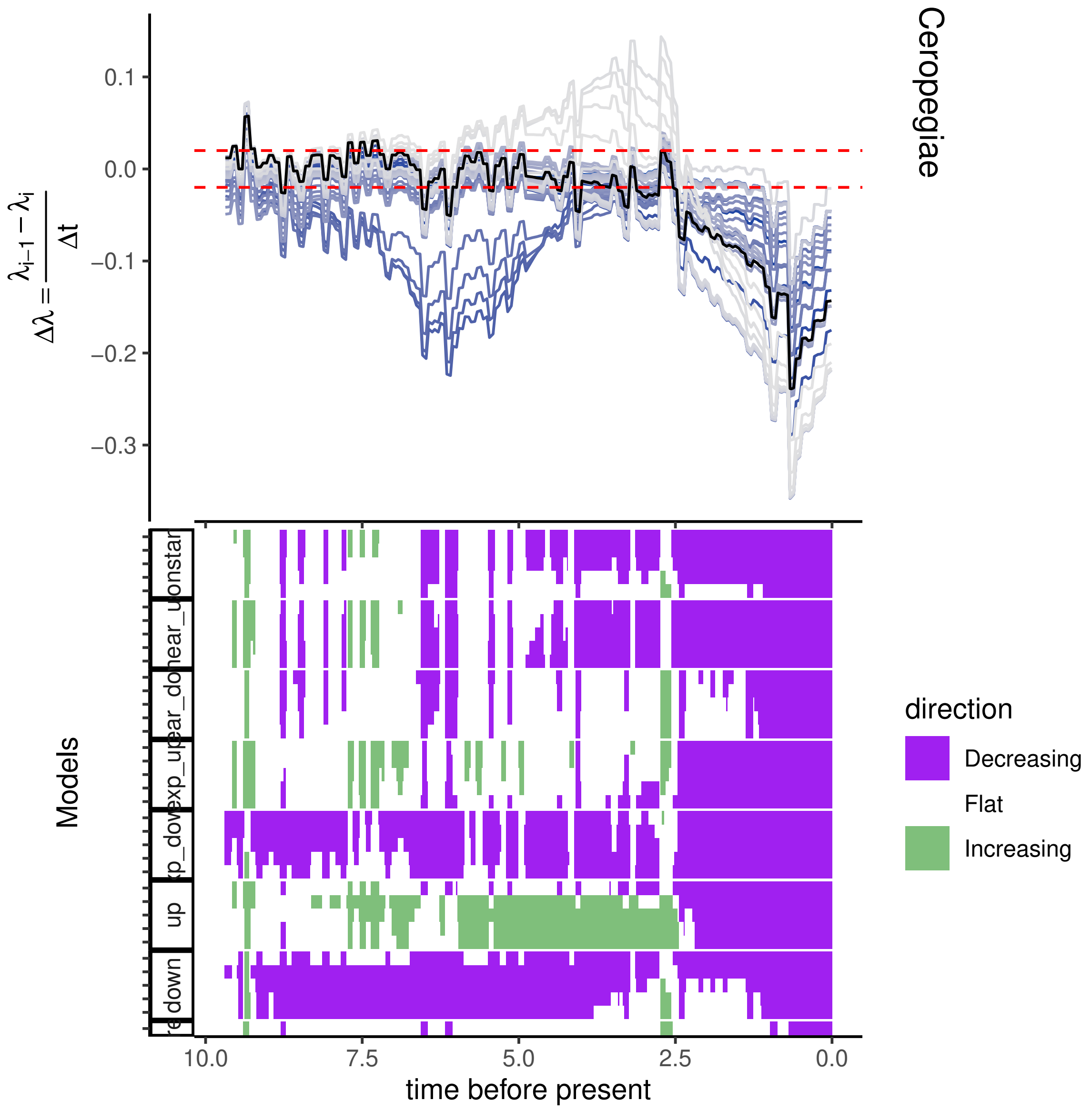
Ceropegiae
0.1
0.0
−0.1
−0.2
−0.3
$\Delta\lambda = \frac{\lambda_{i-1} - \lambda_i}{\Delta t}$
Models
direction
Decreasing
Flat
Increasing
up
down
10.0
7.5
5.0
2.5
0.0
time before present

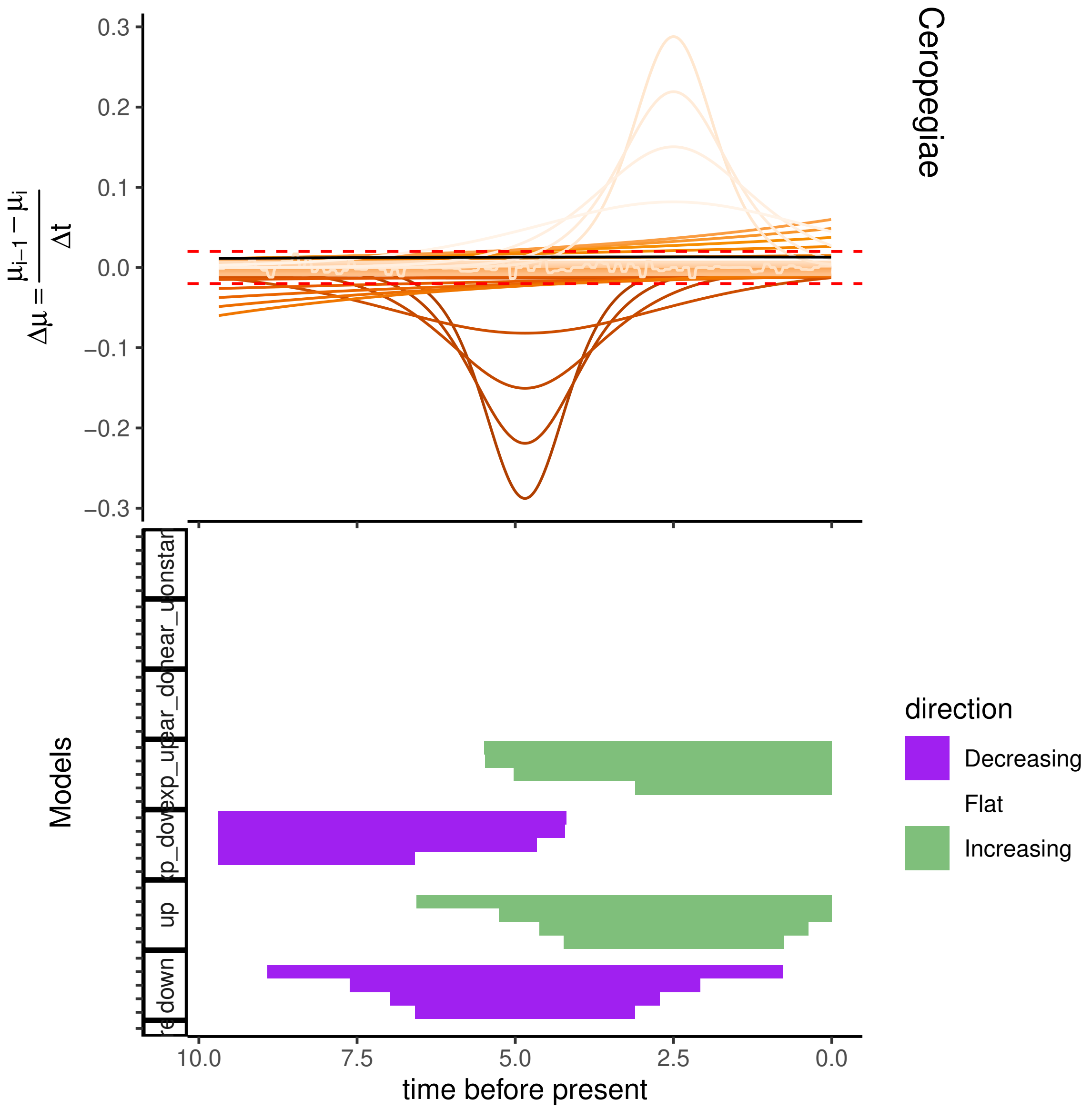

Cerופegiae

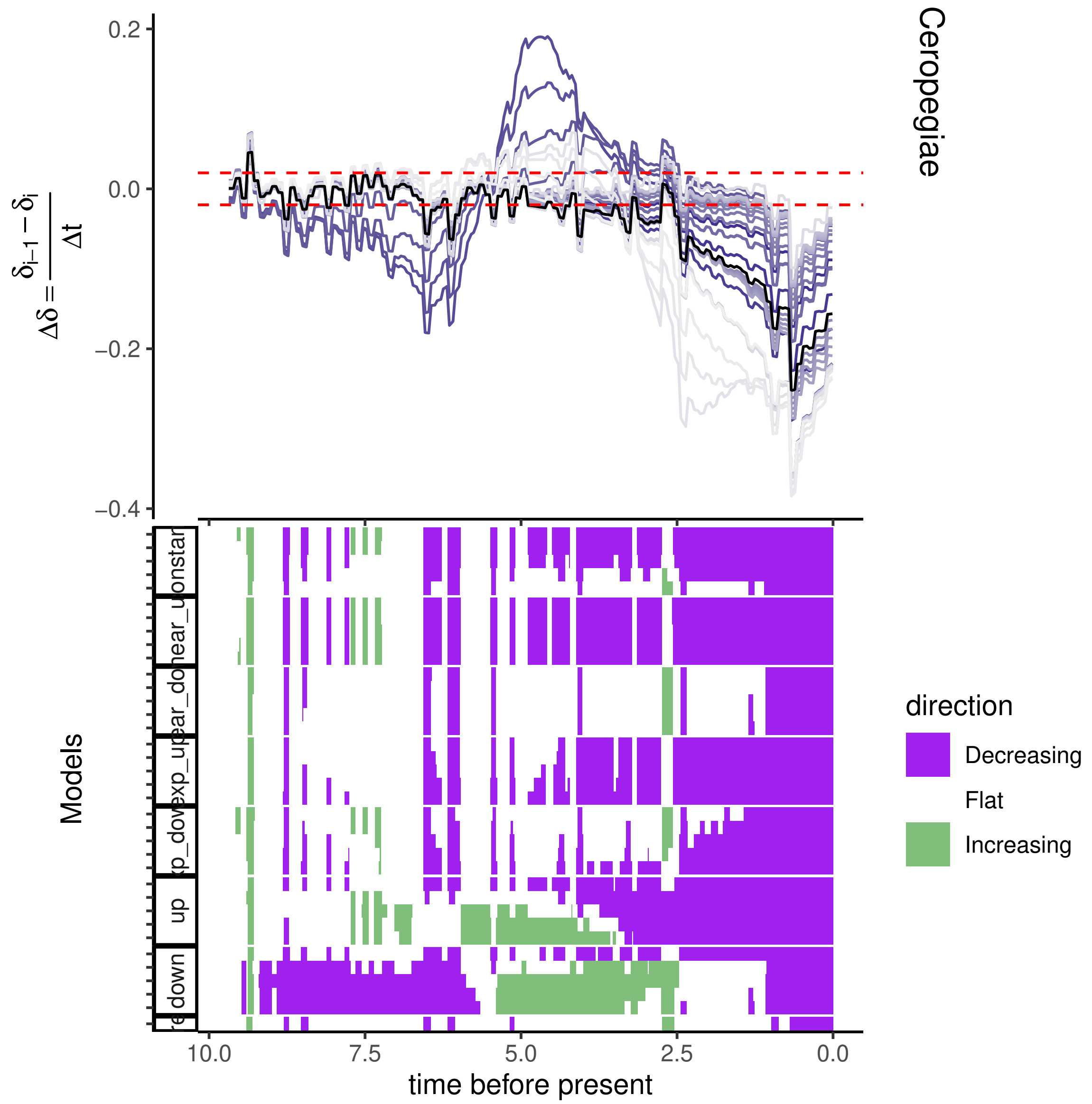
Cerepegiae
direction
Decreasing
Flat
Increasing
Models
up
down
time before present
10.0
7.5
5.0
2.5
0.0
0.2
0.0
−0.2
−0.4

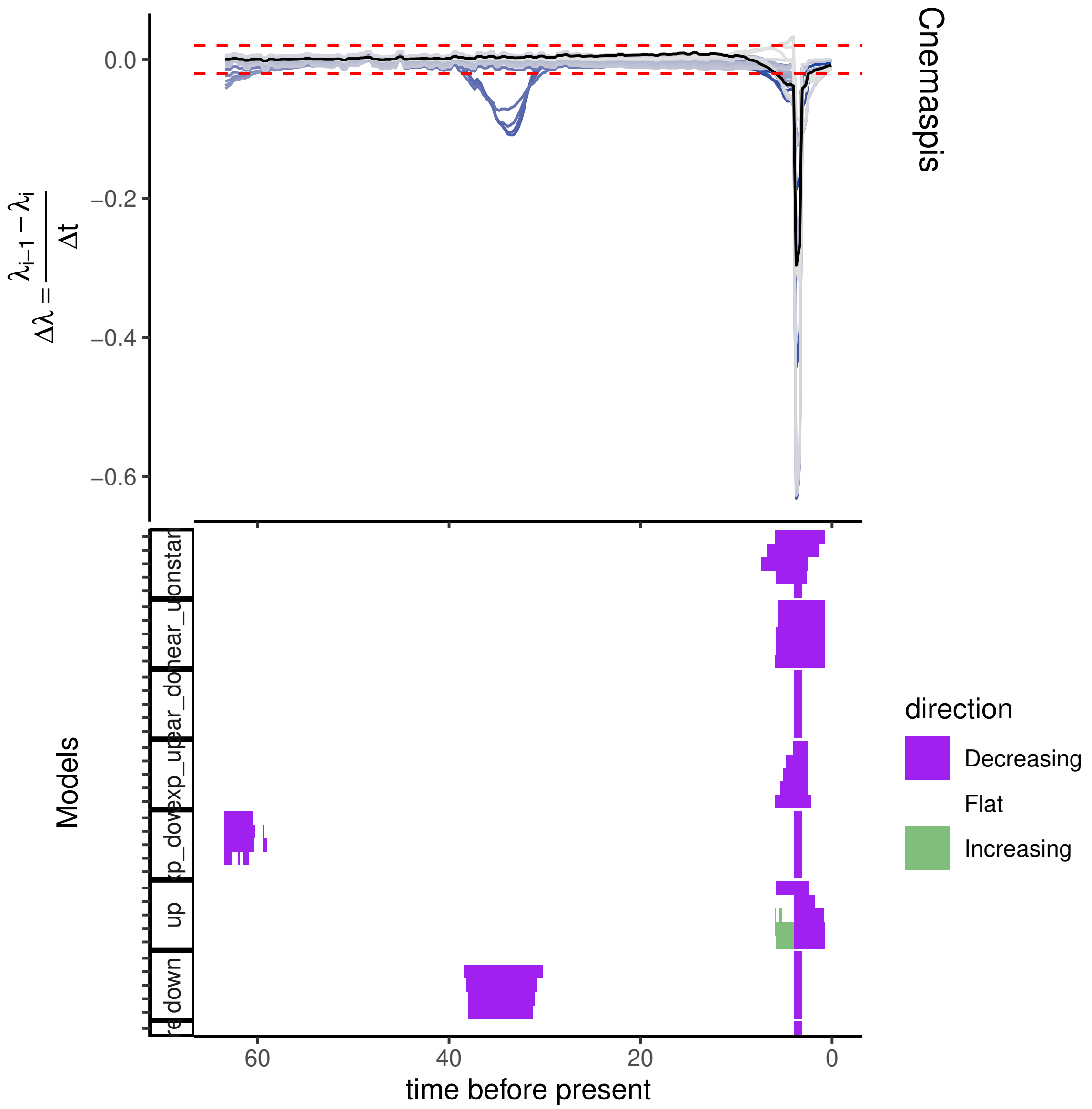

Cnemaspis
$\Delta\lambda = \frac{\lambda_{i-1} - \lambda_i}{\Delta t}$
0.0
−0.2
−0.4
−0.6
Models
up
down
direction
Decreasing
Flat
Increasing
60
40
20
0
time before present

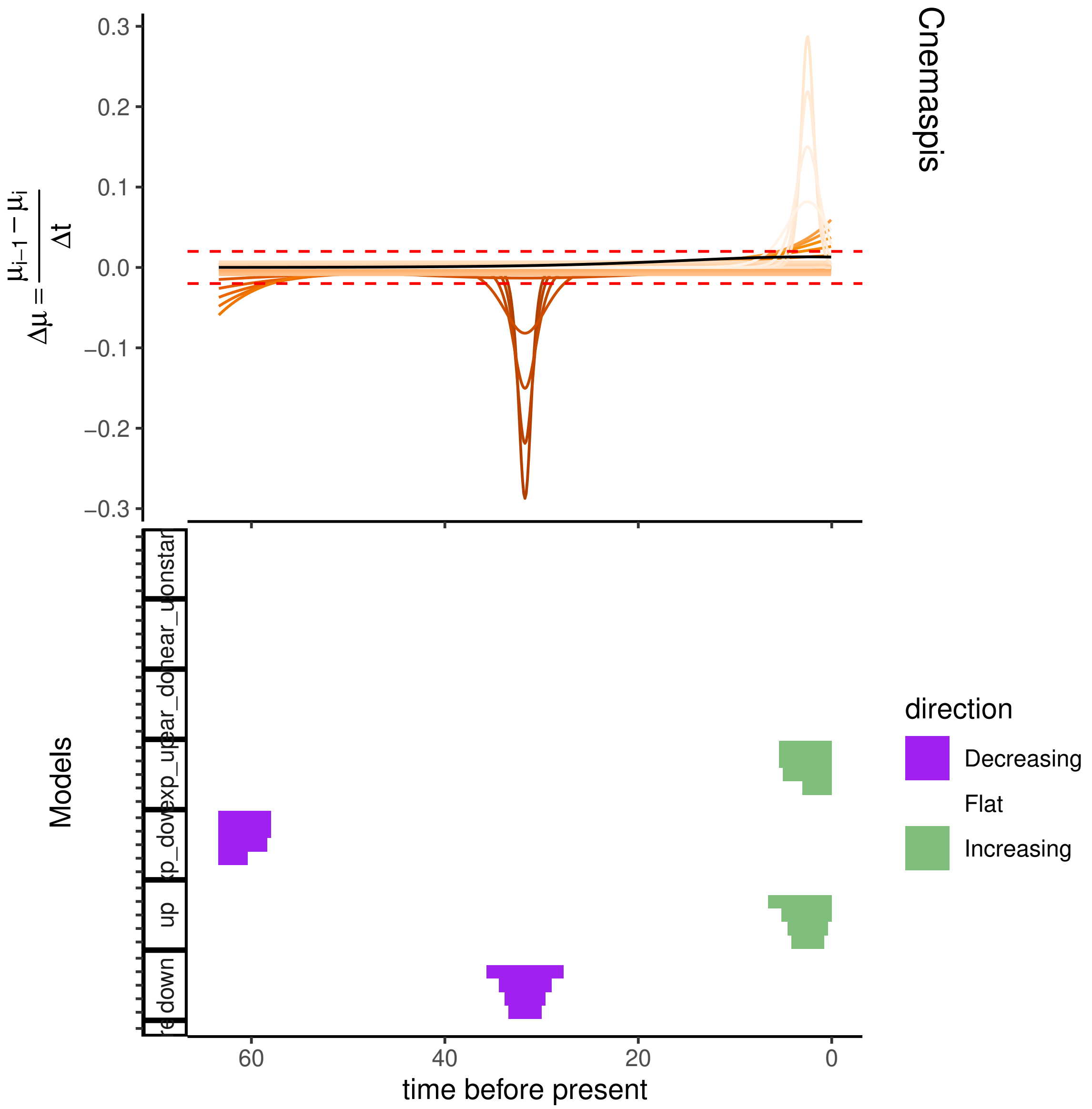

Cnemaspis
$\Delta\mu = \frac{\mu_{i-1} - \mu_i}{\Delta t}$
0.3
0.2
0.1
0.0
−0.1
−0.2
−0.3
Models
onstar
near_u
ear_do
exp_up
xp_dov
up
down
60
40
20
0
time before present
direction
Decreasing
Flat
Increasing

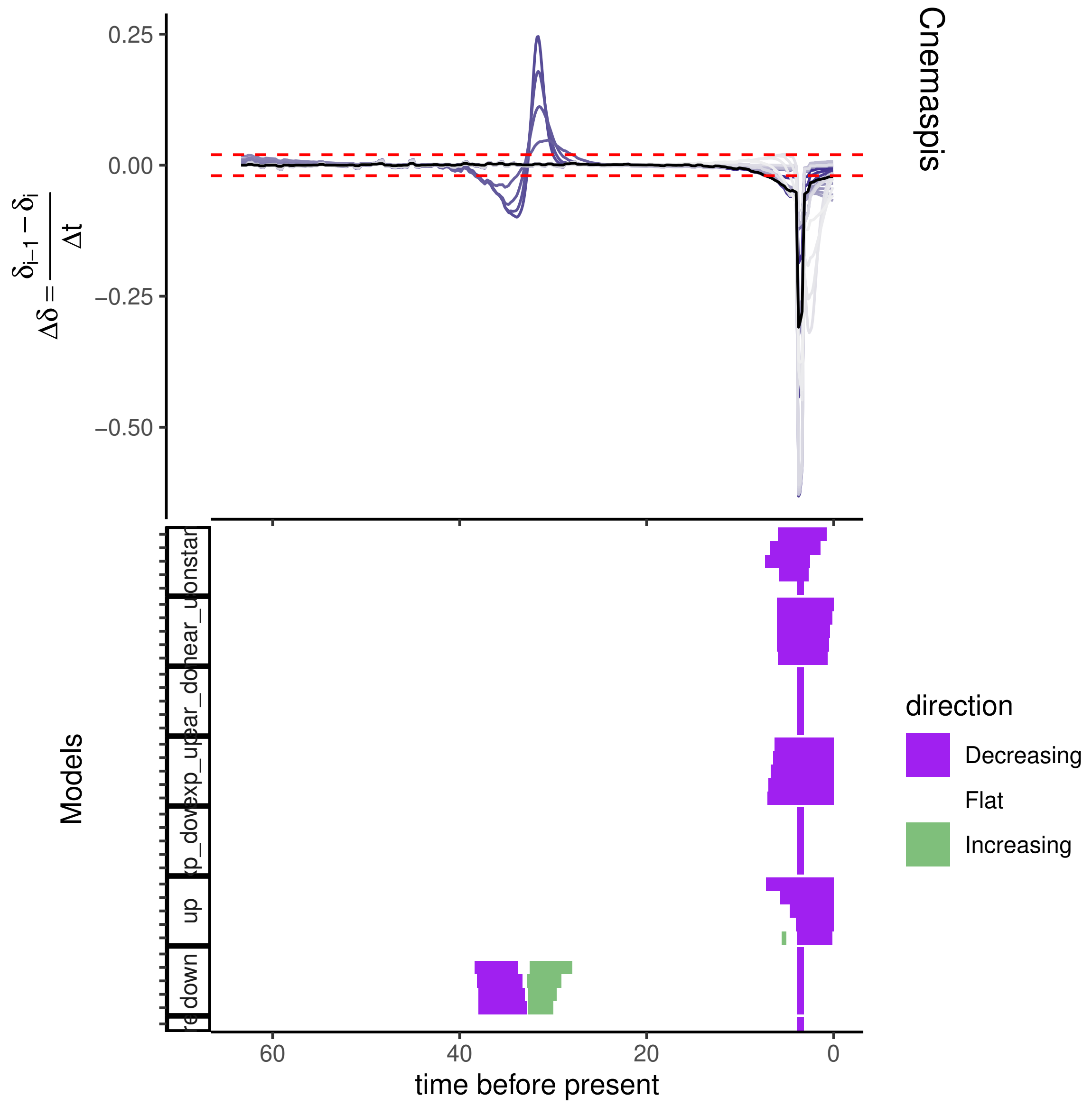

Cnemaspis
0.25
0.00
−0.25
−0.50
$\Delta\delta = \frac{\delta_{i-1} - \delta_i}{\Delta t}$
Models
down
up
direction
Decreasing
Flat
Increasing
60
40
20
0
time before present

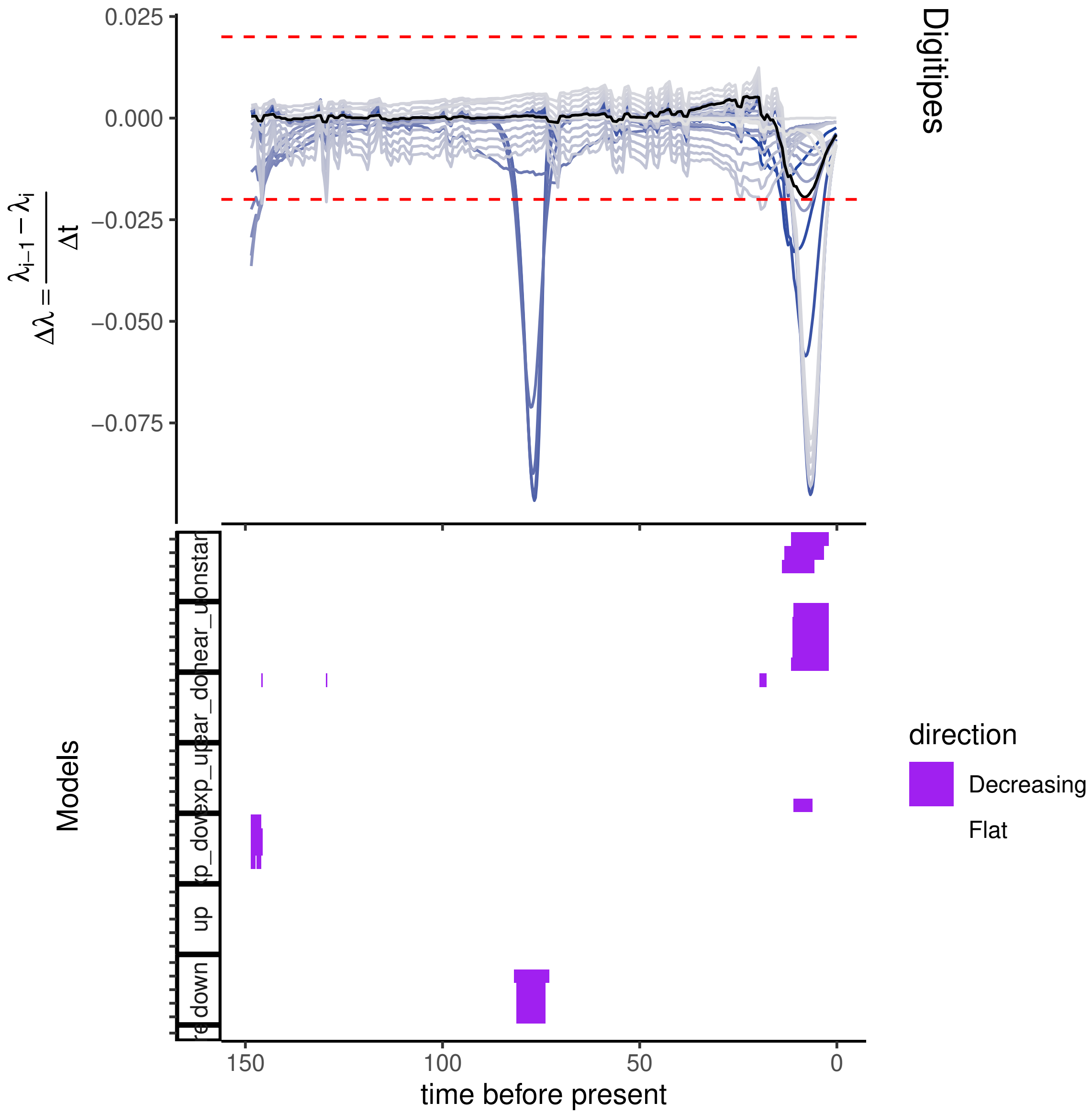

0.025
0.000
−0.025
−0.050
−0.075
$\Delta\lambda = \frac{\lambda_{i-1} - \lambda_i}{\Delta t}$
Digitipes
Models
direction
Decreasing
Flat
150
100
50
0
time before present

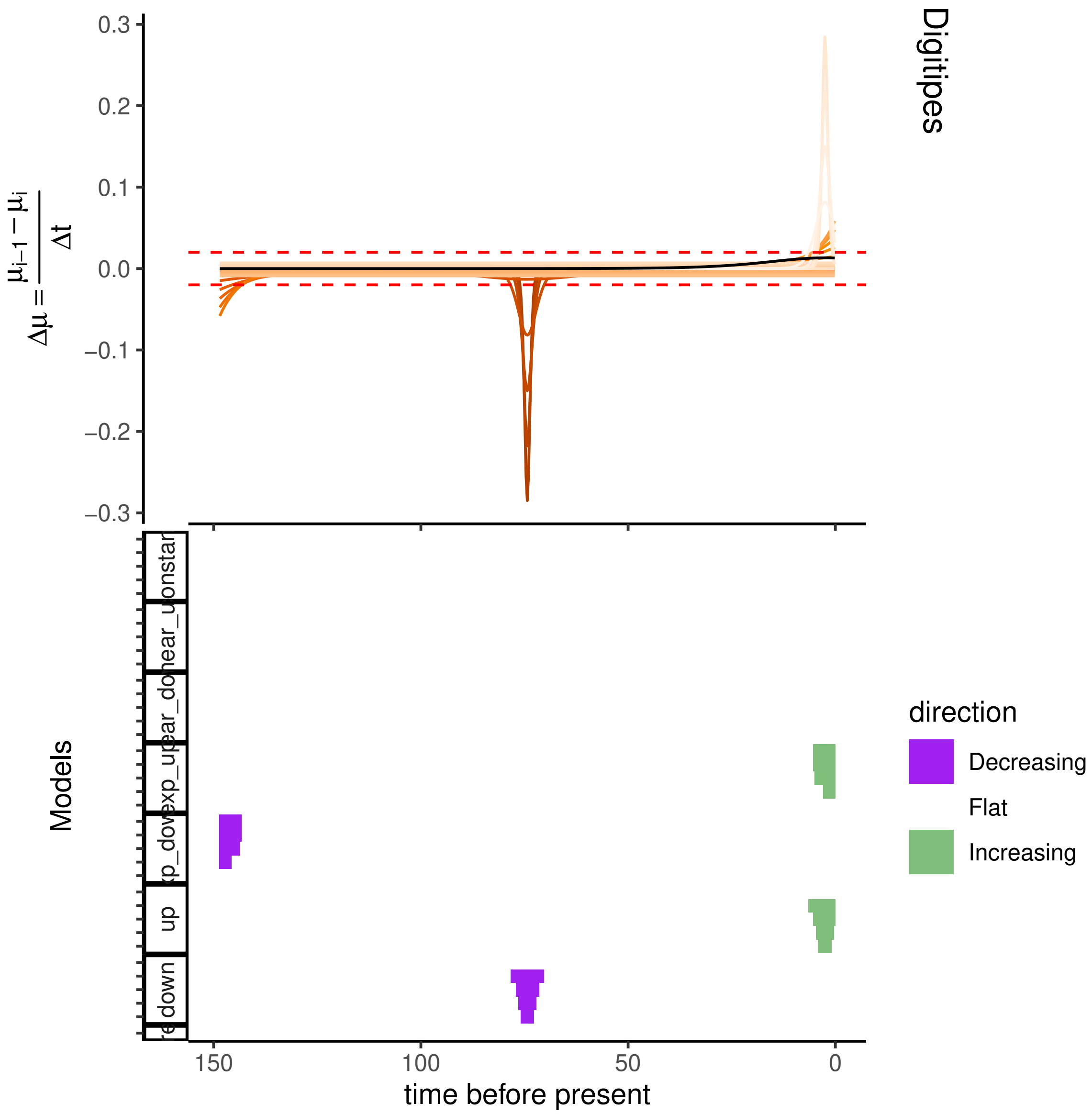

Digitipes
$\Delta\mu = \frac{\mu_{i-1} - \mu_i}{\Delta t}$
0.3
0.2
0.1
0.0
−0.1
−0.2
−0.3
Models
up
down
direction
Decreasing
Flat
Increasing
150
100
50
0
time before present

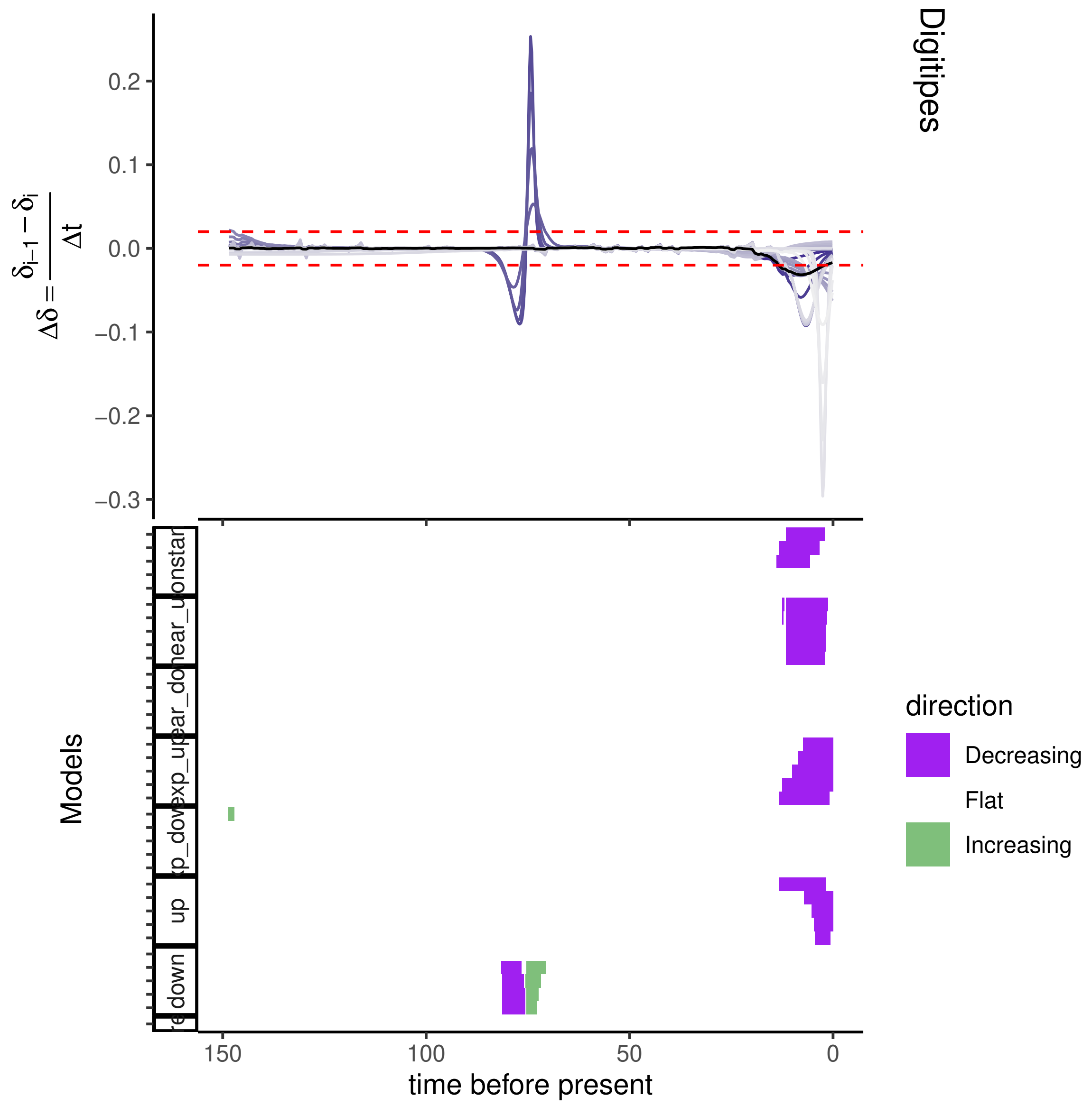

Digitipes
0.2
0.1
0.0
−0.1
−0.2
−0.3
Models
150
100
50
0
time before present
direction
Decreasing
Flat
Increasing
up
down

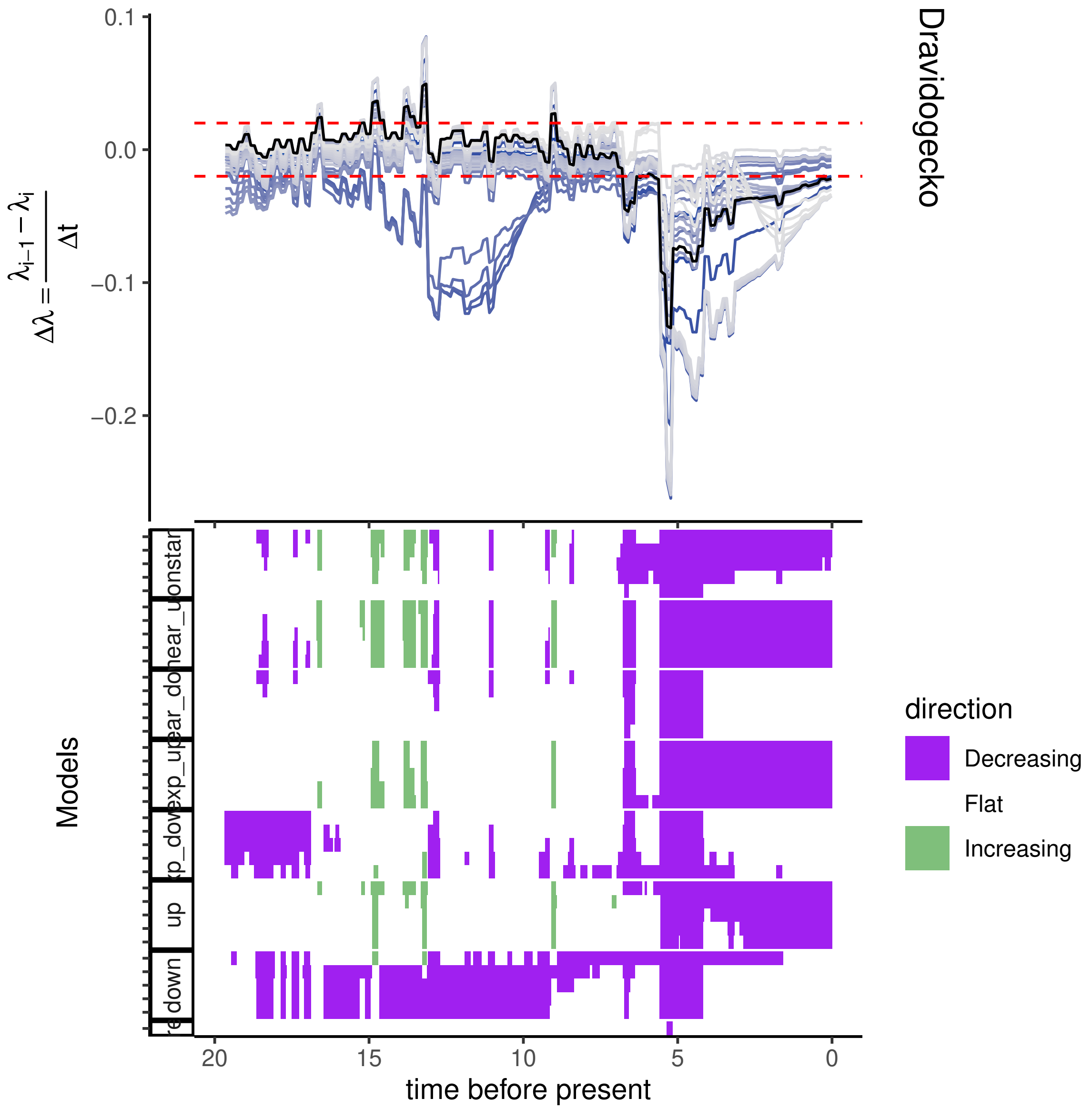

Dravidogecko
$\Delta\lambda = \frac{\lambda_{i-1} - \lambda_i}{\Delta t}$
0.1
0.0
−0.1
−0.2
Models
time before present
20
15
10
5
0
direction
Decreasing
Flat
Increasing

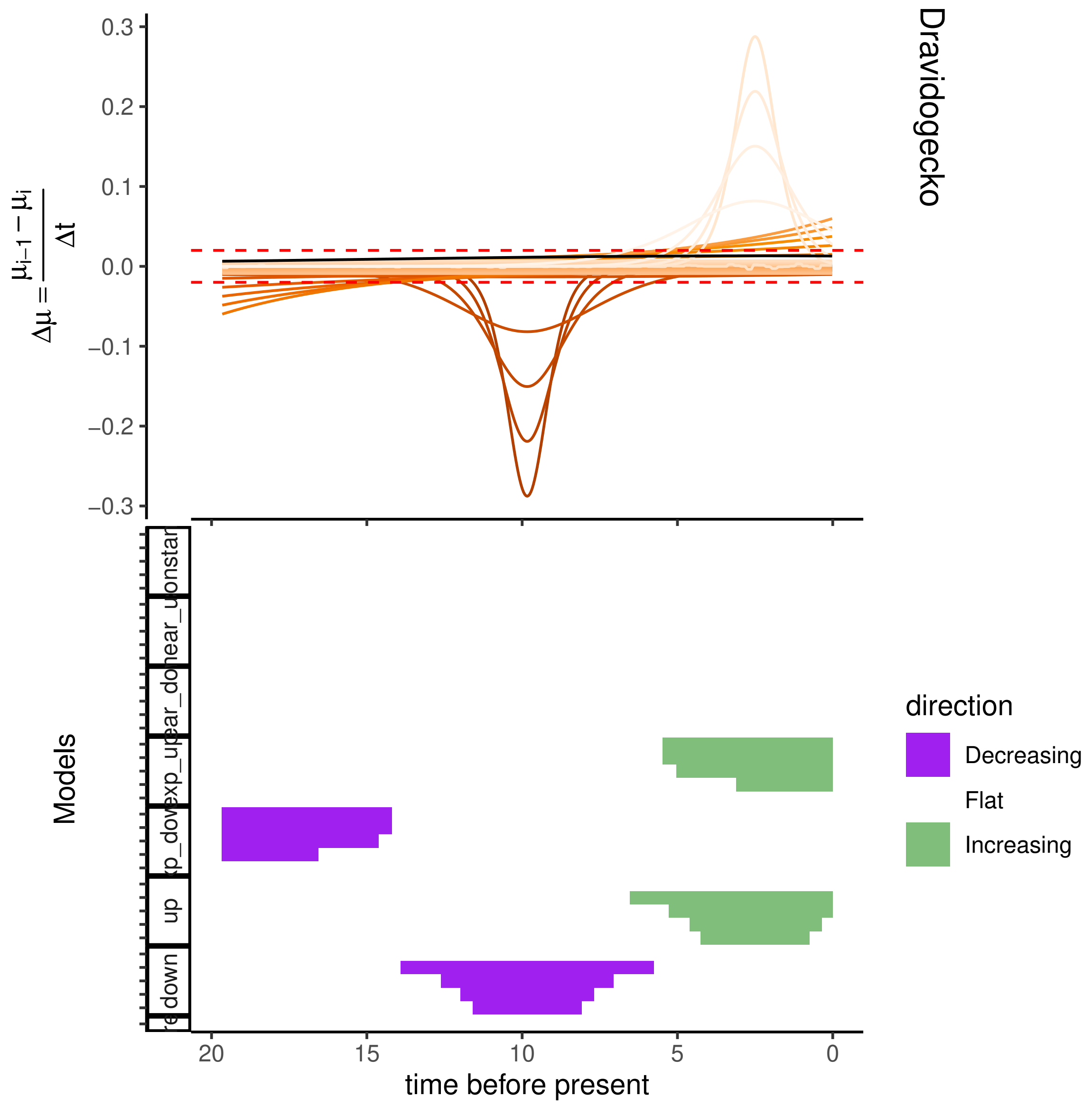

Dravidogecko
$\Delta\mu = \frac{\mu_{i-1} - \mu_i}{\Delta t}$
Models
time before present
direction
Decreasing
Flat
Increasing
up
down

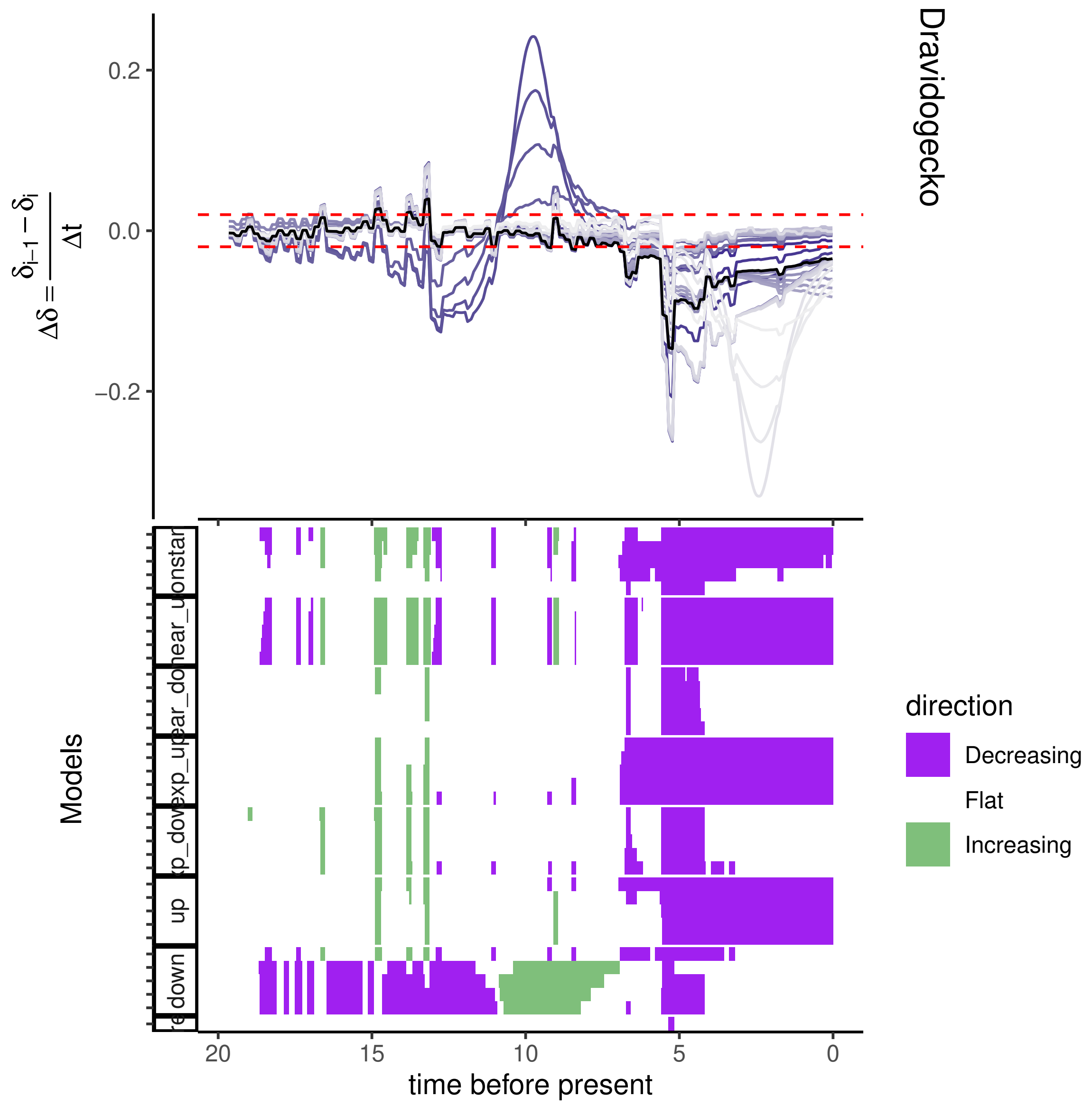
Dravidogecko
0.2
0.0
−0.2
$\Delta\delta = \frac{\delta_{i-1} - \delta_i}{\Delta t}$
Models
direction
Decreasing
Flat
Increasing
20
15
10
5
0
time before present
down
up

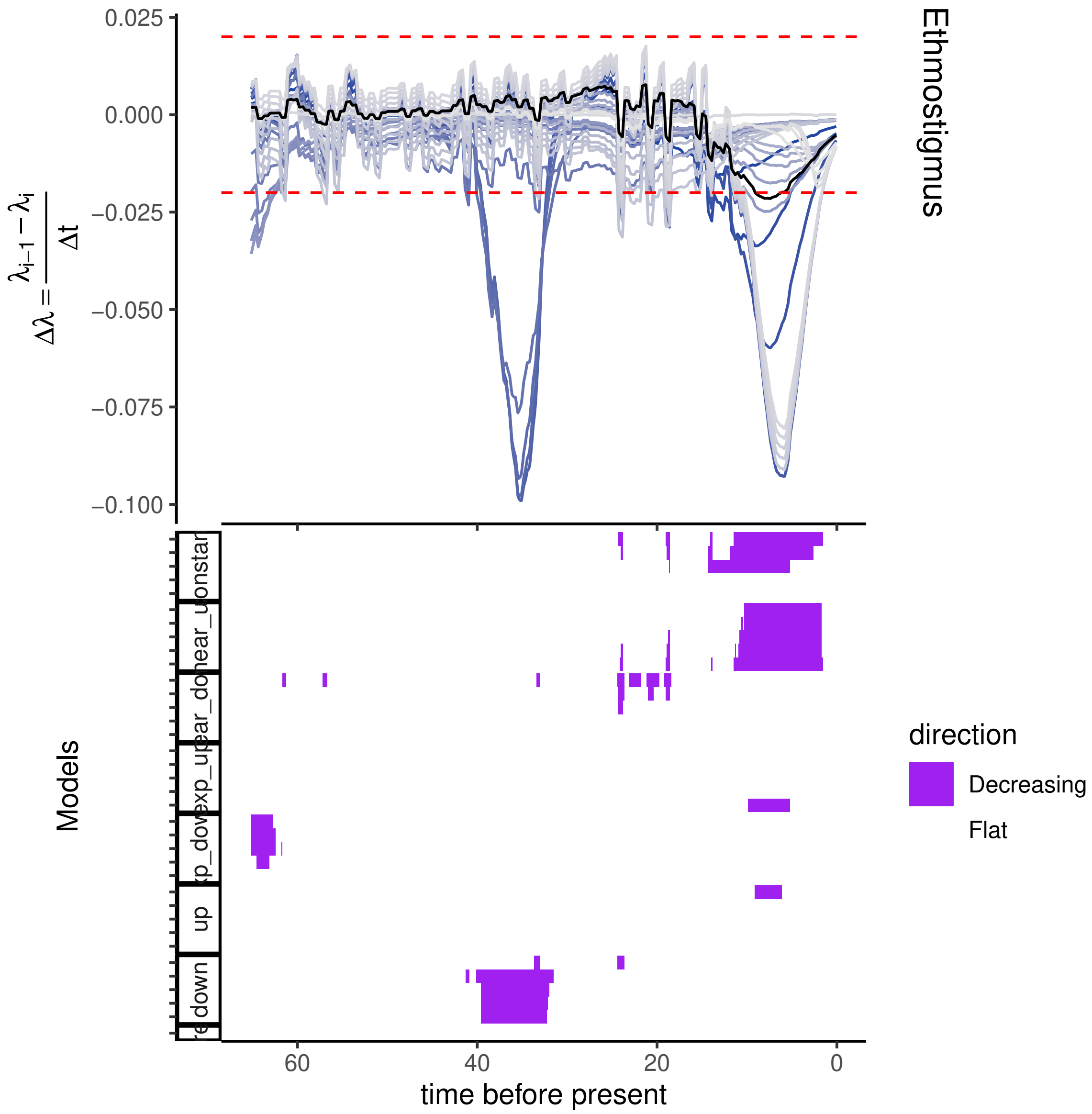

Ethmostigmus
0.025
0.000
−0.025
−0.050
−0.075
−0.100
$\Delta\lambda = \frac{\lambda_{i-1} - \lambda_i}{\Delta t}$
Models
up
down
direction
Decreasing
Flat
60
40
20
0
time before present

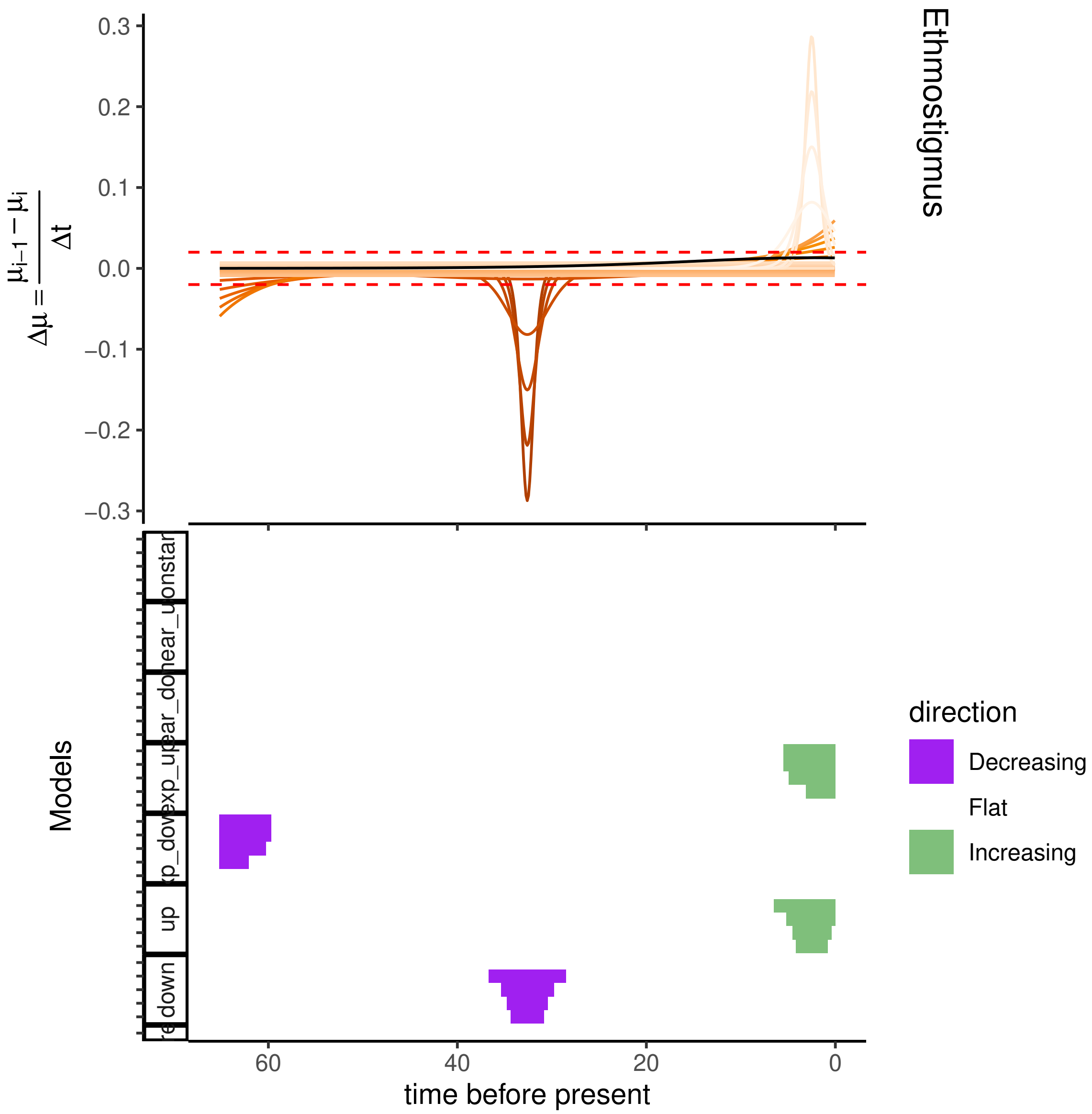

Ethmostigmus
$\Delta\mu = \frac{\mu_{i-1} - \mu_i}{\Delta t}$
0.3
0.2
0.1
0.0
−0.1
−0.2
−0.3
Models
direction
Decreasing
Flat
Increasing
60
40
20
0
time before present

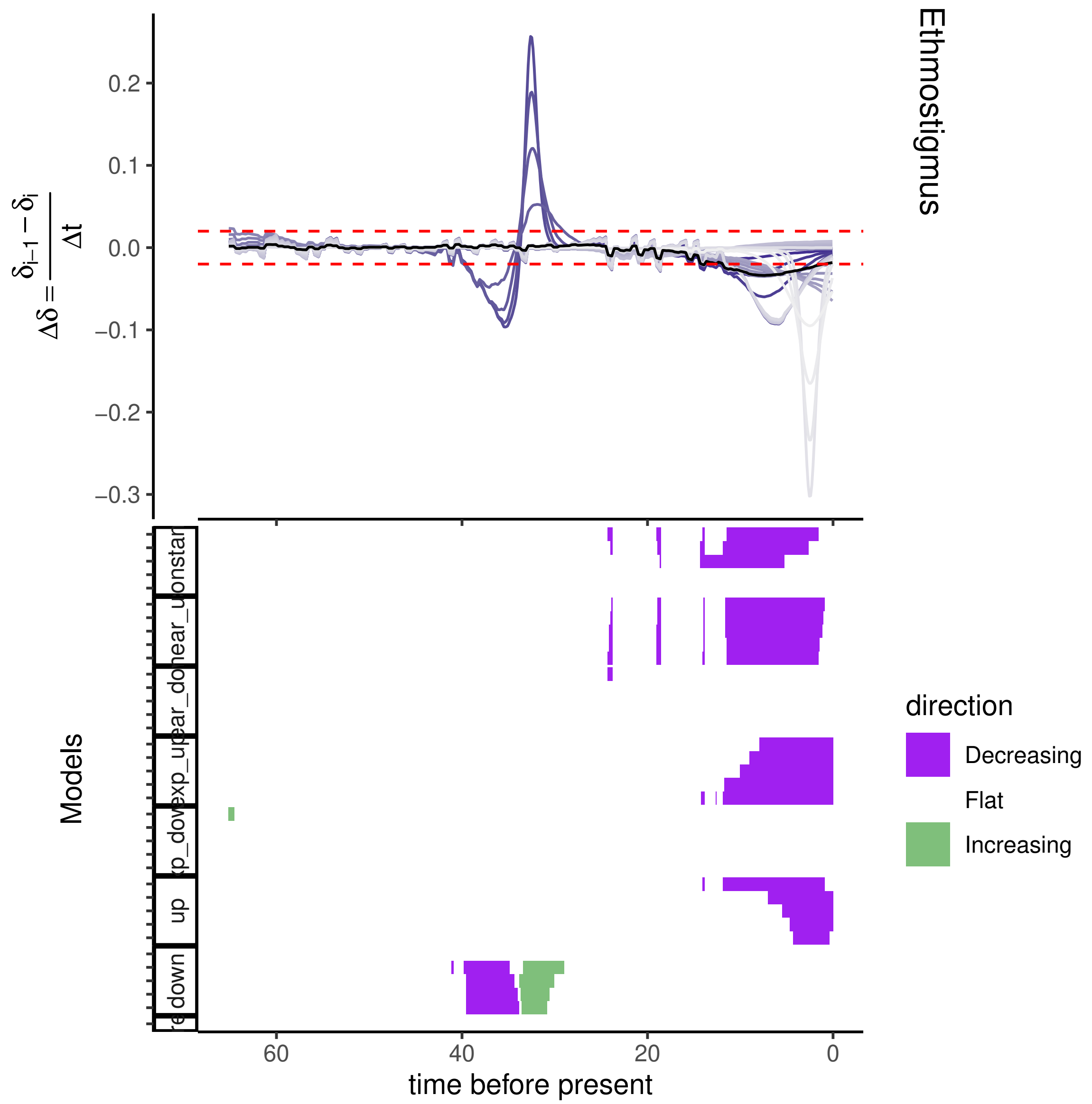
Ethmostigmus
$\Delta\delta = \frac{\delta_{i-1} - \delta_i}{\Delta t}$
0.2
0.1
0.0
−0.1
−0.2
−0.3
Models
constant
linear_down
linear_up
exp_down
exp_up
up
down
60
40
20
0
time before present
direction
Decreasing
Flat
Increasing

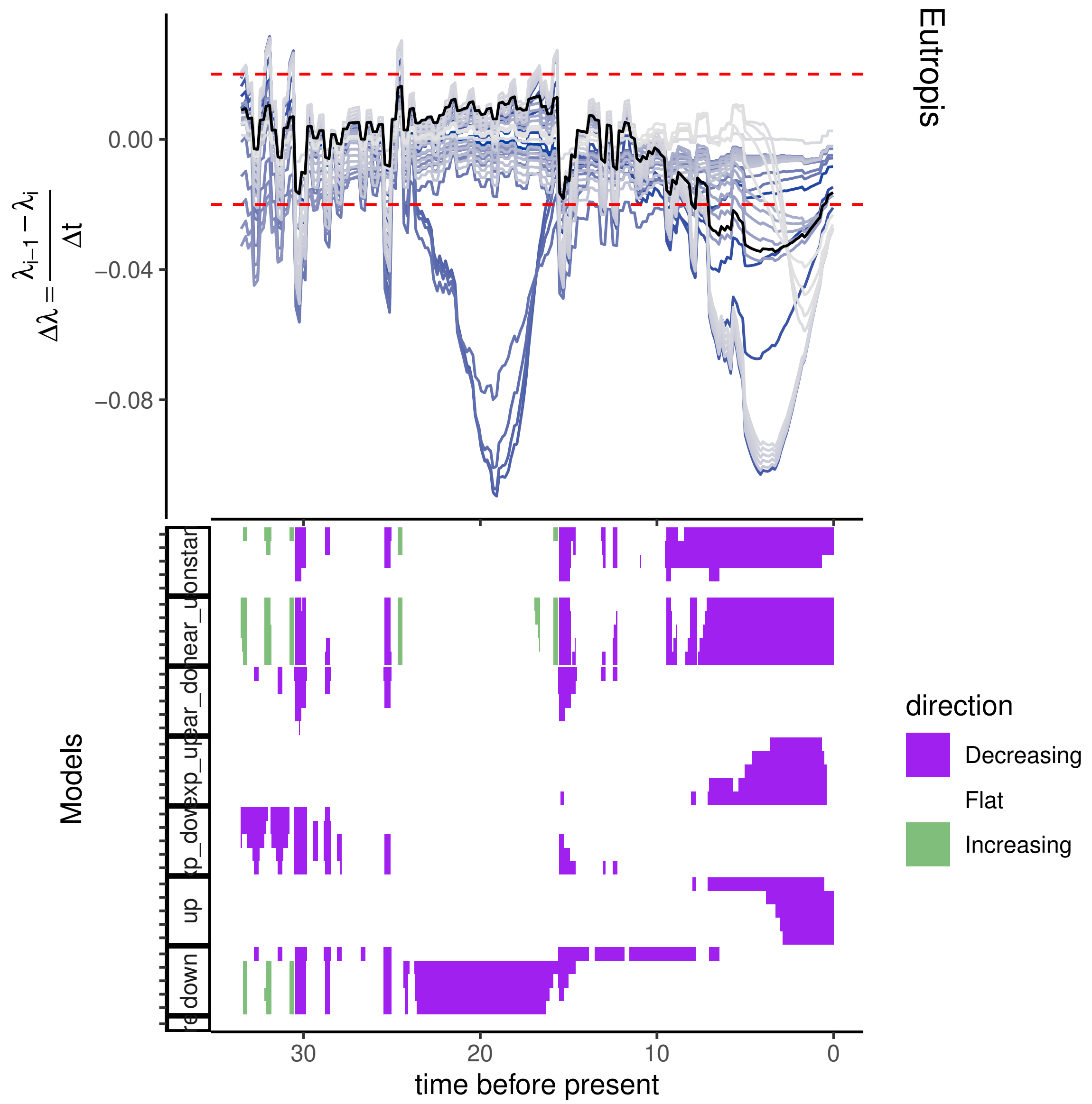
Eutropis
0.00
−0.04
−0.08
Δλ = (λ_{i−1} − λ_i)/Δt
Models
direction
Decreasing
Flat
Increasing
up
down
30
20
10
0
time before present

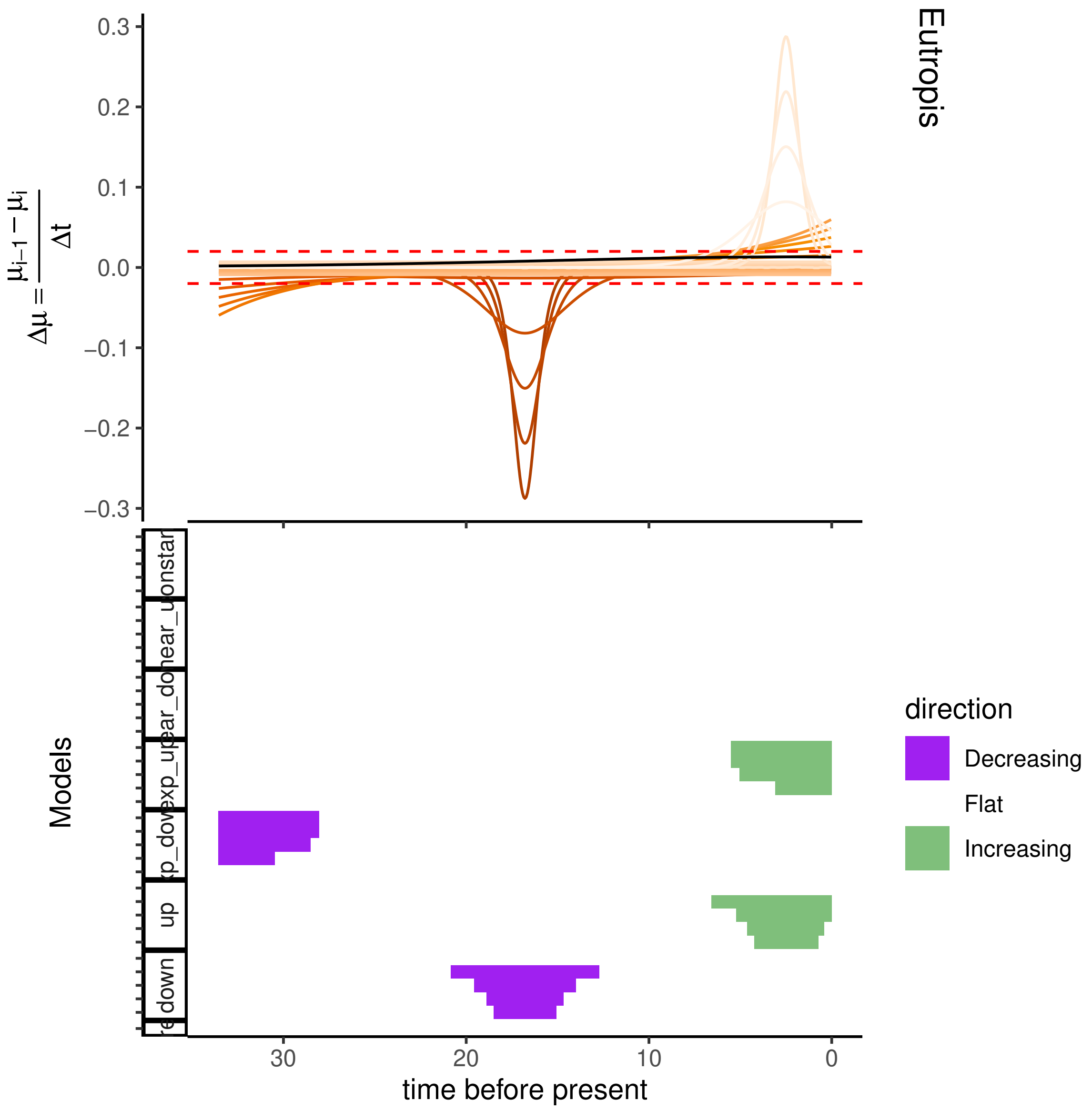

Eutropis
0.3
0.2
0.1
0.0
−0.1
−0.2
−0.3
$\Delta\mu = \frac{\mu_{i-1} - \mu_i}{\Delta t}$
Models
30
20
10
0
time before present
direction
Decreasing
Flat
Increasing

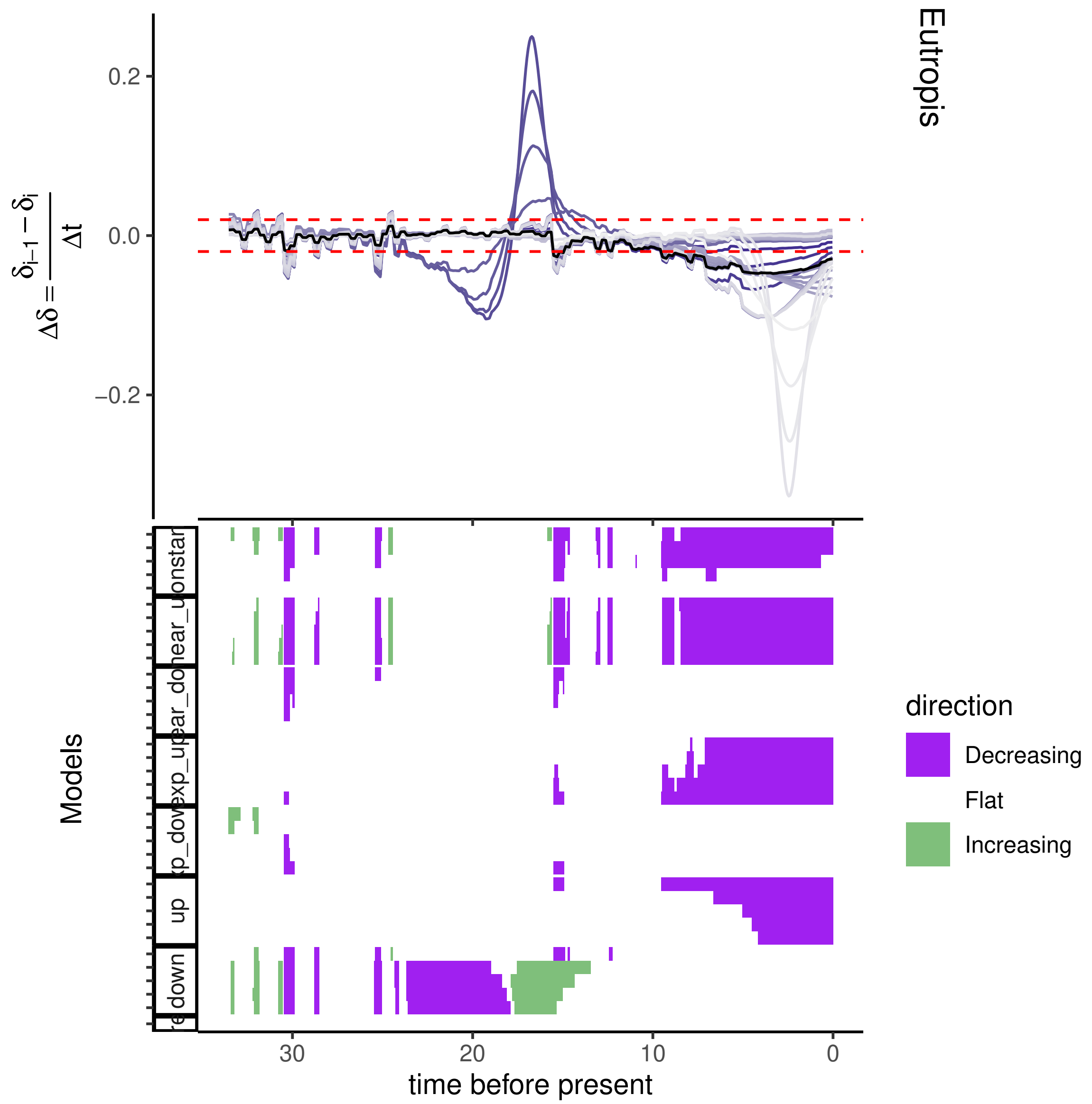
Eutropis
$\Delta\delta = \frac{\delta_{i-1} - \delta_i}{\Delta t}$
0.2
0.0
−0.2
Models
direction
Decreasing
Flat
Increasing
30
20
10
0
time before present

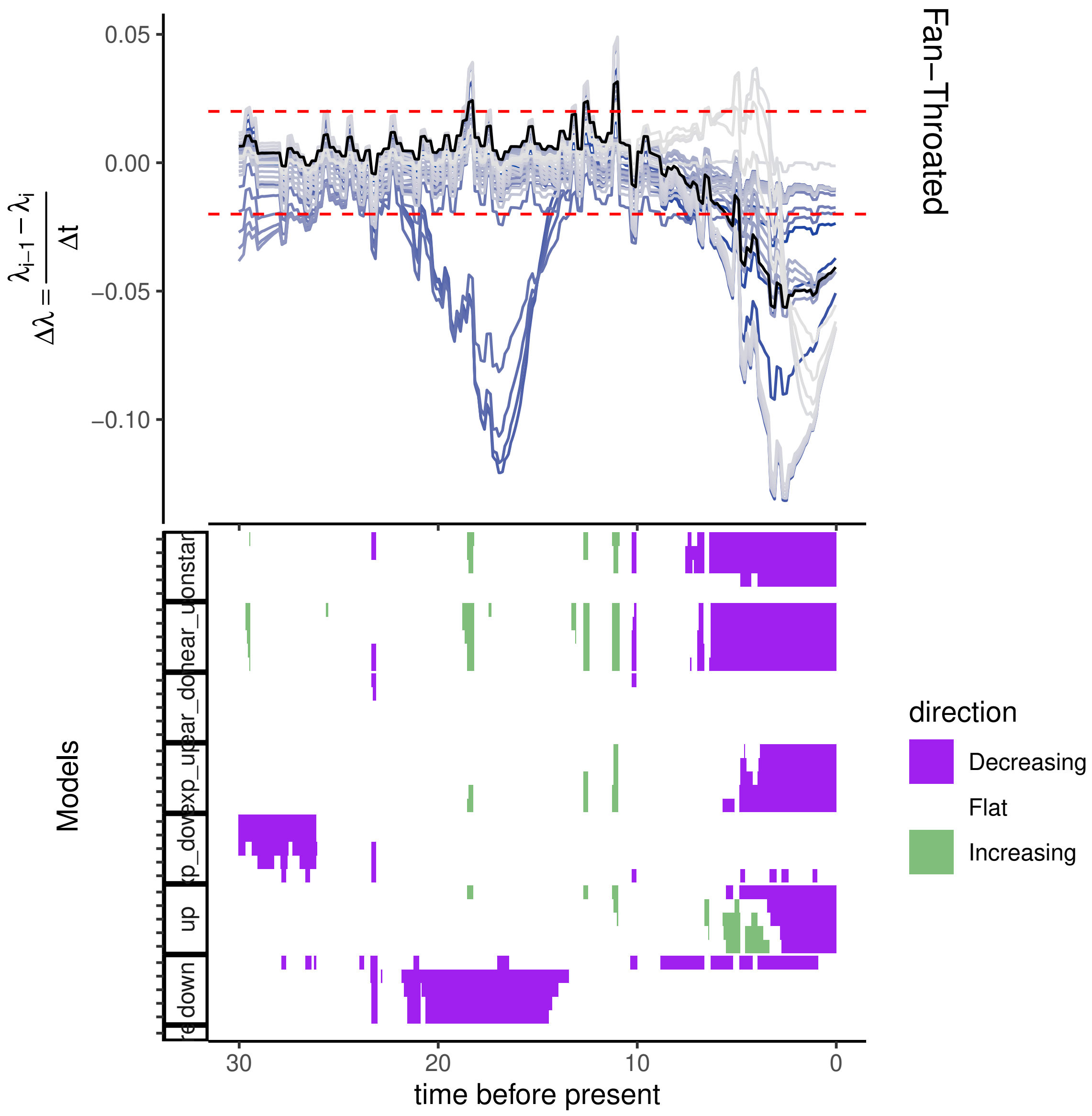
Fan−Throated
0.05
0.00
−0.05
−0.10
$\Delta\lambda = \frac{\lambda_{i-1} - \lambda_i}{\Delta t}$
Models
direction
Decreasing
Flat
Increasing
30
20
10
0
time before present
down
up

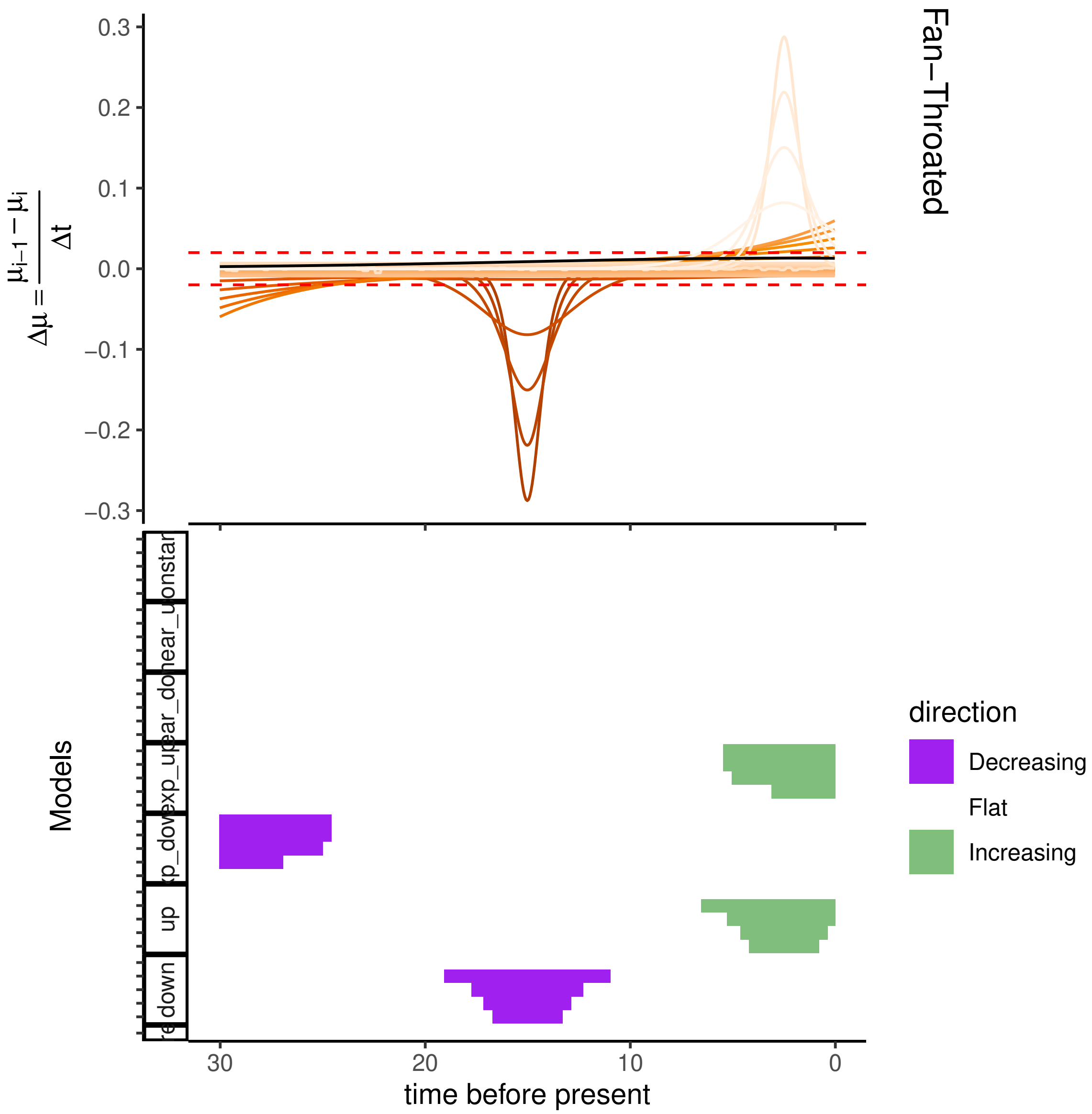

Fan–Throated
0.3
0.2
0.1
0.0
−0.1
−0.2
−0.3
Δμ = (μi−1 − μi)/Δt
Models
up
down
direction
Decreasing
Flat
Increasing
30
20
10
0
time before present

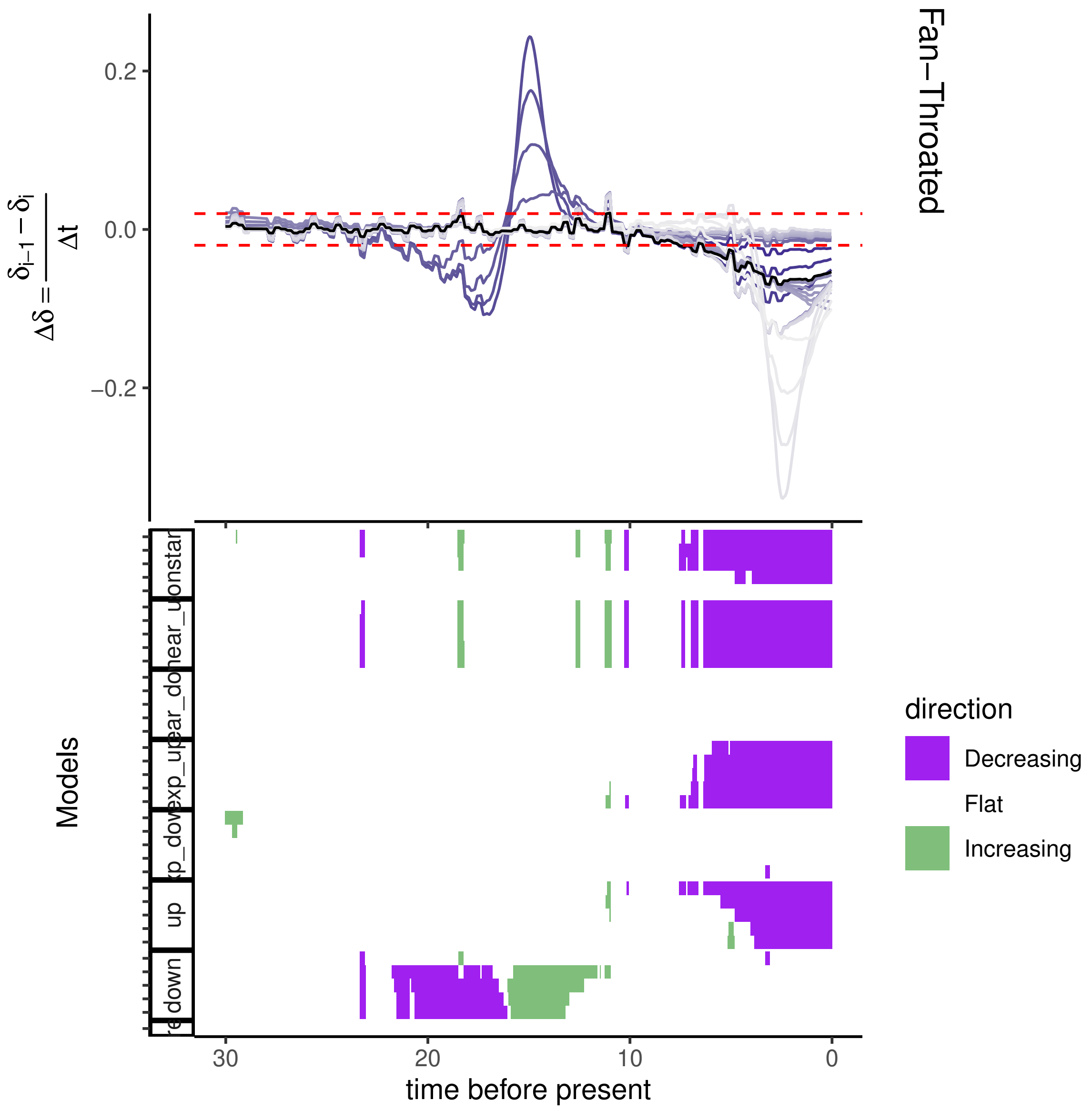
Fan−Throated
0.2
0.0
−0.2
Models
up
down
direction
Decreasing
Flat
Increasing
30
20
10
0
time before present

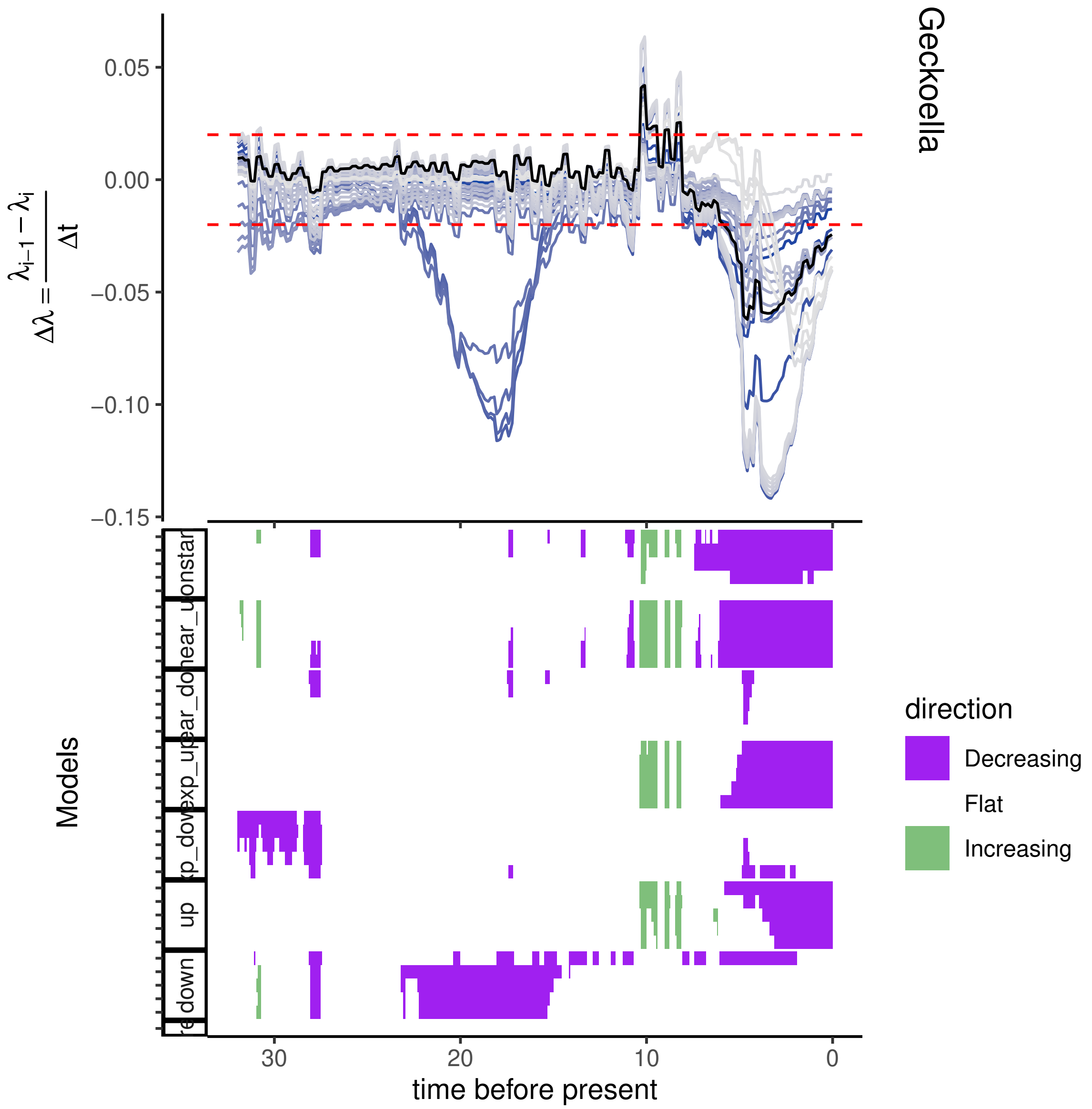
Geckoella
0.05
0.00
−0.05
−0.10
−0.15
$\Delta\lambda = \frac{\lambda_{i-1} - \lambda_i}{\Delta t}$
Models
up
down
30
20
10
0
time before present
direction
Decreasing
Flat
Increasing

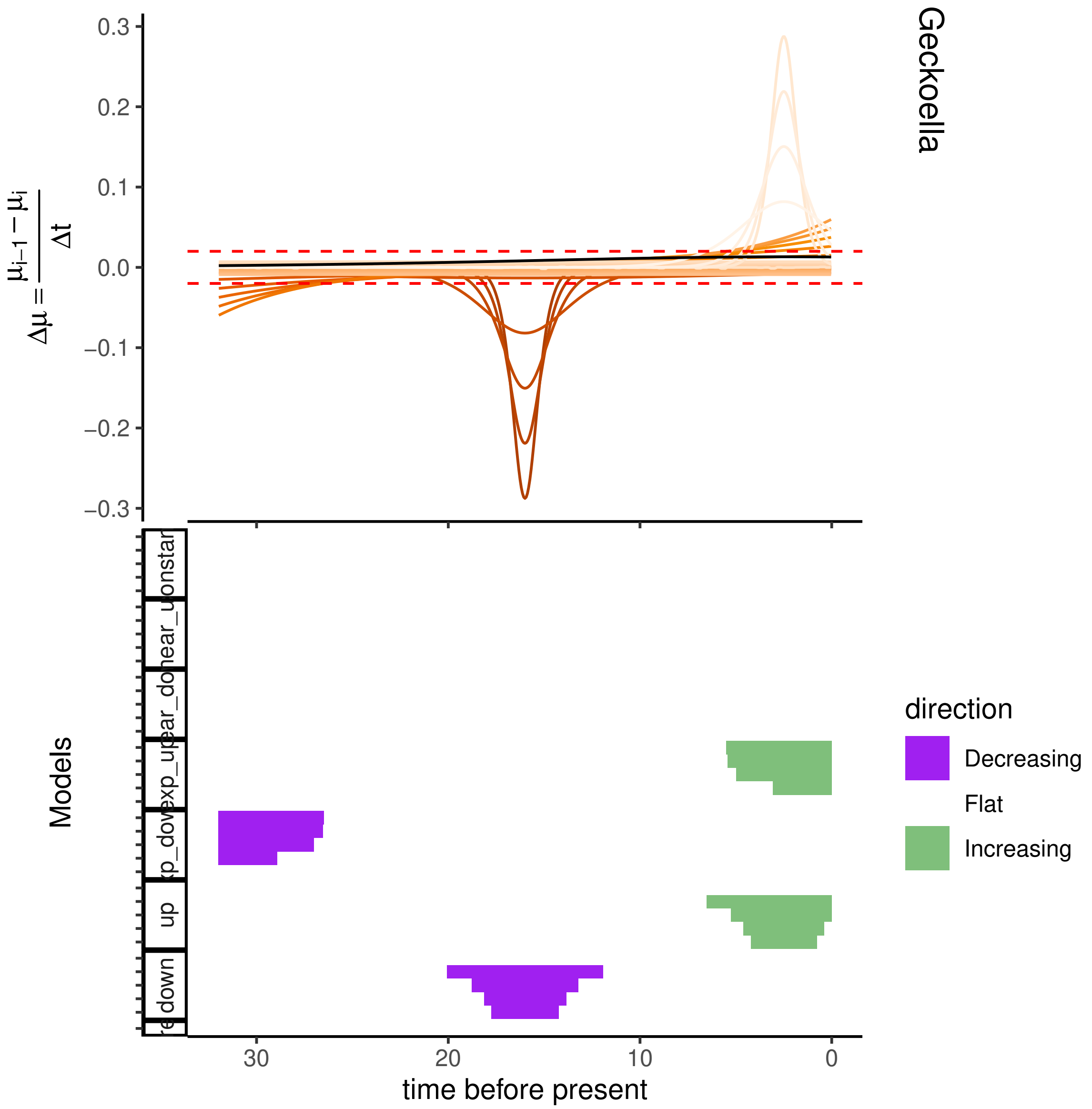

Geckoella
$\Delta\mu = \frac{\mu_{i-1} - \mu_i}{\Delta t}$
0.3
0.2
0.1
0.0
−0.1
−0.2
−0.3
Models
down
up
time before present
30
20
10
0
direction
Decreasing
Flat
Increasing

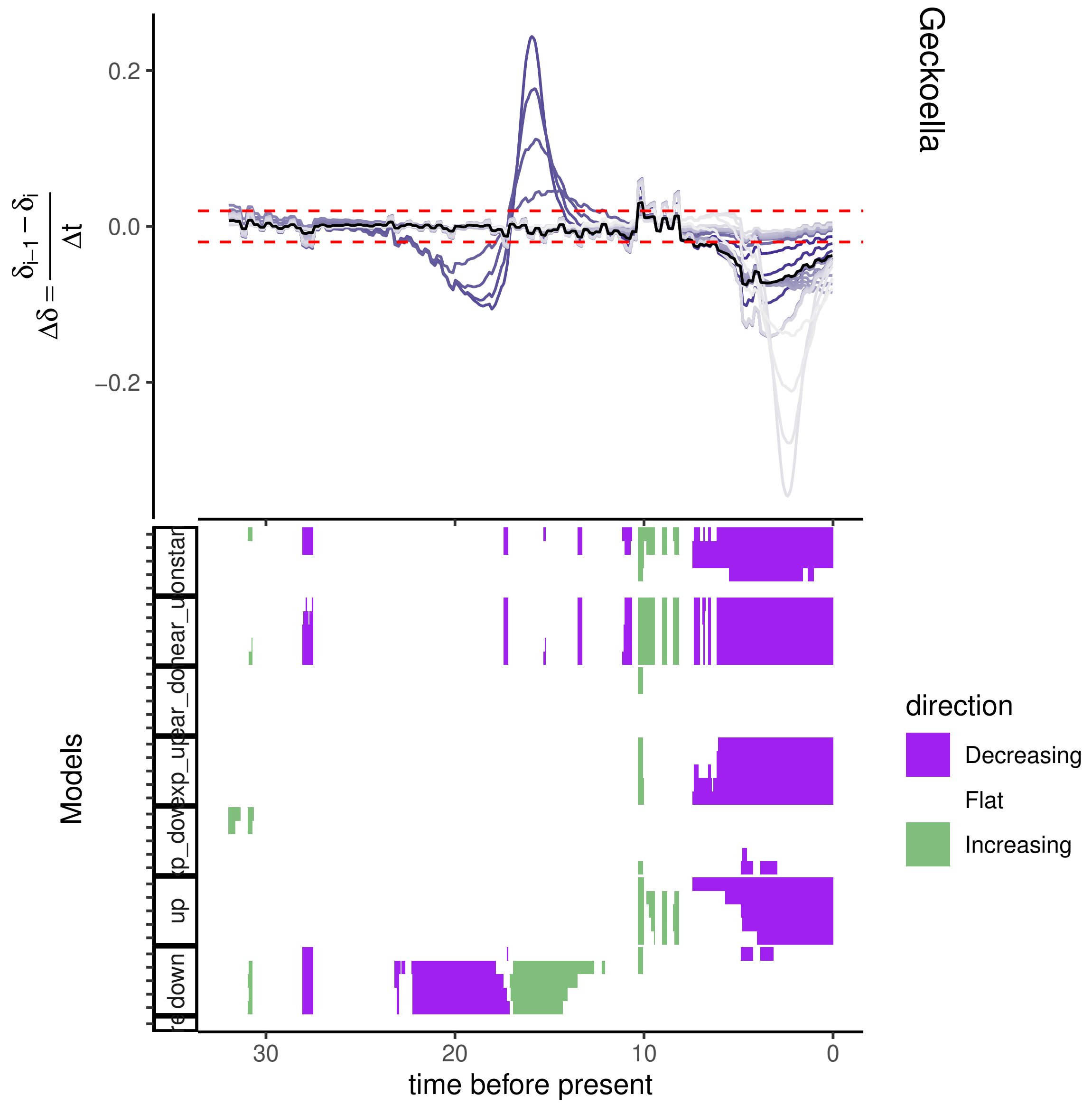

Geckoella
0.2
0.0
−0.2
$\Delta\delta = \frac{\delta_{i-1} - \delta_i}{\Delta t}$
Models
down
up
direction
Decreasing
Flat
Increasing
30
20
10
0
time before present

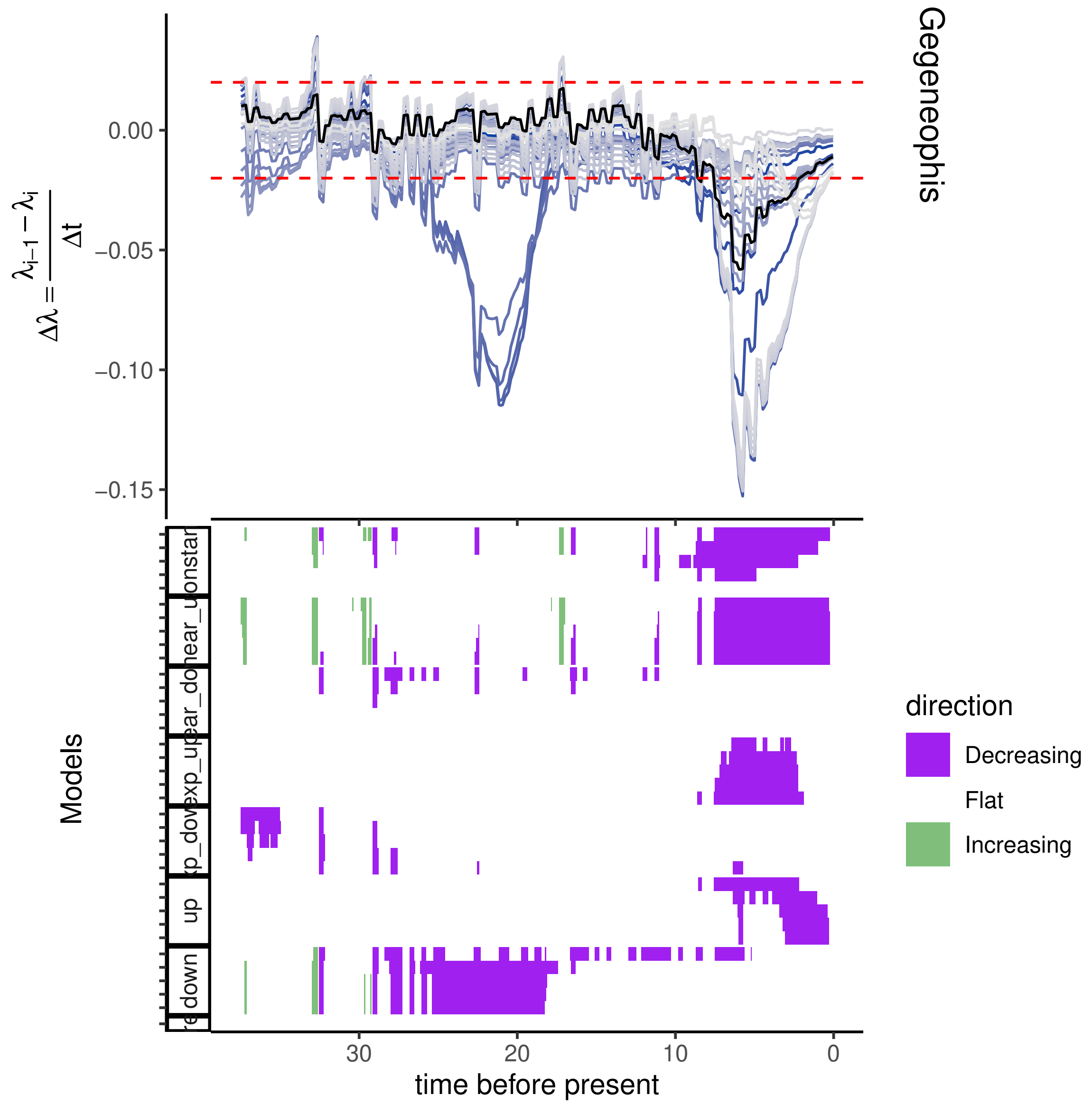

Gegeneophis
$\Delta\lambda = \frac{\lambda_{i-1} - \lambda_i}{\Delta t}$
0.00
−0.05
−0.10
−0.15
Models
constar
linear_u
linear_do
exp_up
exp_dow
up
down
30
20
10
0
time before present
direction
Decreasing
Flat
Increasing

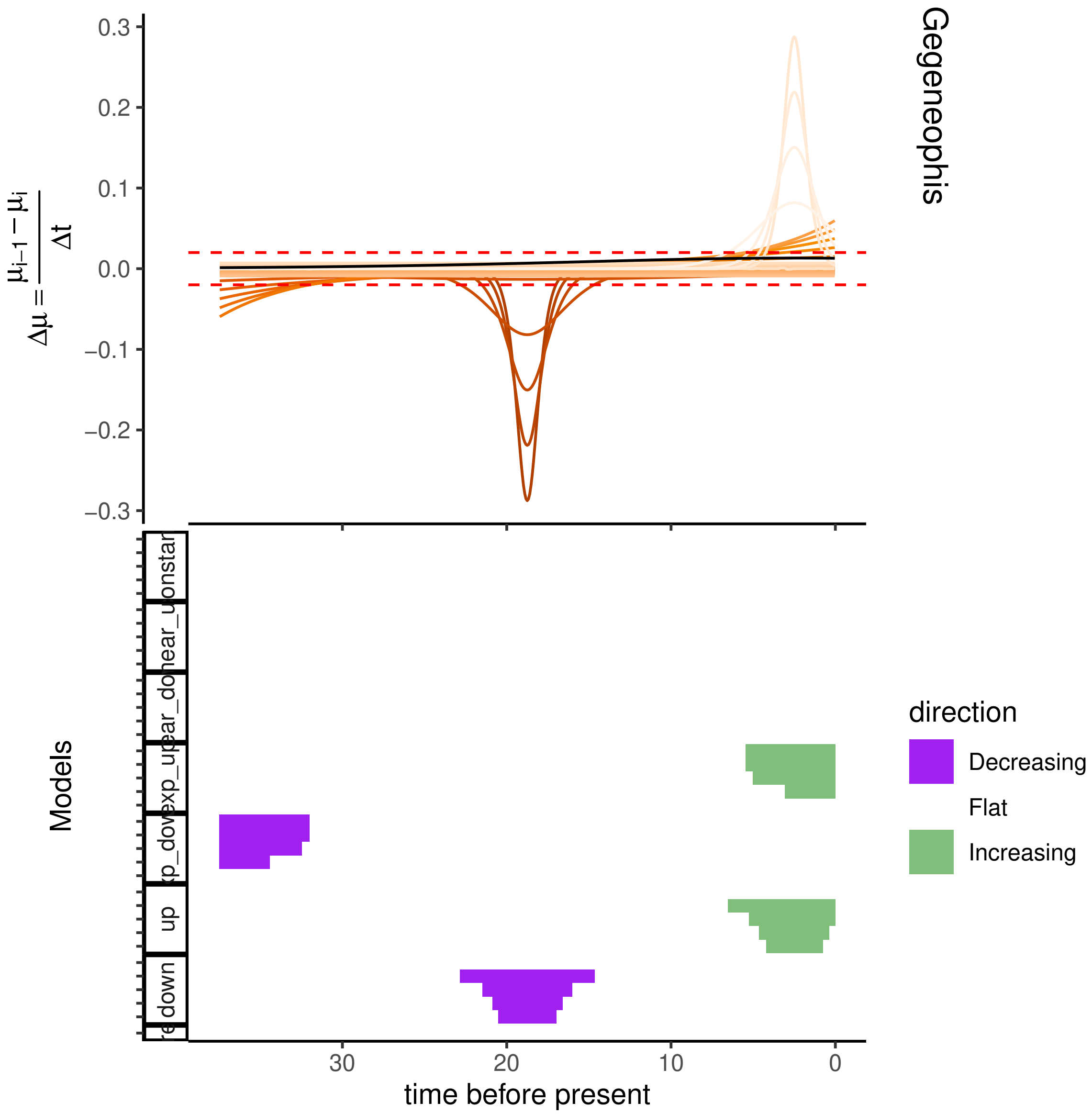

Gegeneophis
$\Delta\mu = \frac{\mu_{i-1} - \mu_i}{\Delta t}$
0.3
0.2
0.1
0.0
−0.1
−0.2
−0.3
Models
up
down
time before present
30
20
10
0
direction
Decreasing
Flat
Increasing

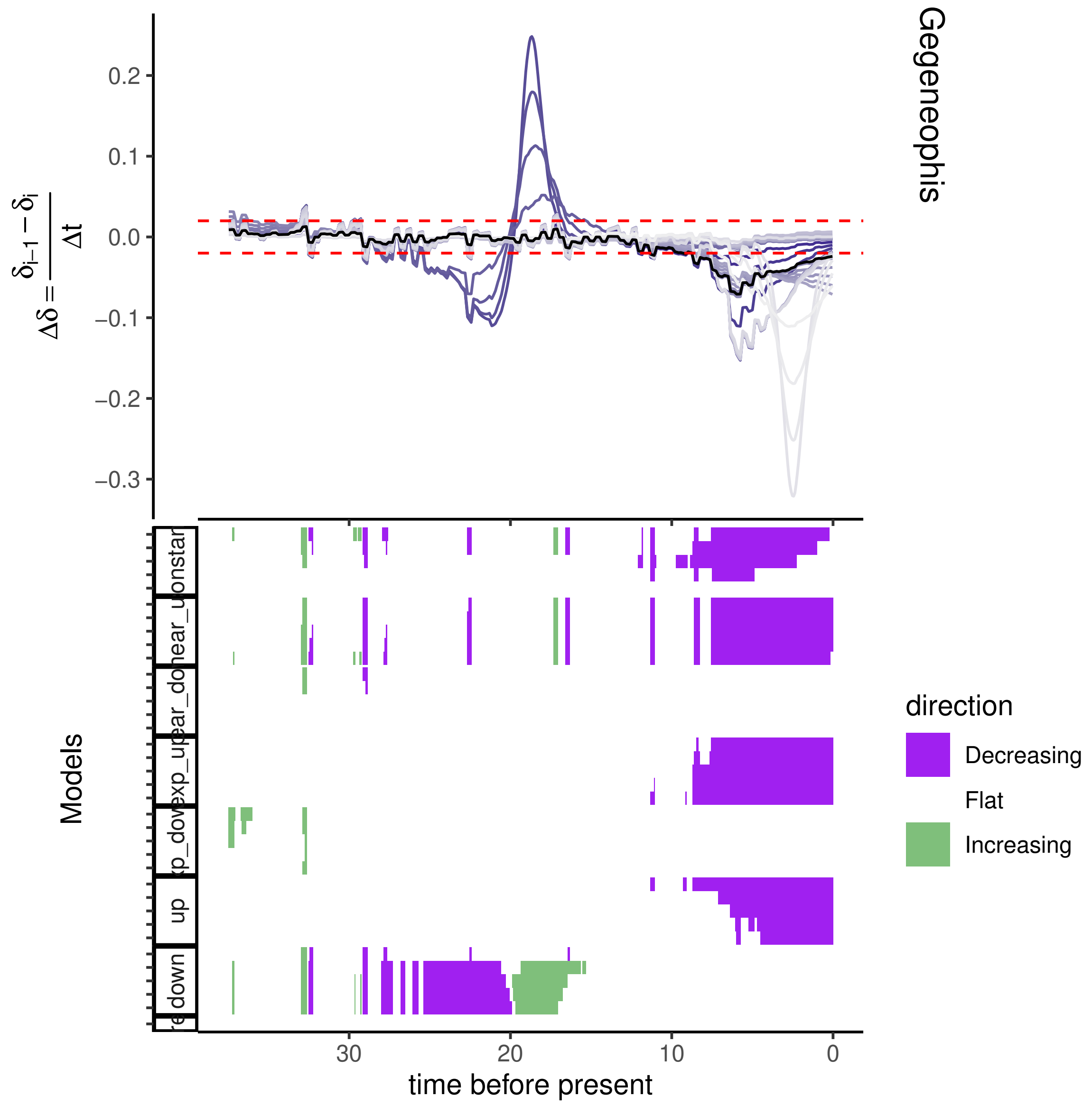

Gegeneophis
$\Delta\delta = \frac{\delta_{i-1} - \delta_i}{\Delta t}$
0.2
0.1
0.0
−0.1
−0.2
−0.3
Models
up
down
30
20
10
0
time before present
direction
Decreasing
Flat
Increasing

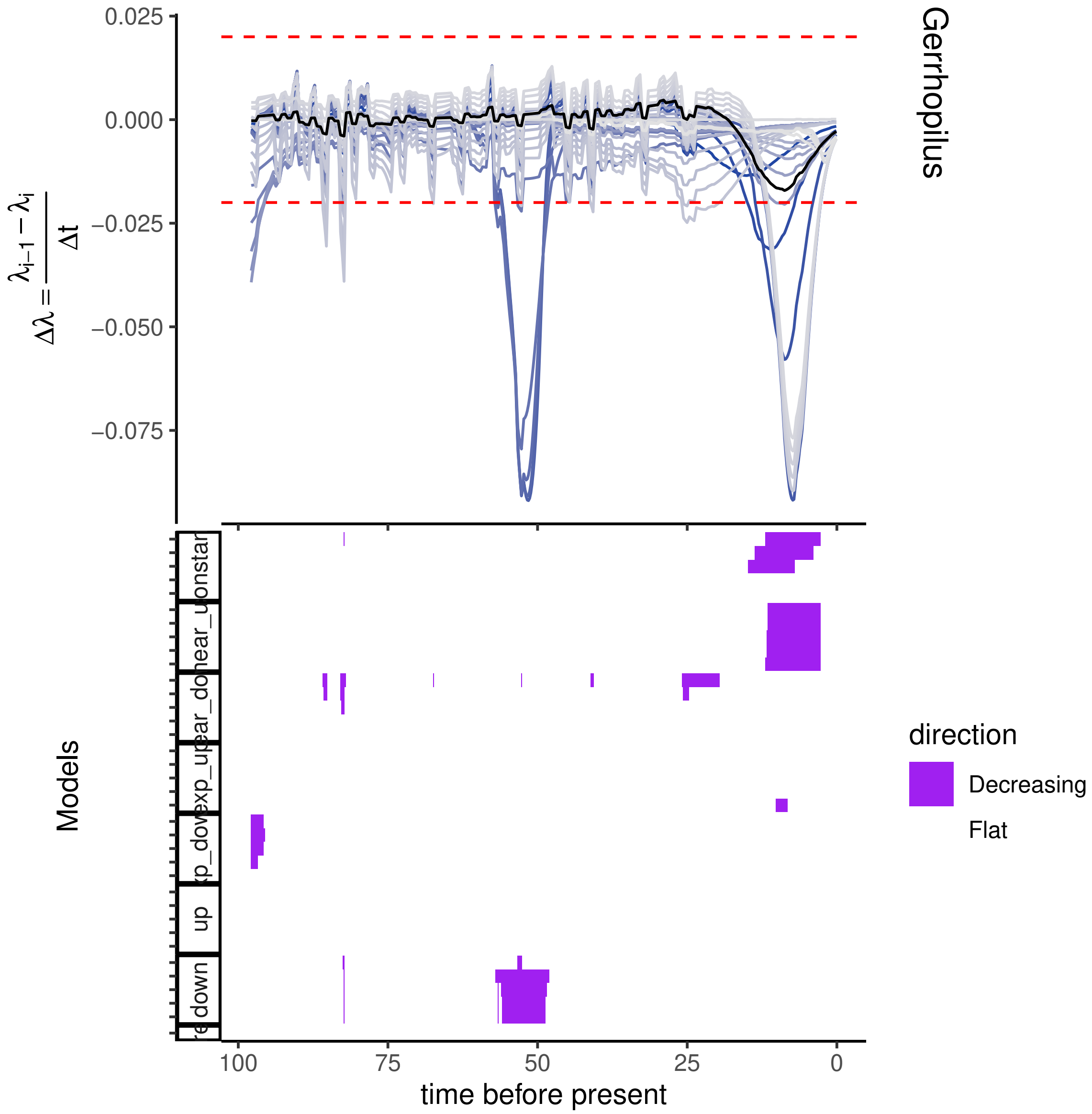
Gerrhopilus
0.025
0.000
−0.025
−0.050
−0.075
$\Delta\lambda = \frac{\lambda_{i-1} - \lambda_i}{\Delta t}$
Models
up
down
time before present
100
75
50
25
0
direction
Decreasing
Flat

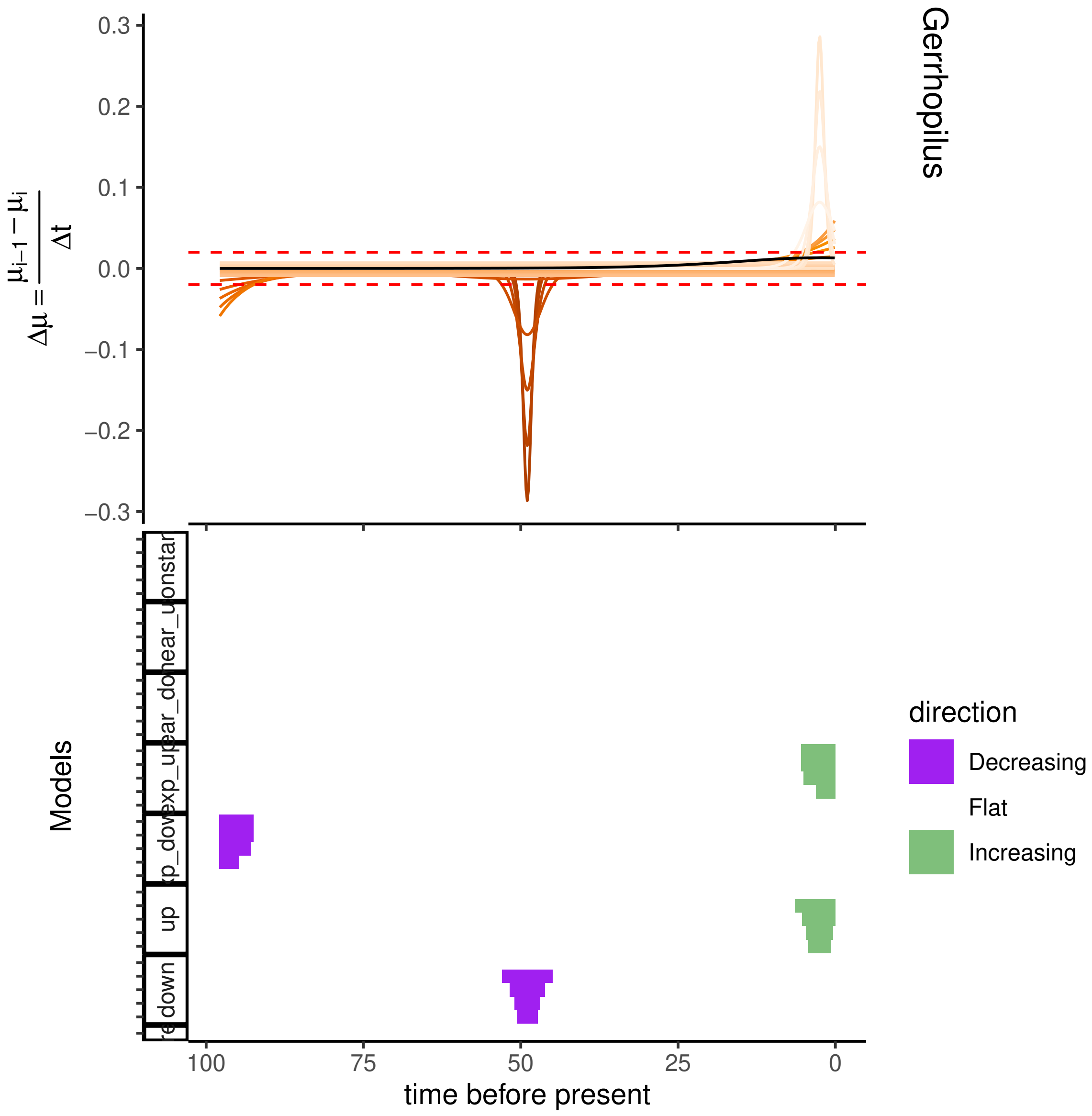

Gerrhopilus
$\Delta\mu = \frac{\mu_{i-1} - \mu_i}{\Delta t}$
0.3
0.2
0.1
0.0
−0.1
−0.2
−0.3
Models
down
up
time before present
100
75
50
25
0
direction
Decreasing
Flat
Increasing

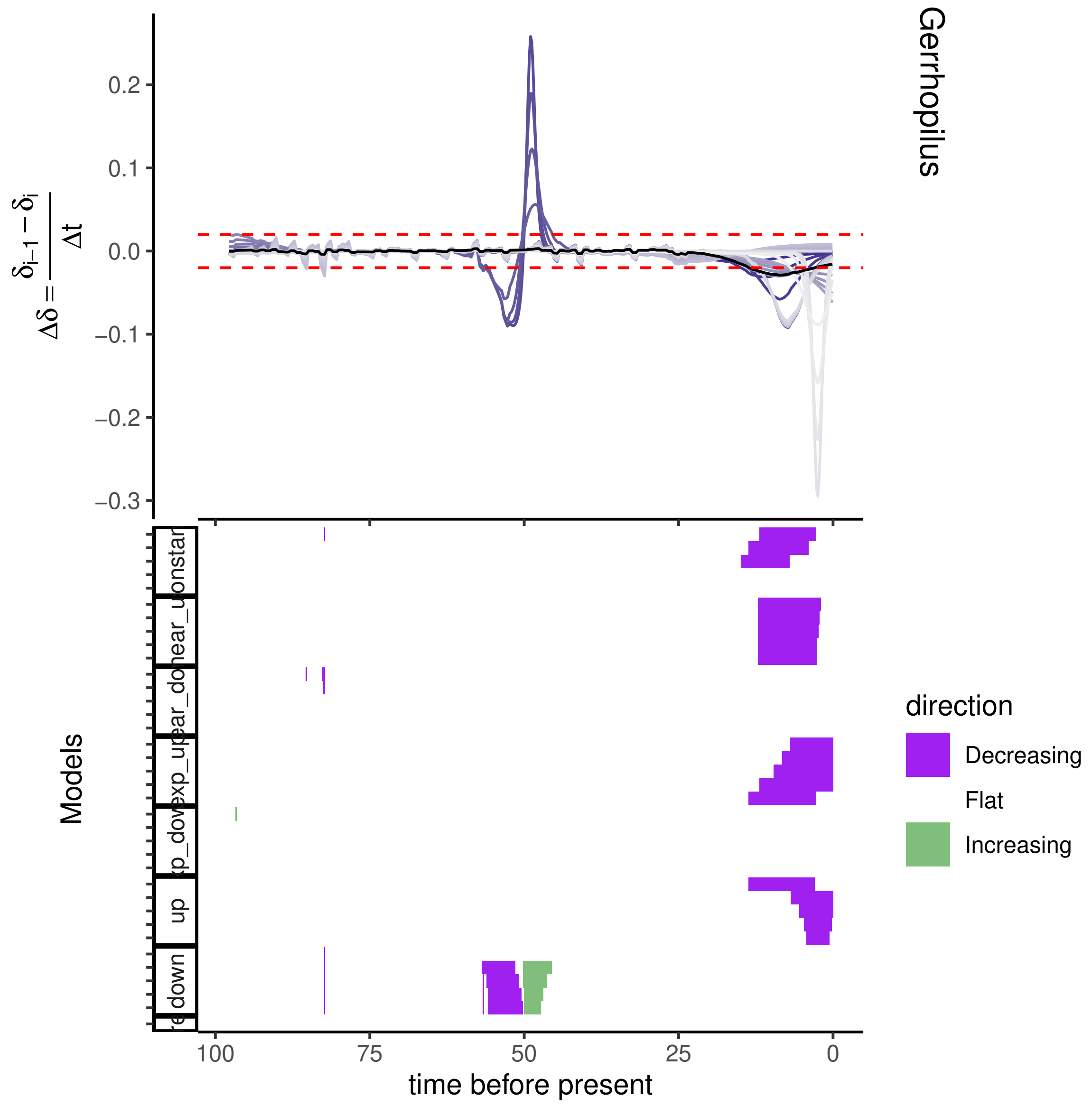
Gerrhopilus
0.2
0.1
0.0
−0.1
−0.2
−0.3
Models
direction
Decreasing
Flat
Increasing
100
75
50
25
0
time before present

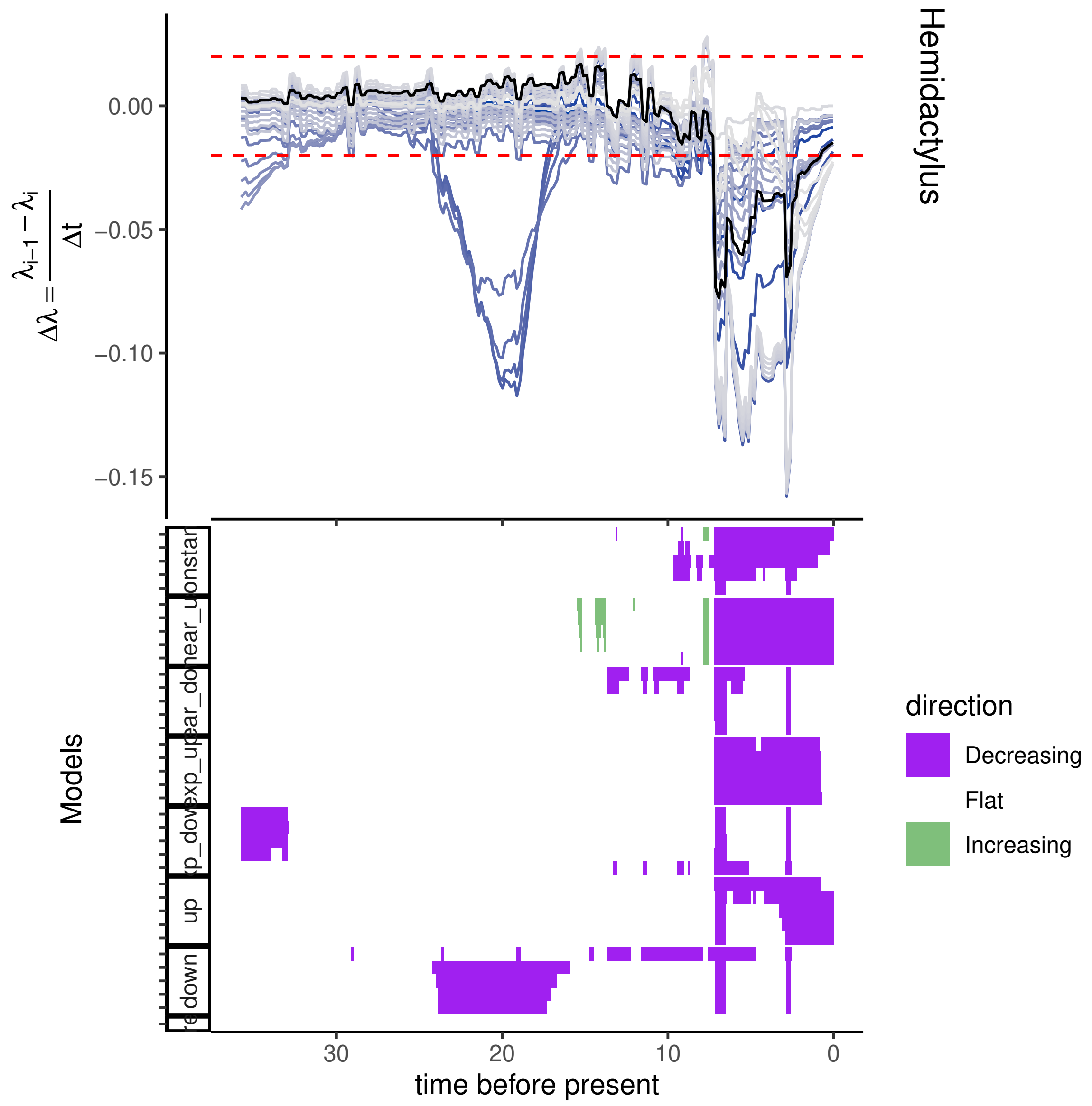

Hemidactylus
$\Delta\lambda = \frac{\lambda_{i-1} - \lambda_i}{\Delta t}$
0.00
−0.05
−0.10
−0.15
Models
direction
Decreasing
Flat
Increasing
up
down
30
20
10
0
time before present

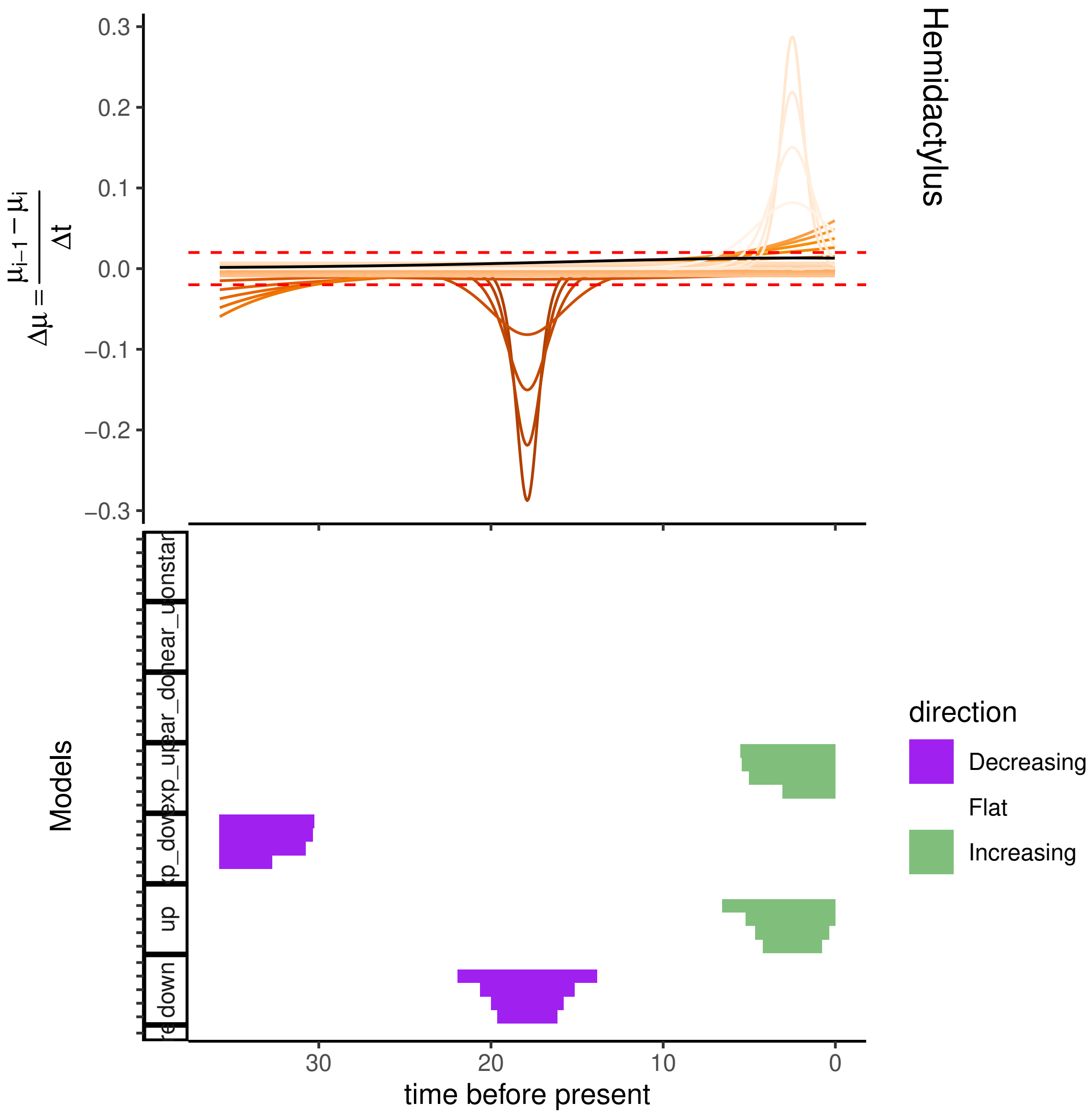
Hemidactylus
$\Delta\mu = \frac{\mu_{i-1} - \mu_i}{\Delta t}$
0.3
0.2
0.1
0.0
−0.1
−0.2
−0.3
Models
onstar
near_u
ear_do
xp_up
xp_dov
up
down
30
20
10
0
time before present
direction
Decreasing
Flat
Increasing

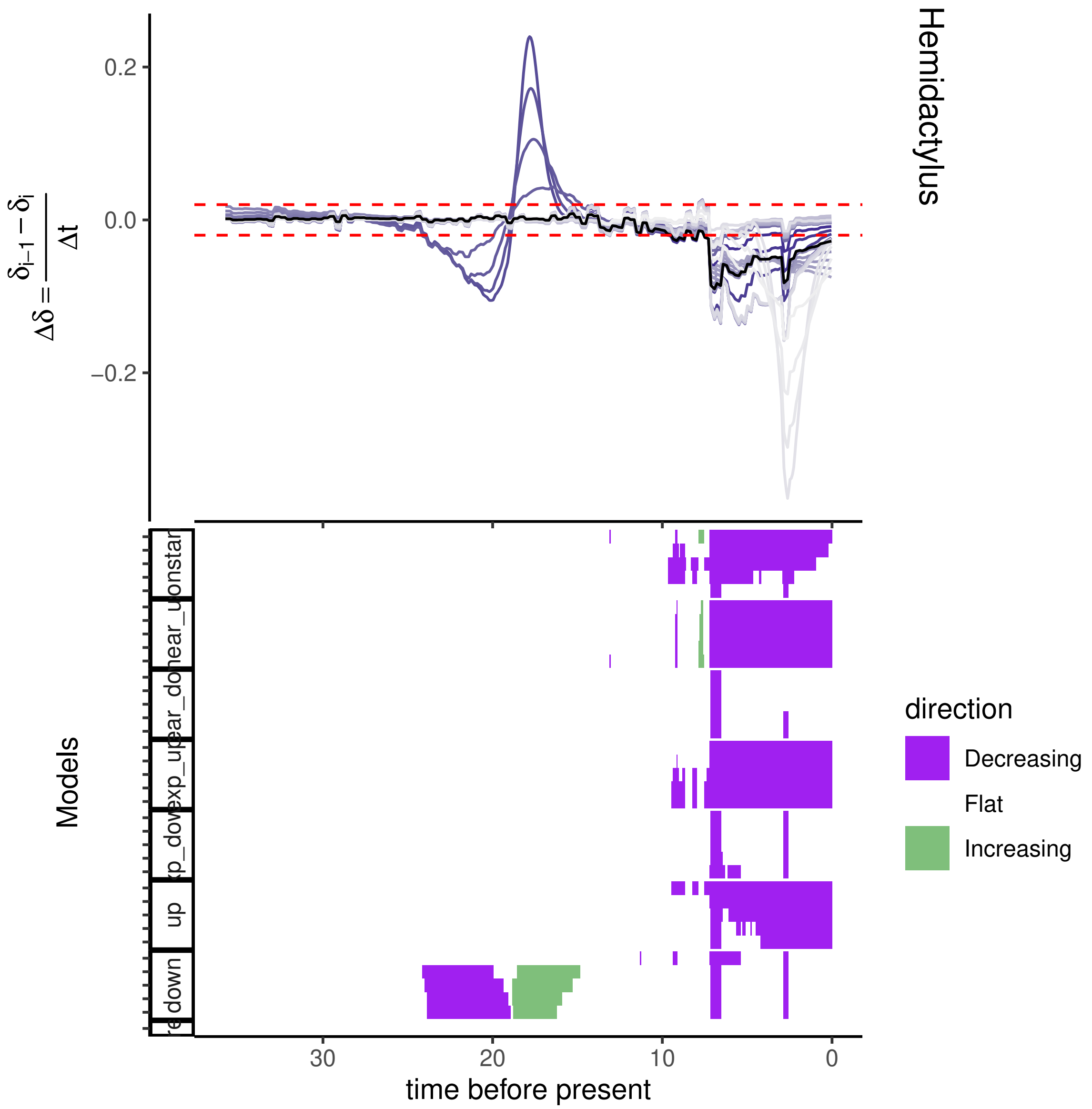
Hemidactylus
0.2
0.0
−0.2
Models
up
down
30
20
10
0
time before present
direction
Decreasing
Flat
Increasing

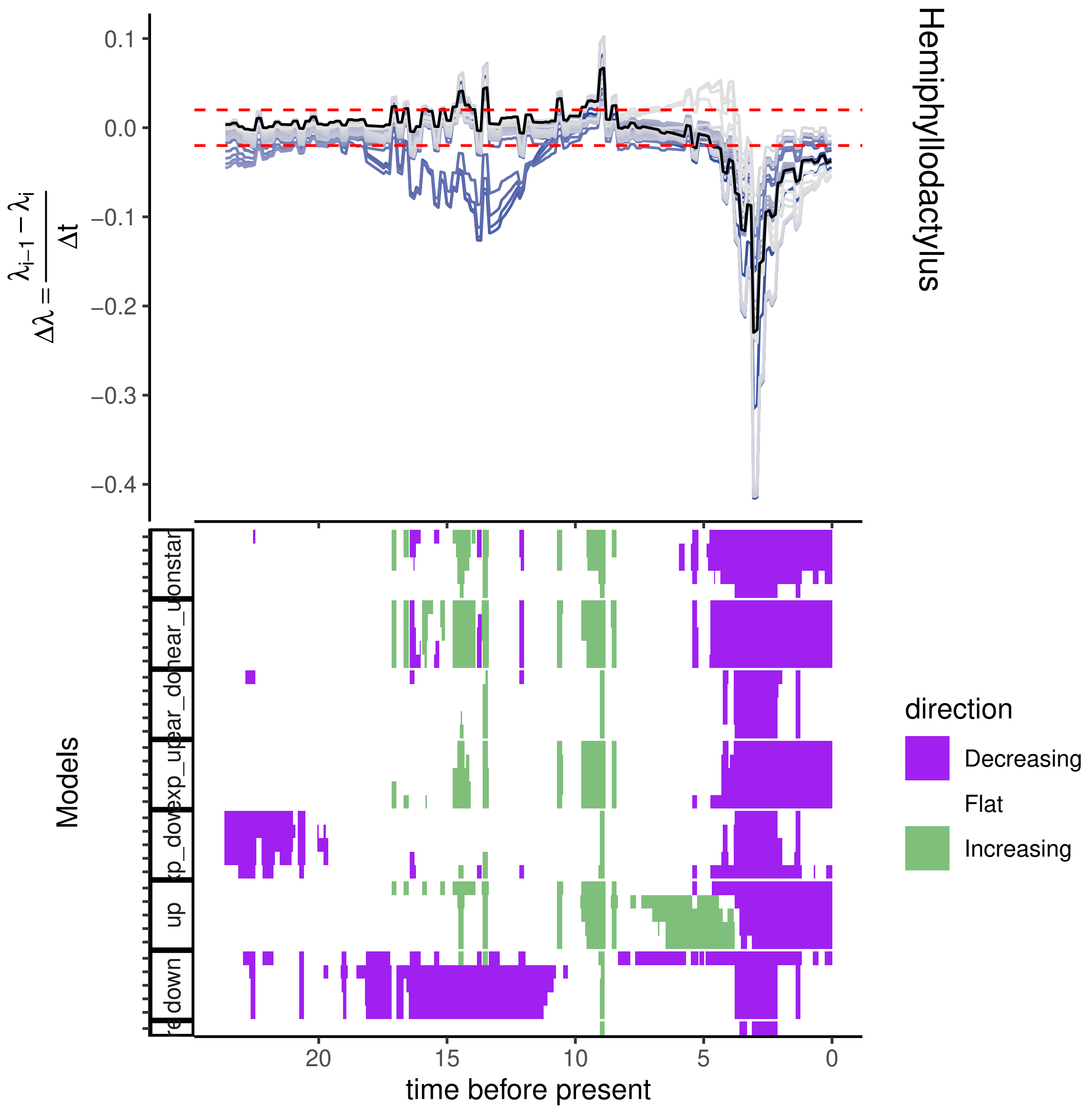
Hemiphyllodactylus
$\Delta\lambda = \frac{\lambda_{i-1} - \lambda_i}{\Delta t}$
0.1
0.0
−0.1
−0.2
−0.3
−0.4
Models
up
down
direction
Decreasing
Flat
Increasing
20
15
10
5
0
time before present

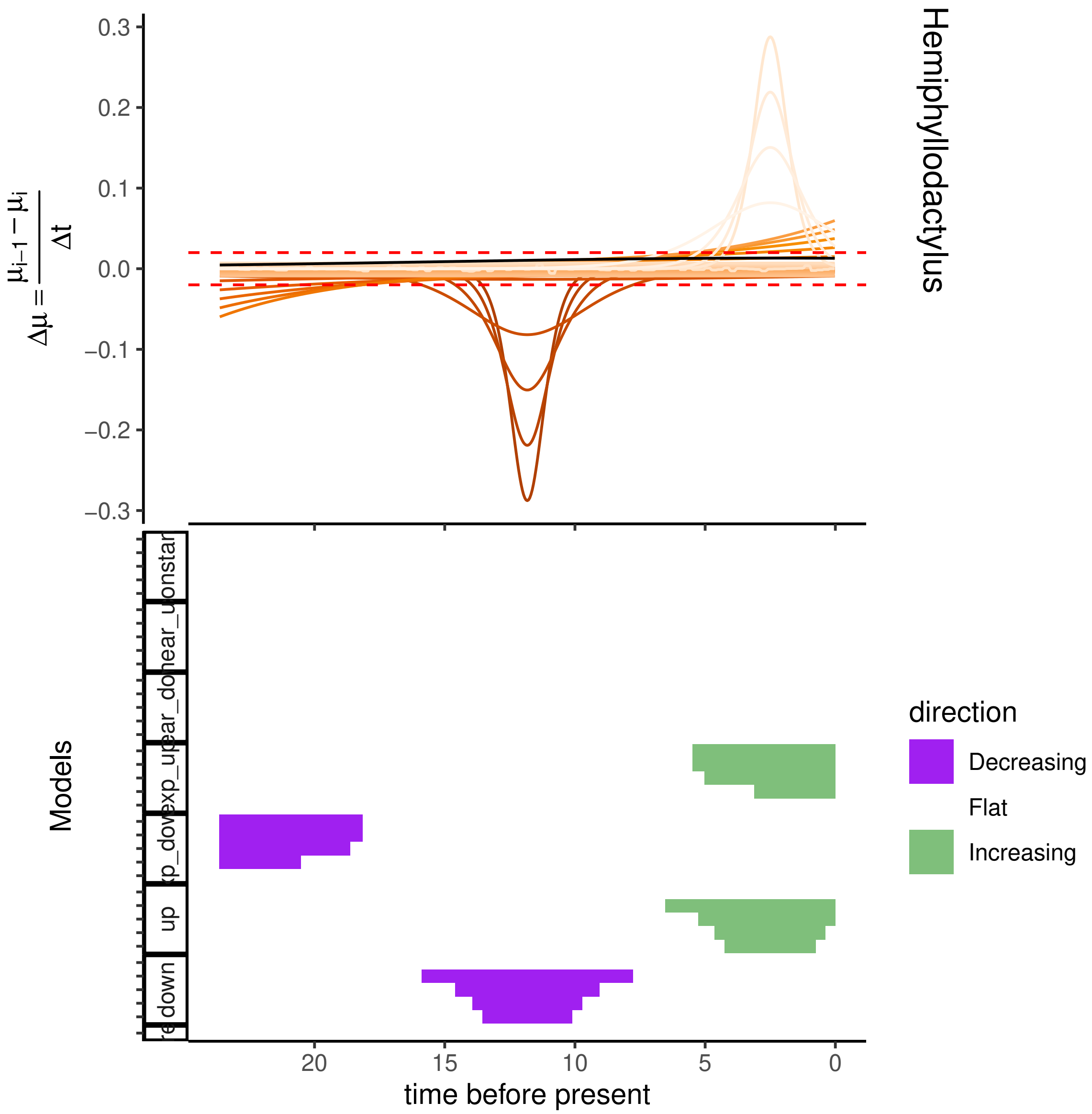
0.3
0.2
0.1
0.0
−0.1
−0.2
−0.3
$\Delta\mu = \frac{\mu_{i-1} - \mu_i}{\Delta t}$
Hemiphyllodactylus
Models
onstar
near_u
near_do
exp_up
xp_dov
up
down
20
15
10
5
0
time before present
direction
Decreasing
Flat
Increasing

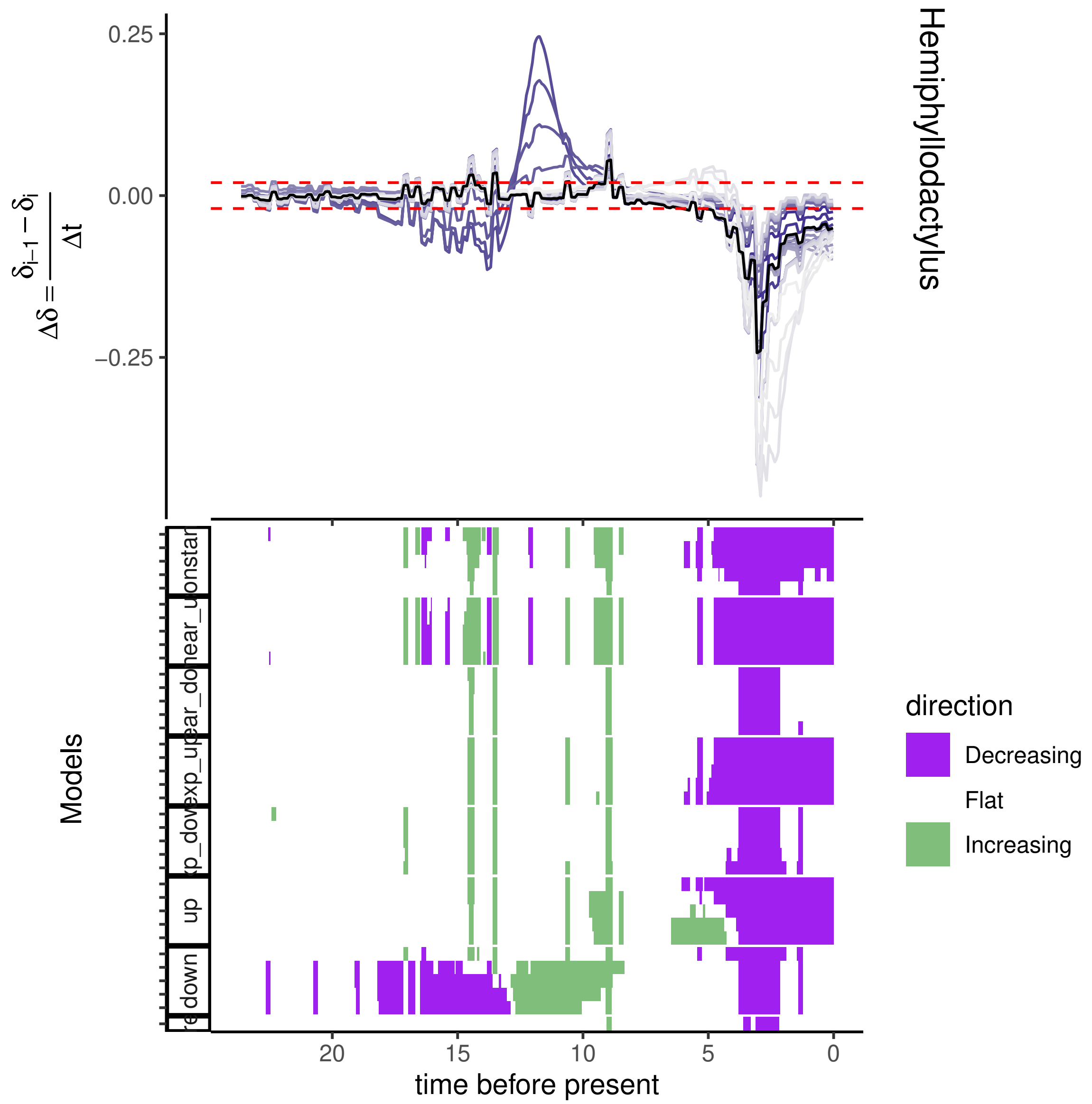

0.25
0.00
−0.25
$\Delta\delta = \frac{\delta_{i-1} - \delta_i}{\Delta t}$
Hemiphyllodactylus
Models
up
down
20
15
10
5
0
time before present
direction
Decreasing
Flat
Increasing

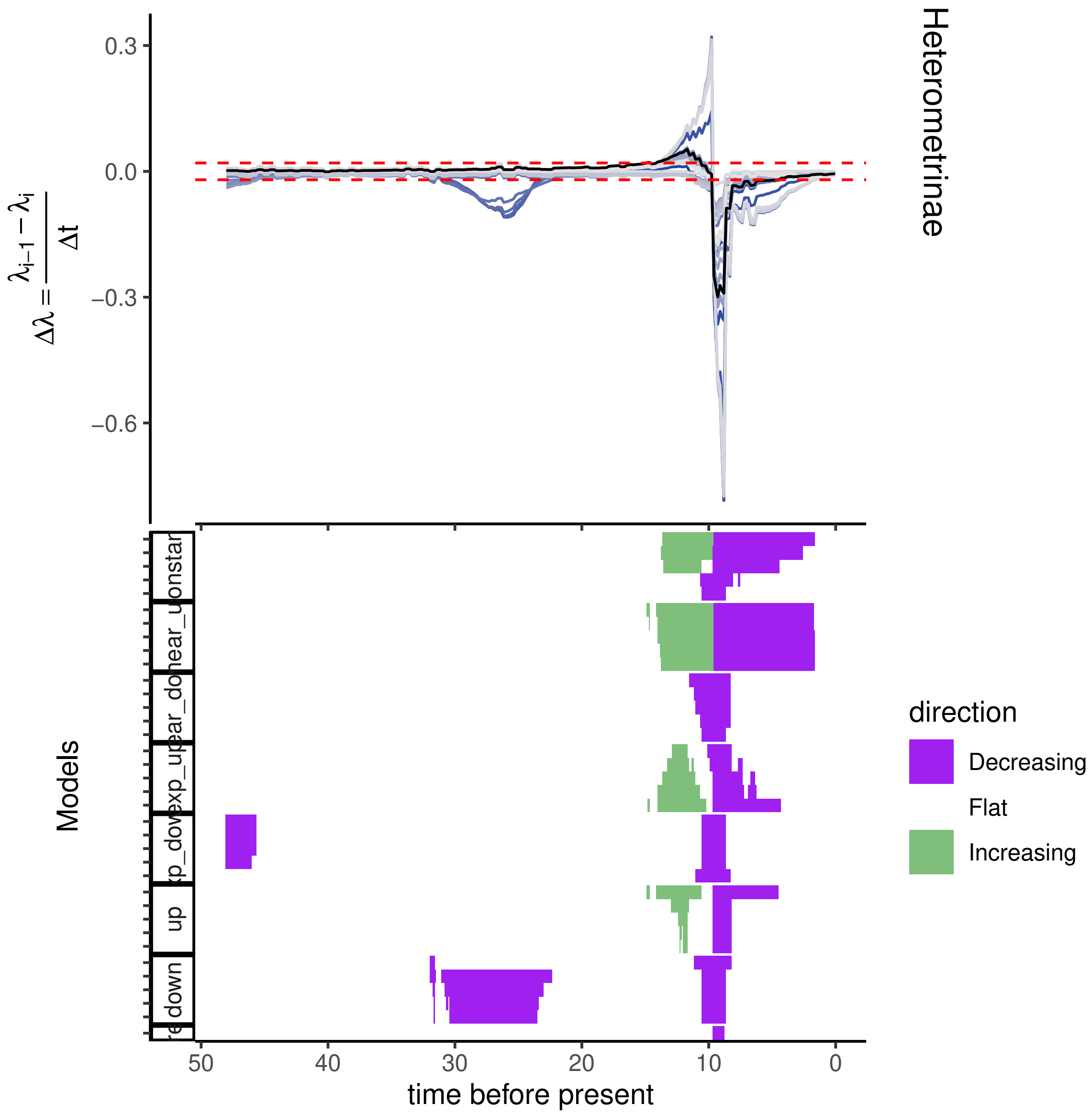
Heterometrinae
0.3
0.0
−0.3
−0.6
Models
direction
Decreasing
Flat
Increasing
up
down
50
40
30
20
10
0
time before present

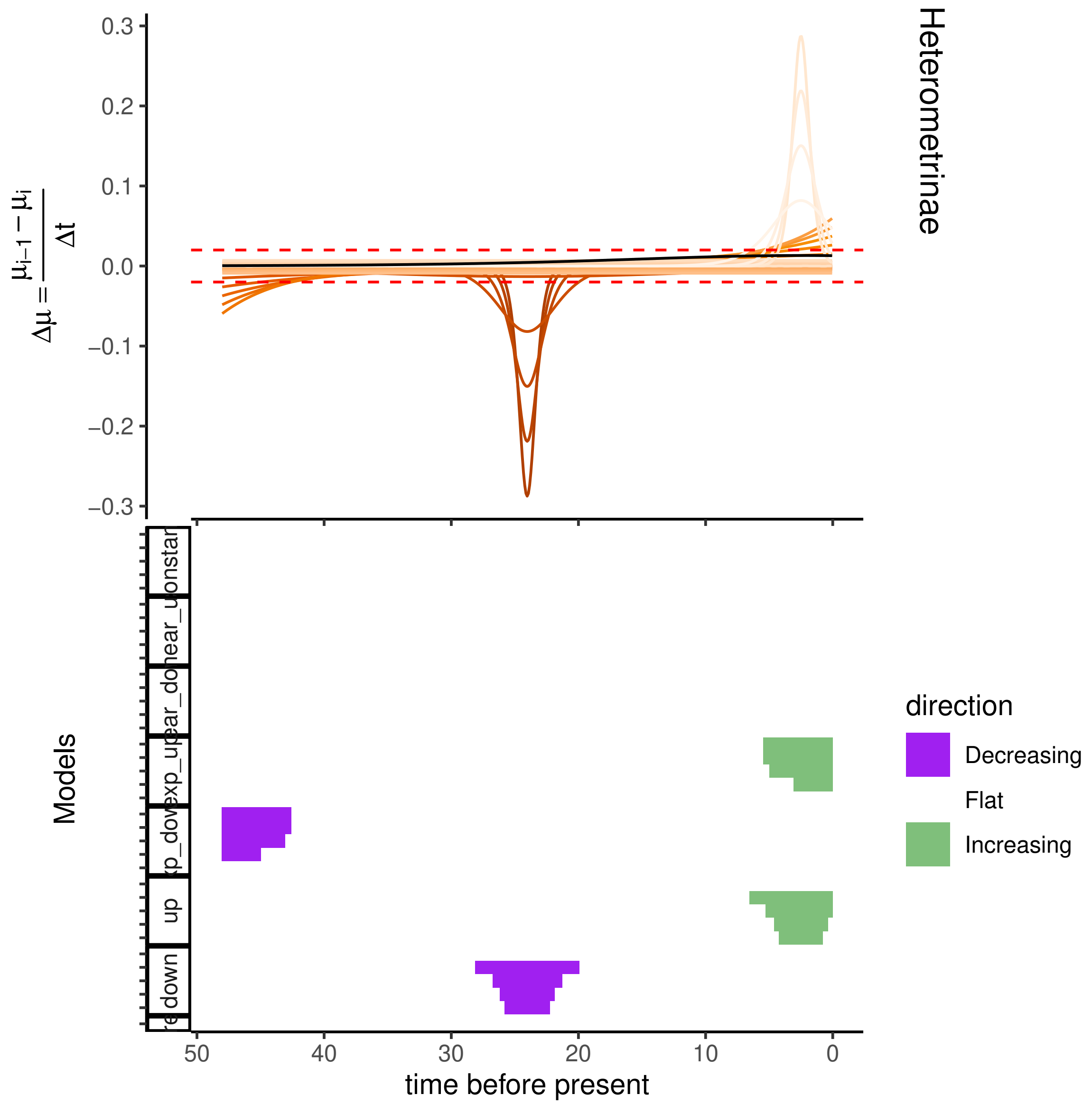

Heterometrinae
$\Delta\mu = \frac{\mu_{i-1} - \mu_i}{\Delta t}$
0.3
0.2
0.1
0.0
−0.1
−0.2
−0.3
Models
up
down
direction
Decreasing
Flat
Increasing
50
40
30
20
10
0
time before present

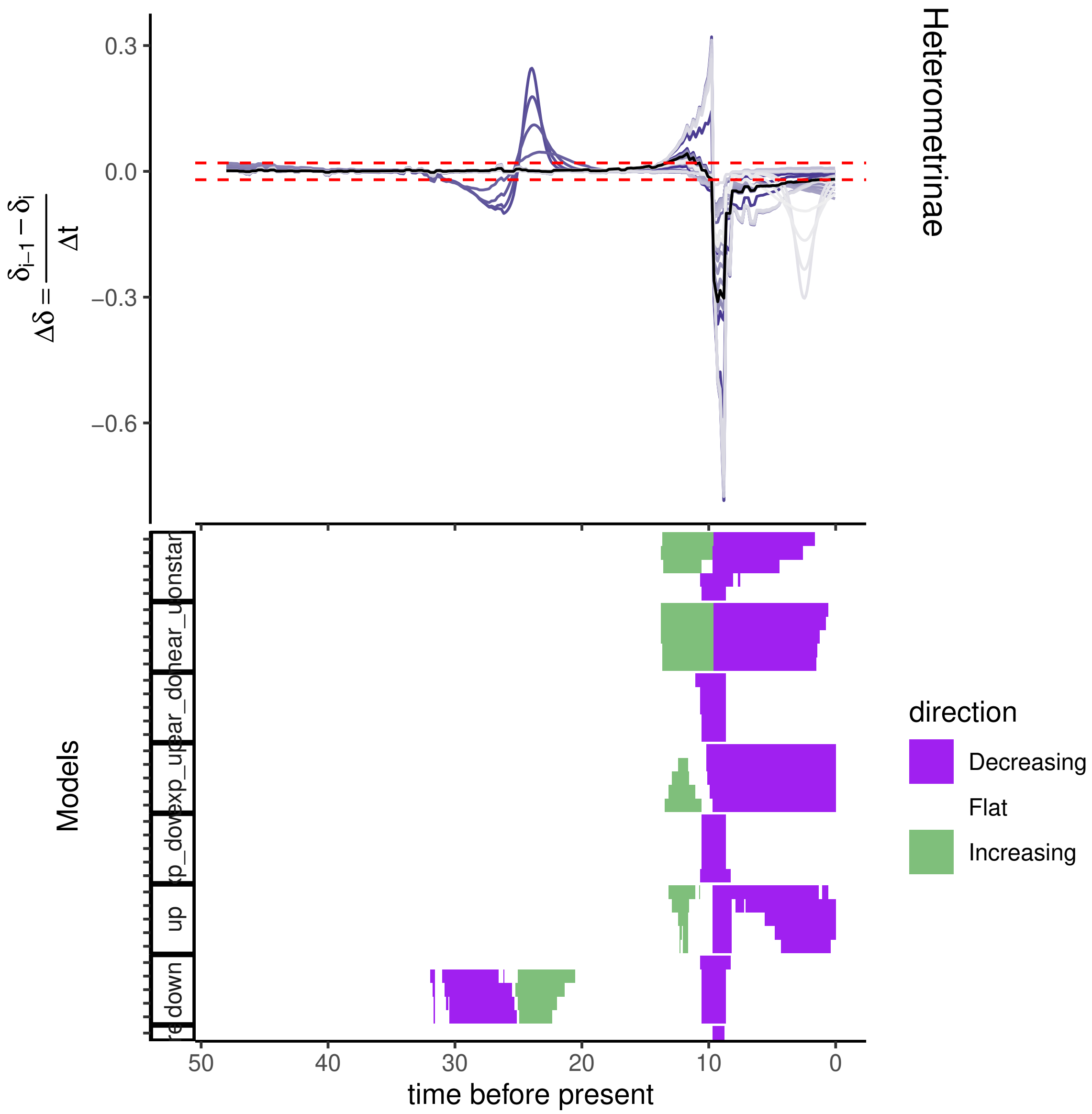

Heterometrinae
$\Delta\delta = \frac{\delta_{i-1} - \delta_i}{\Delta t}$
0.3
0.0
−0.3
−0.6
Models
up
down
time before present
50
40
30
20
10
0
direction
Decreasing
Flat
Increasing

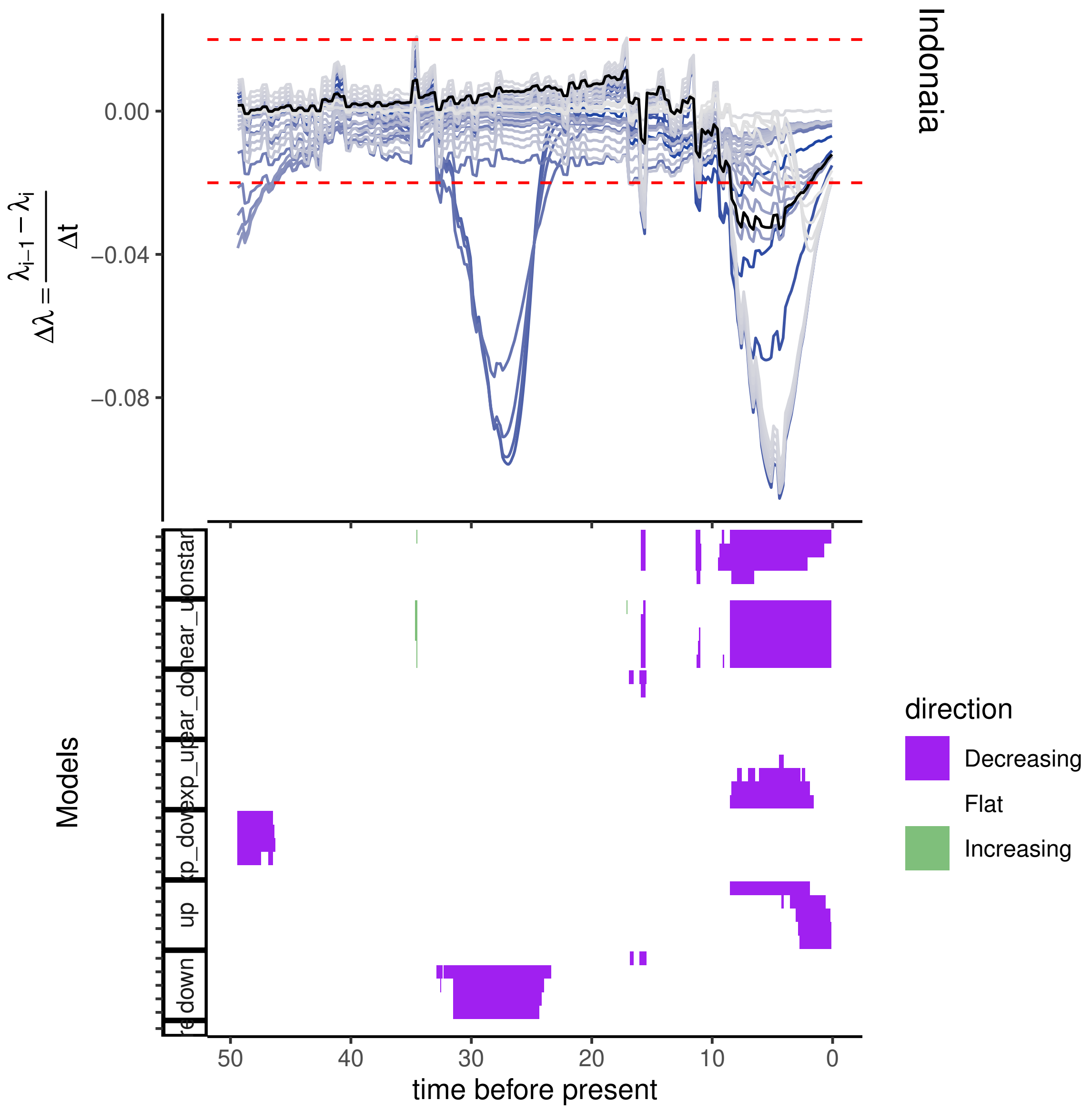

Indonaia
0.00
−0.04
−0.08
Models
up
down
50
40
30
20
10
0
time before present
direction
Decreasing
Flat
Increasing

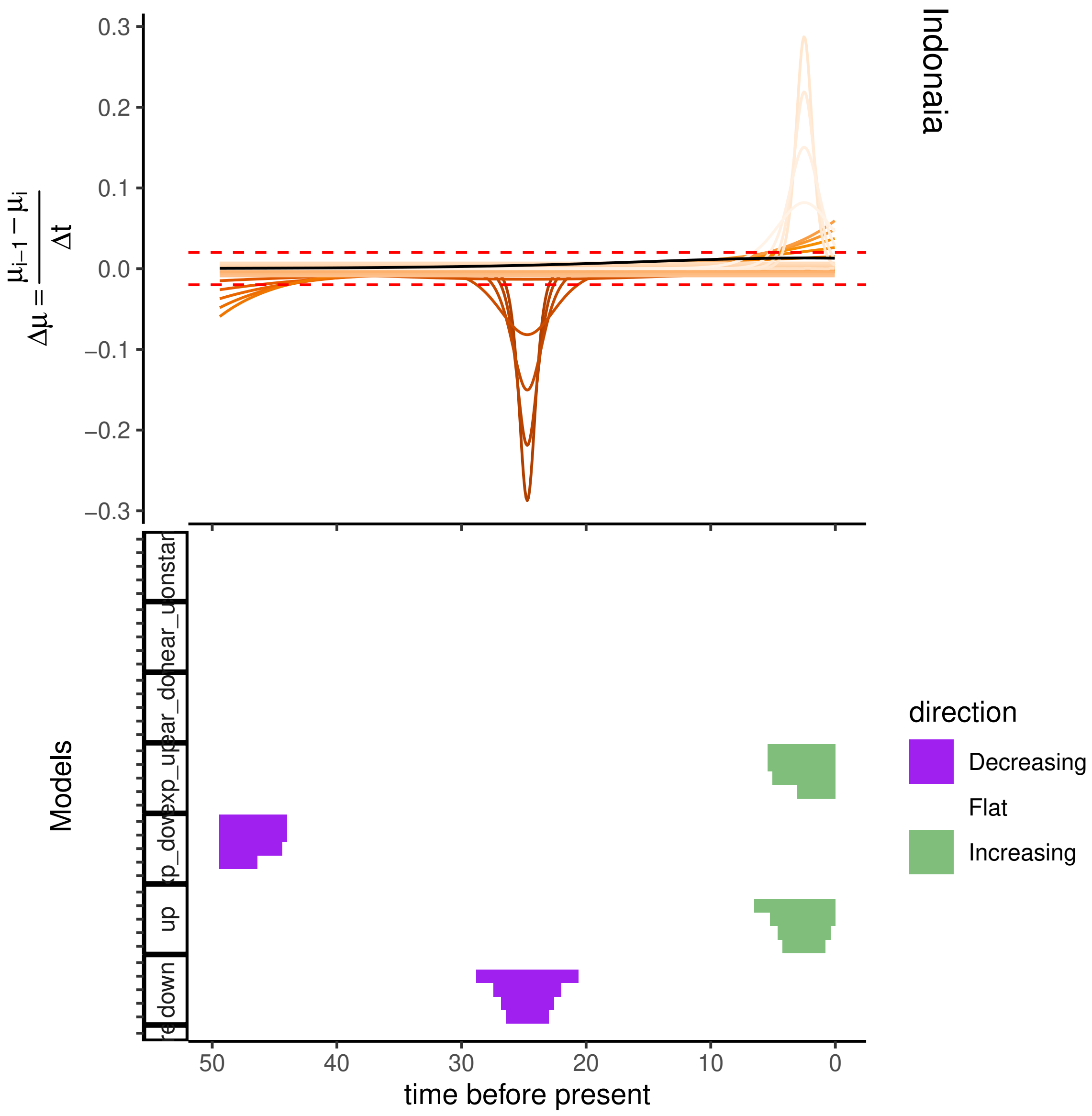
Indonaia
$\Delta\mu = \frac{\mu_{i-1} - \mu_i}{\Delta t}$
0.3
0.2
0.1
0.0
−0.1
−0.2
−0.3
Models
up
down
direction
Decreasing
Flat
Increasing
50
40
30
20
10
0
time before present

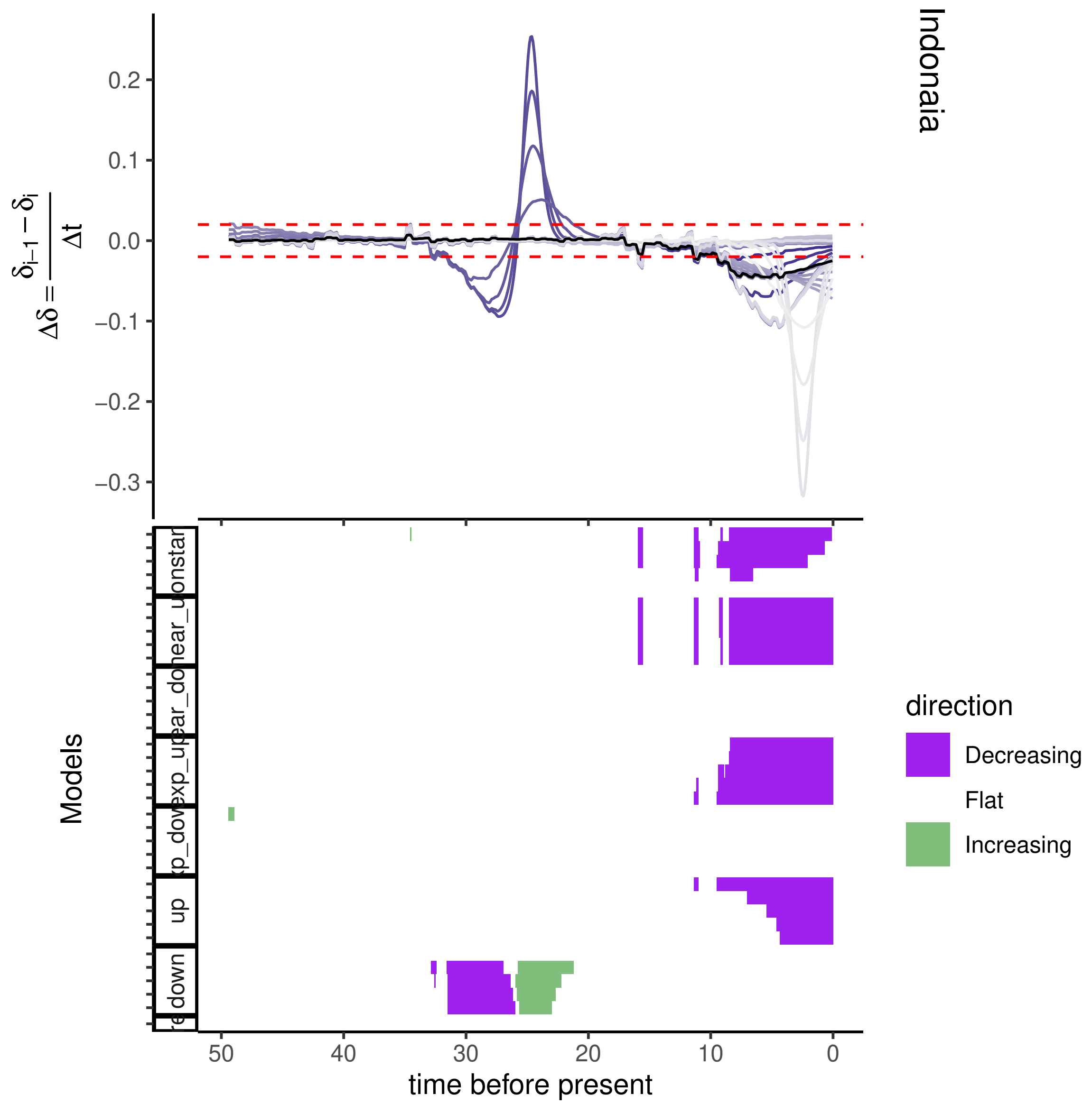
Indonaia
0.2
0.1
0.0
−0.1
−0.2
−0.3
$\Delta\delta = \frac{\delta_{i-1} - \delta_i}{\Delta t}$
Models
up
down
direction
Decreasing
Flat
Increasing
50
40
30
20
10
0
time before present

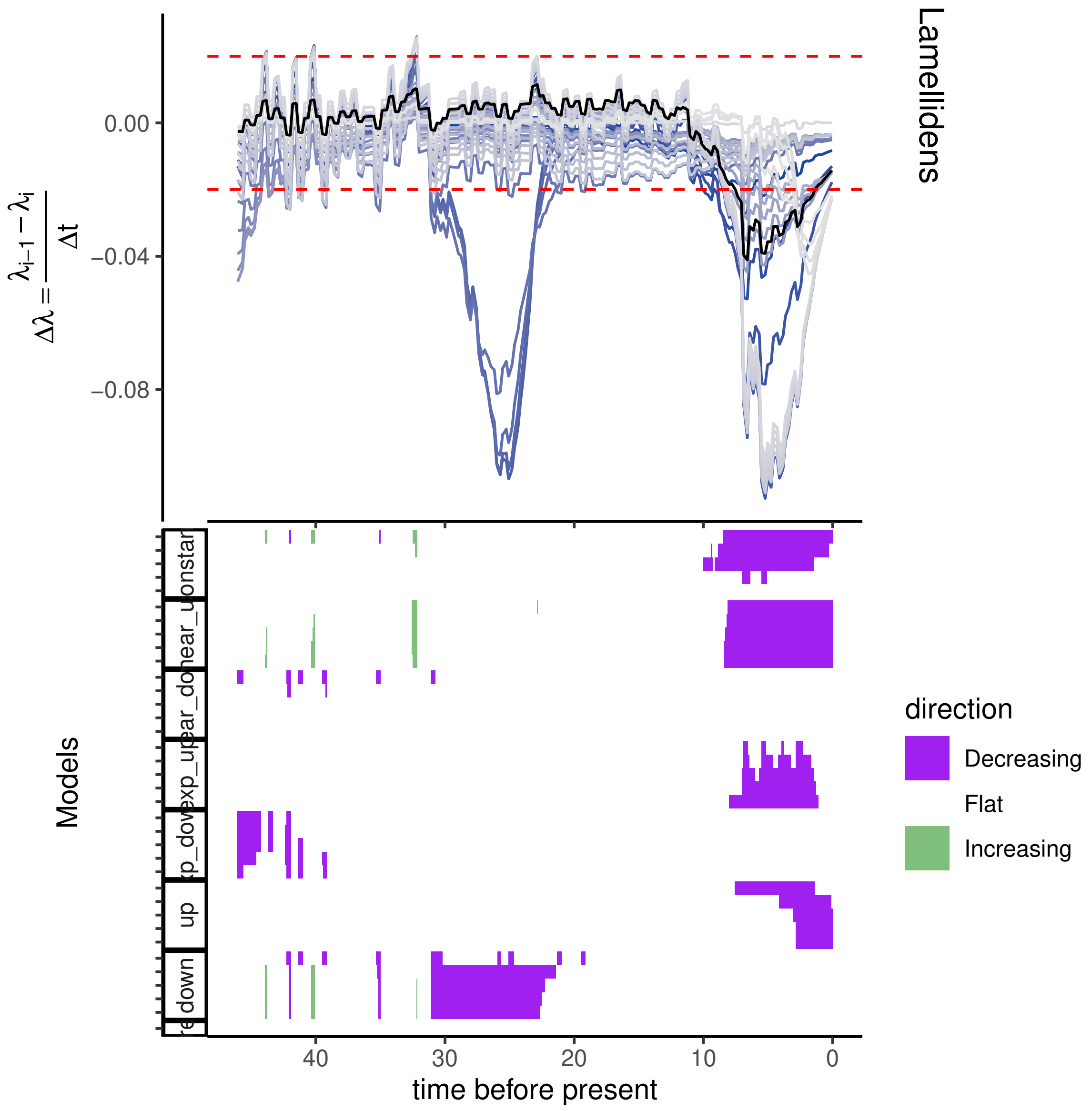

Lamellidens
0.00
−0.04
−0.08
$\Delta\lambda = \frac{\lambda_{i-1} - \lambda_i}{\Delta t}$
Models
direction
Decreasing
Flat
Increasing
40
30
20
10
0
time before present

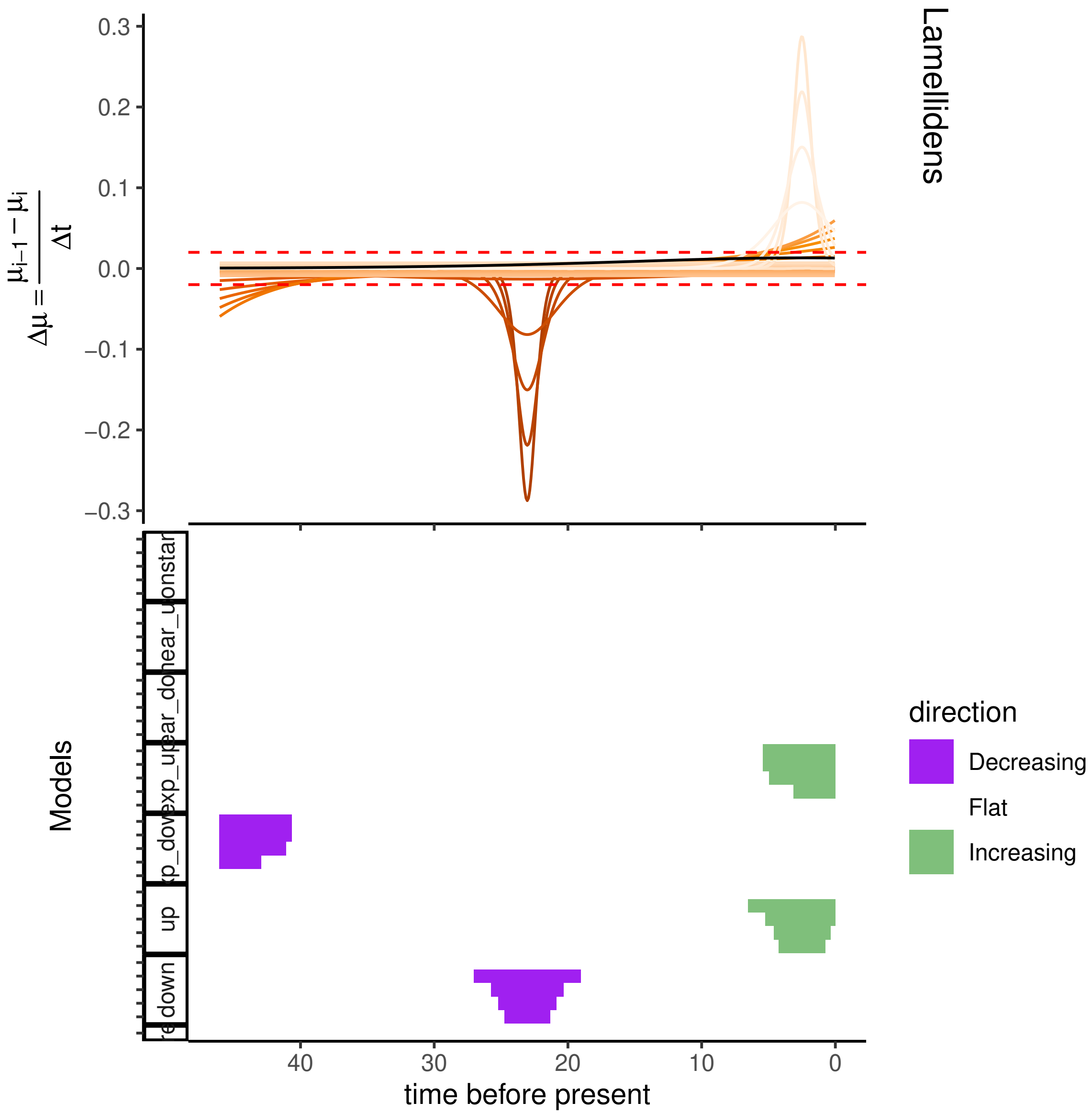

Lamellidens
$\Delta\mu = \frac{\mu_{i-1} - \mu_i}{\Delta t}$
0.3
0.2
0.1
0.0
−0.1
−0.2
−0.3
Models
up
down
direction
Decreasing
Flat
Increasing
40
30
20
10
0
time before present

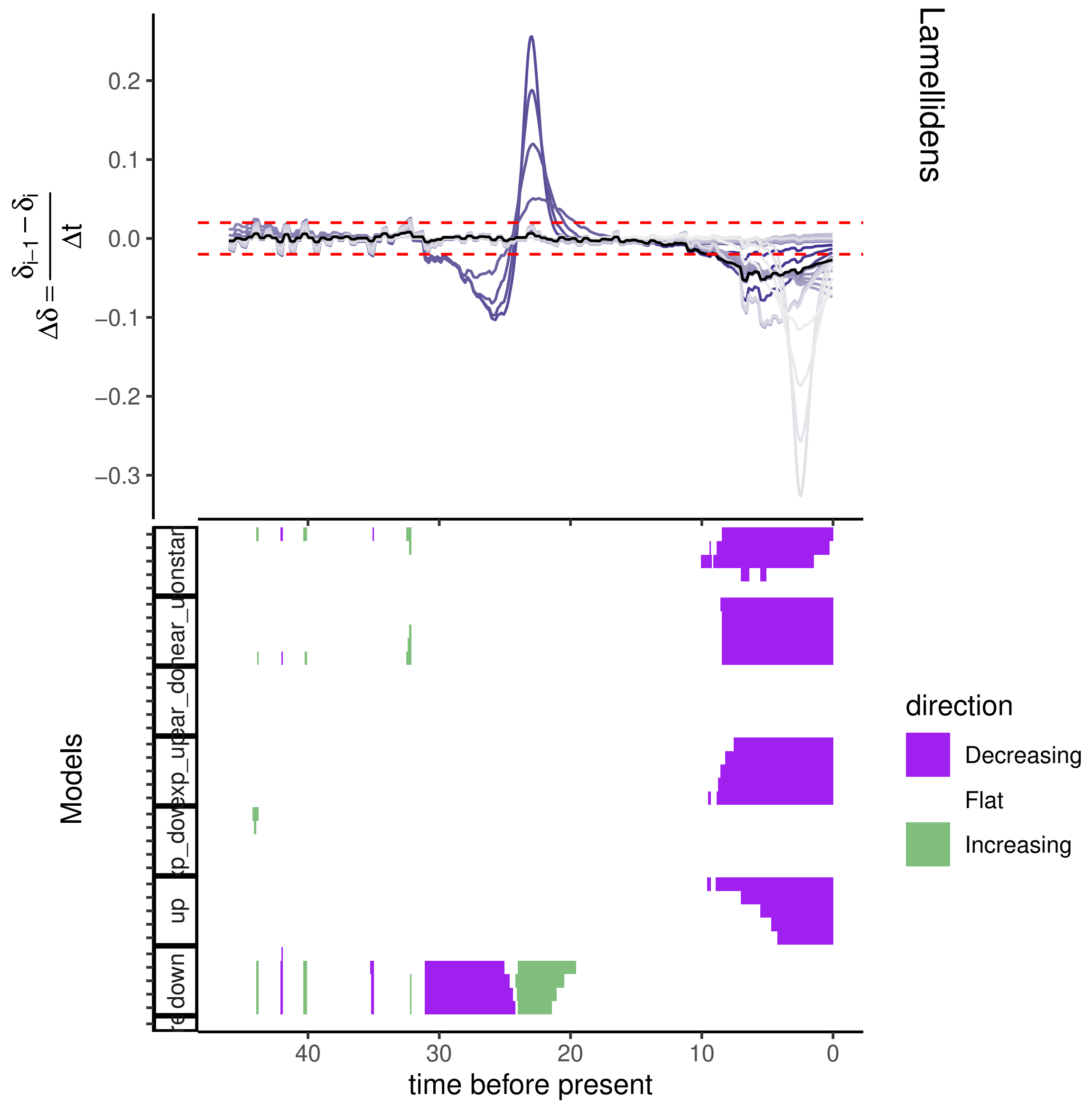

Lamellidens
$\Delta\delta = \frac{\delta_{i-1} - \delta_i}{\Delta t}$
0.2
0.1
0.0
−0.1
−0.2
−0.3
Models
down
up
direction
Decreasing
Flat
Increasing
40
30
20
10
0
time before present

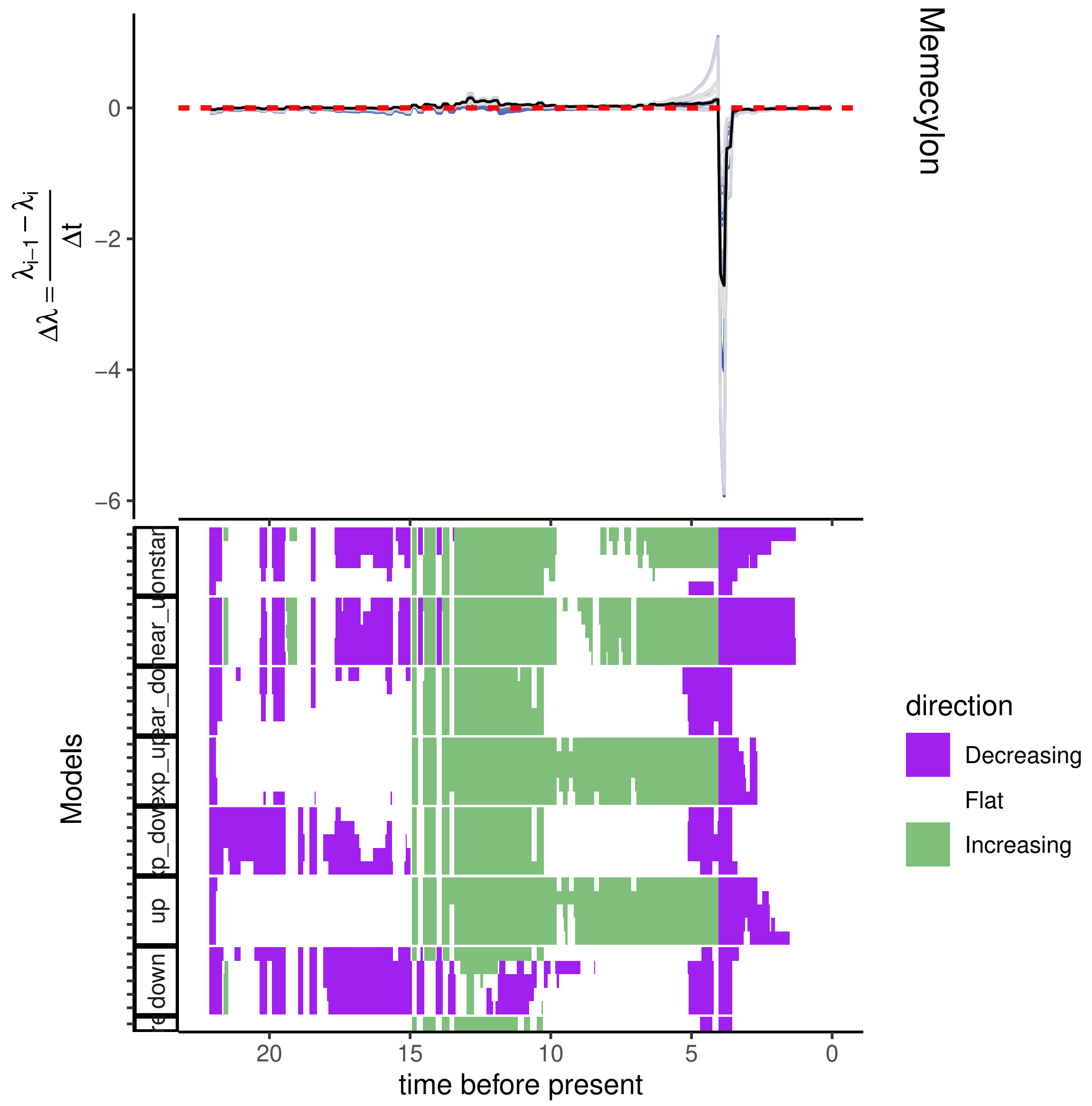

Memecylon
$\Delta\lambda = \frac{\lambda_{i-1} - \lambda_i}{\Delta t}$
0
−2
−4
−6
Models
direction
Decreasing
Flat
Increasing
up
down
20
15
10
5
0
time before present

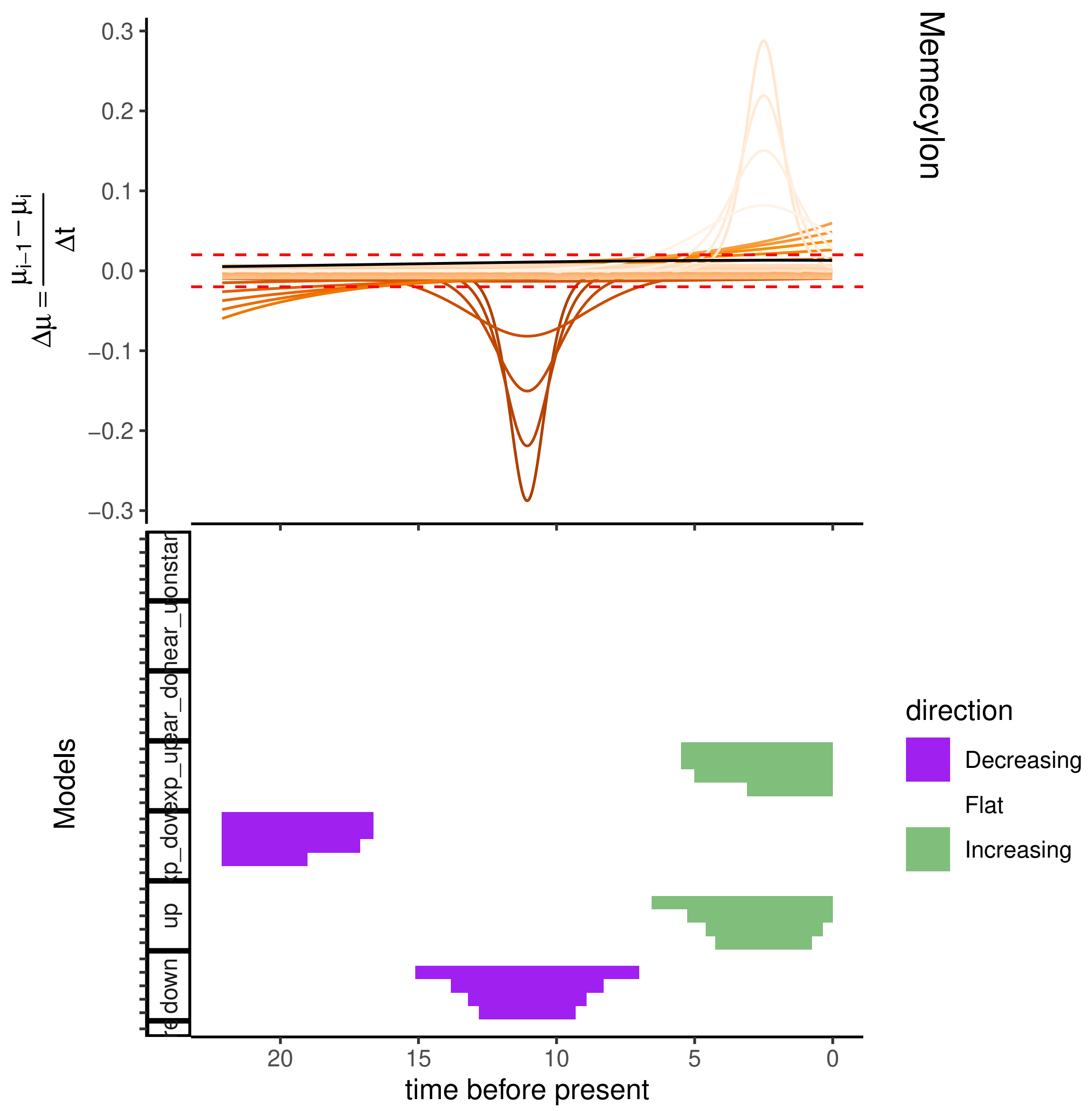
Memecylon
$\Delta\mu = \frac{\mu_{i-1} - \mu_i}{\Delta t}$
0.3
0.2
0.1
0.0
−0.1
−0.2
−0.3
Models
onstar
ear_u
ear_do
xp_up
xp_dow
up
down
20
15
10
5
0
time before present
direction
Decreasing
Flat
Increasing

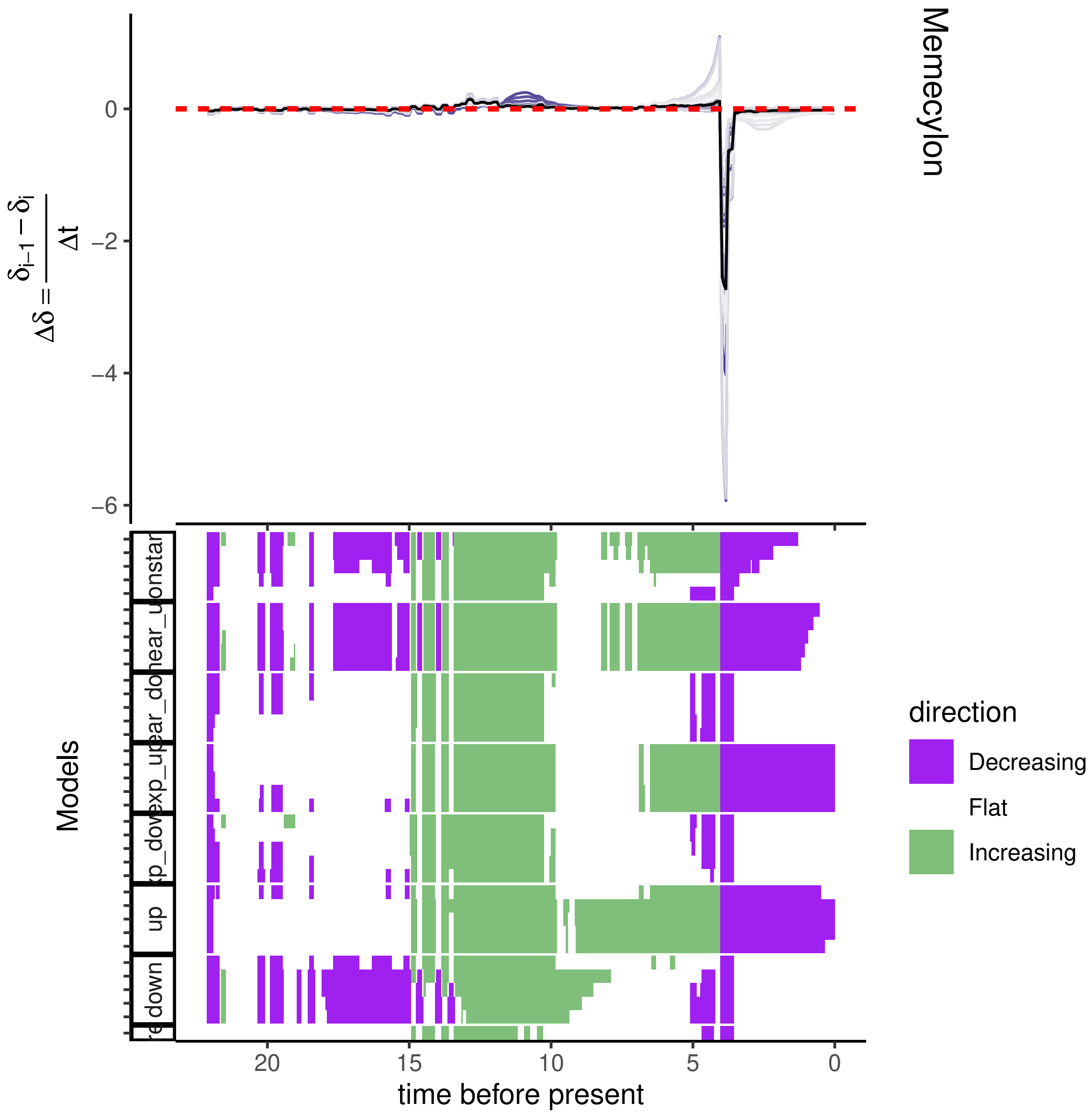

Memecylon
$\Delta\delta = \frac{\delta_{i-1} - \delta_i}{\Delta t}$
0
−2
−4
−6
Models
up
down
direction
Decreasing
Flat
Increasing
20
15
10
5
0
time before present

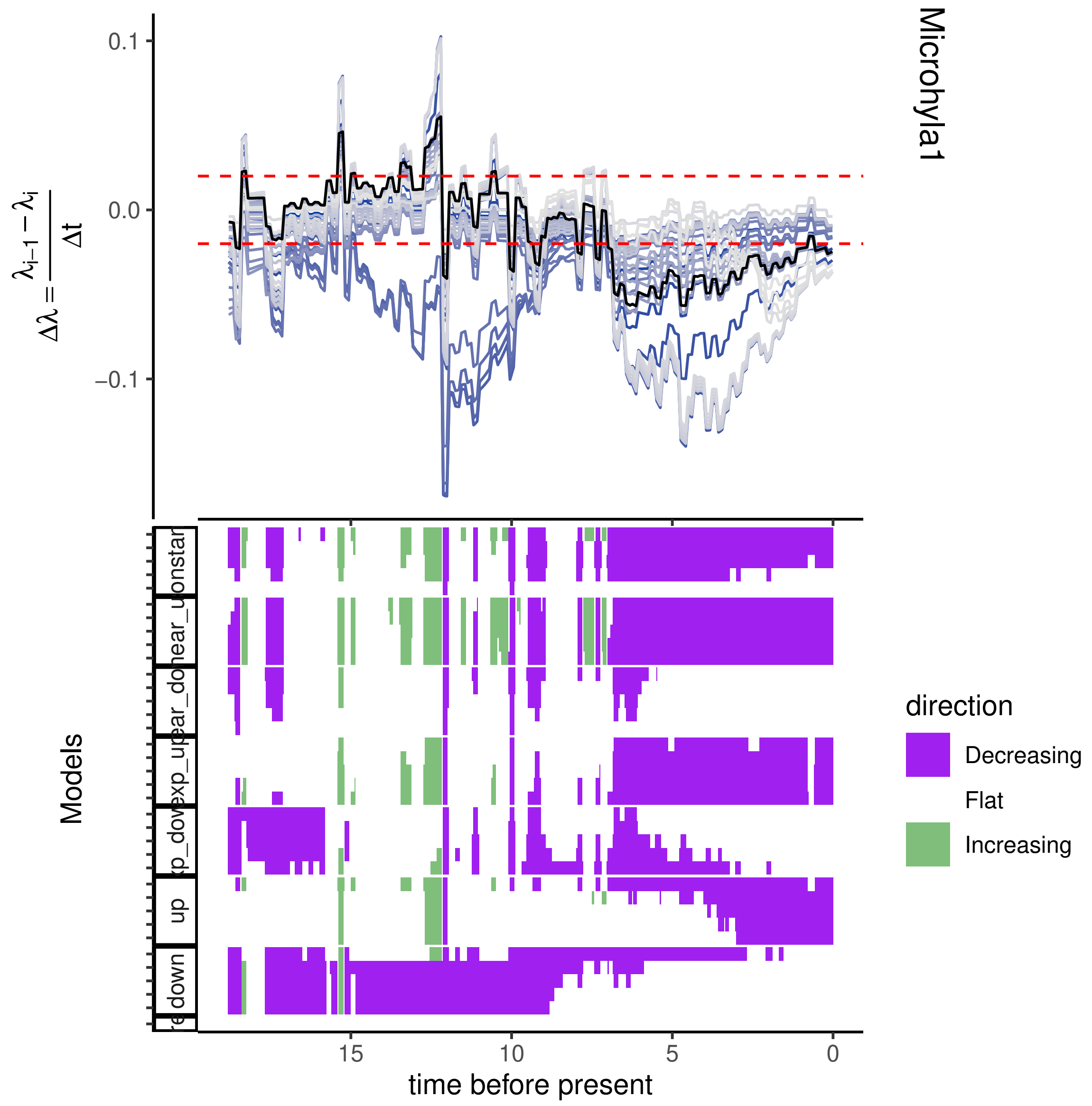
Microhyla1
0.1
0.0
−0.1
$\Delta\lambda = \frac{\lambda_{i-1} - \lambda_i}{\Delta t}$
Models
direction
Decreasing
Flat
Increasing
15
10
5
0
time before present

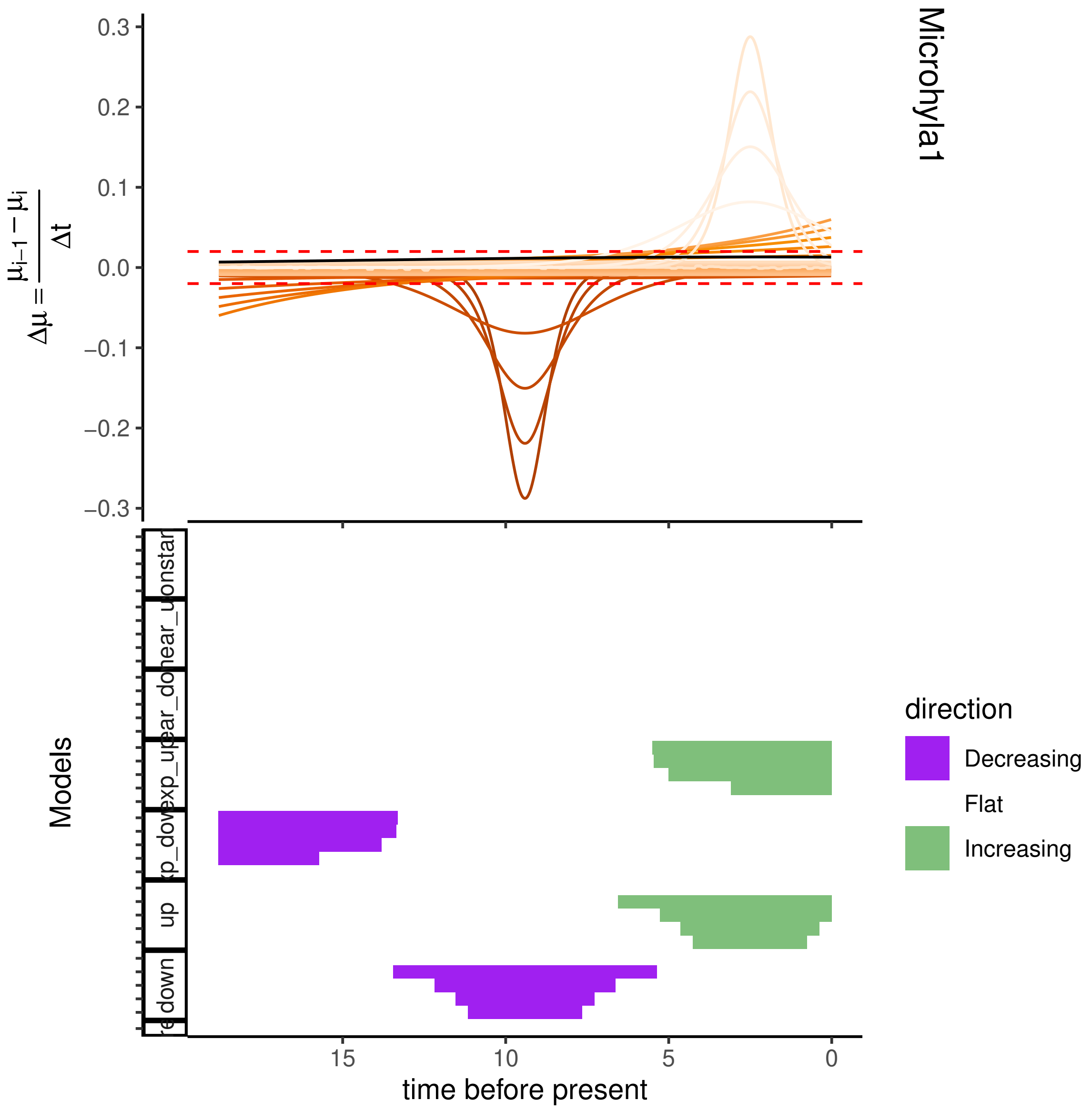
Microhyla1
0.3
0.2
0.1
0.0
−0.1
−0.2
−0.3
$\Delta\mu = \frac{\mu_{i-1} - \mu_i}{\Delta t}$
Models
down
up
direction
Decreasing
Flat
Increasing
15
10
5
0
time before present

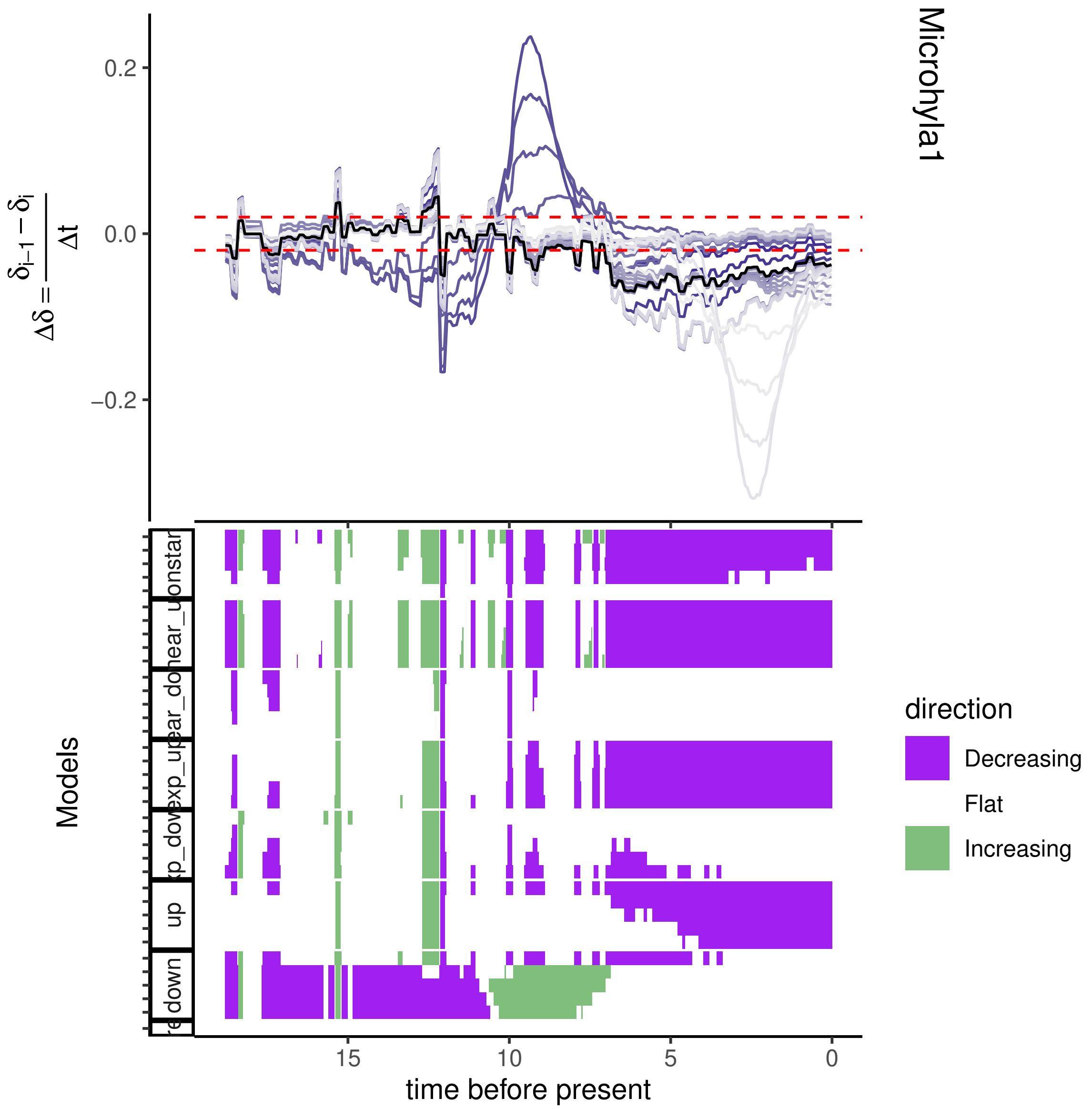
Microhyla1
$\Delta\delta = \frac{\delta_{i-1} - \delta_i}{\Delta t}$
0.2
0.0
−0.2
Models
up
down
15
10
5
0
time before present
direction
Decreasing
Flat
Increasing

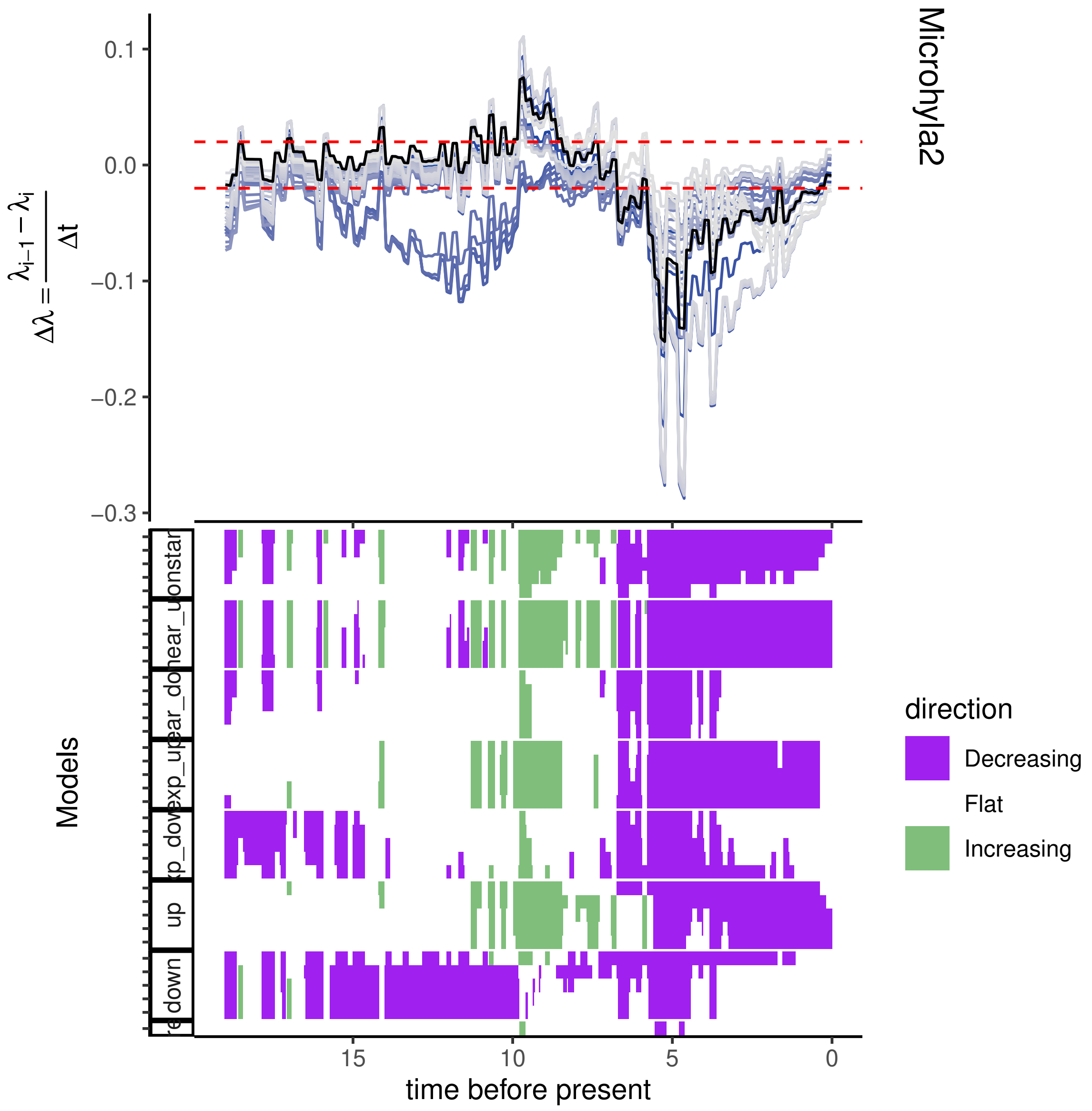
Microhyla2
0.1
0.0
−0.1
−0.2
−0.3
$\Delta\lambda = \frac{\lambda_{i-1} - \lambda_i}{\Delta t}$
Models
direction
Decreasing
Flat
Increasing
15
10
5
0
time before present
up
down

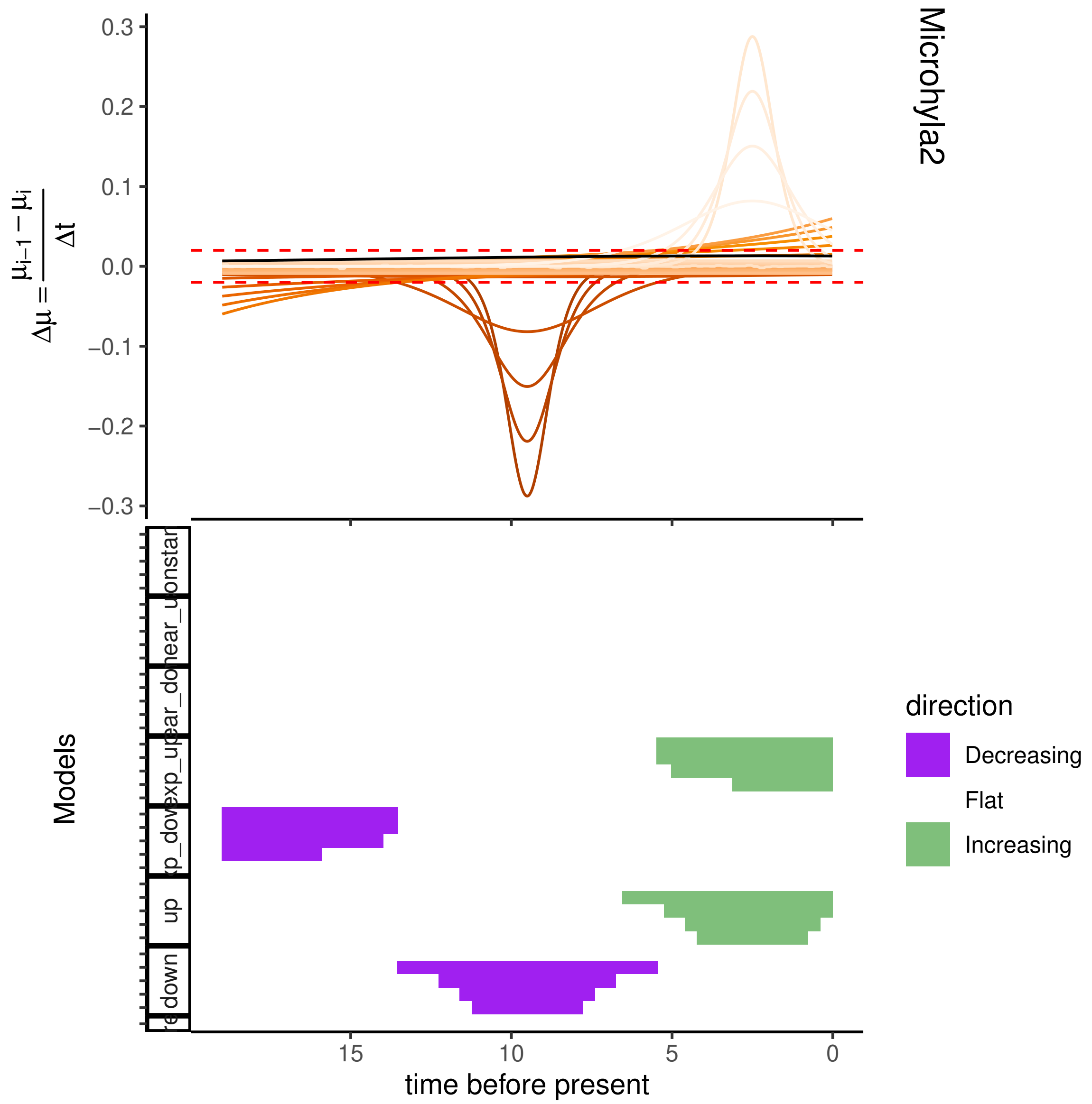

Microhyla2
$\Delta\mu = \frac{\mu_{i-1} - \mu_i}{\Delta t}$
0.3
0.2
0.1
0.0
−0.1
−0.2
−0.3
Models
up
down
direction
Decreasing
Flat
Increasing
15
10
5
0
time before present

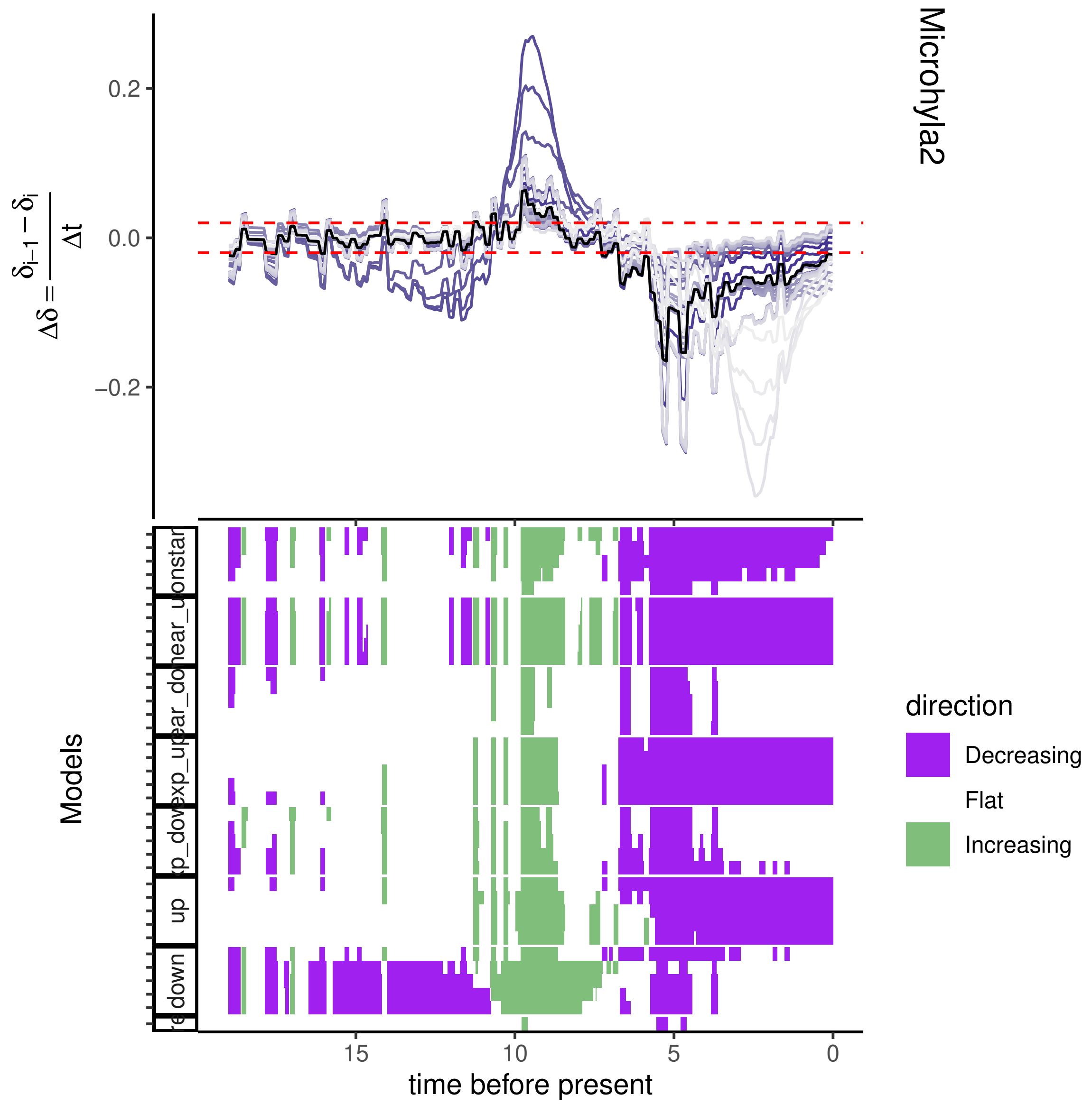
Microhyla2
$\Delta\delta = \frac{\delta_{i-1} - \delta_i}{\Delta t}$
0.2
0.0
−0.2
Models
up
down
direction
Decreasing
Flat
Increasing
15
10
5
0
time before present

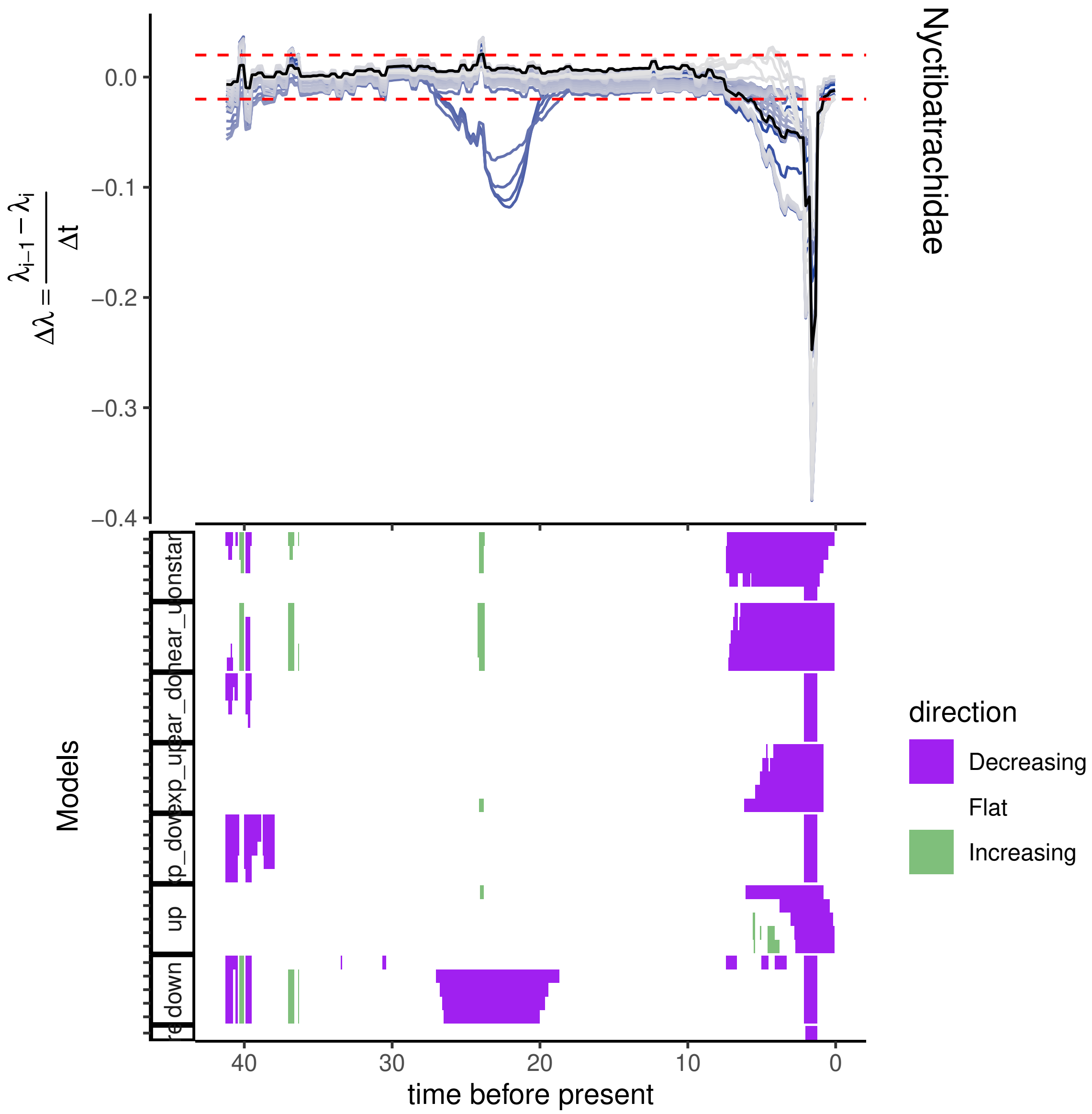
Nyctibatrachidae
$\Delta\lambda = \frac{\lambda_{i-1} - \lambda_i}{\Delta t}$
0.0
−0.1
−0.2
−0.3
−0.4
Models
direction
Decreasing
Flat
Increasing
40
30
20
10
0
time before present

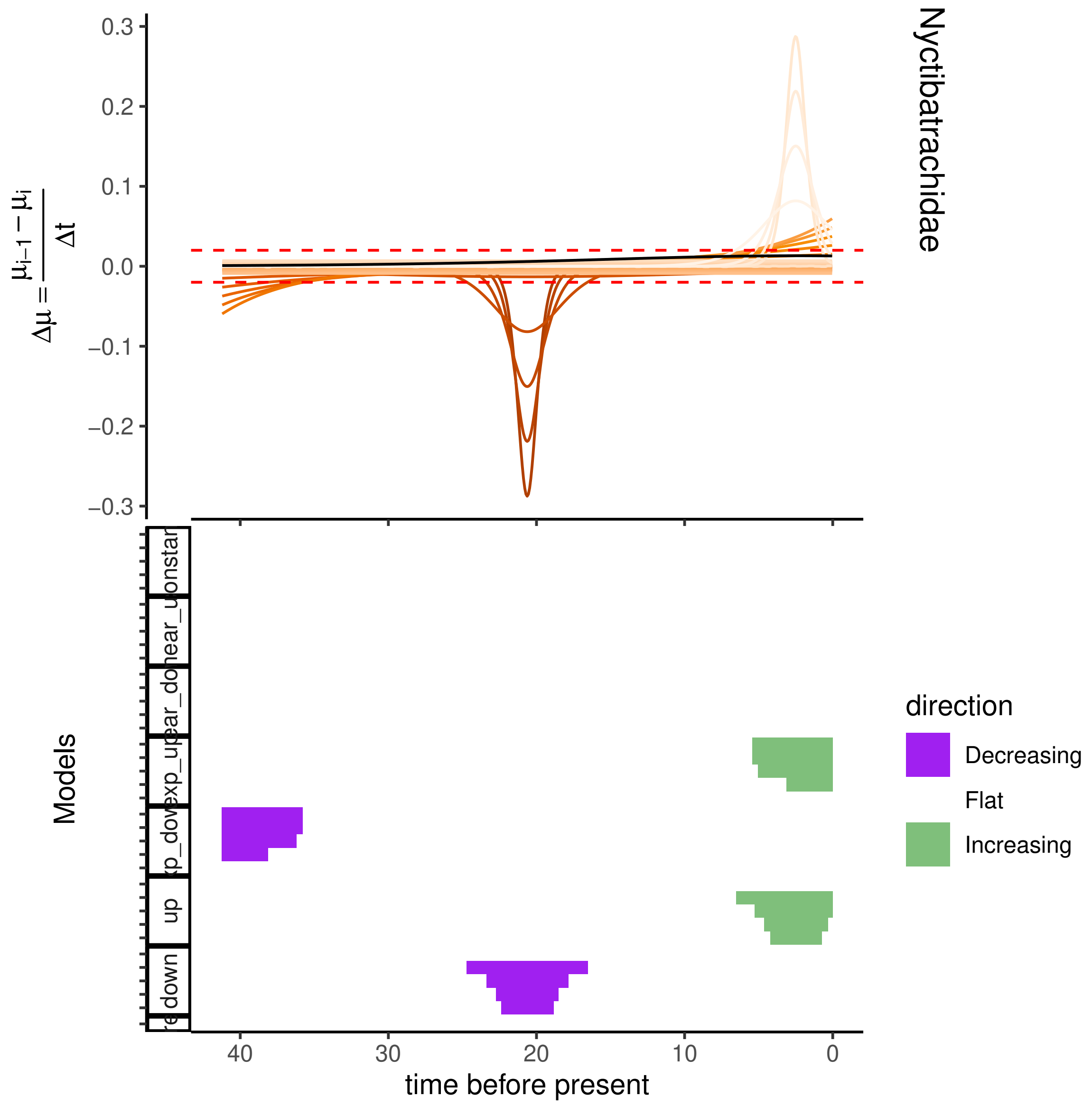

Nyctibatrachidae
0.3
0.2
0.1
0.0
−0.1
−0.2
−0.3
$\Delta\mu = \frac{\mu_{i-1} - \mu_i}{\Delta t}$
Models
40
30
20
10
0
time before present
direction
Decreasing
Flat
Increasing
up
down

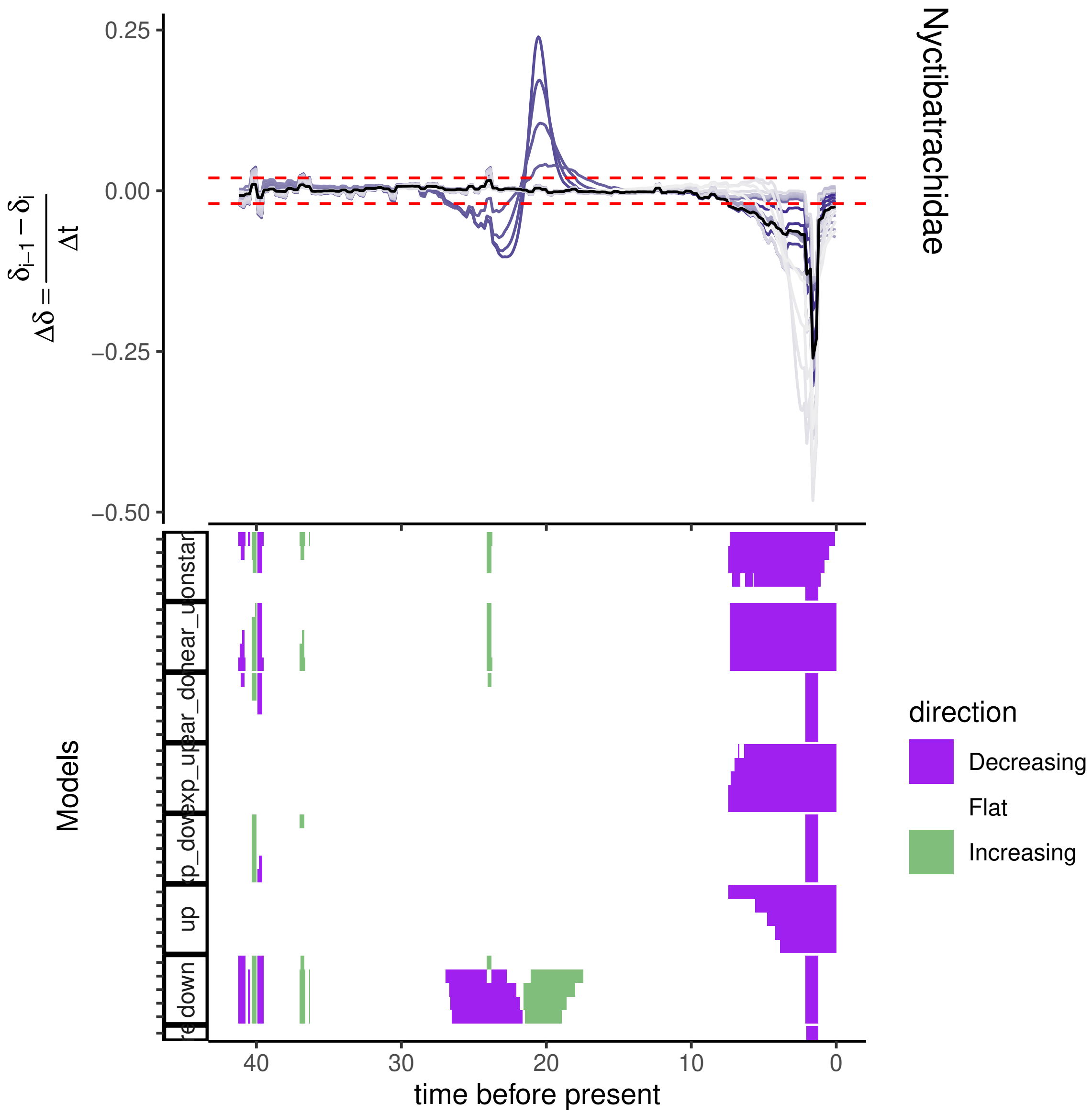
Nyctibatrachidae
0.25
0.00
−0.25
−0.50
$\Delta\delta = \frac{\delta_{i-1} - \delta_i}{\Delta t}$
Models
onstar
near_do
exp_up
ar_do
xp_dov
up
down
40
30
20
10
0
time before present
direction
Decreasing
Flat
Increasing

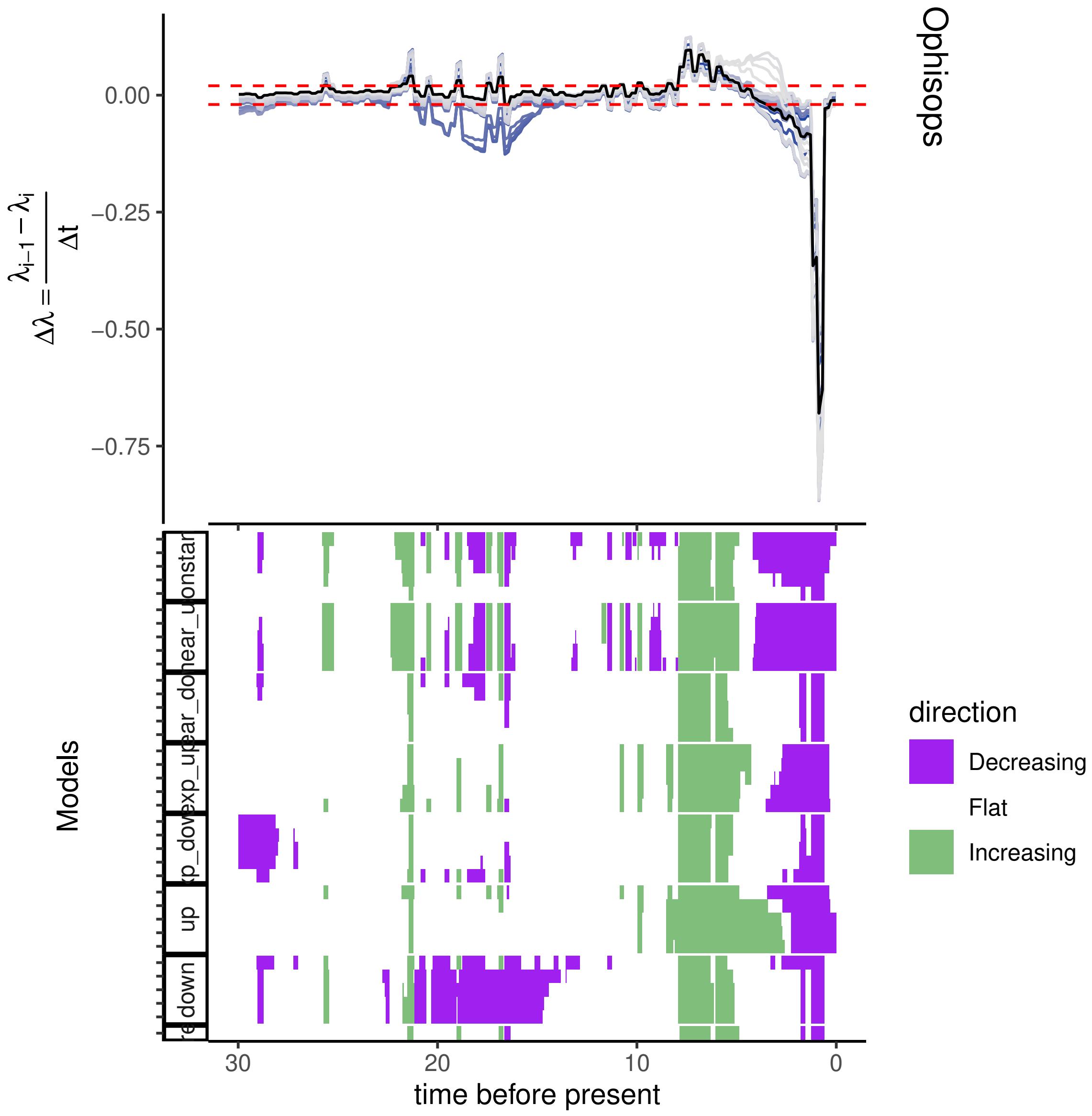

Ophisops
0.00
−0.25
−0.50
−0.75
Models
up
down
30
20
10
0
time before present
direction
Decreasing
Flat
Increasing

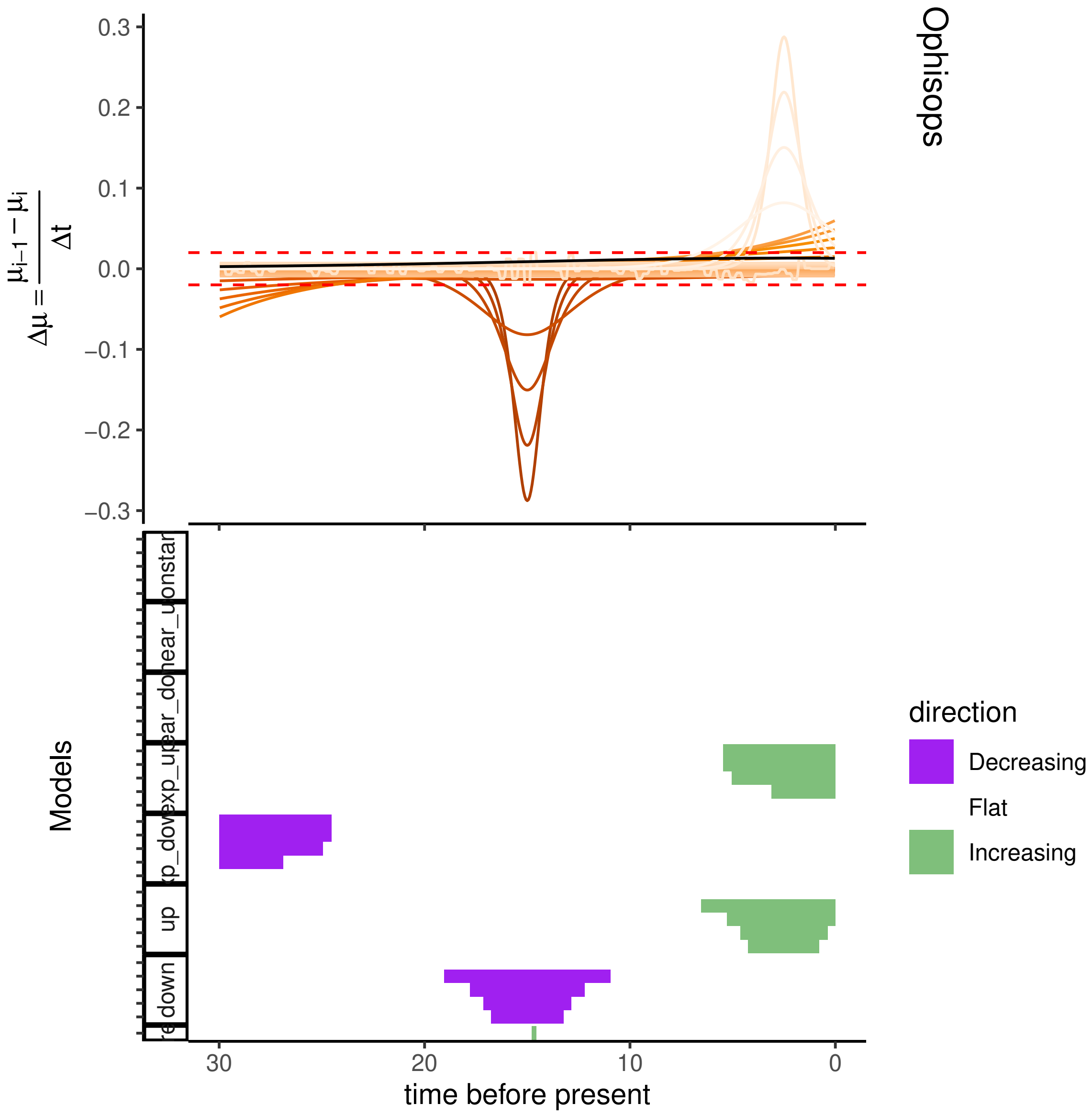

Ophisops
0.3
0.2
0.1
0.0
−0.1
−0.2
−0.3
$\Delta\mu = \frac{\mu_{i-1} - \mu_i}{\Delta t}$
Models
onstar
near_u
ear_do
exp_up
xp_dov
up
down
30
20
10
0
time before present
direction
Decreasing
Flat
Increasing

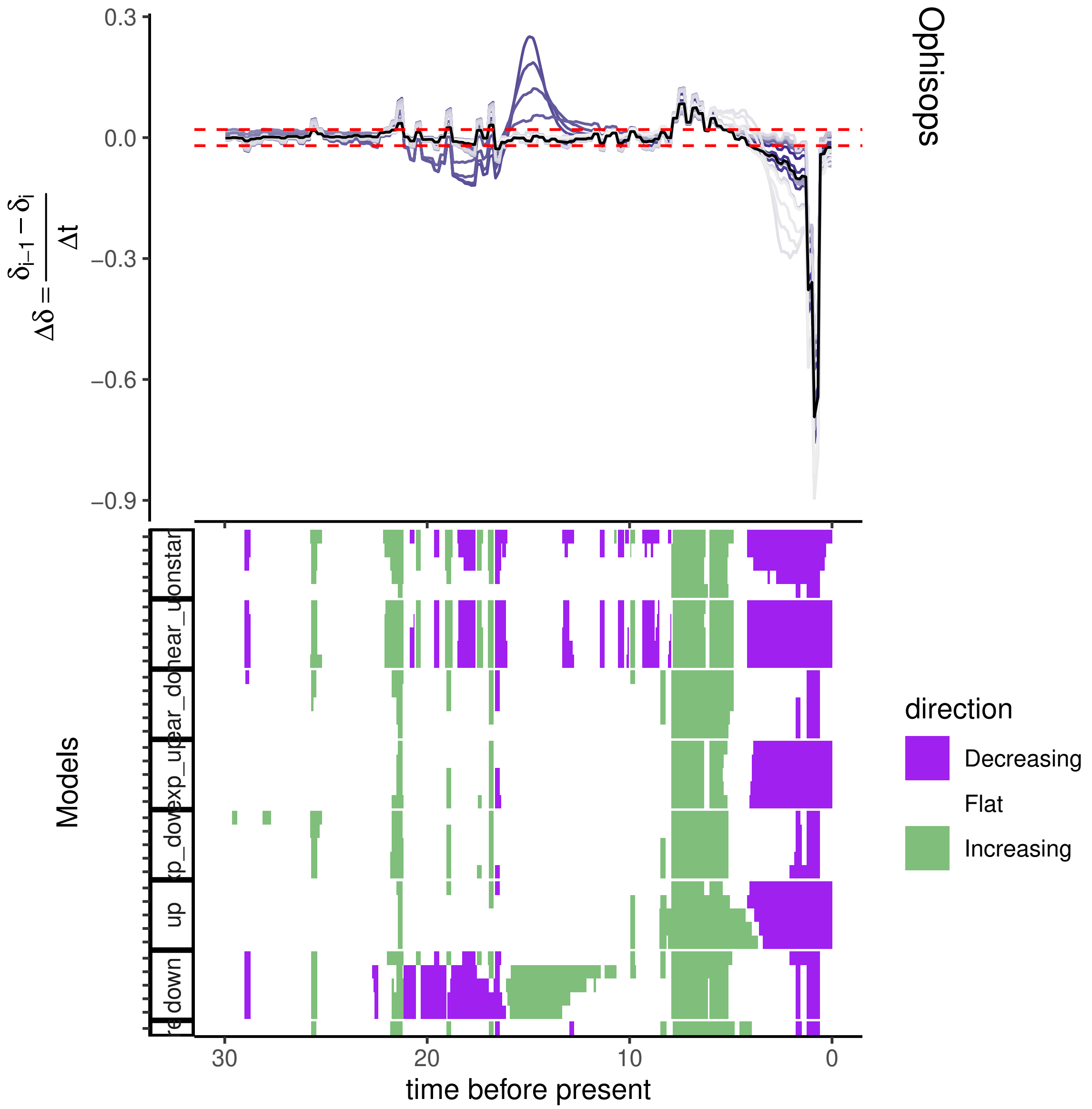

Ophisops
0.3
0.0
−0.3
−0.6
−0.9
$\Delta\delta = \frac{\delta_{i-1} - \delta_i}{\Delta t}$
Models
up
down
30
20
10
0
time before present
direction
Decreasing
Flat
Increasing

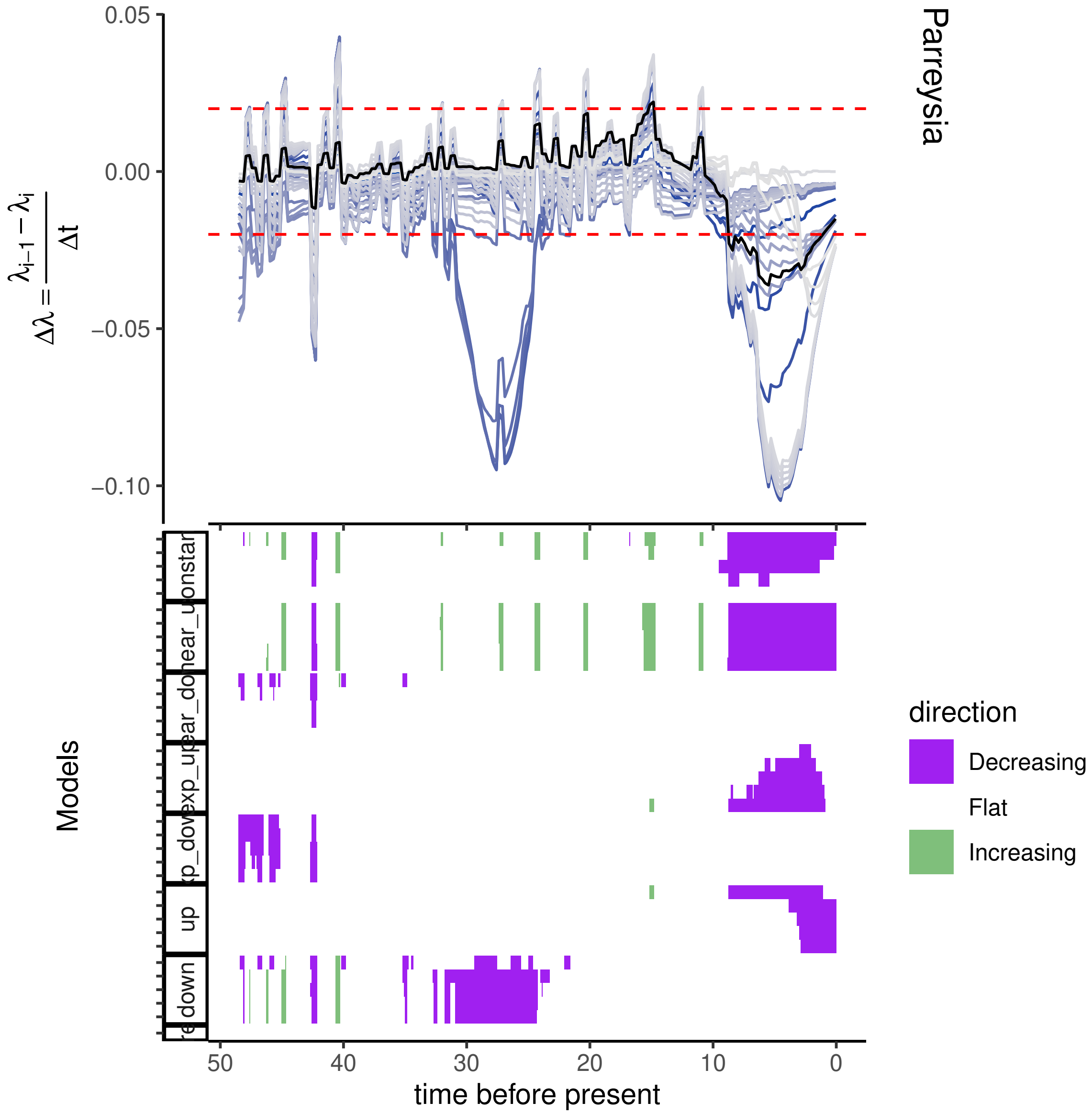

Parreysia
0.05
0.00
−0.05
−0.10
$\Delta\lambda = \frac{\lambda_{i-1} - \lambda_i}{\Delta t}$
Models
down
up
time before present
50
40
30
20
10
0
direction
Decreasing
Flat
Increasing

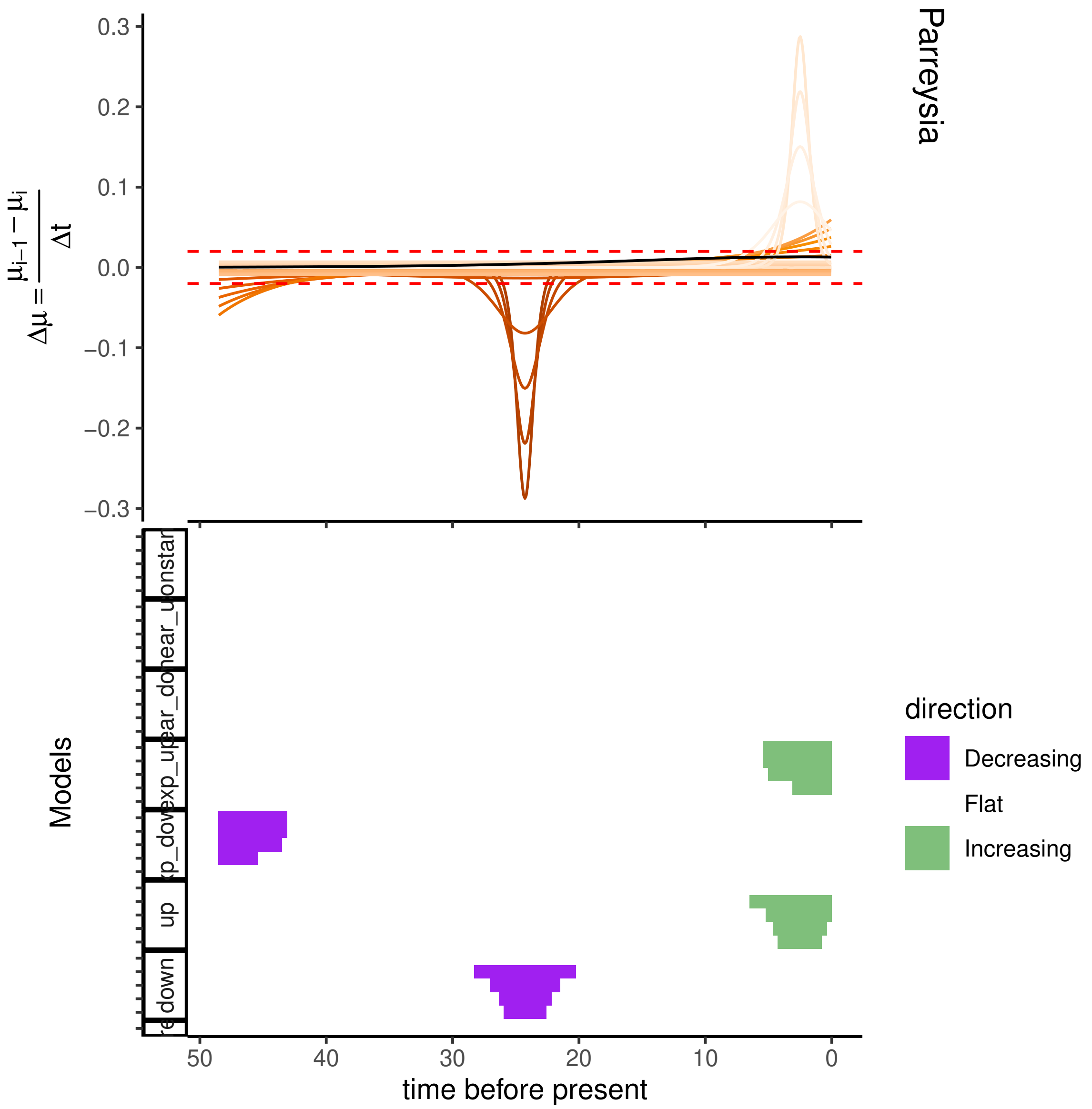
Parreysia
$\Delta\mu = \frac{\mu_{i-1} - \mu_i}{\Delta t}$
0.3
0.2
0.1
0.0
−0.1
−0.2
−0.3
Models
up
down
50
40
30
20
10
0
time before present
direction
Decreasing
Flat
Increasing

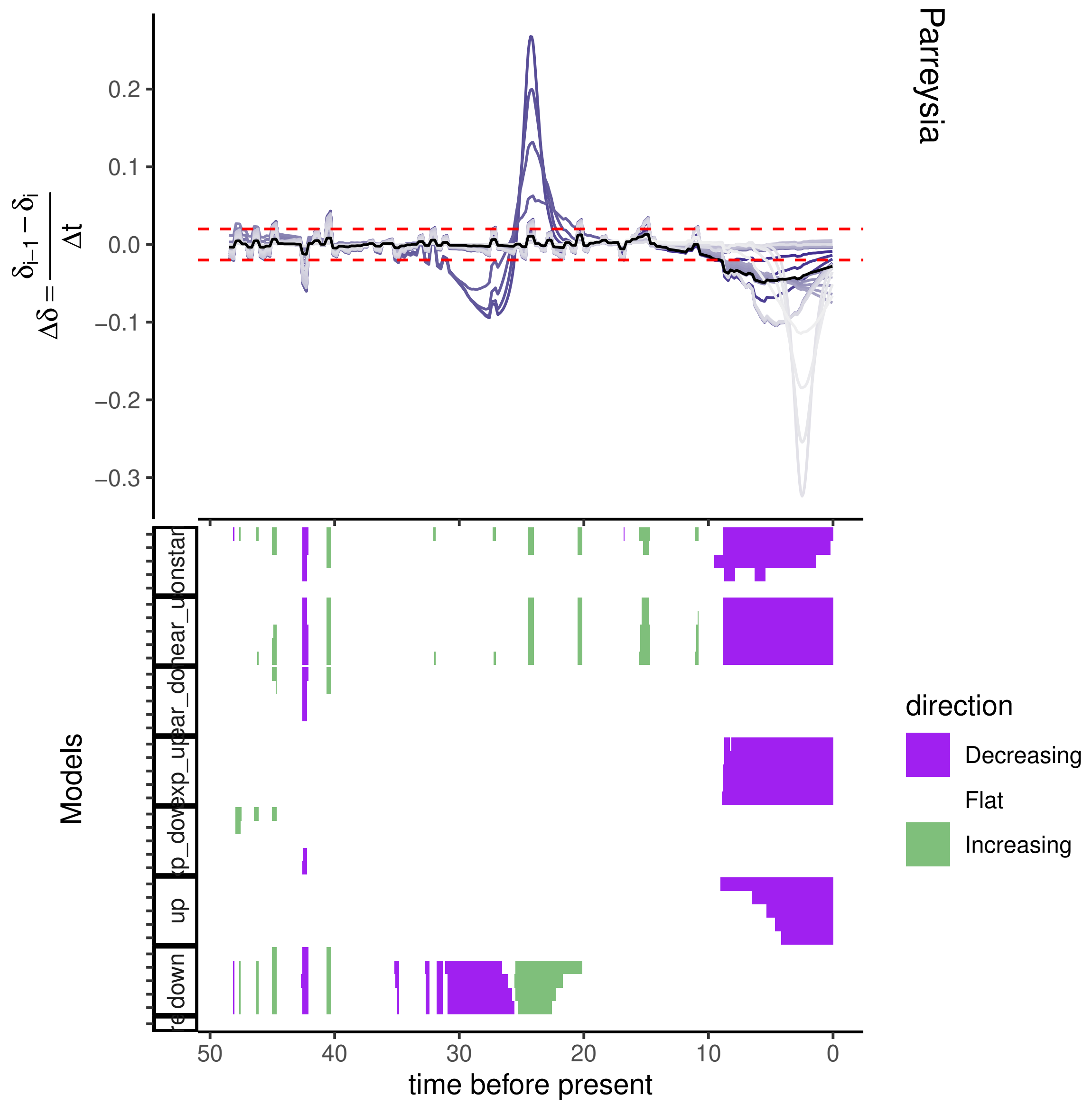

Parreysia
$\Delta\delta = \frac{\delta_{i-1} - \delta_i}{\Delta t}$
0.2
0.1
0.0
−0.1
−0.2
−0.3
Models
up
down
direction
Decreasing
Flat
Increasing
50
40
30
20
10
0
time before present

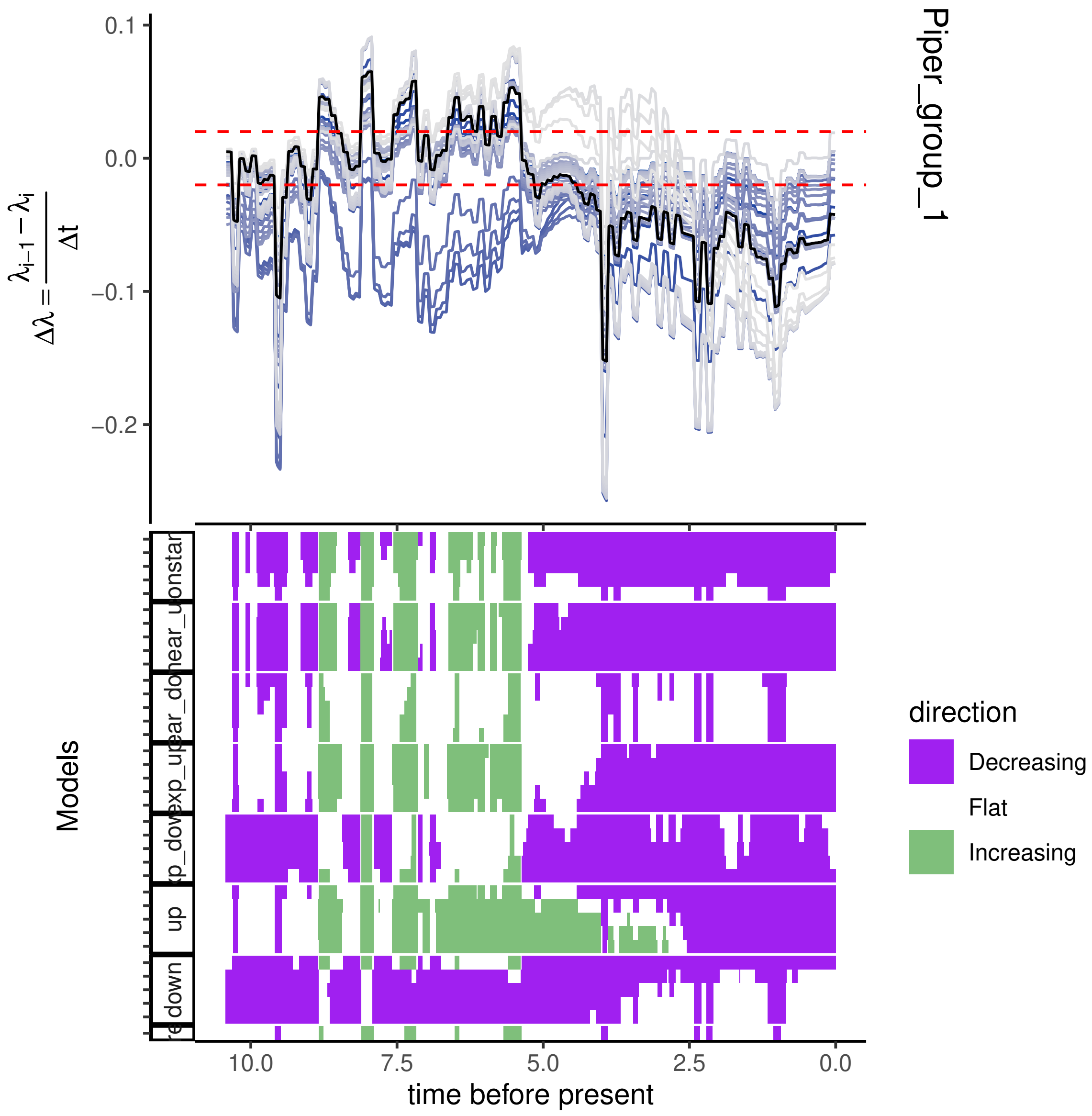

0.1
0.0
−0.1
−0.2
$\Delta\lambda = \frac{\lambda_{i-1} - \lambda_i}{\Delta t}$
Piper_group_1
Models
direction
Decreasing
Flat
Increasing
10.0
7.5
5.0
2.5
0.0
time before present

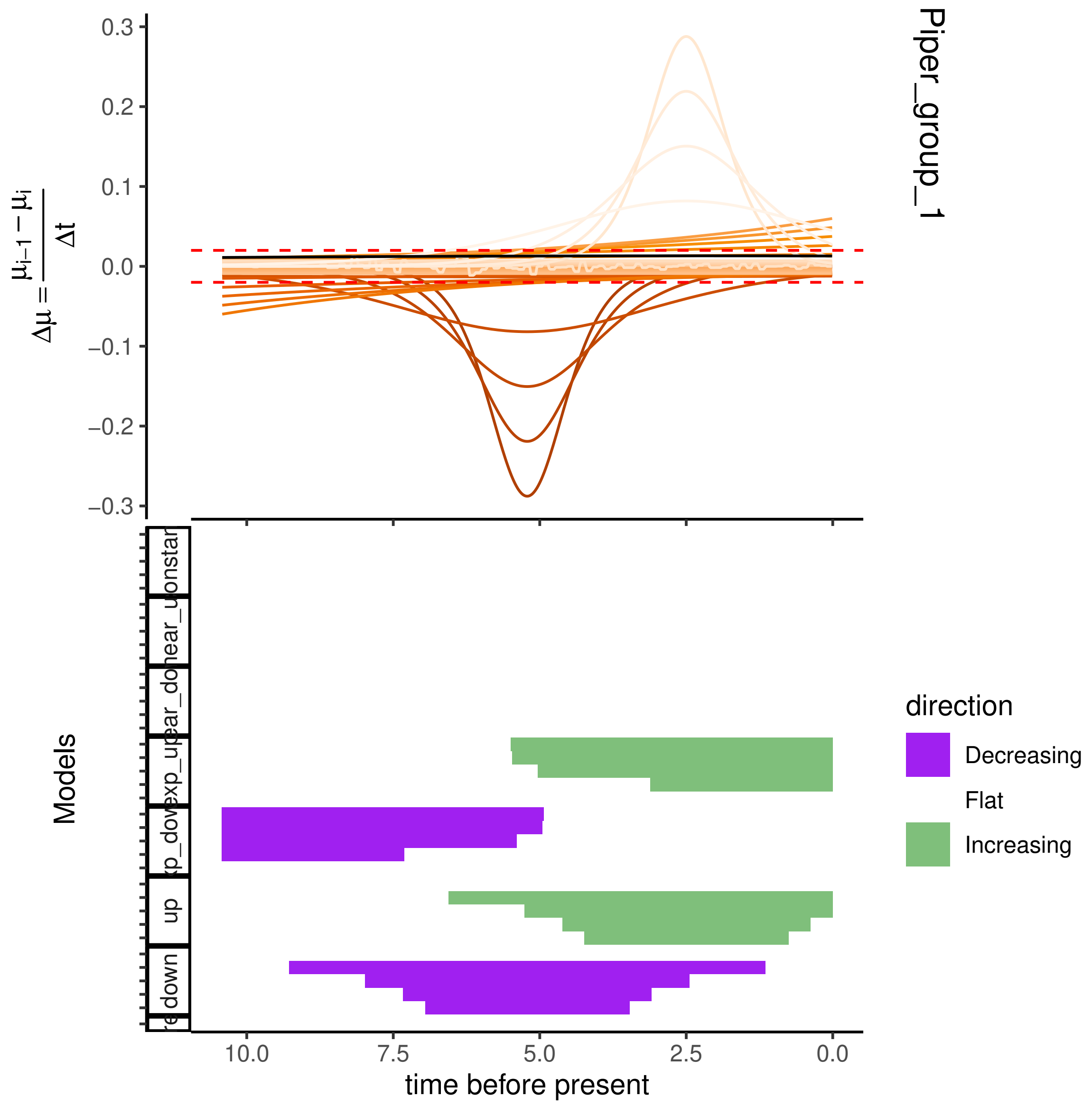
Piper_group_1
$\Delta\mu = \frac{\mu_{i-1} - \mu_i}{\Delta t}$
0.3
0.2
0.1
0.0
−0.1
−0.2
−0.3
Models
direction
Decreasing
Flat
Increasing
up
down
10.0
7.5
5.0
2.5
0.0
time before present

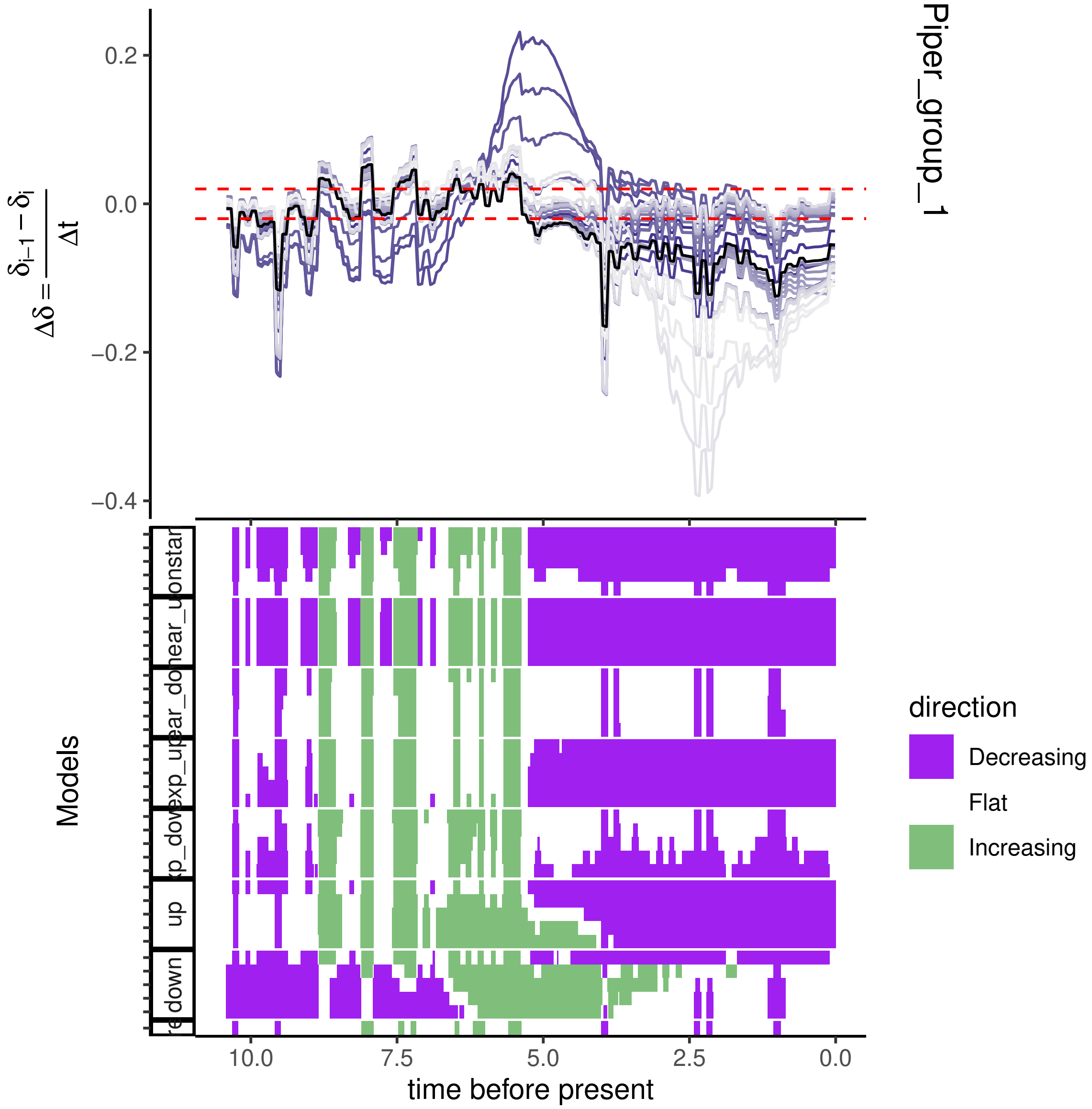
Piper_group_1
0.2
0.0
−0.2
−0.4
$\Delta\delta = \frac{\delta_{i-1} - \delta_i}{\Delta t}$
Models
10.0
7.5
5.0
2.5
0.0
time before present
direction
Decreasing
Flat
Increasing

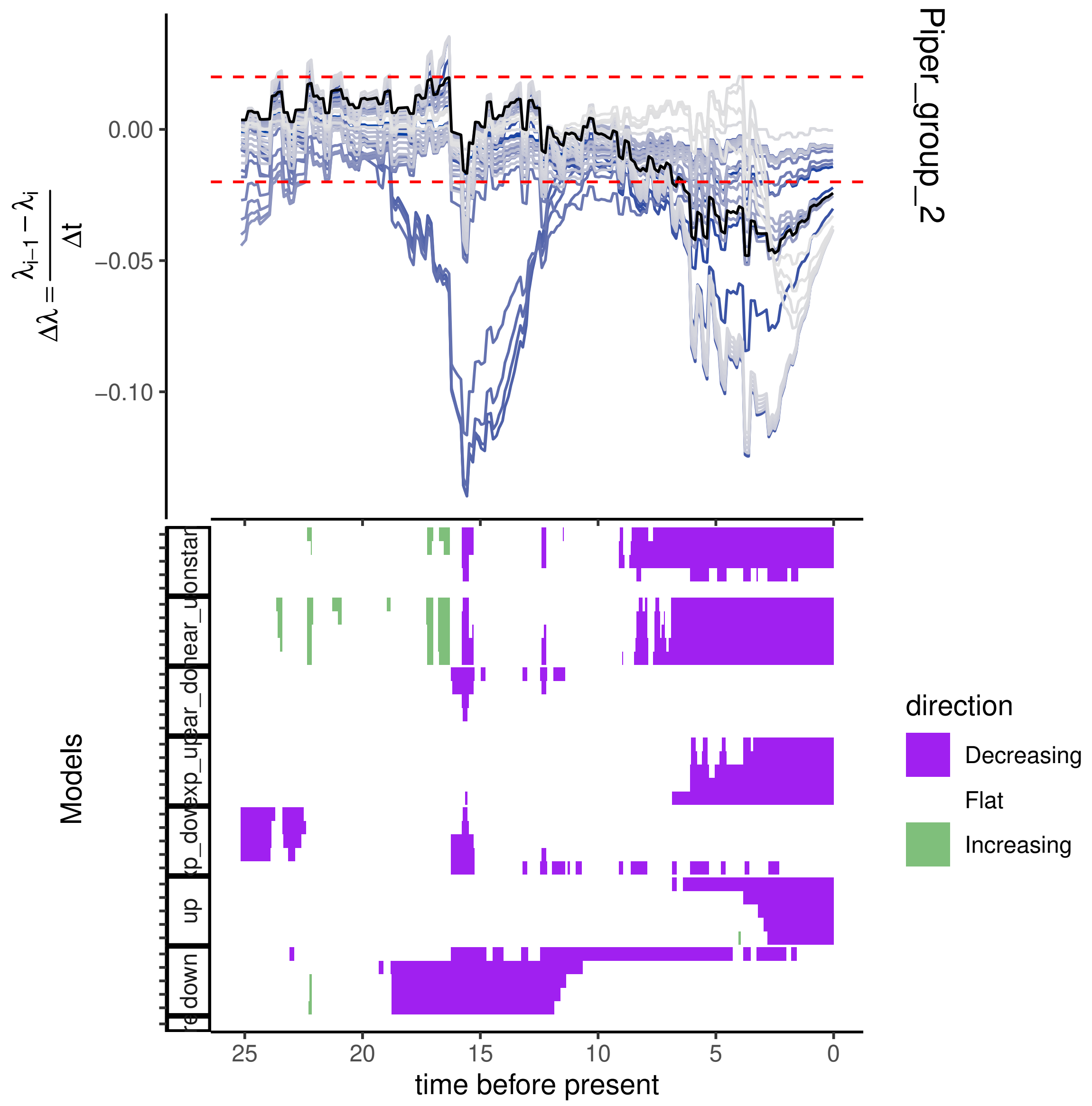

Piper_group_2
$\Delta\lambda = \frac{\lambda_{i-1} - \lambda_i}{\Delta t}$
0.00
−0.05
−0.10
Models
direction
Decreasing
Flat
Increasing
25
20
15
10
5
0
time before present
up
down

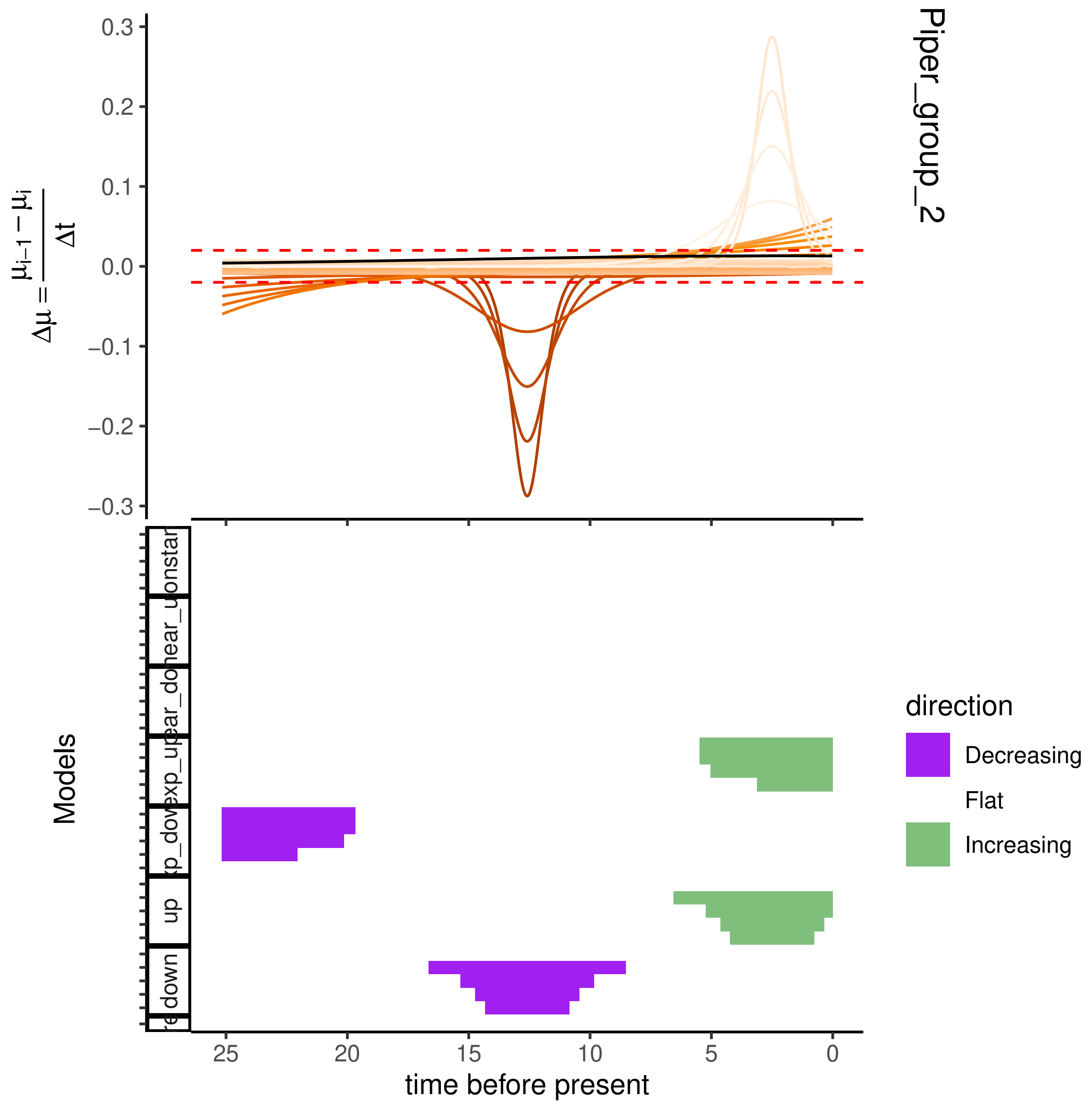
Piper_group_2
$\Delta\mu = \frac{\mu_{i-1} - \mu_i}{\Delta t}$
0.3
0.2
0.1
0.0
−0.1
−0.2
−0.3
Models
direction
Decreasing
Flat
Increasing
25
20
15
10
5
0
time before present
up
down

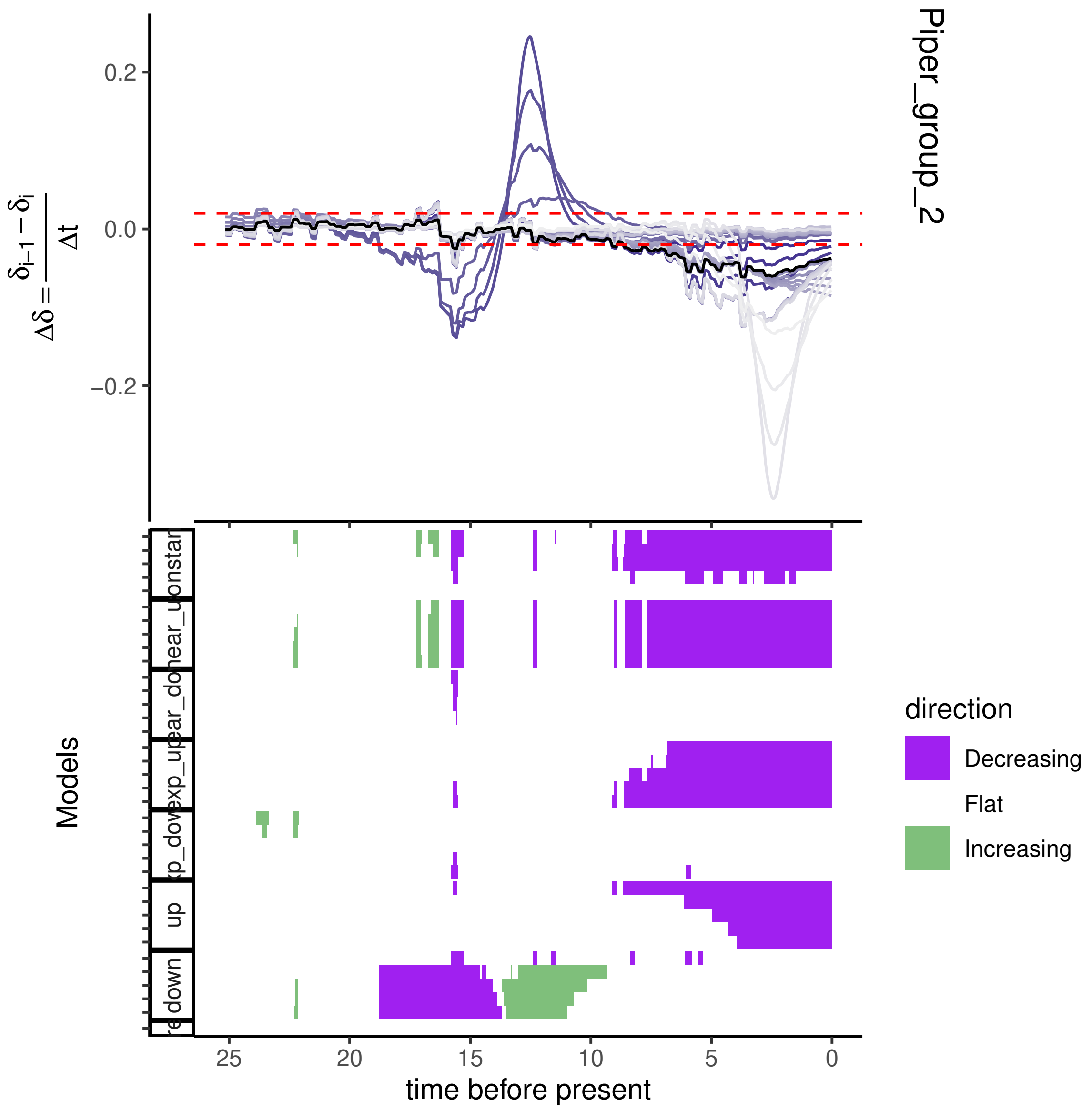

Piper_group_2
0.2
0.0
−0.2
$\Delta\delta = \frac{\delta_{i-1} - \delta_i}{\Delta t}$
Models
direction
Decreasing
Flat
Increasing
25
20
15
10
5
0
time before present

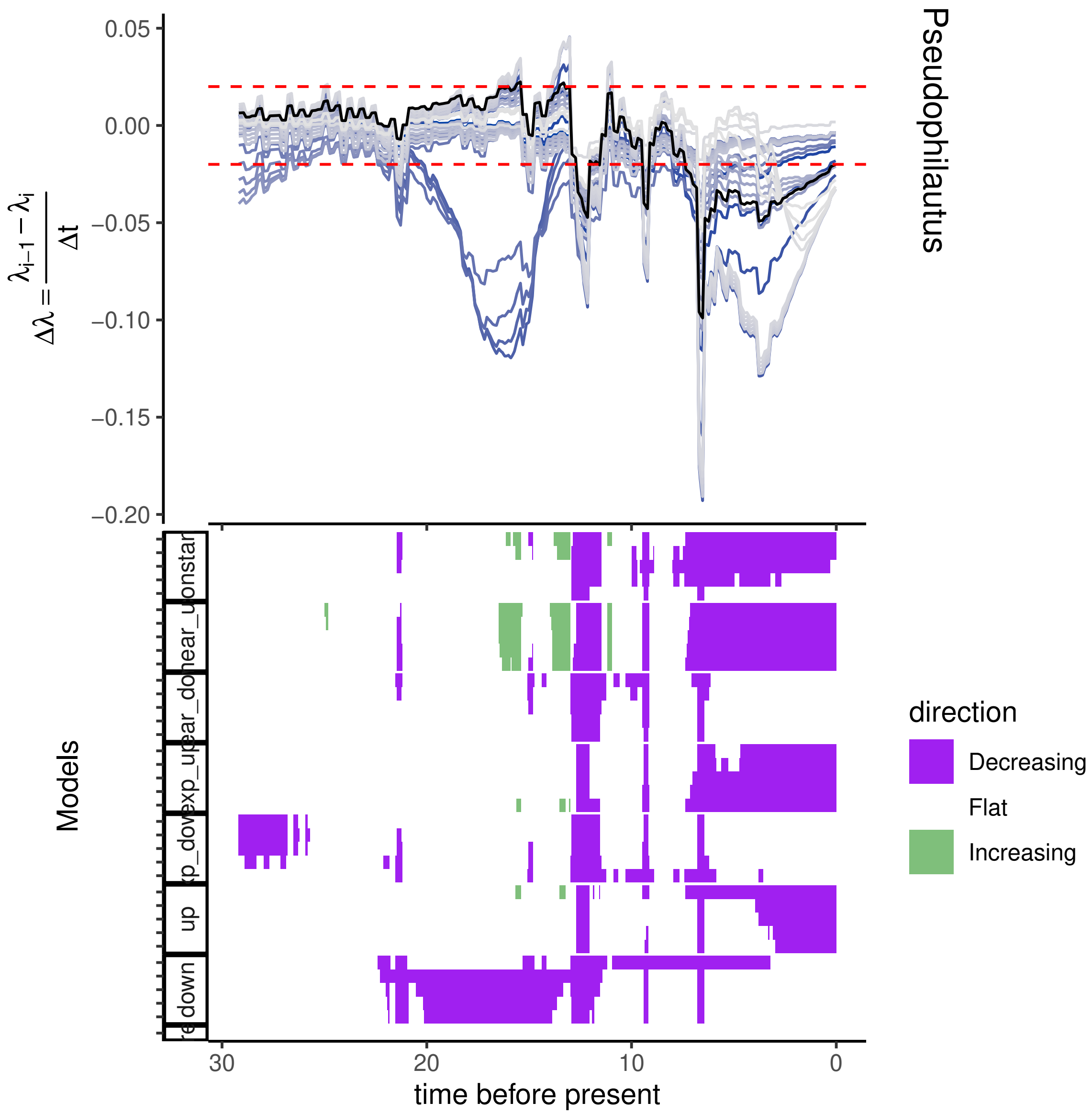

0.05
0.00
−0.05
−0.10
−0.15
−0.20
$\Delta\lambda = \frac{\lambda_{i-1} - \lambda_i}{\Delta t}$
Pseudophilautus
Models
up
down
30
20
10
0
time before present
direction
Decreasing
Flat
Increasing

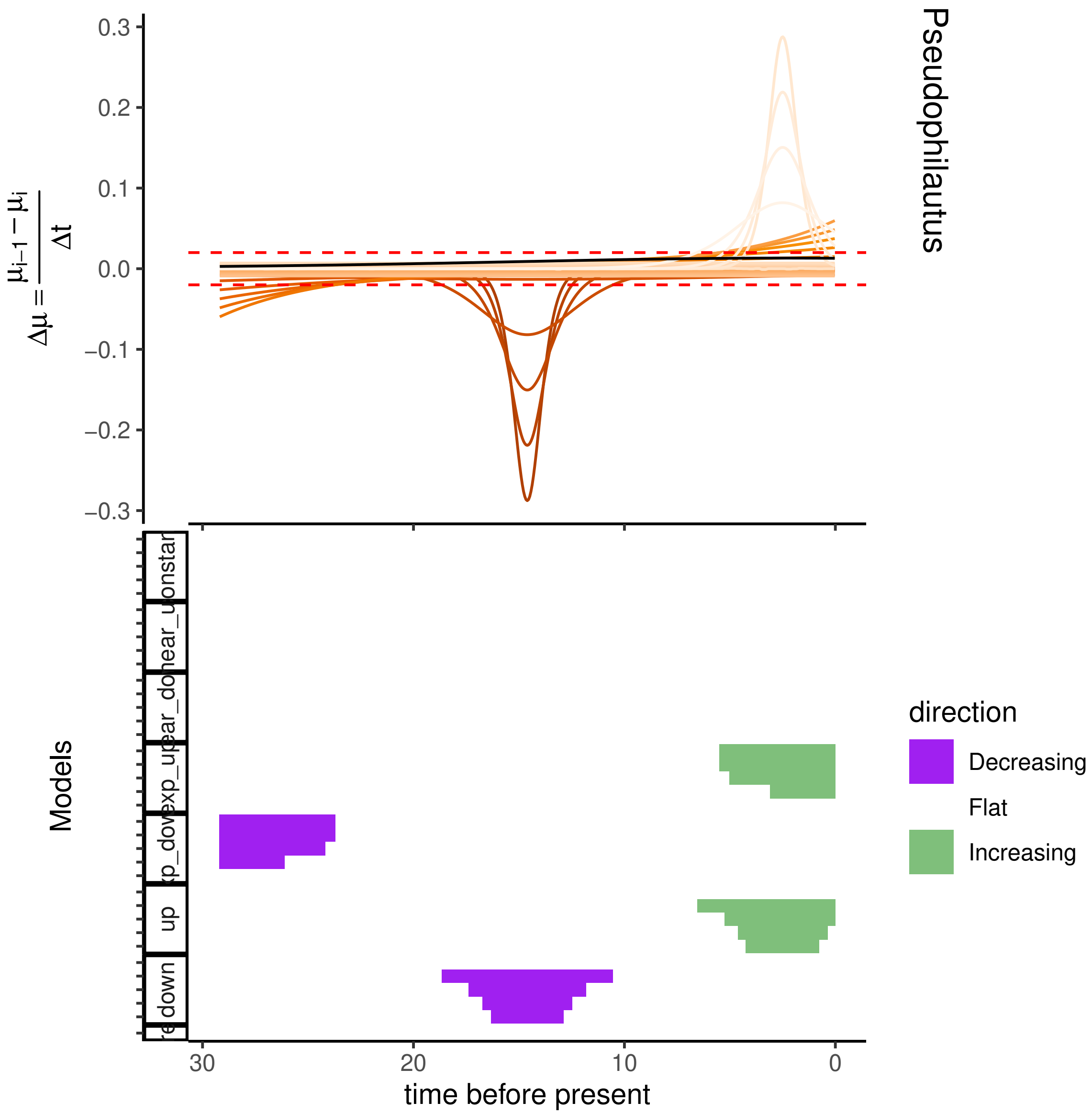
Pseudophilautus
$\Delta\mu = \frac{\mu_{i-1} - \mu_i}{\Delta t}$
0.3
0.2
0.1
0.0
−0.1
−0.2
−0.3
Models
up
down
30
20
10
0
time before present
direction
Decreasing
Flat
Increasing

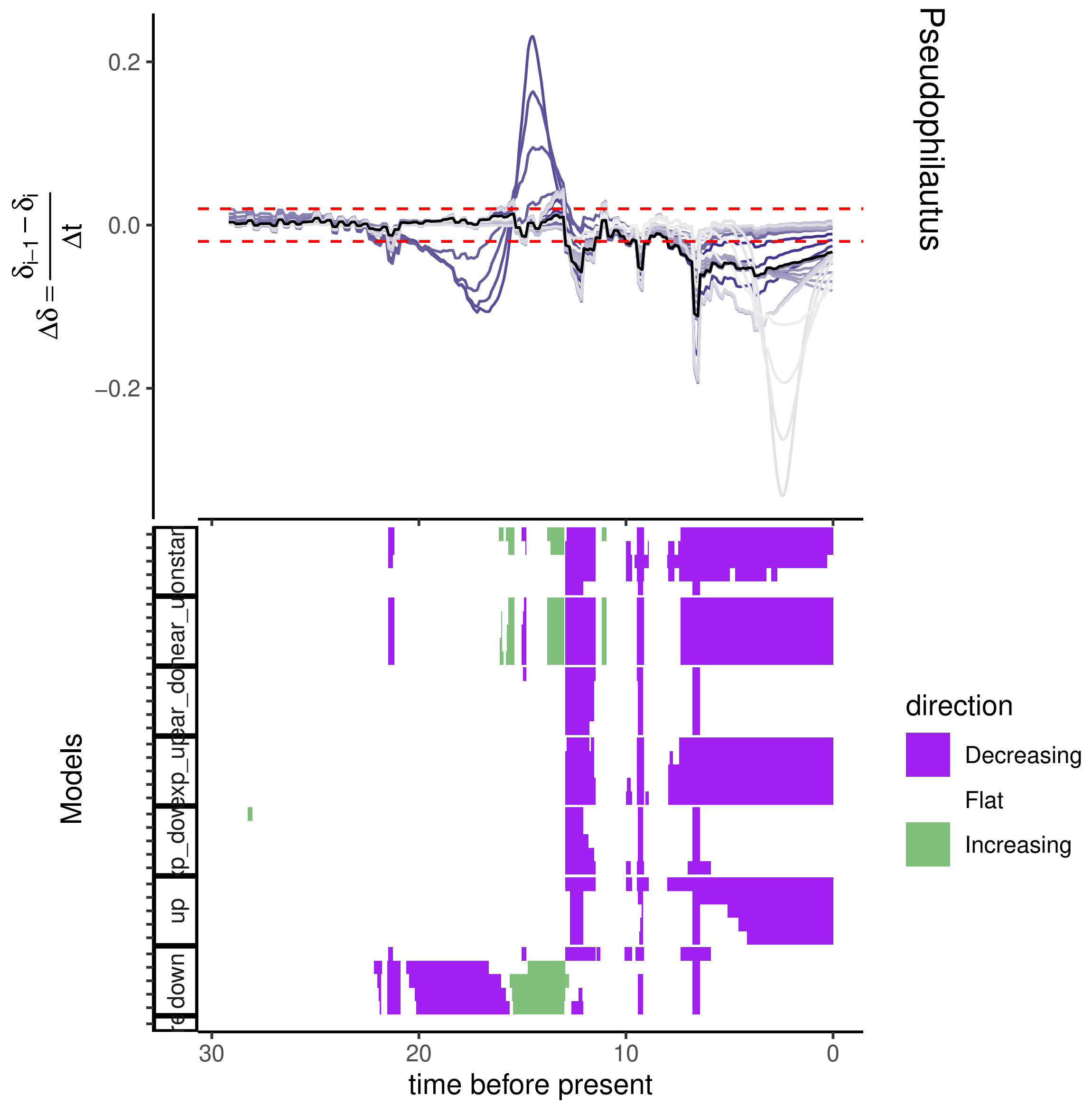

Pseudophilautus
$\Delta\delta = \frac{\delta_{i-1} - \delta_i}{\Delta t}$
0.2
0.0
−0.2
Models
up
down
direction
Decreasing
Flat
Increasing
30
20
10
0
time before present

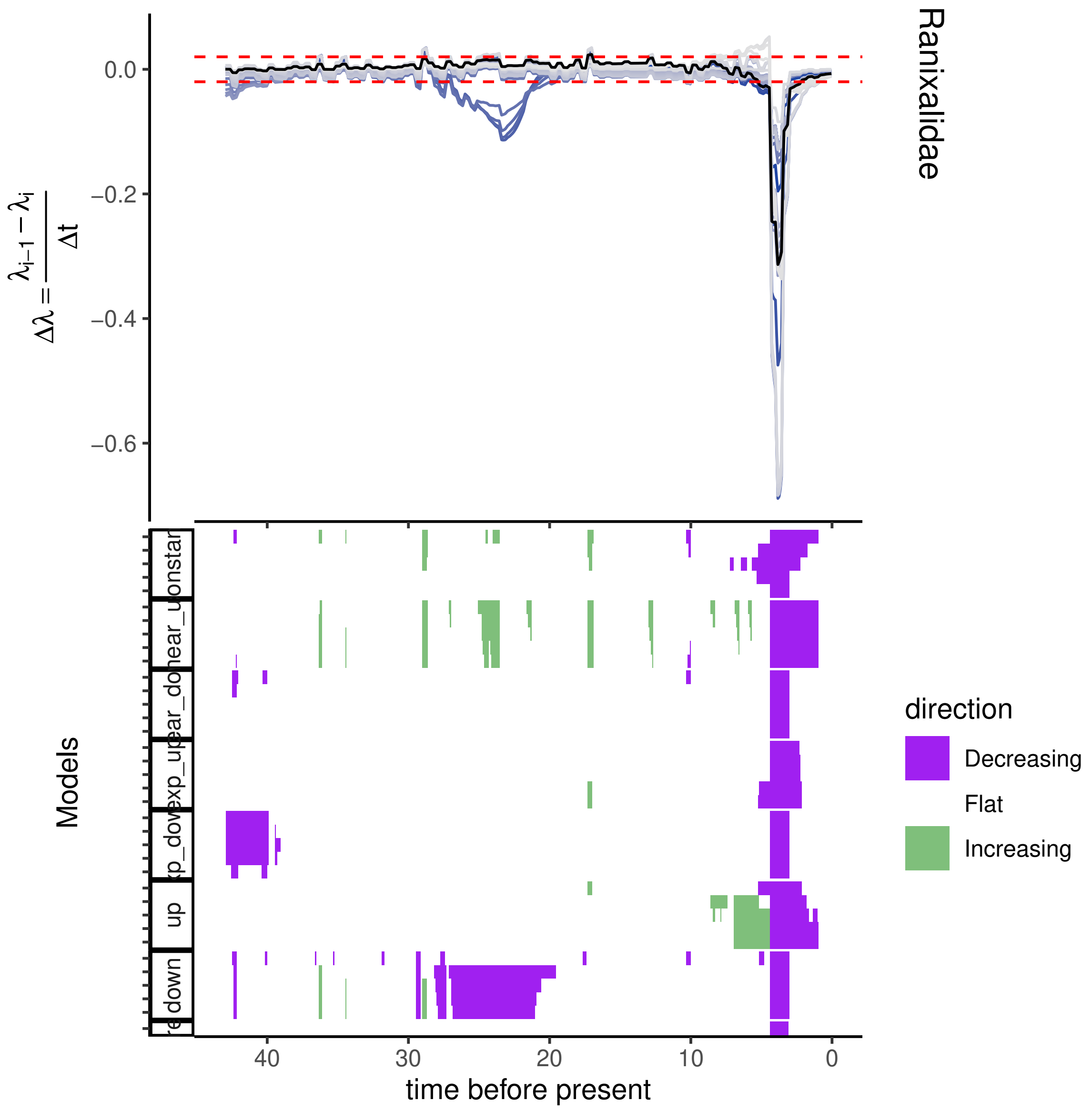

Ranixalidae
$\Delta\lambda = \frac{\lambda_{i-1} - \lambda_i}{\Delta t}$
0.0
−0.2
−0.4
−0.6
Models
time before present
40
30
20
10
0
direction
Decreasing
Flat
Increasing

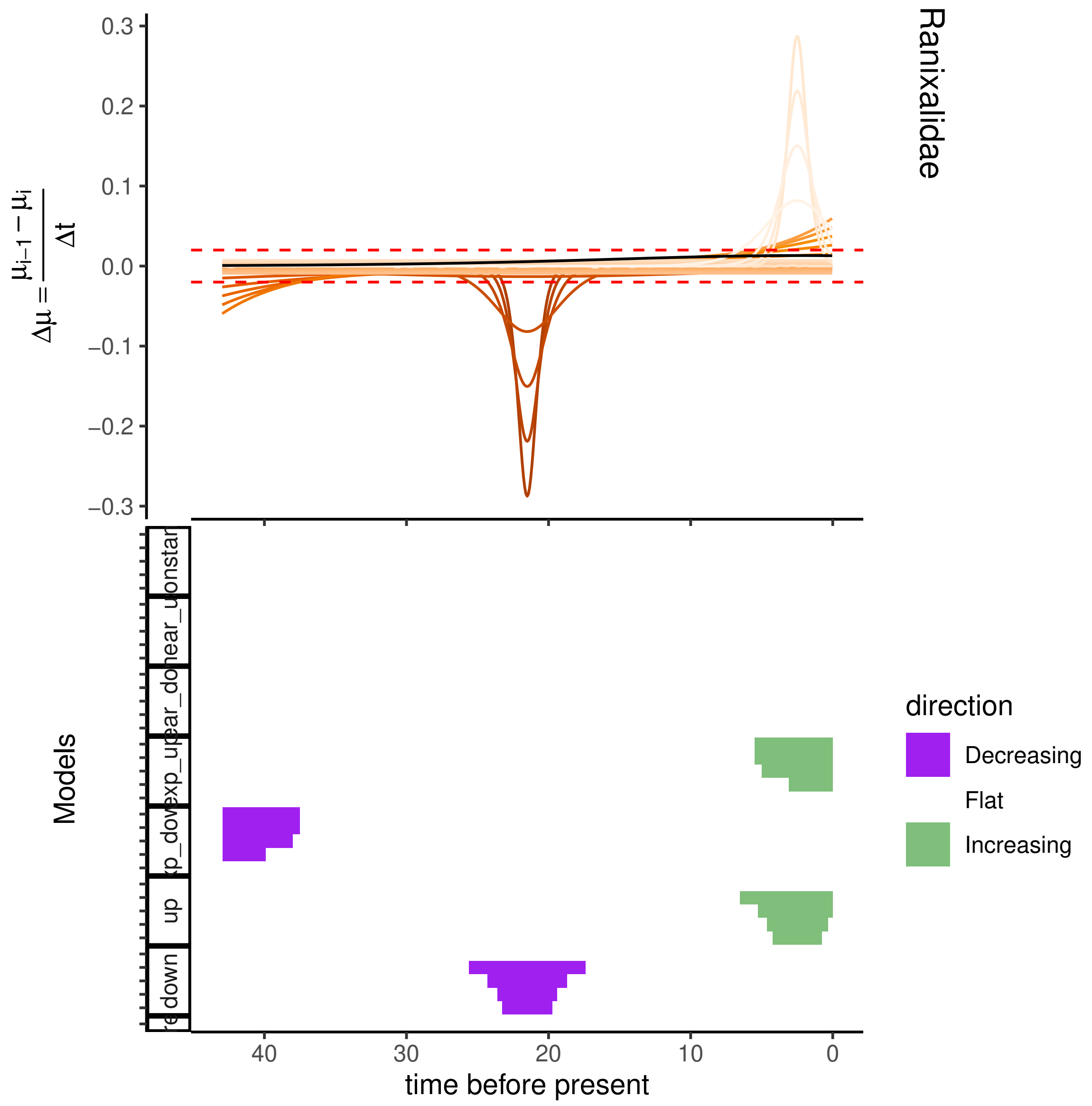
Ranixalidae
0.3
0.2
0.1
0.0
−0.1
−0.2
−0.3
$\Delta\mu = \frac{\mu_{i-1} - \mu_i}{\Delta t}$
Models
down
up
direction
Decreasing
Flat
Increasing
40
30
20
10
0
time before present

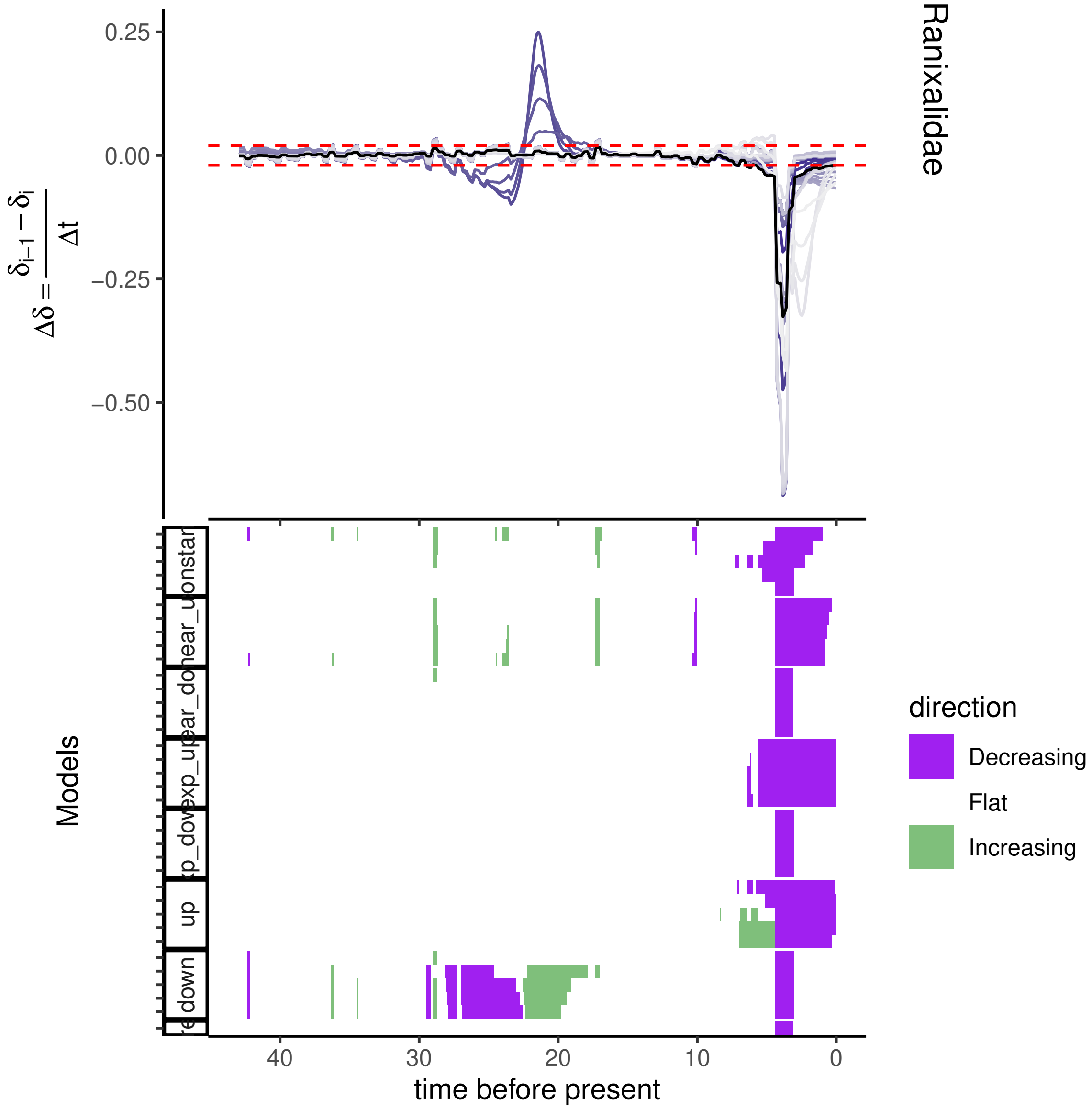

Ranixalidae
0.25
0.00
−0.25
−0.50
$\Delta\delta = \frac{\delta_{i-1} - \delta_i}{\Delta t}$
Models
40
30
20
10
0
time before present
direction
Decreasing
Flat
Increasing
down
up

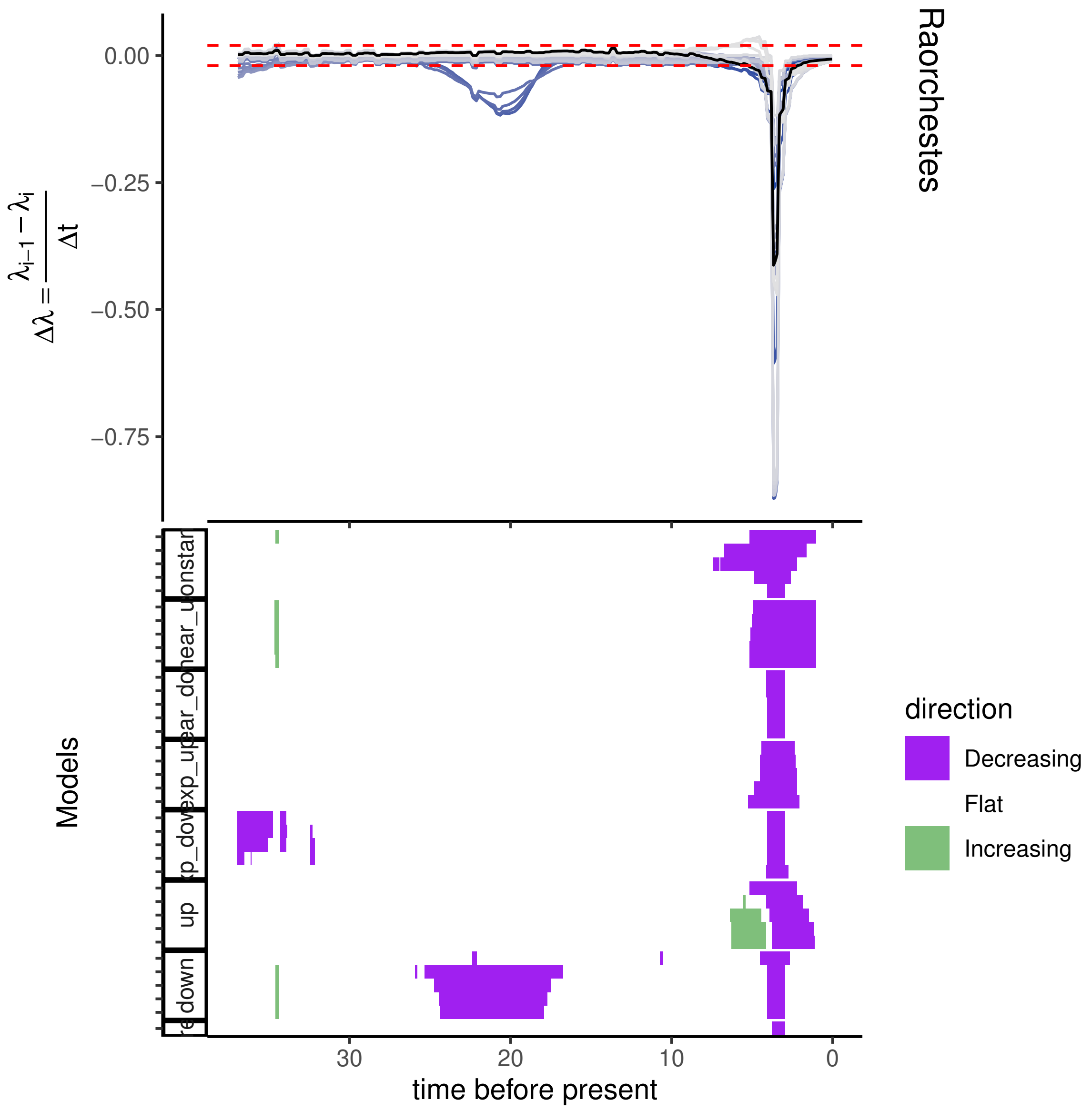

Raorchestes
$\Delta\lambda = \frac{\lambda_{i-1} - \lambda_i}{\Delta t}$
0.00
−0.25
−0.50
−0.75
Models
onstar
hear_u
uear_do
exp_upar
xp_dov
up
down
30
20
10
0
time before present
direction
Decreasing
Flat
Increasing

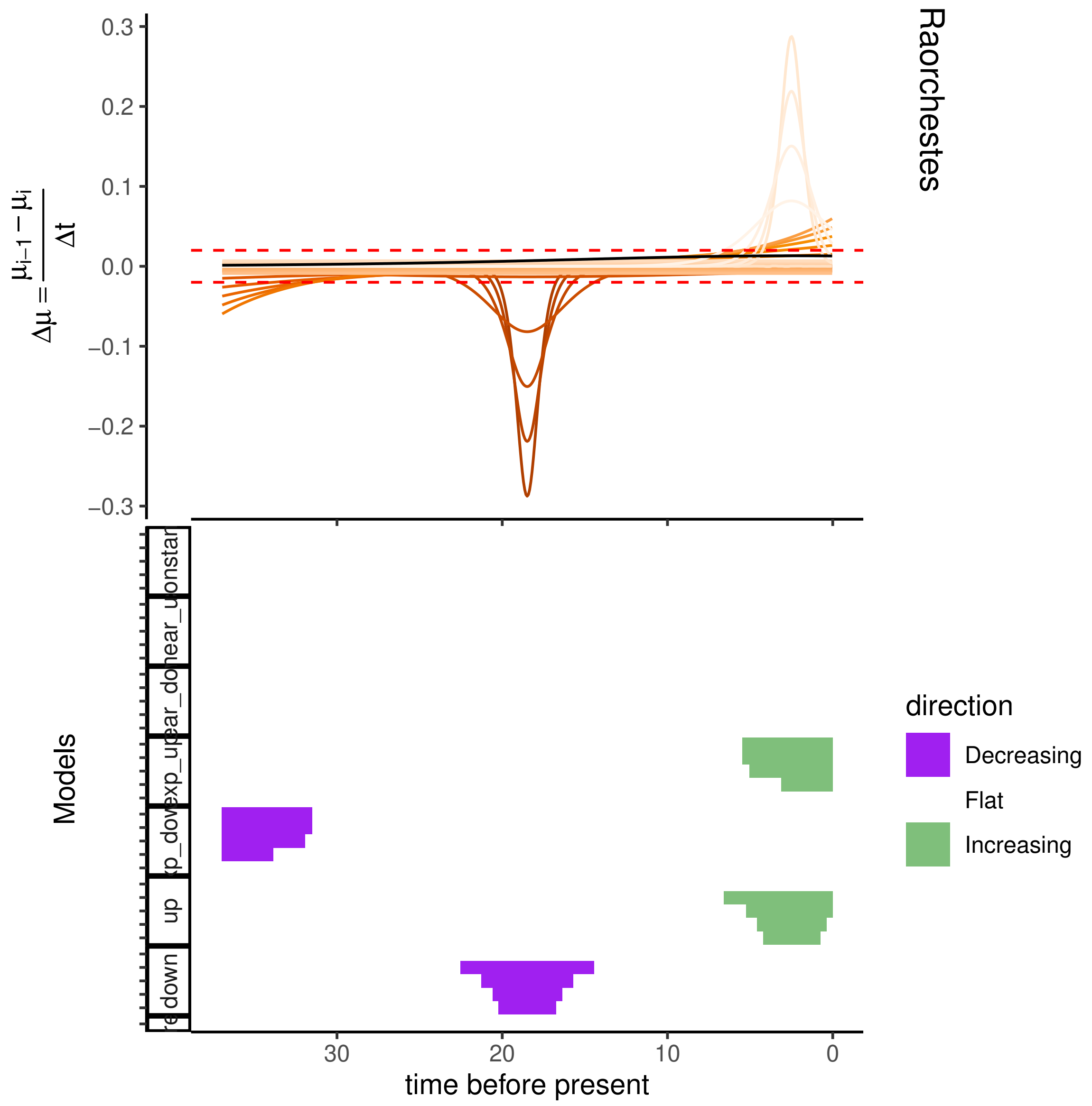
Raorchestes
0.3
0.2
0.1
0.0
−0.1
−0.2
−0.3
$\Delta\mu = \frac{\mu_{i-1} - \mu_i}{\Delta t}$
Models
onstar
near_u
ear_do
exp_up
xp_dov
up
down
30
20
10
0
time before present
direction
Decreasing
Flat
Increasing

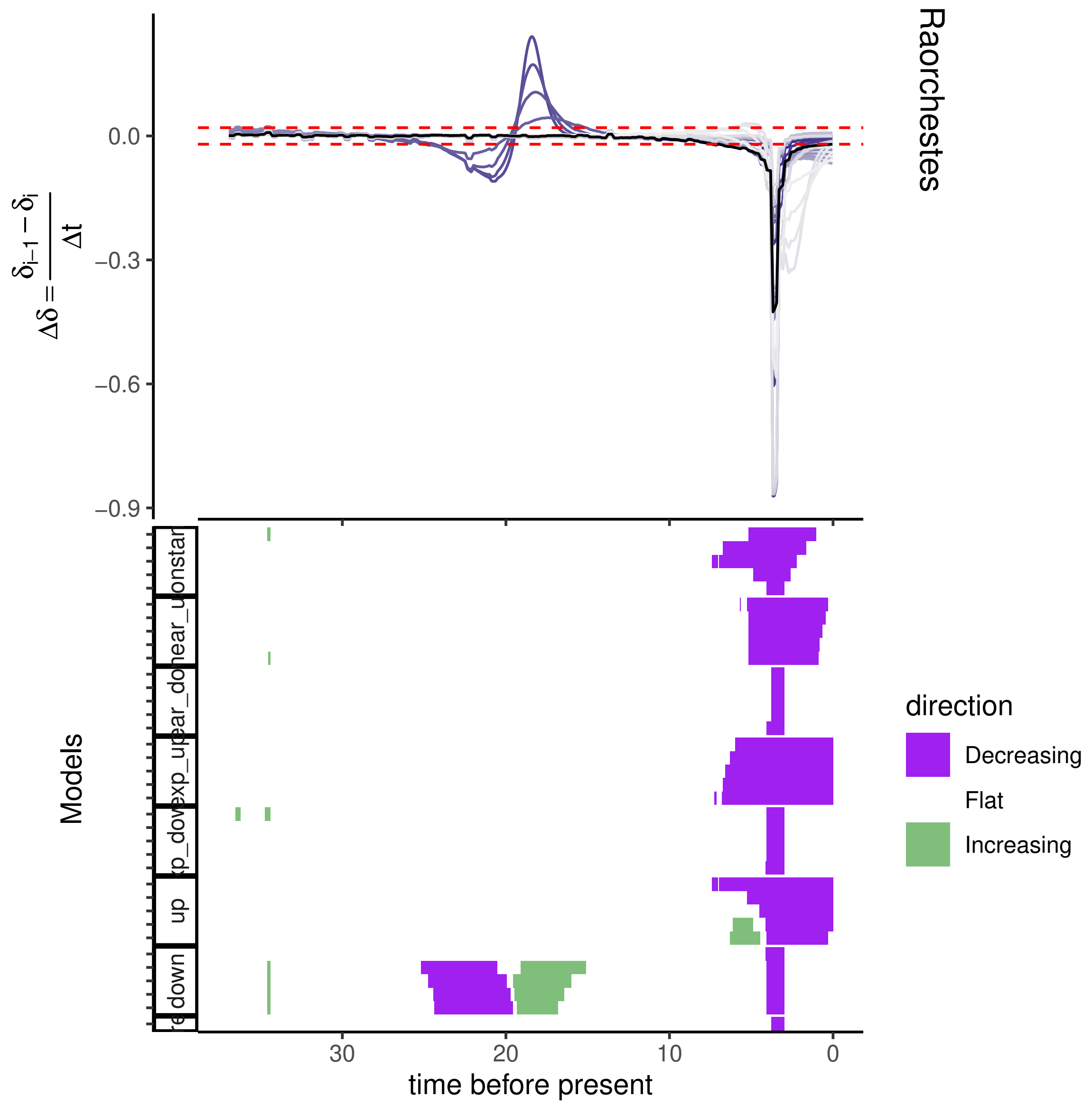

Raorchestes
$\Delta\delta = \frac{\delta_{i-1} - \delta_i}{\Delta t}$
0.0
−0.3
−0.6
−0.9
Models
time before present
30
20
10
0
direction
Decreasing
Flat
Increasing
up
down

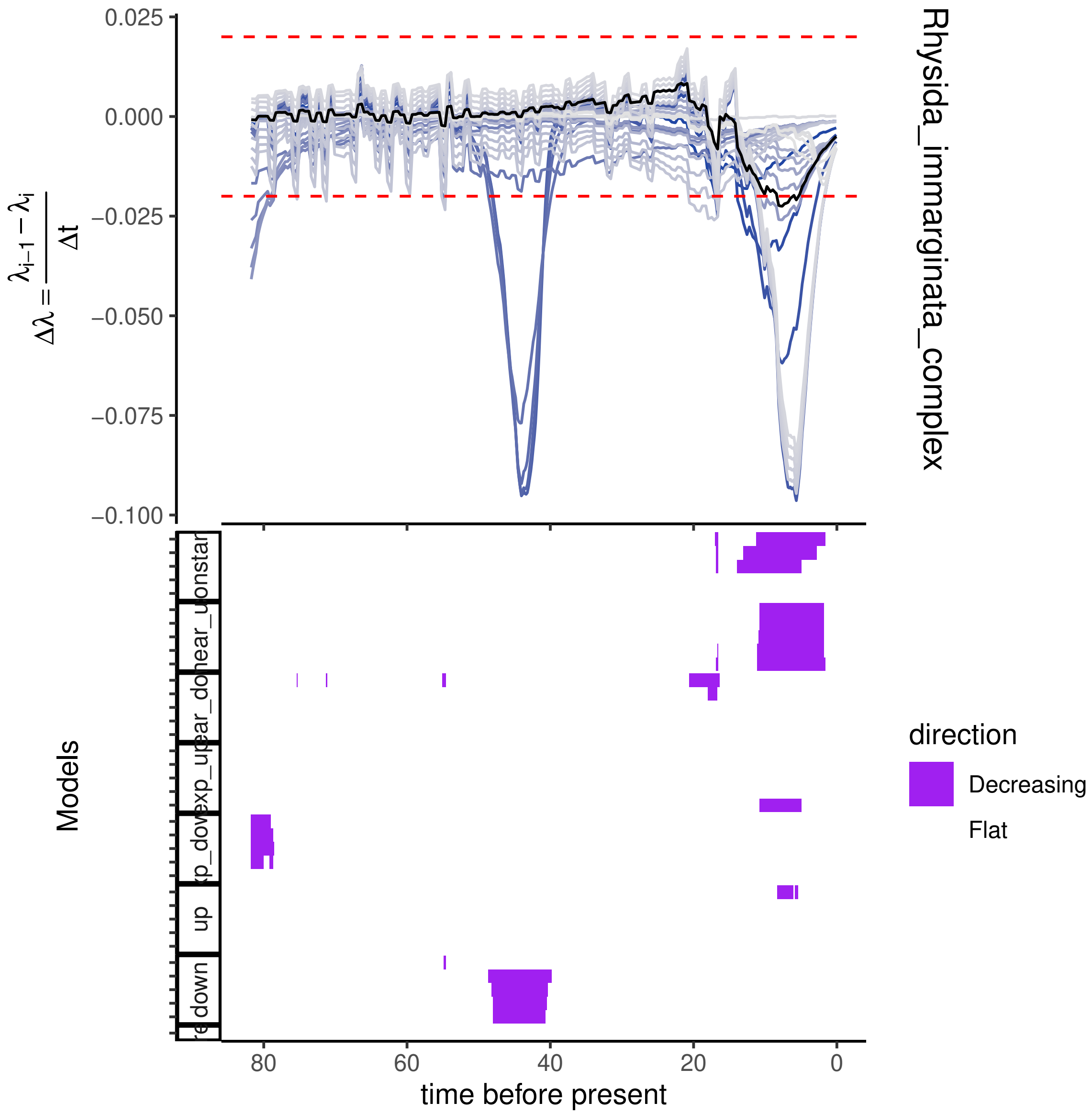
0.025
0.000
−0.025
−0.050
−0.075
−0.100
$\Delta\lambda = \frac{\lambda_{i-1} - \lambda_i}{\Delta t}$
Rhysida_immarginata_complex
Models
direction
Decreasing
Flat
80
60
40
20
0
time before present
up
down

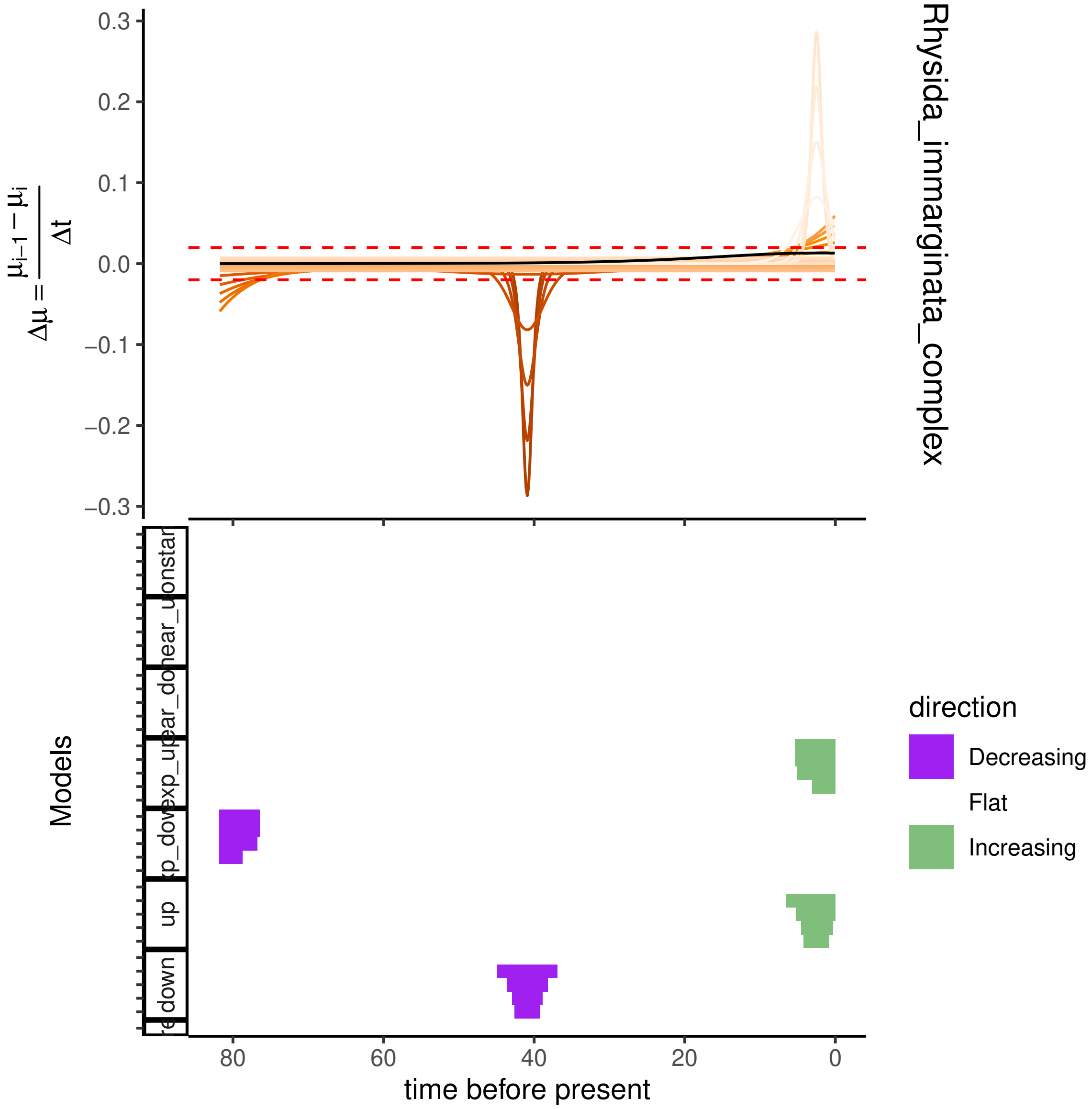
Rhysida_immarginata_complex
$\Delta\mu = \frac{\mu_{i-1} - \mu_i}{\Delta t}$
0.3
0.2
0.1
0.0
−0.1
−0.2
−0.3
Models
down
up
time before present
80
60
40
20
0
direction
Decreasing
Flat
Increasing

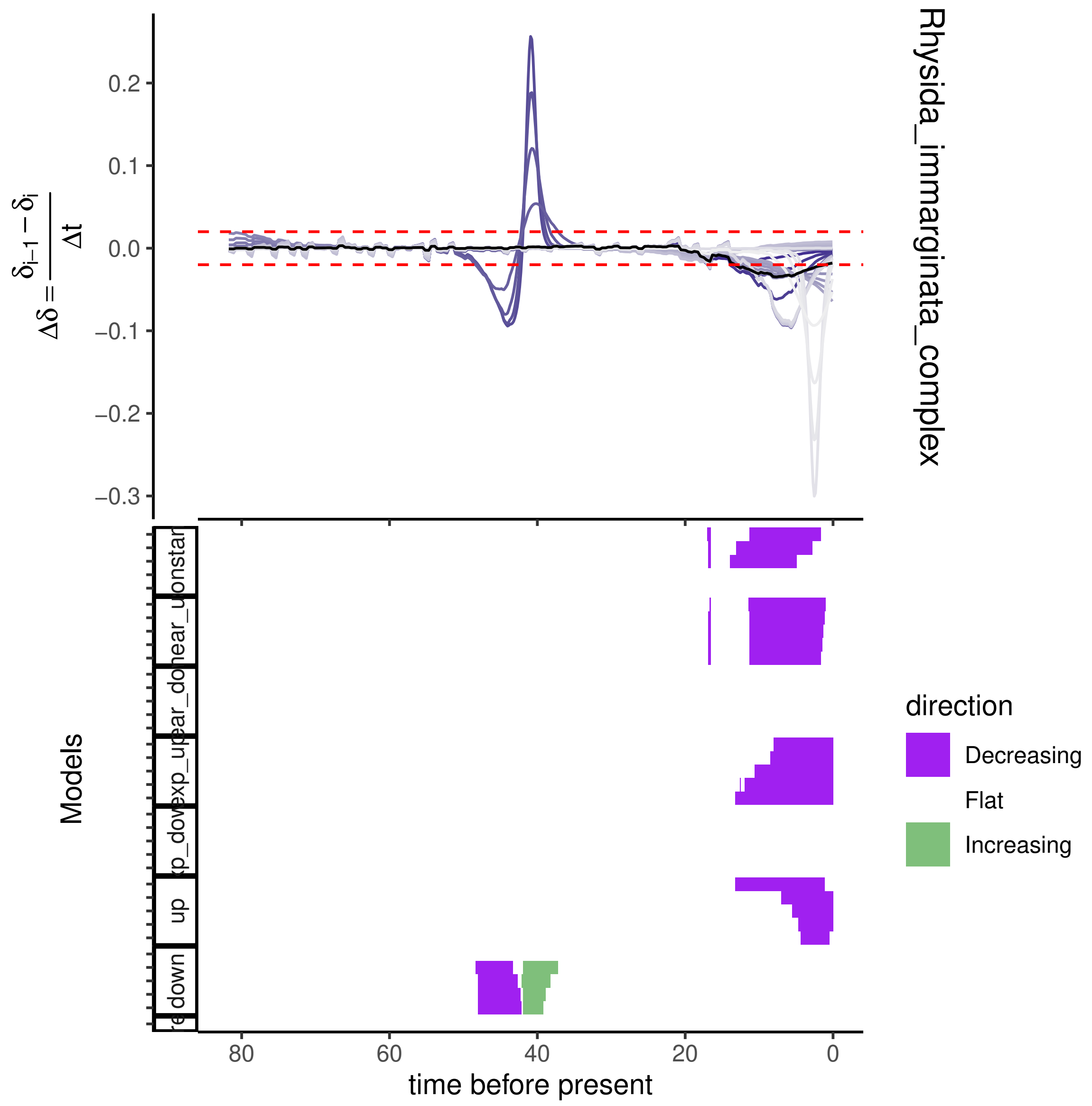

Rhysida_immarginata_complex
$\Delta\delta = \frac{\delta_{i-1} - \delta_i}{\Delta t}$
Models
time before present
direction
Decreasing
Flat
Increasing

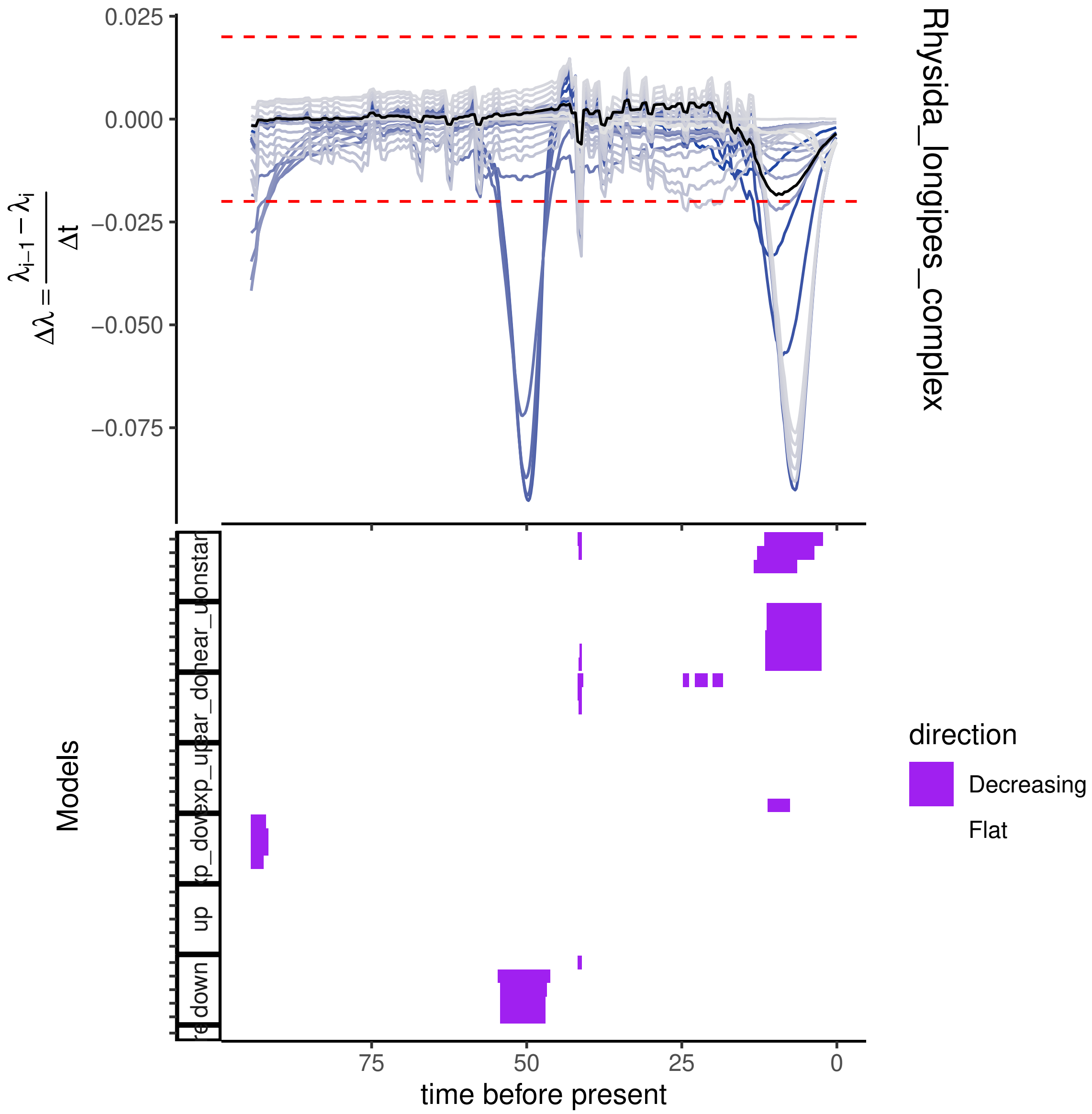

0.025
0.000
−0.025
−0.050
−0.075
$\Delta\lambda = \frac{\lambda_{i-1} - \lambda_i}{\Delta t}$
Rhysida_longipes_complex
Models
down
up
time before present
75
50
25
0
direction
Decreasing
Flat

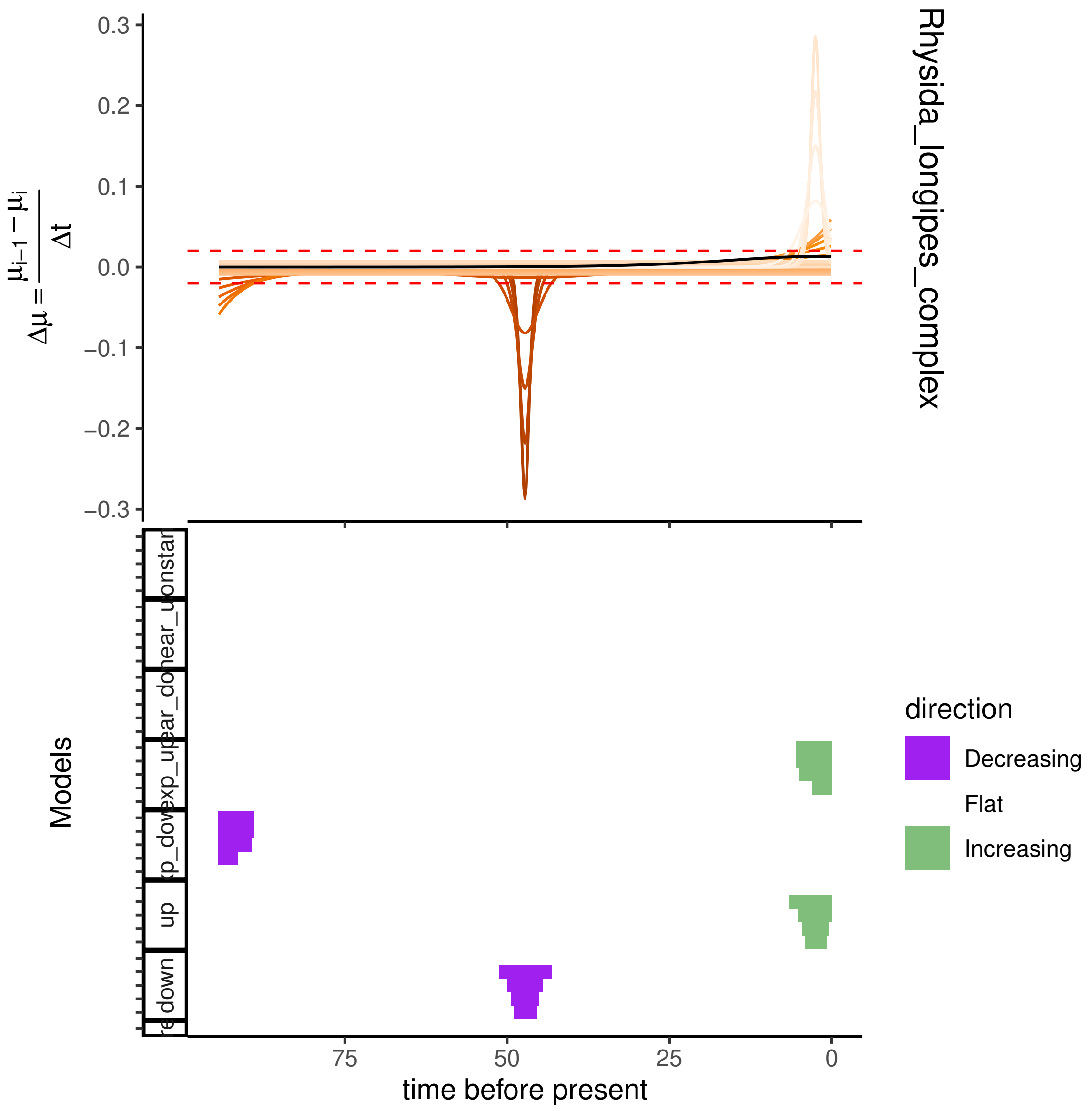

Rhysida_longipes_complex
$\Delta\mu = \frac{\mu_{i-1} - \mu_i}{\Delta t}$
0.3
0.2
0.1
0.0
−0.1
−0.2
−0.3
Models
onstan
near_u
ear_do
exp_up
xp_dow
up
down
75
50
25
0
time before present
direction
Decreasing
Flat
Increasing

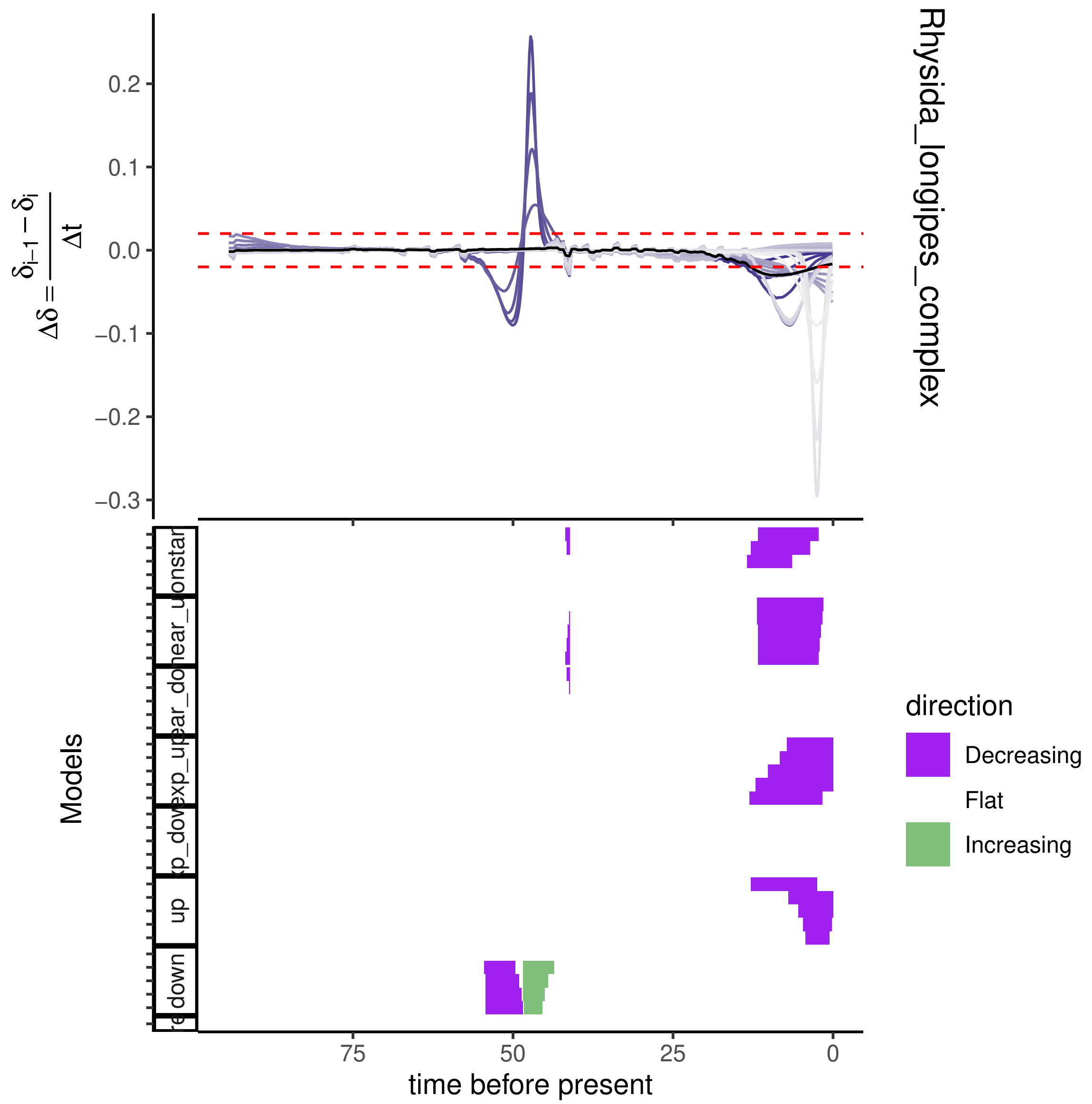

Rhysida_longipes_complex
0.2
0.1
0.0
−0.1
−0.2
−0.3
$\Delta\delta = \frac{\delta_{i-1} - \delta_i}{\Delta t}$
Models
up
down
75
50
25
0
time before present
direction
Decreasing
Flat
Increasing

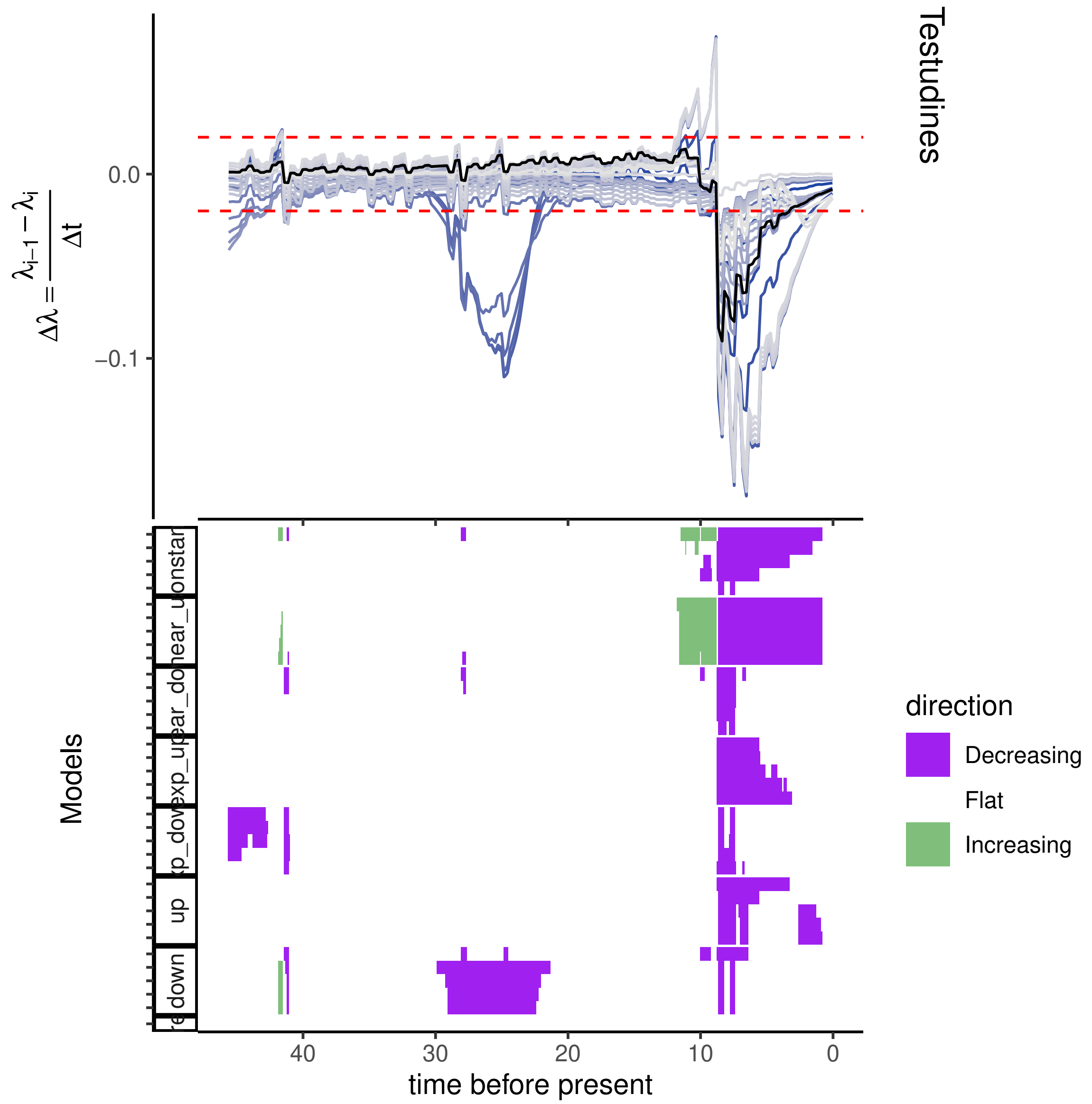
Testudines
0.0
−0.1
$\Delta\lambda = \frac{\lambda_{i-1} - \lambda_i}{\Delta t}$
Models
40
30
20
10
0
time before present
direction
Decreasing
Flat
Increasing
down
up

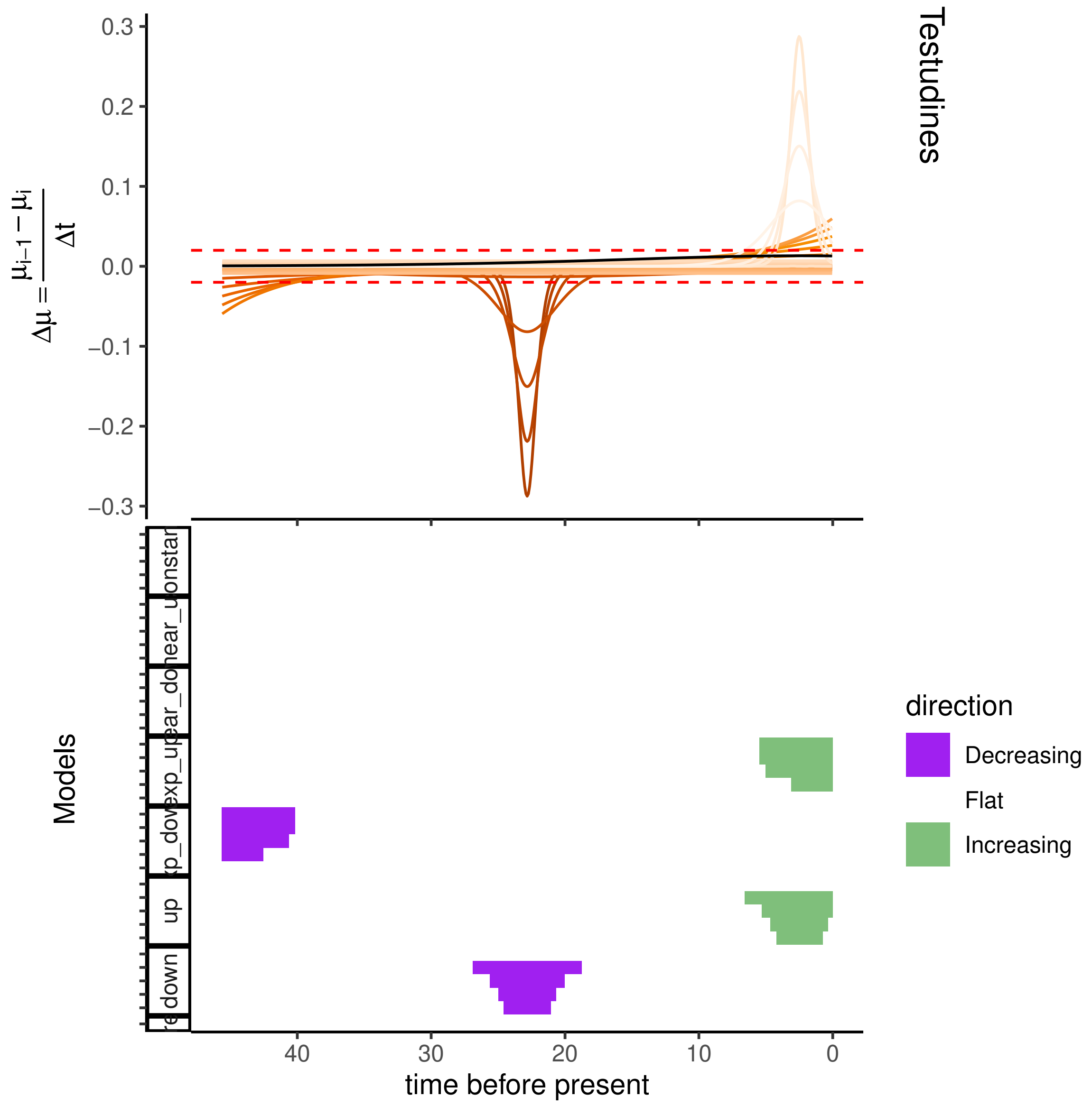
Testudines
$\Delta\mu = \frac{\mu_{i-1} - \mu_i}{\Delta t}$
0.3
0.2
0.1
0.0
−0.1
−0.2
−0.3
Models
direction
Decreasing
Flat
Increasing
40
30
20
10
0
time before present
up
down

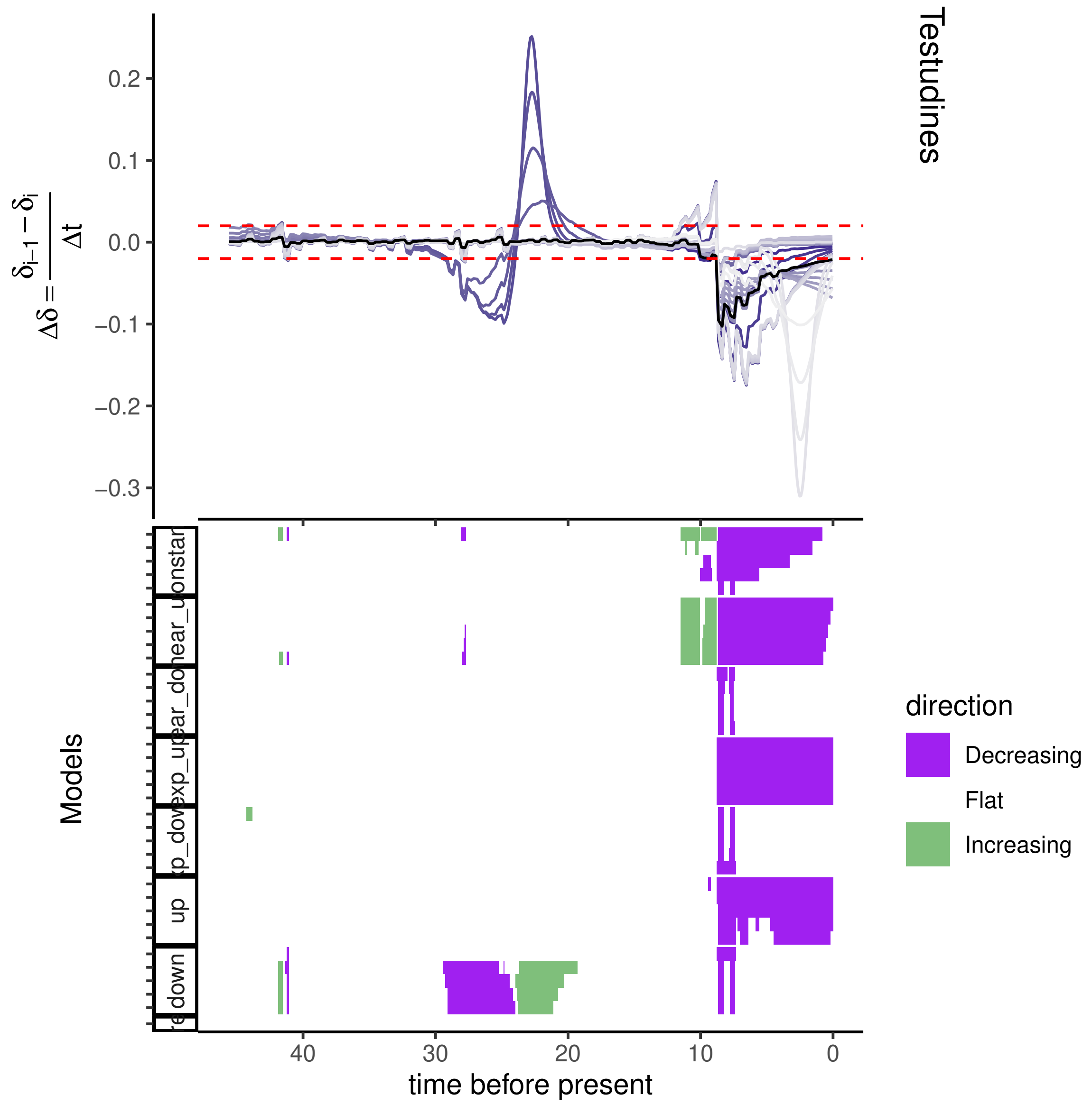

Testudines
0.2
0.1
0.0
−0.1
−0.2
−0.3
$\Delta\delta = \frac{\delta_{i-1} - \delta_i}{\Delta t}$
Models
direction
Decreasing
Flat
Increasing
40
30
20
10
0
time before present
up
down

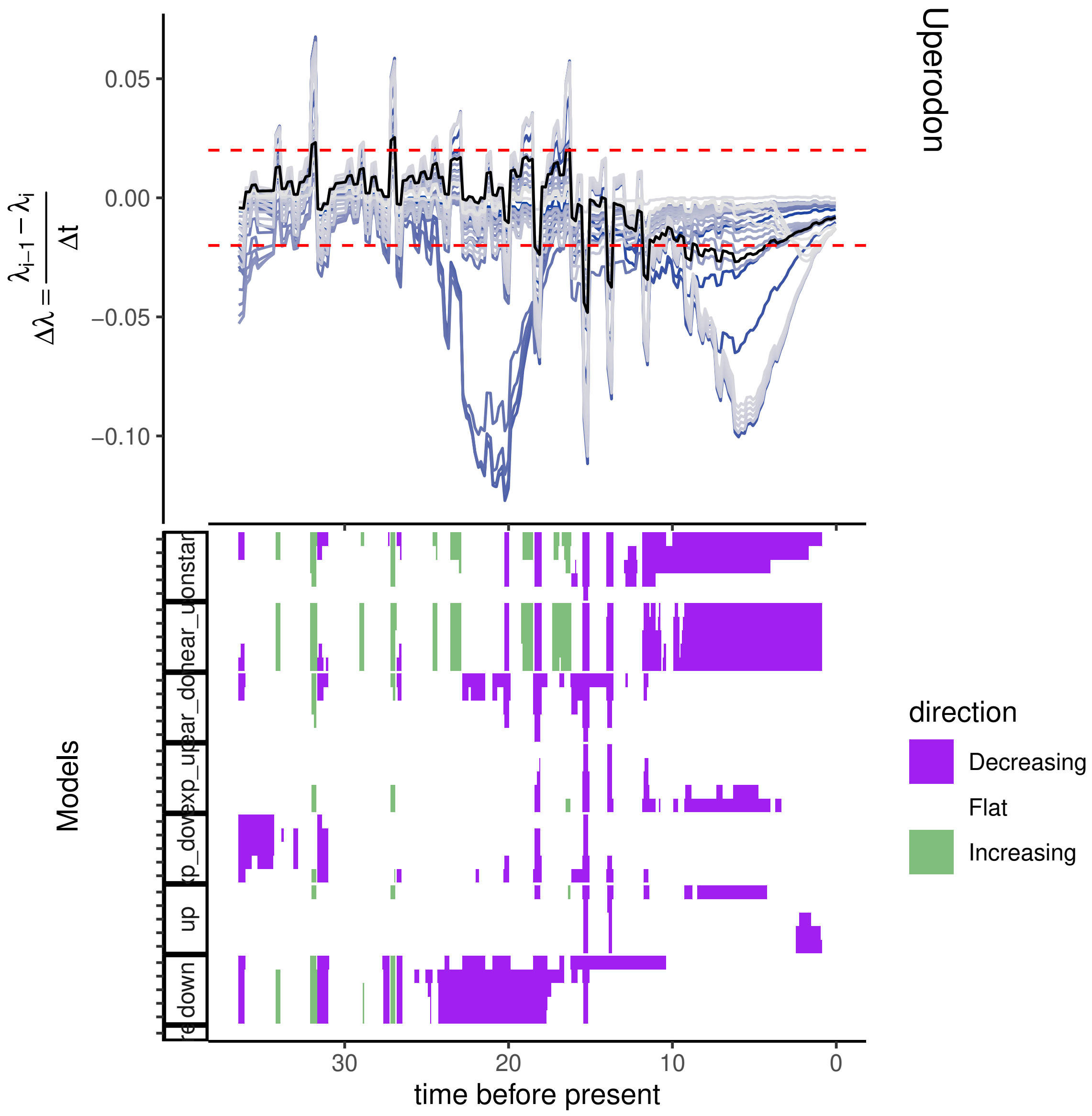
Uperodon
$\Delta\lambda = \frac{\lambda_{i-1} - \lambda_i}{\Delta t}$
0.05
0.00
−0.05
−0.10
Models
direction
Decreasing
Flat
Increasing
30
20
10
0
time before present

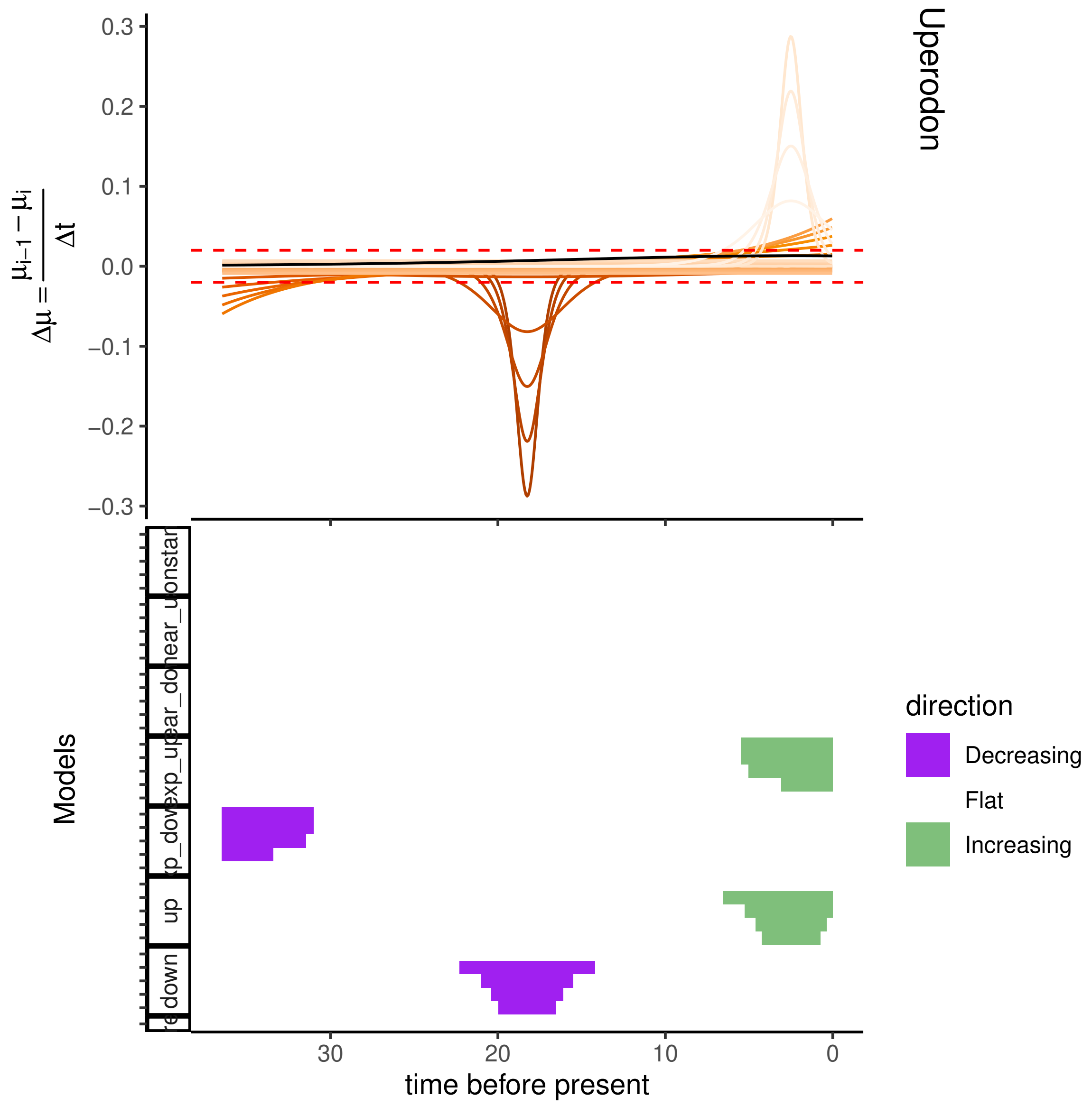

0.3
0.2
0.1
0.0
−0.1
−0.2
−0.3
$\Delta\mu = \frac{\mu_{i-1} - \mu_i}{\Delta t}$
Uperodon
Models
direction
Decreasing
Flat
Increasing
up
down
30
20
10
0
time before present

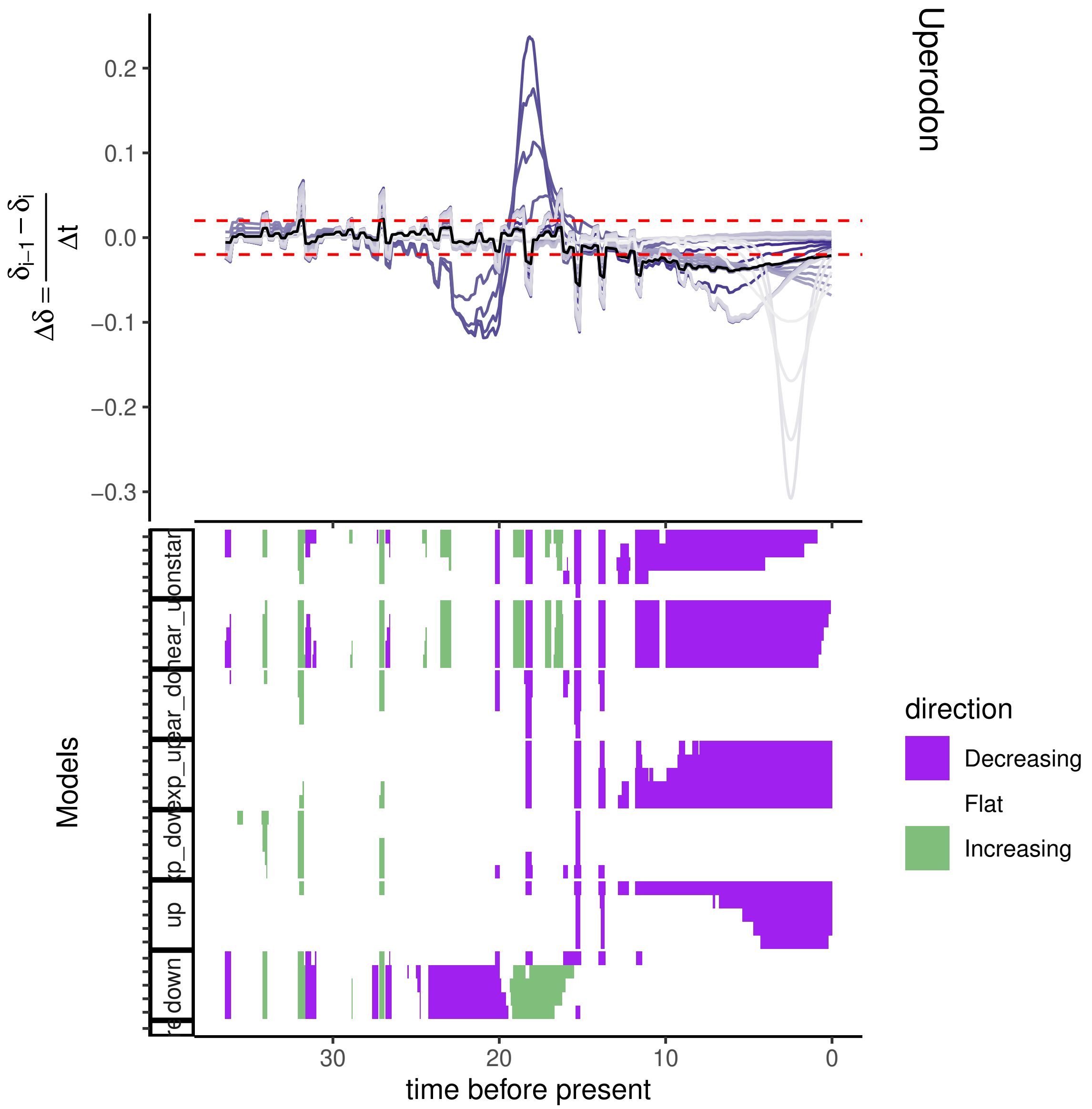
Uperodon
0.2
0.1
0.0
−0.1
−0.2
−0.3
$\Delta\delta = \frac{\delta_{i-1} - \delta_i}{\Delta t}$
Models
up
down
direction
Decreasing
Flat
Increasing
30
20
10
0
time before present

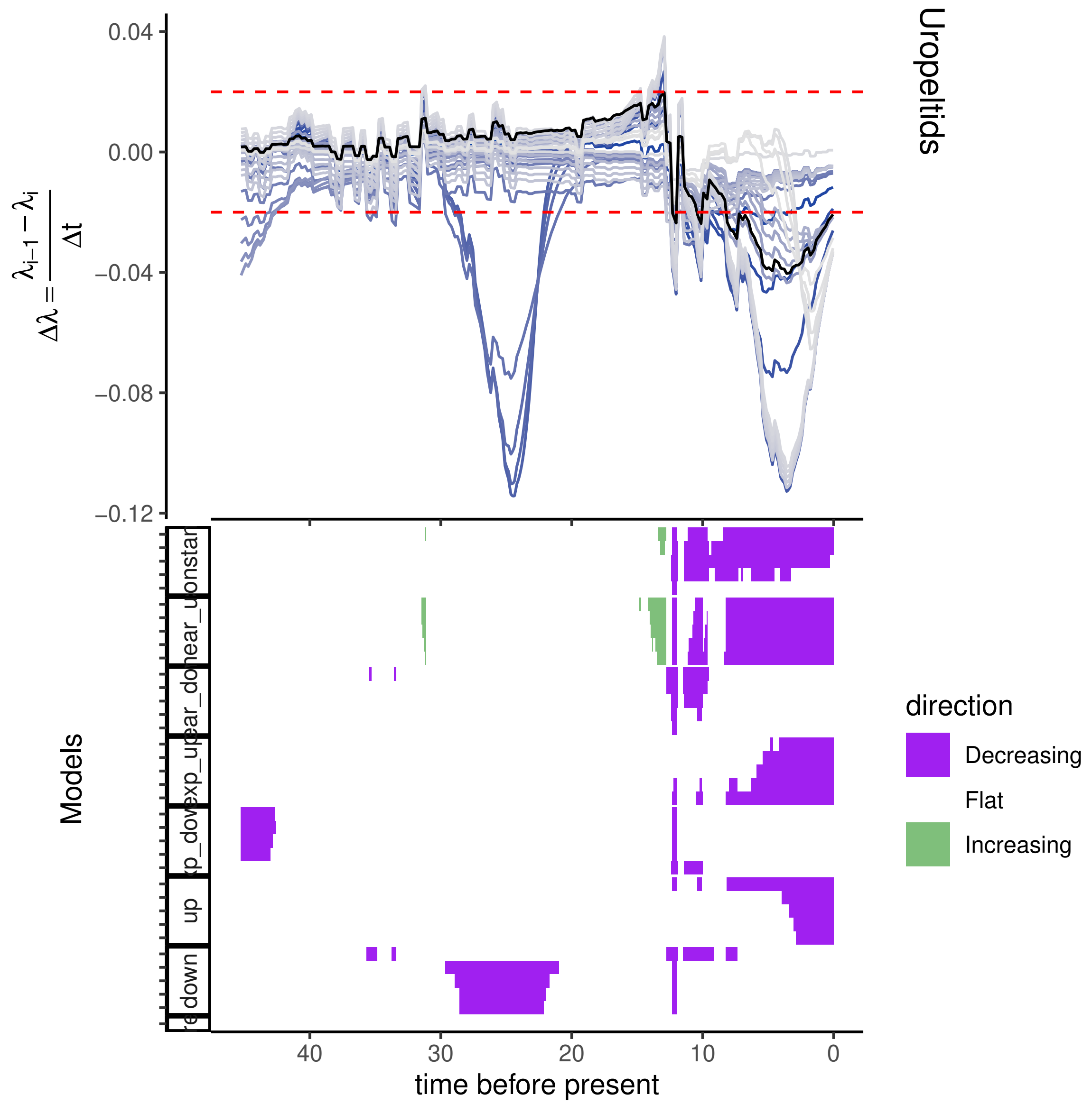
Uropeltids
0.04
0.00
−0.04
−0.08
−0.12
$\Delta\lambda = \frac{\lambda_{i-1} - \lambda_i}{\Delta t}$
Models
up
down
40
30
20
10
0
time before present
direction
Decreasing
Flat
Increasing

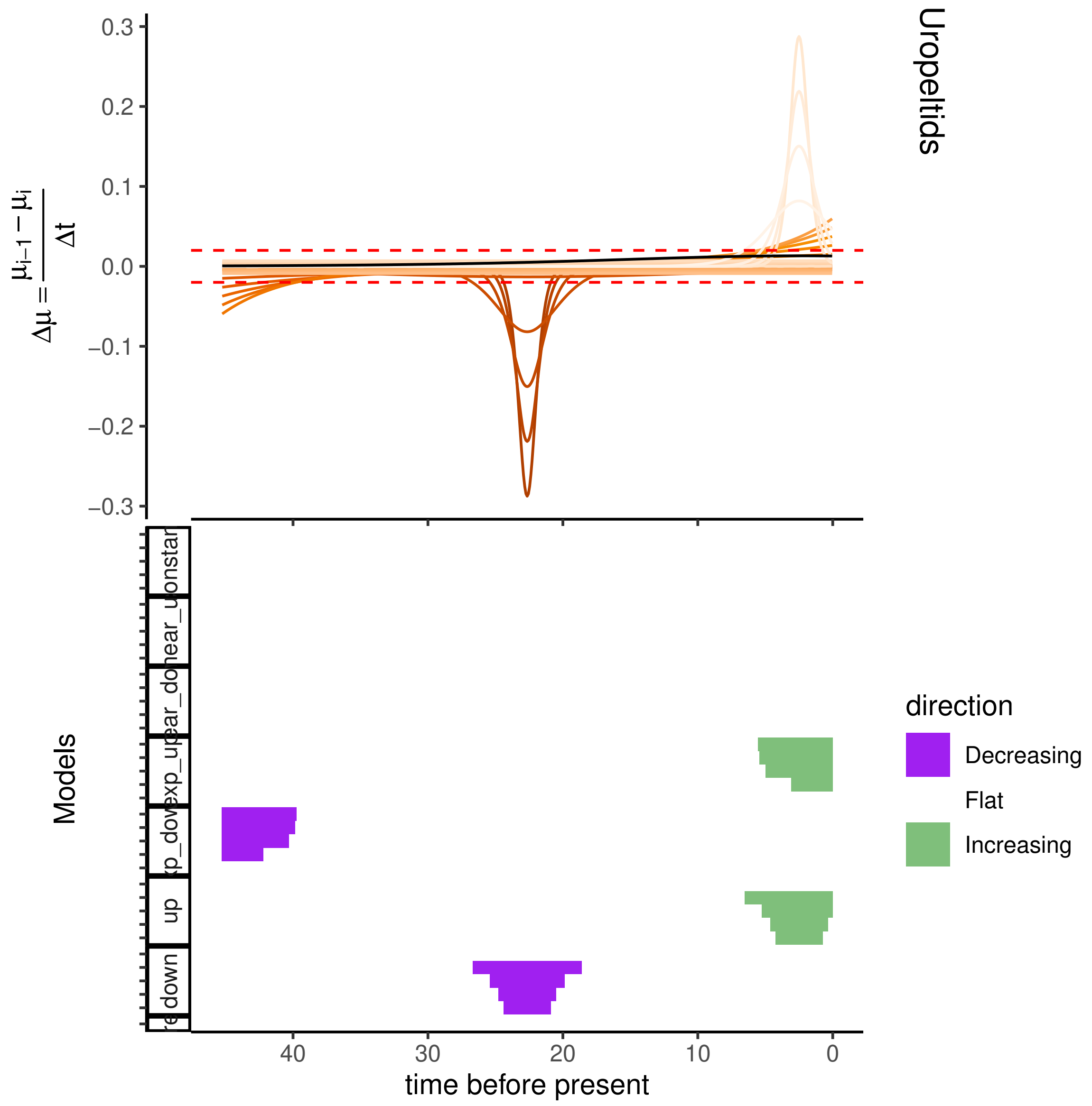
Uropeltids
0.3
0.2
0.1
0.0
−0.1
−0.2
−0.3
Models
up
down
40
30
20
10
0
time before present
direction
Decreasing
Flat
Increasing

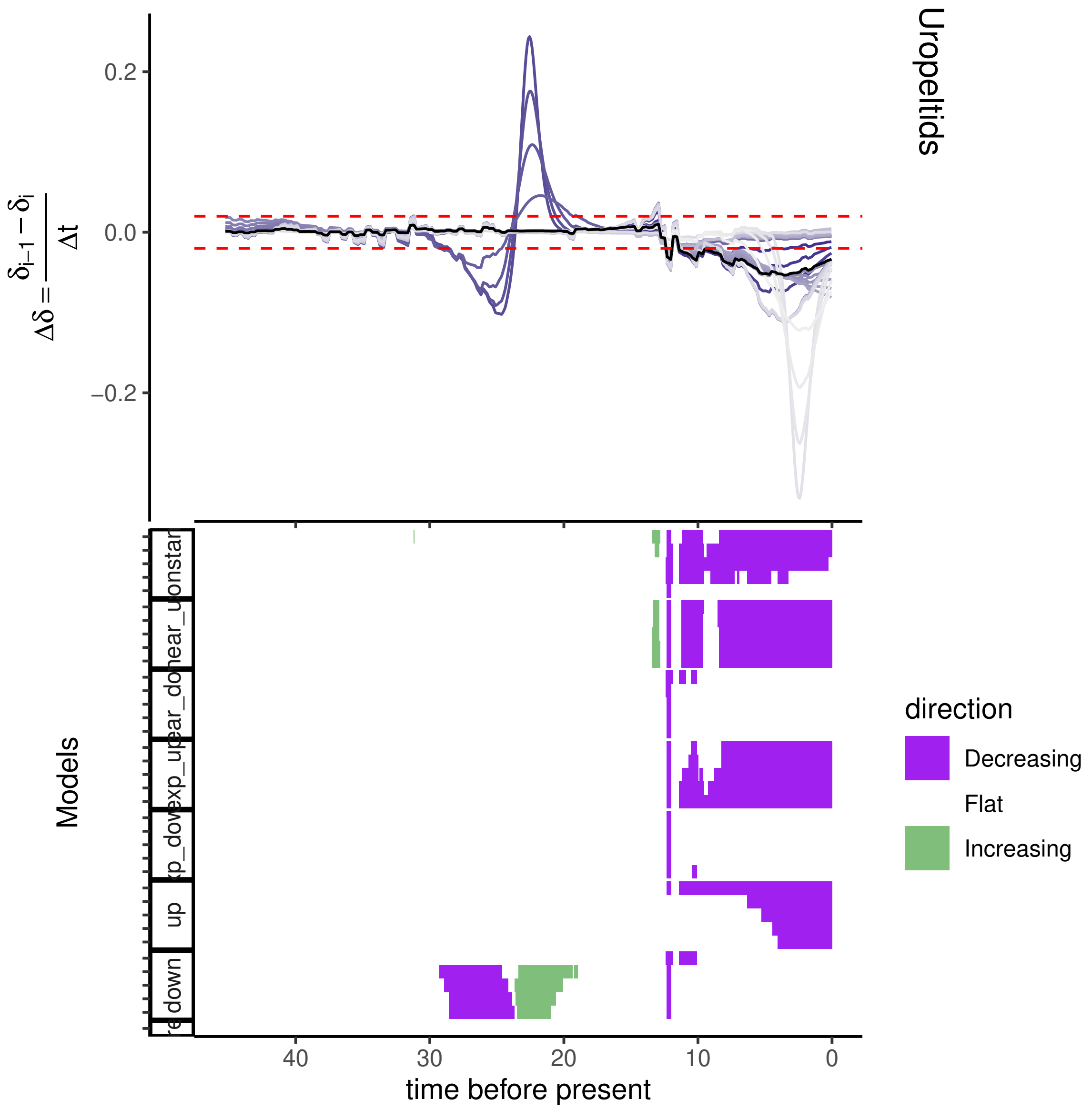
Uropeltids
0.2
0.0
−0.2
$\Delta\delta = \frac{\delta_{i-1} - \delta_i}{\Delta t}$
Models
direction
Decreasing
Flat
Increasing
down
up
40
30
20
10
0
time before present

**Figure S5: The priors (page 28) and results of the CRABS analysis (pages 29 - 124)** [25]. The different prior functions and values were used for extinction rates to obtain the corresponding pulled speciation rates and vice-versa that constitute the congruence class and this further helped generate the pulled diversification rates. Functions include constant rates and time-varying rates (linearly, exponentially and sigmoidally). Within pages 29 - 124, each consecutive triplet of pages correspond to one lineage, the name of which is written on the top right. Illustrations of the dynamics in the different congruent speciation ($\lambda$), extinction ($\mu$) and diversification ($\delta$) models have been provided. The upper panel plots the change in the slopes of each model and the lower panel summarises the trends in the change of slope (i.e. flat or constant, increasing and decreasing) for each time bin.

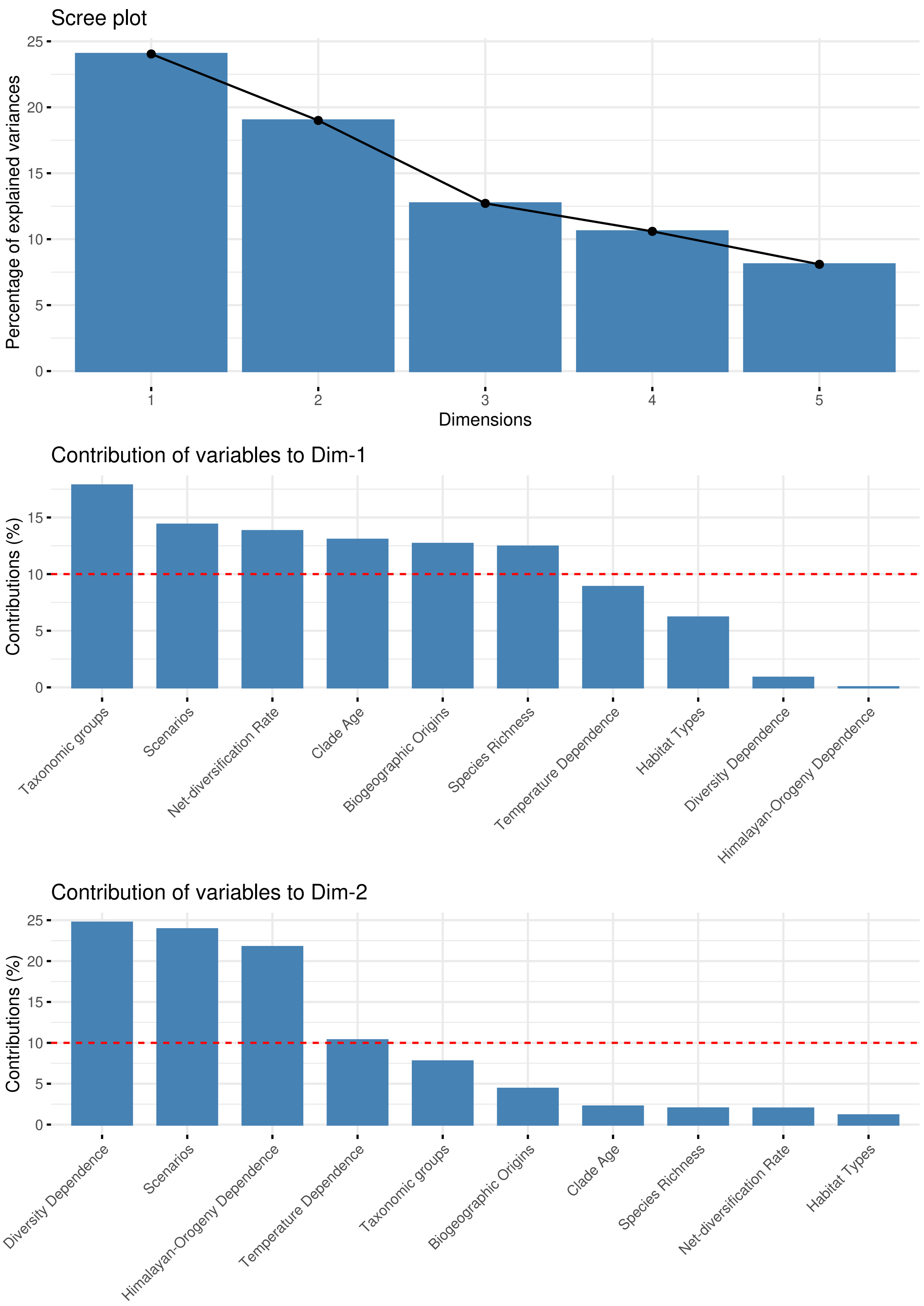

**(a)** Scree plot and dimension contributions for FAMD

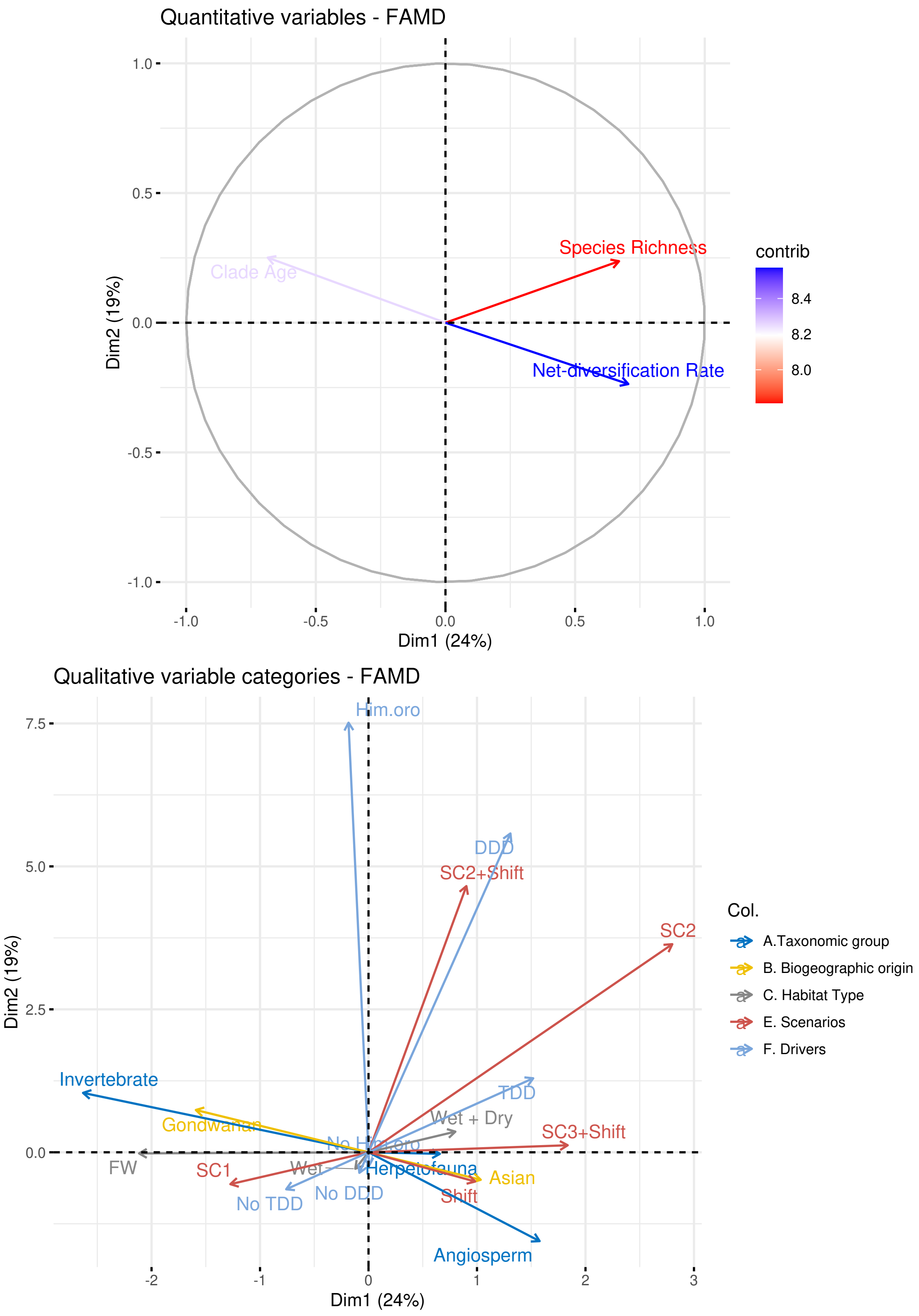

**(b)** Quantitative and qualitative variables used in FAMD

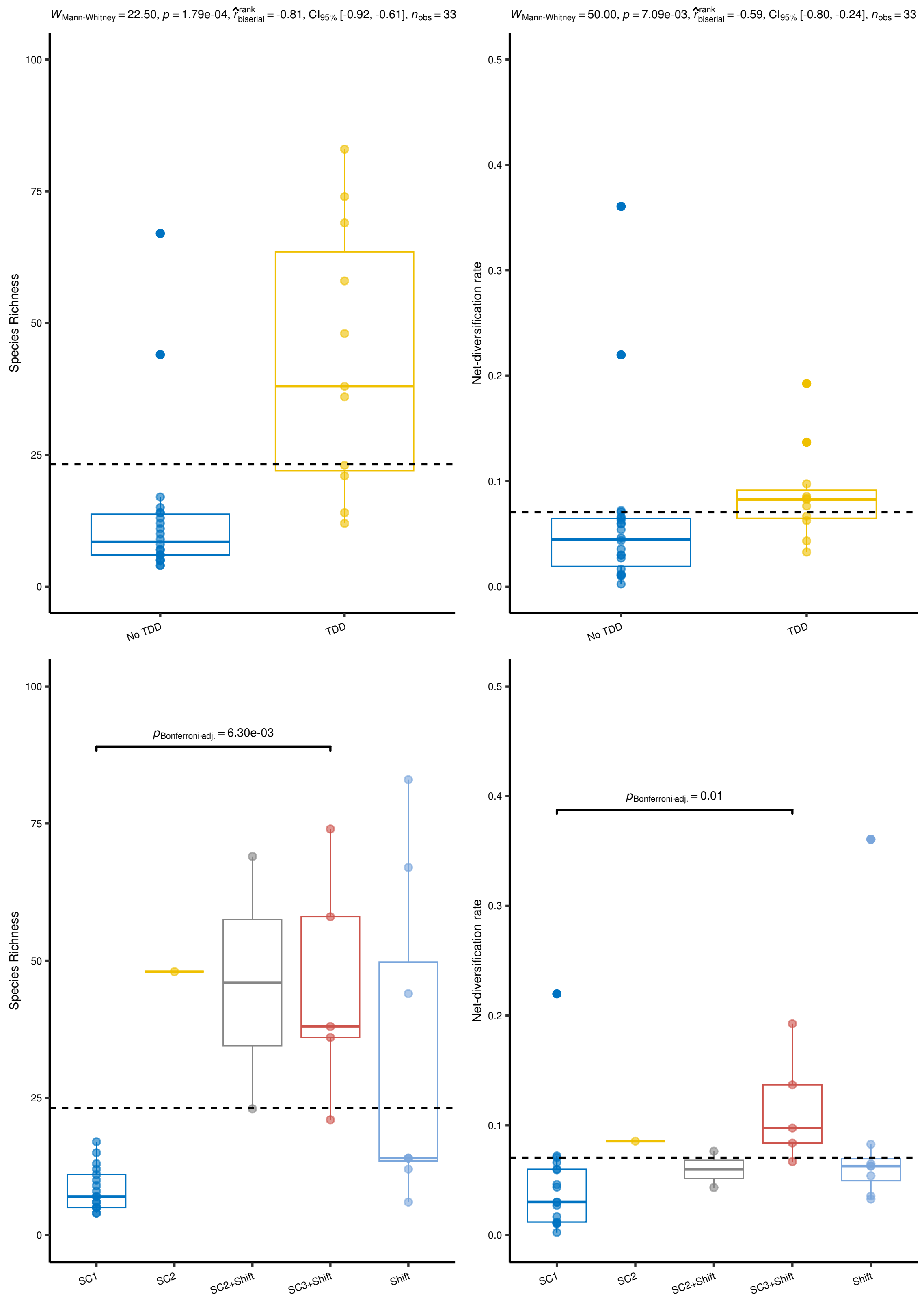

**(c)** Boxplots comparing species richness and net-diversification rates (ClaDS) across scenarios and drivers

**Figure S6: Results from the Factor Analysis of Mixed Data** [26, 27]

**(a) (page 126) (upper panel)** Scree plot indicates the percentages of total variance in the dataset that each ordination dimension explains, **(middle and bottom panel)** variables used in the analysis and their individual contributions to dimension-1 and dimension-2 respectively. Red dotted line indicates the average contribution by each variable to the corresponding ordination dimension.

**(b) (page 127)** Quantitative and Qualitative variables on the reduced two-dimensional ordination space. (upper panel) Three quantitative variables – species richness, net-diversification rates and clade ages represented by arrows. Colours indicate the contribution (percentage) of each variable to the total variance of the dataset (legend provided in figure). Species richness seems to be more strongly correlated to net-diversification rate than to clade age and as has been shown in earlier figures, clade age and net-diversification rates seem to be inversely correlated. (lower panel) Positions of different categorical(qualitative) variables in the ordination space indicating their correlations in with each other.

**(c) (page 128)** The FAMD analysis indicates high congruence in the clusters for temperature-dependence in diversification and for different diversification scenarios. And, temperature-dependence and scenarios have high contribution to Dimension 1 of the ordination which also is substantially contributed by net-diversification rates and species richness. Hence, we expected to observe some structure in species richness and diversification rates across different scenarios and groups of lineages showing varying support for temperature-dependence. **(upper panel)** Boxplots comparing species richness and net-diversifcation rates (Mann-Whitney U test) of lineages across lineages that show temperature-dependent diversification and the ones that do not. Lineages showing TDD seem to show significantly higher species richness and net-diversification rates than lineages that do not (p-values have been provided at the top the respective plots). **(bottom panel)** Boxplots comparing species richness and net-diversification rates across different diversification scenarios. Lineages following gradual accumulation (SC1) seem to show lower species richness and net-diversification rates than the ones following other diversification scenarios (tested using the Dunn's test following Kruskal-Wallis test). Significant comparisons have been highlighted and their corresponding corrected p-values have been provided.

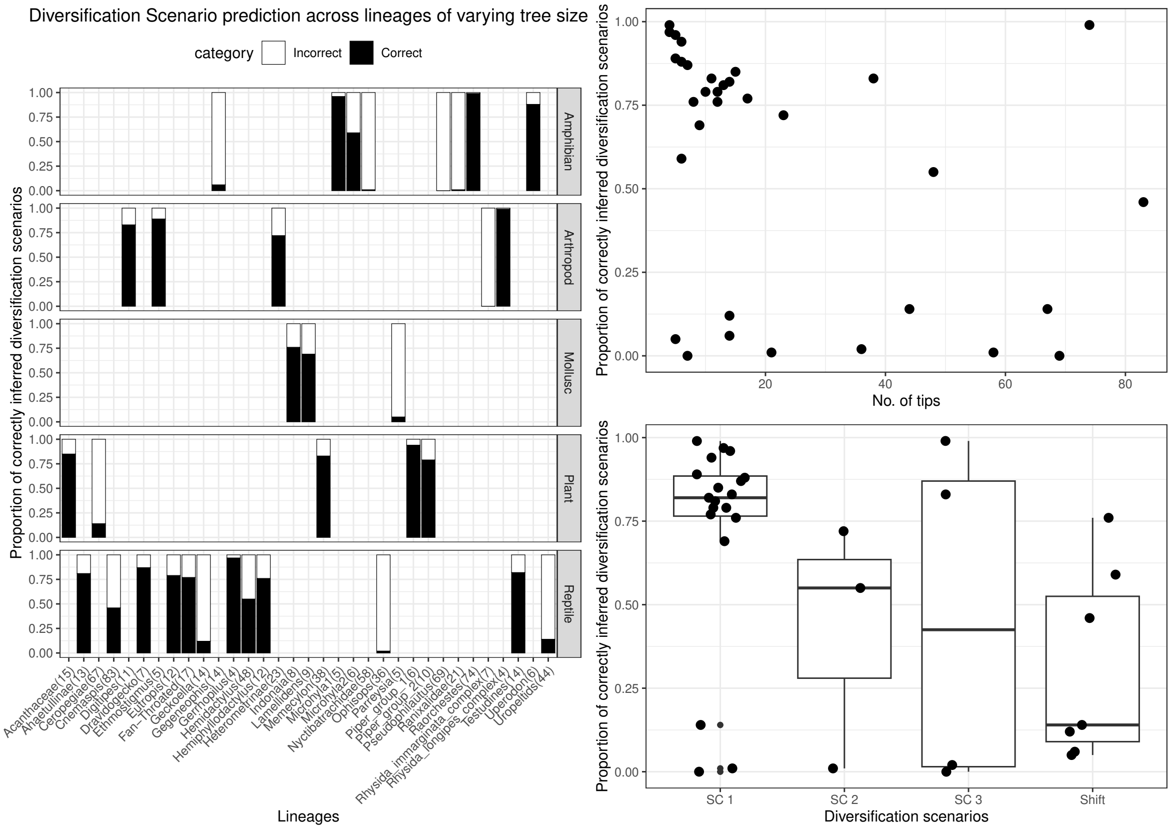

**(a)** Diversification scenario estimation of simulated trees by RPANDA.

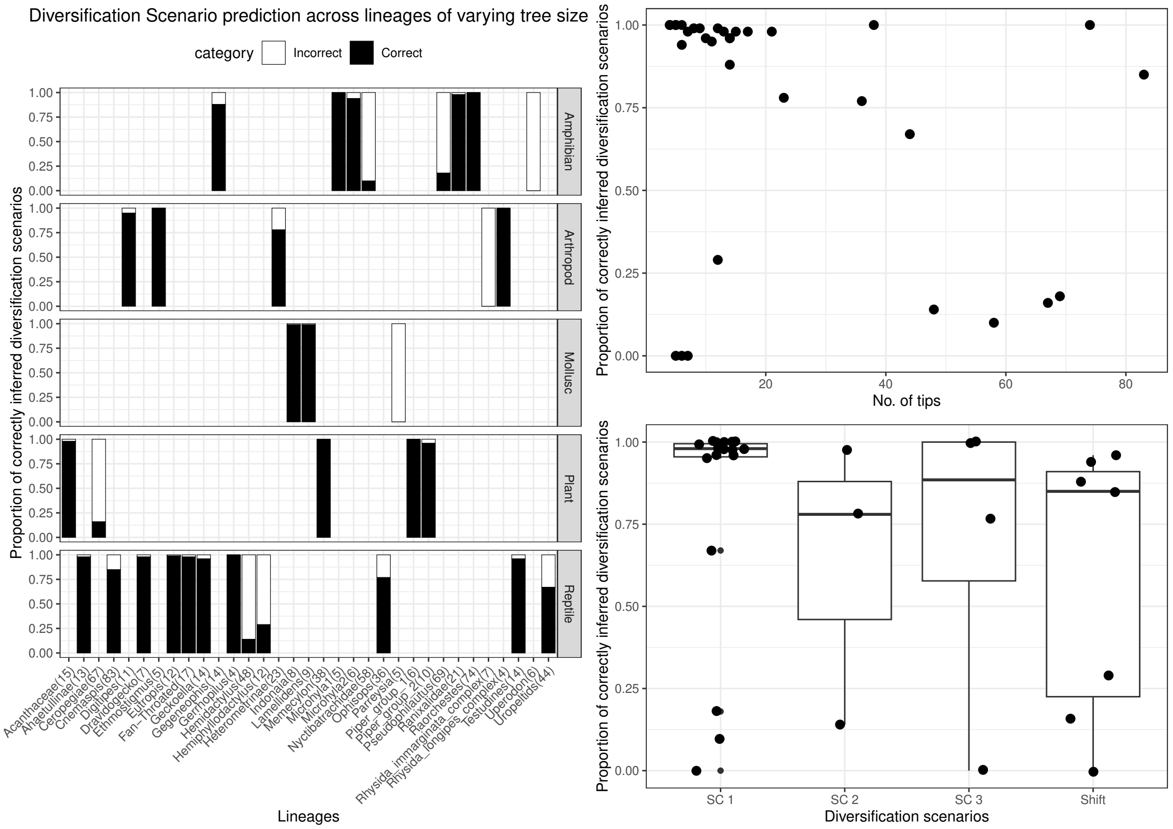

**(b)** Diversification scenario estimation of simulated trees by CoMET.

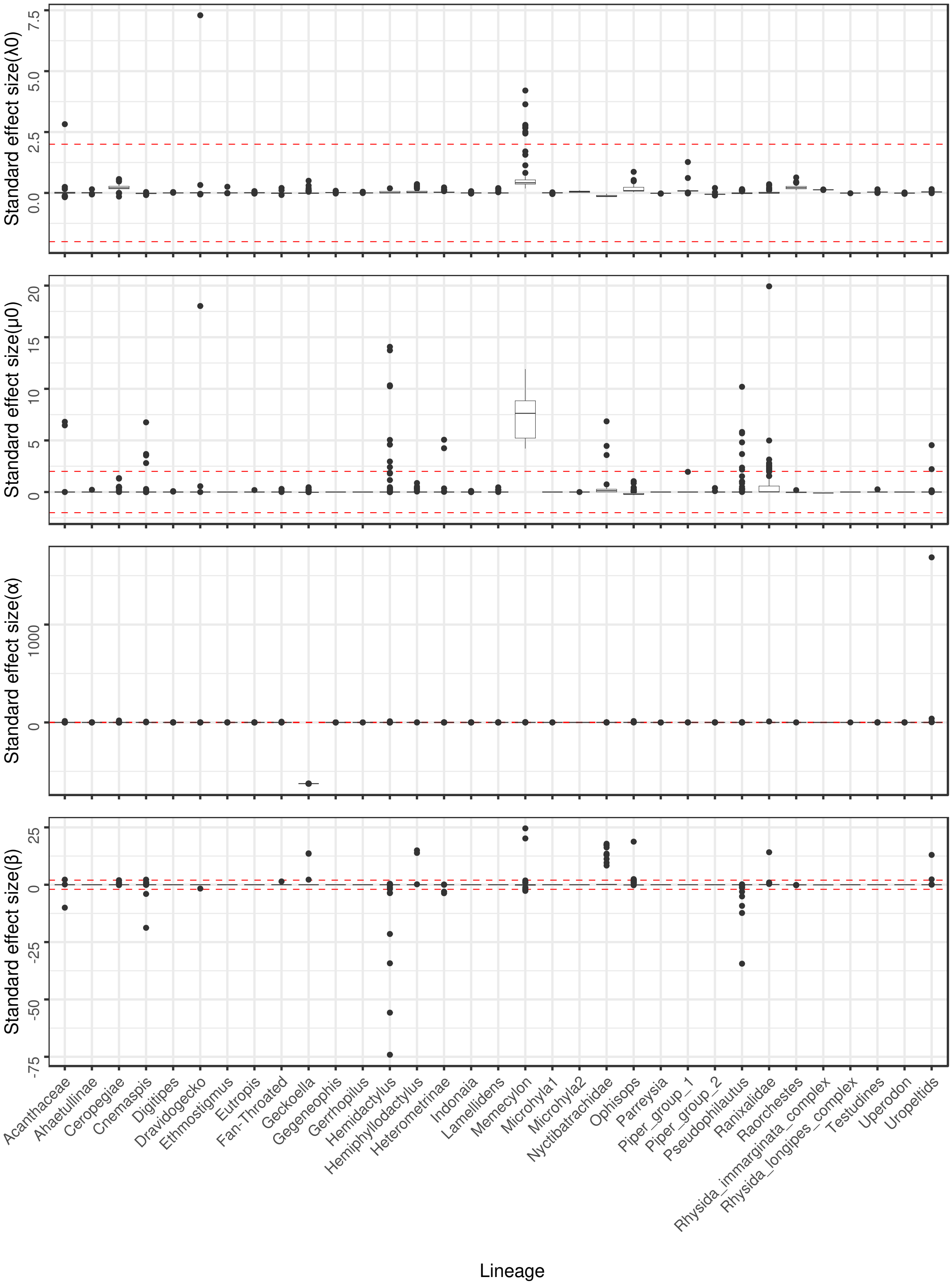

**(c)** Best-fitting birth-death model parameters of simulated trees estimated by RPANDA.

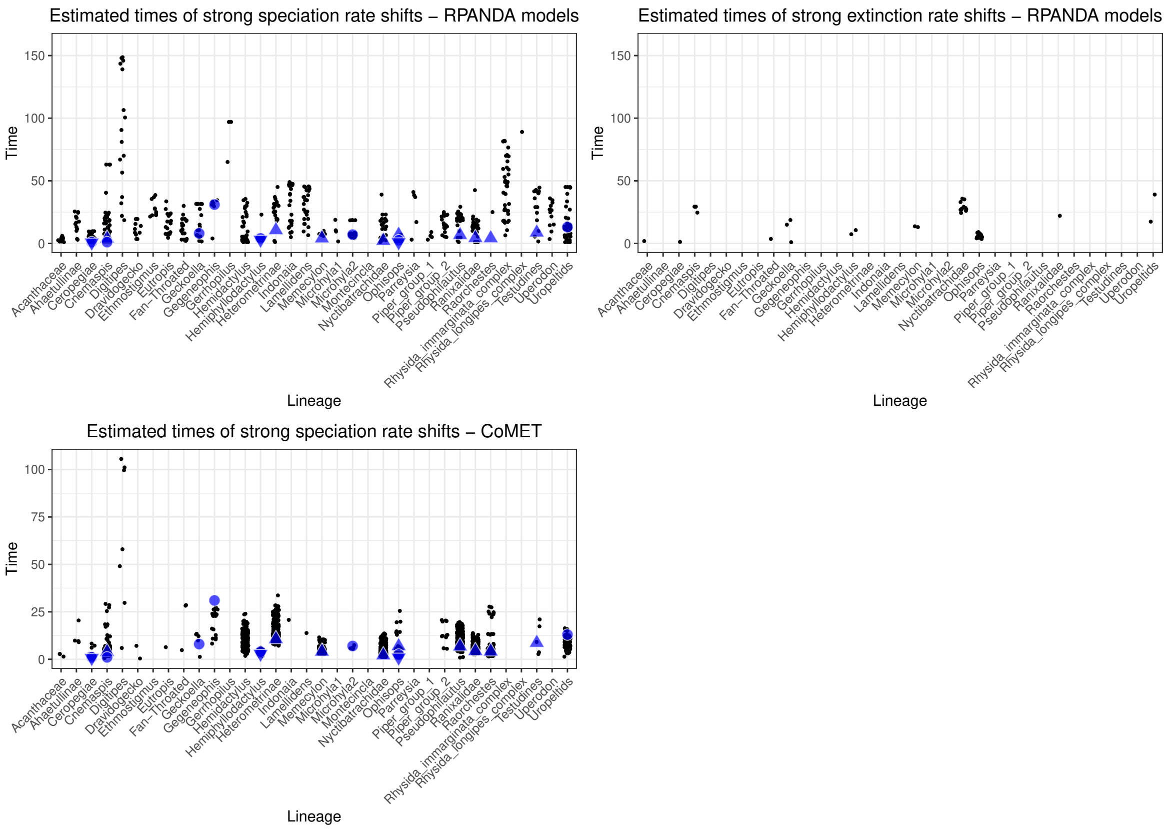

**(d)** Times of episodic rate shifts in simulated trees estimated by the SES model (in RPANDA) and CoMET.

**Figure S7: Credibility in estimates through simulations**. Credibility of the best-fitting models was inferred through simulating 100 trees with the corresponding parameter estimates, conditioning on the crown ages of the lineages. Proportion of correctness in scenario estimations were assessed by analysing the diversification dynamics of the simulated trees using **(a)** **(page 130)** the birth-death models in RPANDA and by performing the **(b)** **(page 131)** CoMET analysis. Stacked bar plots in **(a)** and **(b)** indicate the proportions of correctly (black) and incorrectly (white) inferred diversification scenarios. Scatter plots indicate no apparent trend in the proportion of correctly inferred scenarios with the number of tips. **(c)** **(page 132)** Boxplots indicates the distributions of errors in estimating the empirical parameters - $\lambda_0$, $\alpha$, $\mu_0$ and $\beta$ through simulations. **(d)** **(page 133)** Dots indicate estimated times of significant diversification rate shifts in the simulated trees estimated using the SES model and CoMET analysis and the triangles and circles indicate estimated times of episodic rate shifts in empirical trees estimated using CoMET and the SES models, respectively.

### References

[1] Hélène Morlon et al. "RPANDA: an R package for macroevolutionary analyses on phylogenetic trees". In: *Methods in Ecology and Evolution* 7.5 (2016), pp. 589–597.

[2] Sebastian Höhna, Michael R. May, and Brian R. Moore. "TESS: an R package for efficiently simulating phylogenetic trees and performing Bayesian inference of lineage diversification rates". en. In: *Bioinformatics* 32.5 (Mar. 2016), pp. 789–791. ISSN: 1367-4811, 1367-4803. DOI: 10.1093/bioinformatics/btv651. URL: https://academic.oup.com/bioinformatics/article/32/5/789/1744433 (visited on 06/20/2023).

[3] Odile Maliet and Hélène Morlon. "Fast and Accurate Estimation of Species-Specific Diversification Rates Using Data Augmentation". en. In: *Systematic Biology* 71.2 (Feb. 2022). Ed. by James Rosindell, pp. 353–366. ISSN: 1063-5157, 1076-836X. DOI: 10.1093/sysbio/syab055. URL: https://academic.oup.com/sysbio/article/71/2/353/6316269 (visited on 07/20/2024).

[4] R Core Team. *R: A Language and Environment for Statistical Computing.* R Foundation for Statistical Computing. Vienna, Austria, 2024. URL: https://www.R-project.org/.

[5] Alexis Dinno. "dunn. test: Dunn's test of multiple comparisons using rank sums". In: *R package version* 1.5 (2017), p. 1.

[6] Michael L. Collyer and Dean C. Adams. "RRPP: An r package for fitting linear models to high-dimensional data using residual randomization". en. In: *Methods in Ecology and Evolution* 9.7 (2018). _eprint: https://onlinelibrary.wiley.com/doi/pdf/10.1111/2041-210X.13029, pp. 1772–1779. ISSN: 2041-210X. DOI: 10.1111/2041-210X.13029. URL: https://onlinelibrary.wiley.com/doi/abs/10.1111/2041-210X.13029 (visited on 12/22/2024).

[7] Sudhir Kumar et al. "TimeTree 5: an expanded resource for species divergence times". In: *Molecular biology and evolution* 39.8 (2022), msac174.

[8] Emmanuel Paradis, Julien Claude, and Korbinian Strimmer. "APE: Analyses of Phylogenetics and Evolution in R language". In: *Bioinformatics* 20.2 (Jan. 2004), pp. 289–290. ISSN: 1367-4803. DOI: 10.1093/bioinformatics/btg412. URL: https://doi.org/10.1093/bioinformatics/btg412 (visited on 09/05/2023).

[9] Mark Pagel. "Inferring the historical patterns of biological evolution". In: *Nature* 401.6756 (1999), pp. 877–884.

[10] Liam J. Revell. "phytools 2.0: an updated R ecosystem for phylogenetic comparative methods (and other things)." In: *PeerJ* 12 (2024), e16505. DOI: 10.7717/peerj.16505.

[11] David Orme et al. *caper: Comparative Analyses of Phylogenetics and Evolution in R.* R package version 1.0.3. 2023. URL: https://CRAN.R-project.org/package=caper.

[12] Indrajeet Patil. "Visualizations with statistical details: The 'ggstatsplot' approach". en. In: *Journal of Open Source Software* 6.61 (May 2021), p. 3167. ISSN: 2475-9066. DOI: 10.21105/joss.03167. URL: https://joss.theoj.org/papers/10.21105/joss.03167 (visited on 12/22/2024).

[13] Hadley Wickham. "ggplot2". en. In: *WIREs Computational Statistics* 3.2 (2011). _eprint: https://onlinelibrary.wiley.com/doi/pdf/10.1002/wics.147, pp. 180–185. ISSN: 1939-0068. DOI: 10.1002/wics.147. URL: https://onlinelibrary.wiley.com/doi/abs/10.1002/wics.147 (visited on 09/05/2023).

[14] Hélène Morlon, Matthew D. Potts, and Joshua B. Plotkin. "Inferring the Dynamics of Diversification: A Coalescent Approach". en. In: *PLoS Biology* 8.9 (Sept. 2010). Ed. by Paul H. Harvey, e1000493. ISSN: 1545-7885. DOI: 10.1371/journal.pbio.1000493. URL: https://dx.plos.org/10.1371/journal.pbio.1000493 (visited on 06/20/2023).

[15] Fabien L. Condamine, Jonathan Rolland, and Hélène Morlon. "Assessing the causes of diversification slowdowns: temperature-dependent and diversity-dependent models receive equivalent support". en. In: *Ecology Letters* 22.11 (Nov. 2019). Ed. by Rampal Etienne, pp. 1900–1912. ISSN: 1461-023X, 1461-0248. DOI: 10.1111/ele.13382. URL: https://onlinelibrary.wiley.com/doi/10.1111/ele.13382 (visited on 06/20/2023).

[16] Hélène Morlon, Jonathan Rolland, and Fabien L. Condamine. "Response to technical comment 'A cautionary note for users of linear diversification dependencies'". en. In: *Ecology Letters* 23.7 (July 2020). Ed. by Rampal Etienne, pp. 1172–1174. ISSN: 1461-023X, 1461-0248. DOI: 10.1111/ele.13513. URL: https://onlinelibrary.wiley.com/doi/10.1111/ele.13513 (visited on 06/20/2023).

[17] Roy C. Spencer. "Properties of the Witch of Agnesi—Application to Fitting the Shapes of Spectral Lines". en. In: *Journal of the Optical Society of America* 30.9 (Sept. 1940), pp. 415–419. ISSN: 0030-3941. DOI: 10.1364/JOSA.30.000415. URL: https://opg.optica.org/abstract.cfm?URI=josa-30-9-415 (visited on 08/21/2023).

[18] Kenneth P Burnham and David R Anderson. "Multimodel inference: understanding AIC and BIC in model selection". In: *Sociological methods & research* 33.2 (2004), pp. 261–304.

[19] Sebastian Höhna. "The time-dependent reconstructed evolutionary process with a key-role for mass-extinction events". en. In: *Journal of Theoretical Biology* 380 (Sept. 2015), pp. 321–331. ISSN: 00225193. DOI: 10.1016/j.jtbi.2015.06.005. URL: https://linkinghub.elsevier.com/retrieve/pii/S0022519315002842 (visited on 06/20/2023).

[20] Michael R. May, Sebastian Höhna, and Brian R. Moore. "A Bayesian approach for detecting the impact of mass-extinction events on molecular phylogenies when rates of lineage diversification may vary". en. In: *Methods in Ecology and Evolution* 7.8 (2016), pp. 947–959. ISSN: 2041-210X. DOI: 10.1111/2041-210X.12563. URL: https://onlinelibrary.wiley.com/doi/abs/10.1111/2041-210X.12563 (visited on 11/12/2023).

[21] Harold Jeffreys. *The Theory of Probability*. en. Google-Books-ID: vh9Act9rtzQC. OUP Oxford, Aug. 1998. ISBN: 978-0-19-158967-6.

[22] Jacob Cohen. *Statistical power analysis for the behavioral sciences*. routledge, 2013.

[23] Geoff Cumming. *Understanding the new statistics: Effect sizes, confidence intervals, and meta-analysis*. Routledge, 2013.

[24] Andrea S Meseguer et al. “Diversification dynamics in the Neotropics through time, clades, and biogeographic regions”. en. In: *eLife* 11 (Oct. 2022), e74503. ISSN: 2050-084X. DOI: 10.7554/eLife.74503. URL: https://elifesciences.org/articles/74503 (visited on 06/20/2023).

[25] Sebastian Höhna, Bjørn T. Kopperud, and Andrew F. Magee. “CRABS: Congruent rate analyses in birth–death scenarios”. en. In: *Methods in Ecology and Evolution* 13.12 (Dec. 2022), pp. 2709–2718. ISSN: 2041-210X, 2041-210X. DOI: 10.1111/2041-210X.13997. URL: https://besjournals.onlinelibrary.wiley.com/doi/10.1111/2041-210X.13997 (visited on 07/20/2024).

[26] Irnawati Irnawati et al. “The use of software packages of R factoextra and FactoMineR and their application in principal component analysis for authentication of oils”. en. In: *Indonesian Journal of Chemometrics and Pharmaceutical Analysis* (Feb. 2021), pp. 1–10. ISSN: 2723-4630. DOI: 10.22146/ijcpa.482. URL: https://journal.ugm.ac.id/v3/IJCPA/article/view/482 (visited on 09/05/2023).

[27] Jérôme Pagès. “Analyse factorielle de donnees mixtes: principe et exemple d’application”. In: *Revue de statistique appliquée* 52.4 (2004), pp. 93–111.
